# PACAP induces light aversion in mice by an inheritable mechanism independent of CGRP

**DOI:** 10.1101/2020.08.20.255968

**Authors:** Adisa Kuburas, Bianca N. Mason, Benjamin Hing, Alyssa S. Reis, Levi P. Sowers, Cristina Moldovan Loomis, Leon F. Garcia-Martinez, Andrew F. Russo

## Abstract

The neuropeptides CGRP and PACAP have emerged as mediators of migraine, yet the potential overlap of their mechanisms remains unknown. Infusion of PACAP, like CGRP, can cause migraine in people, and both peptides share similar vasodilatory and nociceptive functions. In this study, we have used light aversion in mice as a surrogate for migraine-like photophobia to compare CGRP and PACAP and ask whether CGRP or PACAP actions were dependent on each other. Similar to CGRP, PACAP induced light aversion in outbred CD-1 mice. The light aversion was accompanied by increased resting in the dark, but not anxiety in a light-independent open field assay. Unexpectedly, about a third of the CD-1 mice did not respond to PACAP, which was not seen with CGRP. The responder and nonresponder phenotypes were stable, inheritable, and not sex-linked, although there was generally a trend for greater responses among male mice. RNA-seq analysis of trigeminal ganglia yielded hieriechial clustering of responder and nonresponder mice and revealed a number of candidate genes, including greater expression of pituitary hormones and receptors in a subset of responder mice. Importantly, an anti-PACAP monoclonal antibody could block PACAP-induced light aversion but not CGRP-induced light aversion. Conversely, an anti-CGRP antibody could not block PACAP-induced light aversion. Thus, we propose that CGRP and PACAP act by independent convergent pathways that cause a migraine-like symptom in mice.

**Significance:** The relationship between the neuropeptides CGRP and PACAP in migraine is relevant given that both peptides can induce migraine in people, yet to date only drugs that target CGRP are available. Using an outbred strain of mice, we were able to show that most, but not all, mice respond to PACAP in a preclinical photophobia assay. Our finding that CGRP and PACAP monoclonal antibodies do not cross-inhibit the other peptide indicates that CGRP and PACAP actions are independent and suggests that PACAP-targeted drugs may be effective in patients who do not respond to CGRP-based therapeutics.

## Introduction

Migraine is a sensory disorder that is more than just a severe headache. It is one of the most incapacitating neurological disorders in the world (Collaborators, 2018). One diagnostic criterion for migraine is photophobia, an often debilitating response to normal levels of light (Digre and Brennan, 2012). Despite the high prevalence of migraine and recent advances in treatments, the underlying mechanisms have yet to be fully defined. However, it is now accepted that migraine involves the neuropeptide calcitonin gene-related peptide (CGRP) (Russo, 2015; Ashina et al., 2019; Edvinsson, 2019). CGRP is upregulated during migraine attacks (Goadsby and Edvinsson, 1993; Ramon et al., 2017), infusion of CGRP can induce migraine in about 70% of migraineurs (Lassen et al., 2002), and antibodies that block CGRP or its canonical receptor can effectively prevent migraine in about 50% of patients (Scuteri et al., 2019; Tringali and Navarra, 2019). While key, it is clear that CGRP is not the only player and other sensory peptides have been suggested as candidates (Russo, 2017). In particular, the neuropeptide pituitary adenylate cyclase-activating polypeptide (PACAP) has been linked to migraine pathogenesis (Tuka et al., 2013). Notably, PACAP levels in plasma are increased during migraine attacks and infusion of either the PACAP-38 or PACAP-27 isoforms causes migraine in people (Tuka et al., 2013; Ghanizada et al., 2020).

PACAP shares many functions with CGRP (Kaiser and Russo, 2013). PACAP is a multifunctional neuropeptide involved in nociception, neurogenic inflammation, and neurovascular modulation (Vaudry et al., 2000; Hashimoto et al., 2006). PACAP belongs to the vasoactive intestinal polypeptide (VIP)-secretin-glucagon superfamily of neuropeptides (Arimura, 1992). It was first identified as a 38-amino acid form (PACAP-38) (Miyata et al., 1989) and was later found to have a truncated isoform (PACAP-27) (Miyata et al., 1990). Both forms have equivalent receptor binding affinities and biological activities (Nilsson et al., 1994). PACAP-38 is the dominant form of the two PACAP peptides, representing approximately 90% of the PACAP peptide found in circulation (Arimura et al., 1991). For this reason, we primarily focused on PACAP-38. The actions of PACAP are mediated through activation of a family of G protein-coupled receptors: VIP-PACAP1 (VPAC1), VPAC2, PACAP1 (PAC1). PACAP and VIP have equal affinity for VPAC1 and VPAC2, whereas PAC1 is preferentially activated by PACAP (Harmar et al., 2012).

Like CGRP, PACAP and its receptors are found in trigeminal ganglia neurons, but PACAP is found in considerably fewer neurons than CGRP (Frederiksen et al., 2018). Instead, the predominant sites of PACAP and its receptor are in the sphenopalatine ganglia (Uddman et al., 1999; Steinberg et al., 2016), an extracranial parasympathetic ganglion (Khan et al., 2014). Stimulation of sphenopalatine ganglia is likely to contribute to autonomic symptoms of migraine since it can increase cerebral blood flow, intracranial and extracranial vasodilation, and dural plasma protein extravasation (Schoenen, 2015). Interestingly, PACAP can also induce release of CGRP from trigeminal neurons (Jansen-Olesen et al., 2014), which suggests the possibility of cross-talk between the sphenopalatine and trigeminal systems.

To better understand the relationship of PACAP and CGRP in migraine, we have used light aversive behavior of mice as a surrogate for photophobia (Russo et al., 2009; Kaiser et al., 2012; Mason et al., 2017). In this study, we have demonstrated that peripheral administration of PACAP causes light aversion in outbred CD-1 mice similar to peripheral administration of CGRP (Mason et al., 2017). However, unlike CGRP, PACAP induces light aversion in only a sub-population of mice and their offspring. The light aversion in the responder population of mice that received PACAP could be attenuated with a monoclonal anti-PACAP antibody, but not by anti-CGRP antibody. Likewise, light aversion elicited by peripheral CGRP could not be attenuated by an anti-PACAP antibody. These data suggest that PACAP and CGRP can act by distinct pathways that converge downstream of the receptors to cause a migraine-like symptom.

## Materials and Methods

### Animals

CD-1 mice were obtained from Charles River (Raleigh, NC and Kingston, NY). Equivalent numbers of adult male and female mice, aged 10-20 weeks, were used in all experiments. Mice were housed in groups of 3-5 per cage, unless otherwise indicated, on a 12 h light cycle with food and water *ad libitum*. All behavioral experiments were performed between 7:00 A.M. and 4:00 P.M. For all experiments, investigators were blinded. Animal procedures were approved by the University of Iowa Animal Care and Use Committee and performed in accordance with the standards set by the National Institutes of Health.

### Intraperitoneal drug and antibody administration

Drugs were diluted with Dulbecco PBS (Hyclone), which was used as the vehicle in all experiments. The amounts injected were as follows: 0.1 mg/kg rat α-CGRP (Sigma-Aldrich), 0.3 mg/kg PACAP-38 (Bachem), 0.2 mg/kg PACAP-27 (Bachem), 30 mg/kg anti-PACAP monoclonal antibody, 30 mg/kg anti-CGRP monoclonal antibody (ALD405), and 30 mg/kg monoclonal IgG control antibody. The antibodies were provided by Alder Biopharmaceuticals. The antibody dose corresponded to 8 nmol per mouse, which is approximately 3-fold excess antibody over exogenous PACAP-38 (2.6 nmol) and 8-fold excess over CGRP (1 nmol). Antibody, PACAP and CGRP were administered at 10μl/g bodyweight with a 30 g x 0.5 in needle. All injections were performed by either A.K. or B.N.M. Animals were held gently but not anesthetized during injection. After PACAP or CGRP (i.p.) injection, mice were allowed to recover for 30 min in their home cages before testing as previously described (Mason et al., 2017). Antibodies were injected i.p. 24 h prior to peptide injections.

### Light aversion and motility assays

Light/dark boxes with infrared beam tracking were used (Med Associates). CD-1 mice were pre-exposed to the chamber once and three days later exposed to the light/dark box following treatment. In addition these mice were tested using bright light (25-27,000 lux), as previously described (Kaiser et al., 2012). Data were collected for 30 min and analyzed in sequential 5 min intervals, as well as average time spent on each side of the chamber per 5 min interval.

Motility outcomes were measured as described previously (Kaiser et al., 2012). To account for the variation in the amount of time mice spent in each zone, data were normalized to time spent in the dark and light zones.

### Open field assay

This assay was performed as previously described (Kaiser et al., 2012). Mice were placed in the center of the chamber and tested for 30 min. The periphery was defined as 4.22 cm from the border with the remaining 18.56 × 18.56 cm area as the center.

### RNA-seq library prep and sequencing

Trigeminal ganglia were dissected and flash frozen in liquid nitrogen. Tissue was stored in −80C till shipping. RNA extraction, library preparation and sequencing were performed by Genewiz (South Plainfiled, NJ). Briefly, good quality RNA (RIN ≥ 8) was extracted and used for library preparation using polyA selection. Sequencing was performed using Illumina HiSeq with 150 bp paired-end sequencing.

### Gene expression, variant identification, and statistical analyses

Gene expression analysis was done using FASTQ files processed using Trim Galore! version 0.5.0 (Krueger, 2018) to remove Illumina adapters and trimmed to a read length of 149 bp to remove bias bases at the 3’ end. For RNA-seq analysis, post-processed FASTQ files were aligned to the mouse genome (Gencode M19; GRCm38) using STAR (version 2.6.0a) program with --twopassMode Basic option and --sjdbOverhang 148 (Dobin and Gingeras, 2015) with high mapping efficiency of ~93% giving ~30 million mapped reads. Alignments were then processed using StringTie (version 1.3.5) for transcript assembly and subsequent quantification of read counts and transcript abundance (Pertea et al., 2016). Data were subsequently imported into R using tximport (version 1.10.1) (Soneson et al., 2015) gene level analysis using DESeq2 as described in the vignette (version 1.22.1) (Love et al., 2014). Data were analyzed using a negative binomial generalized linear model whereby gene level counts was the response variable and treatment group was the explanatory variable. Sex was included in the model as a covariate. Since library preparation was performed on all samples at the same time and samples were sequenced together, this eliminated the requirement for controlling for batch effects. Wald test was used to evaluate statistical difference between treatment groups. The Benjamini-Hochberg procedure was used to adjust p-values to correct for multiple testing (https://www.statisticshowto.datasciencecentral.com/benjamini-hochberg-procedure/). Pathway and gene ontology analyses were performed using ConsensusPathDB (Kamburov et al., 2012).

Variant calling and genetic association analysis was done using the GATK RNA-seq variant calling workflow as described (GATK, 2014). Briefly, STAR 2-pass mapping procedure using regenerated genome workflow was performed using default settings of STAR program with --sjdbOverhang 148 with high mapping efficiency as mentioned above. Read group was assigned followed by subsequent sorting by chromosome, marking of duplicates and indexing using Picard. Following this, tools from The Genome Analysis Toolkit (GATK; version 3.8-1-0-gf15c1c3ef) were used for subsequent processing. SplitNCigarReads was used to split reads into exon segments, hard-clip sequences that overhang into the intronic regions and reassign mapping qualities produced by STAR to be compatible with GATK workflow. This was followed by IndelRealigner for local realignment of reads around indels and BaseRecalibrator to detect and adjust systematic errors in base quality scores. Joint genotyping as previously mentioned (Brouard et al., 2019) was performed using HaplotypeCaller with options -ERC GVCF, -dontUseSoftClippedBases and -stand_call_conf 20.0 to produce one gVCF file per sample. GenotypeGVCFs was then used for multi-sample aggregation to produce a combined variant calling file (McKenna et al., 2010). Given the small sample size, we used a conservative approach whereby only biallelic sites where all samples met previously established thresholds, (i.e. DP >= 10, GQ >= 20, FS < 30 and QD >= 2) were retained for subsequent analysis (GATK, 2014; Song et al., 2016). As such, 62,854 sites remained for the study. None of these sites showed significant deviation from Hardy-Weinberg equilibrium after multiple correction by Bonferroni method. Genome-wide association analysis was performed using factored spectrally transformed linear mixed model (FaST-LMM) to estimate and account for genetic relatedness (Lippert et al., 2011). FaST-LMM removed 15,608 sites that were invariant leaving 47,246 sites for the analysis. Correction for multiple testing was performed using Bonferroni. Single nucleotide polymorphic sites (SNPs) were annotated using SnpEff which also predicts the impact of those SNPs (Cingolani et al., 2012).

### Experimental design and statistical analyses

Each experiment was conducted separately with a new cohort of mice, except that mice used in the open field assay were previously tested for light aversion. For light aversion experiments, data are expressed as time spent in the light as a function of time in the 30 min assay with the average for each cohort per 5 min interval (line graph) and as the average time per 5 min over the entire 30 min assay for each individual mouse (scatter plot). For open field experiments, data are expressed as the percentage of time the mouse spent in the center zone for the entire 30 min duration of the assay. For resting time, rearing (vertical beam breaks), and transitions, data are represented in the same way as the light aversion assay, except normalized to time spent in light and dark zones for resting and rearing. Individual numbers of animals used for each experiment can be found in the legend of each figure. For these studies, 881 mice were tested, 12 of which were excluded for reasons described below. Hence, data are from a total of 869 mice (433 female, 436 male).

A two-way, repeated measures ANOVA (factors: treatment and observation time) was used for data plotted as a function of time. A one-way ANOVA was used to determine whether overall significant effects were observed in bar graphs with individual points. Bonferroni multiple-comparisons test was used as the post hoc analysis. Data are reported as mean ± SEM. Data were analyzed using GraphPad Prism 8, version 8.4.1 software. Significance was set at p<0.05 for all analyses. G power was used to perform a power analysis. Expected effect sizes were based on previously published light aversion data with CGRP (Mason et al., 2017). All sample sizes for light aversion were greater than or equal to the suggested sample size of 9 calculated in G power for a 0.8 desired power. Exclusions were applied to the dataset from the pre-exposure data (before treatments) for the following reasons: never leaving the light zone, an overall resting time >90%, and spending 1 standard deviation less time in the light than the average baseline of the entire cohort tested on that day. Exclusions were also applied to the dataset after treatment for the following reasons: never leaving the light zone during 30 min of testing, an overall resting time >90%, and statistical outliers according to the GraphPad Prism ROUT method (Q=1%). A total of 12 out of 881 mice post-treatment were excluded, 11 for >90% resting time and 1 as a statistical outlier. All behavioral statistics are reported in Tables 1 and 1-1. All data are available upon request.

**Table 1.**
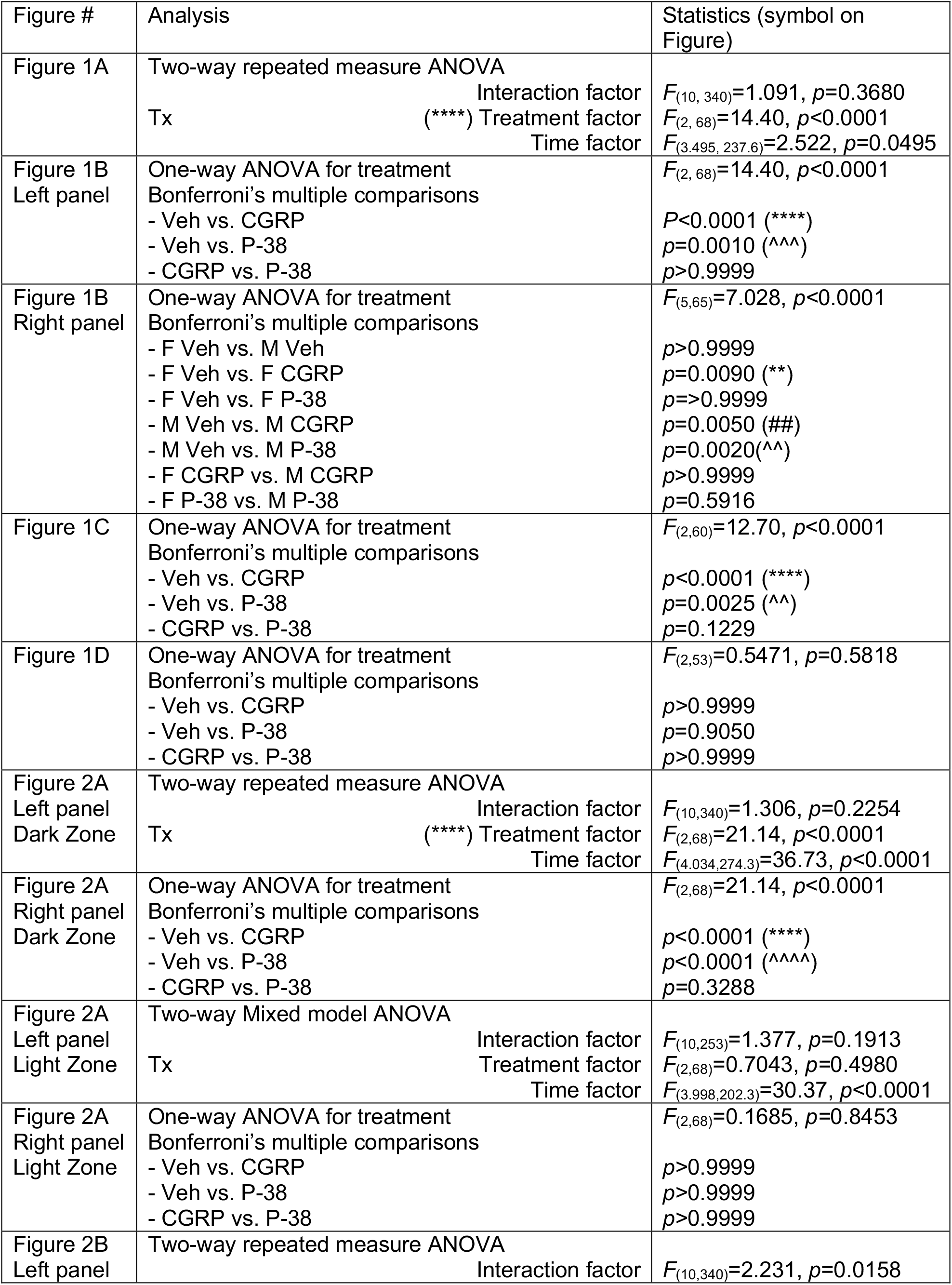
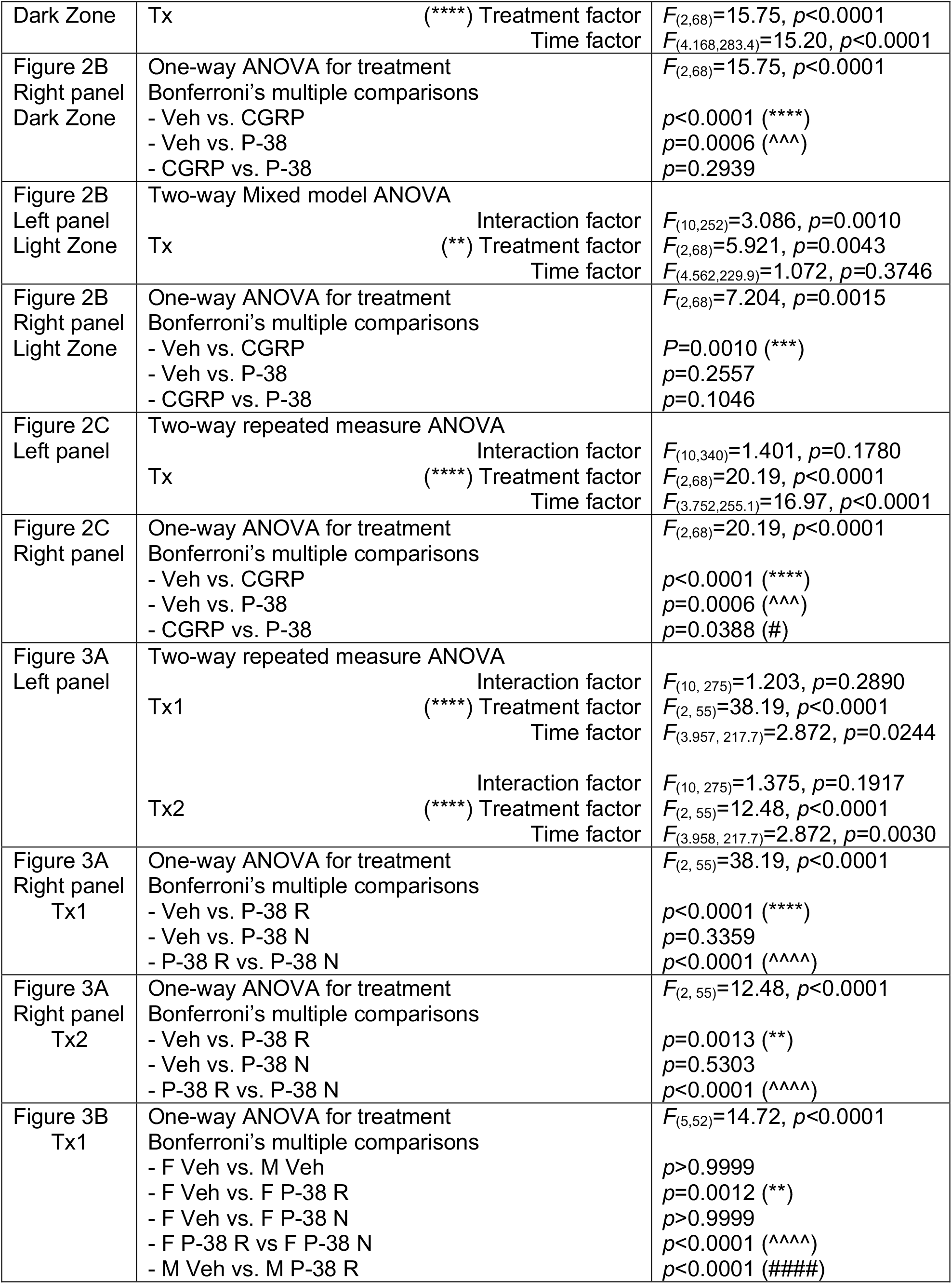
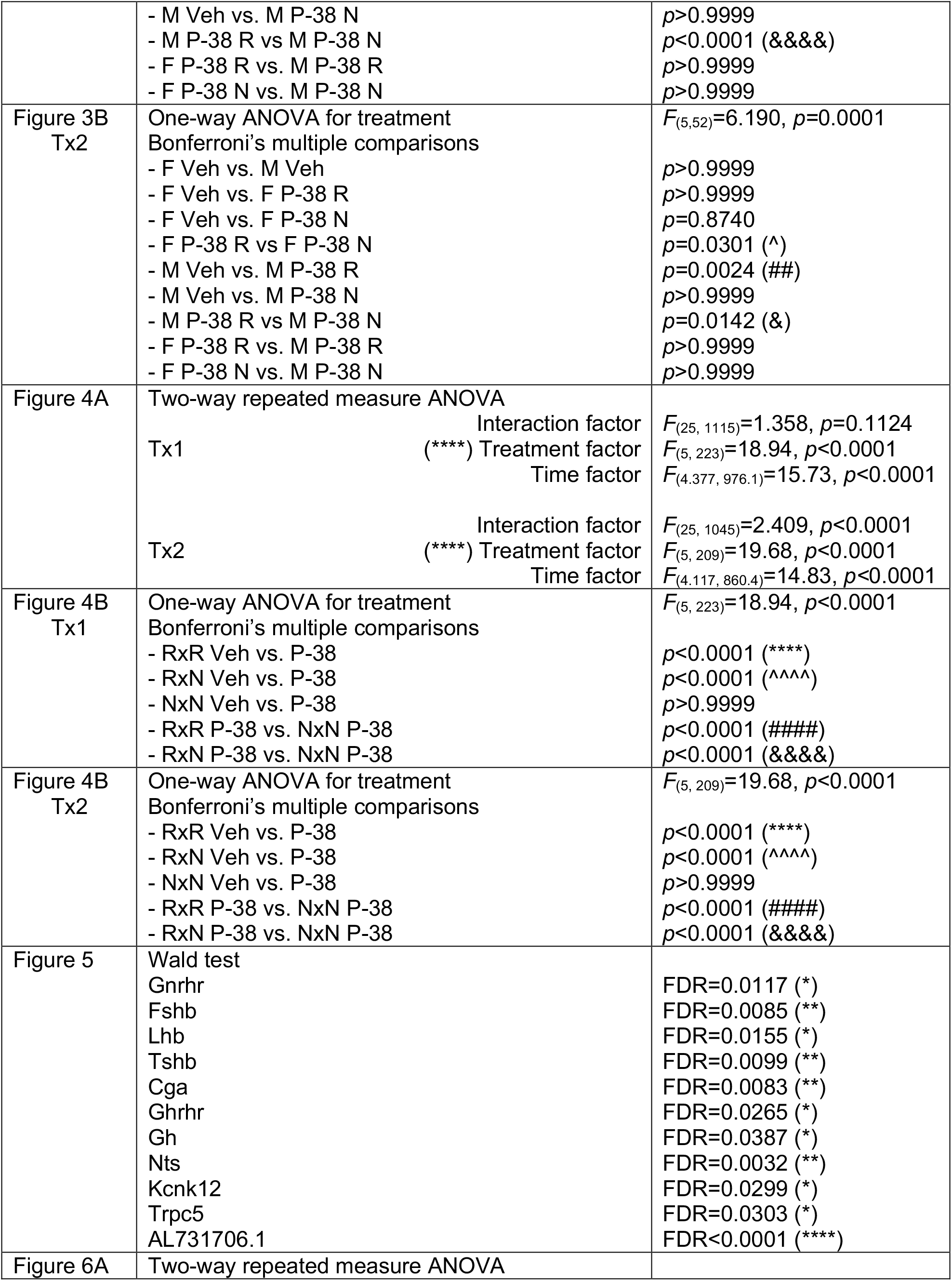
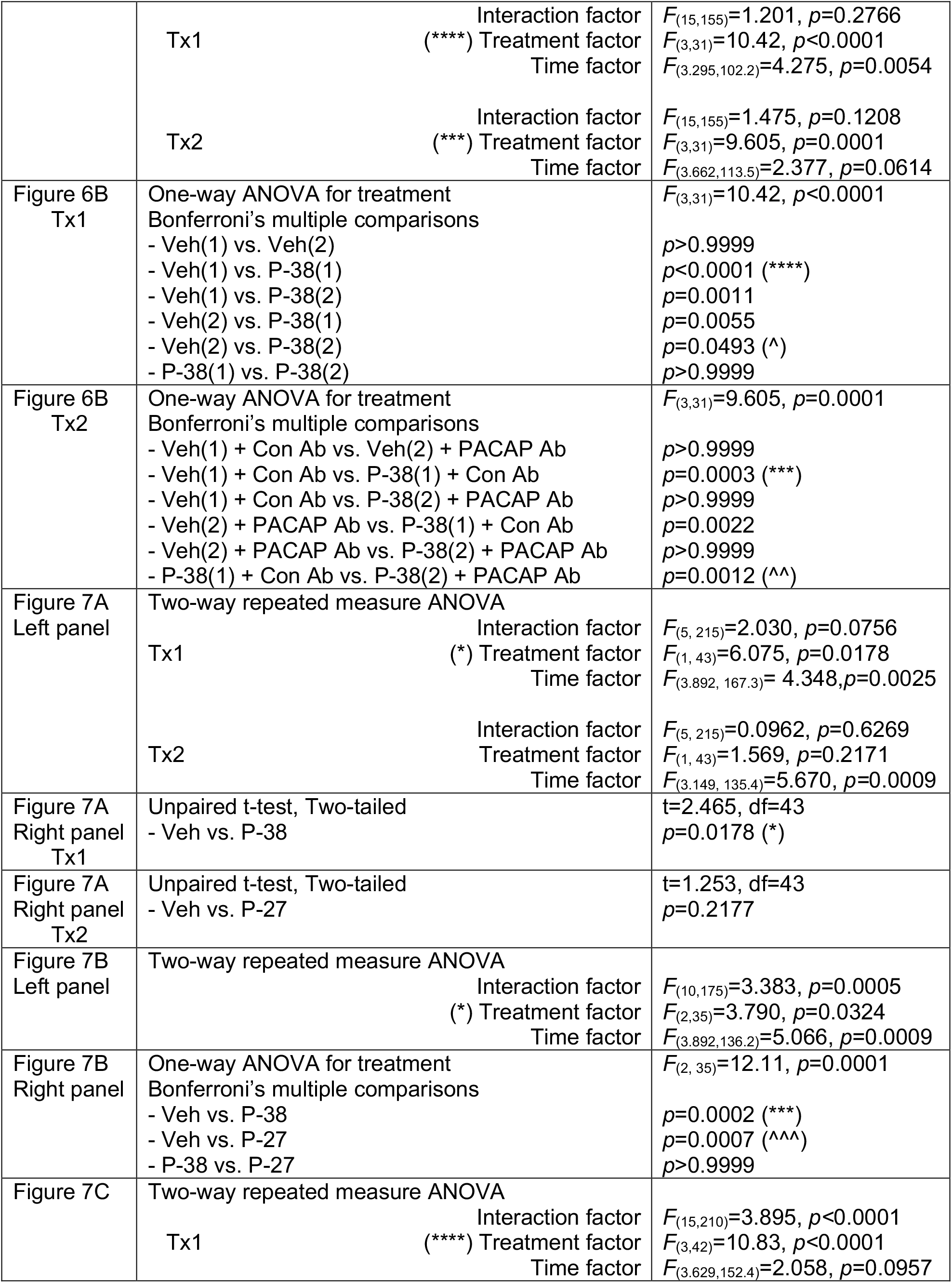
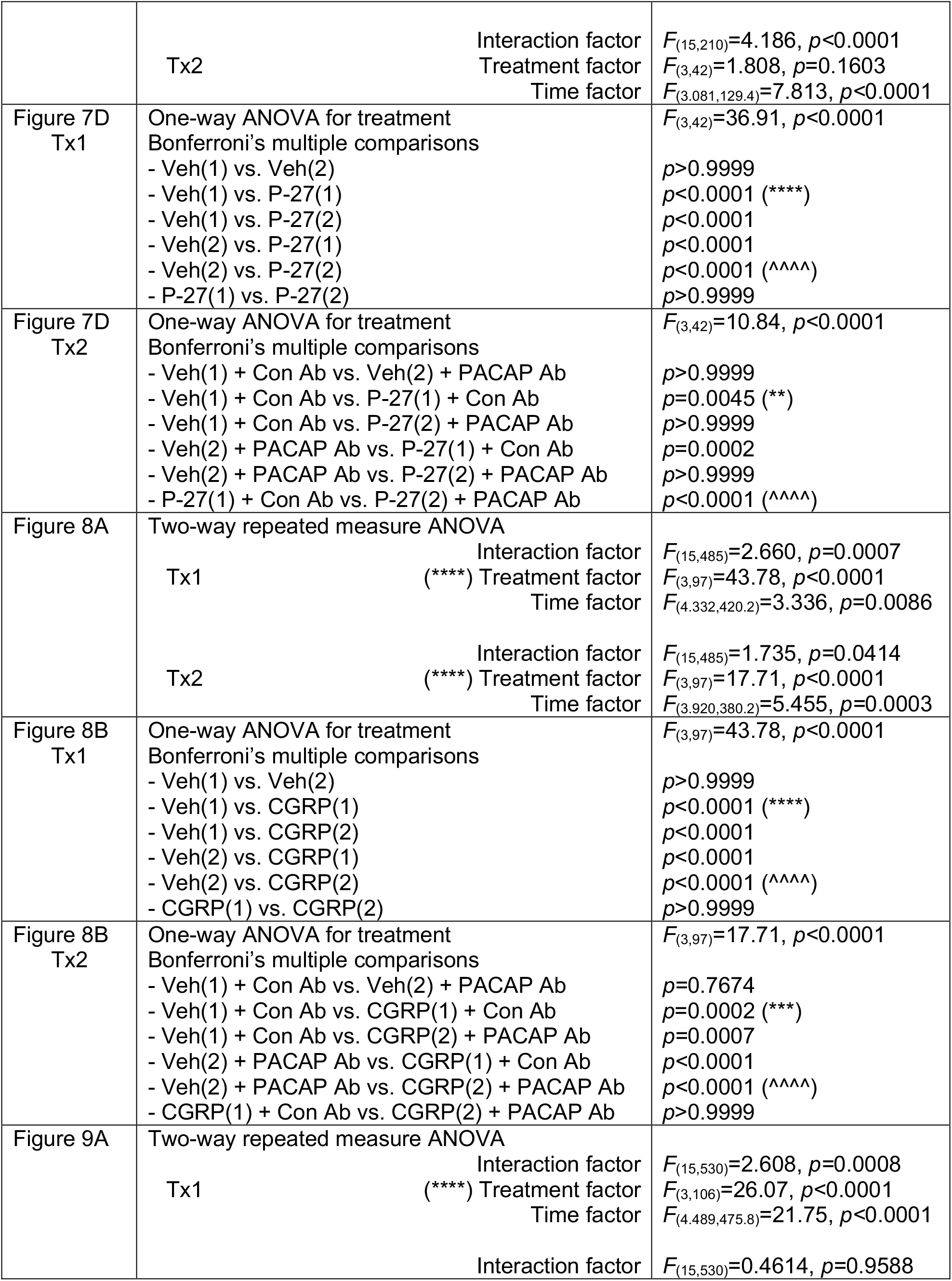
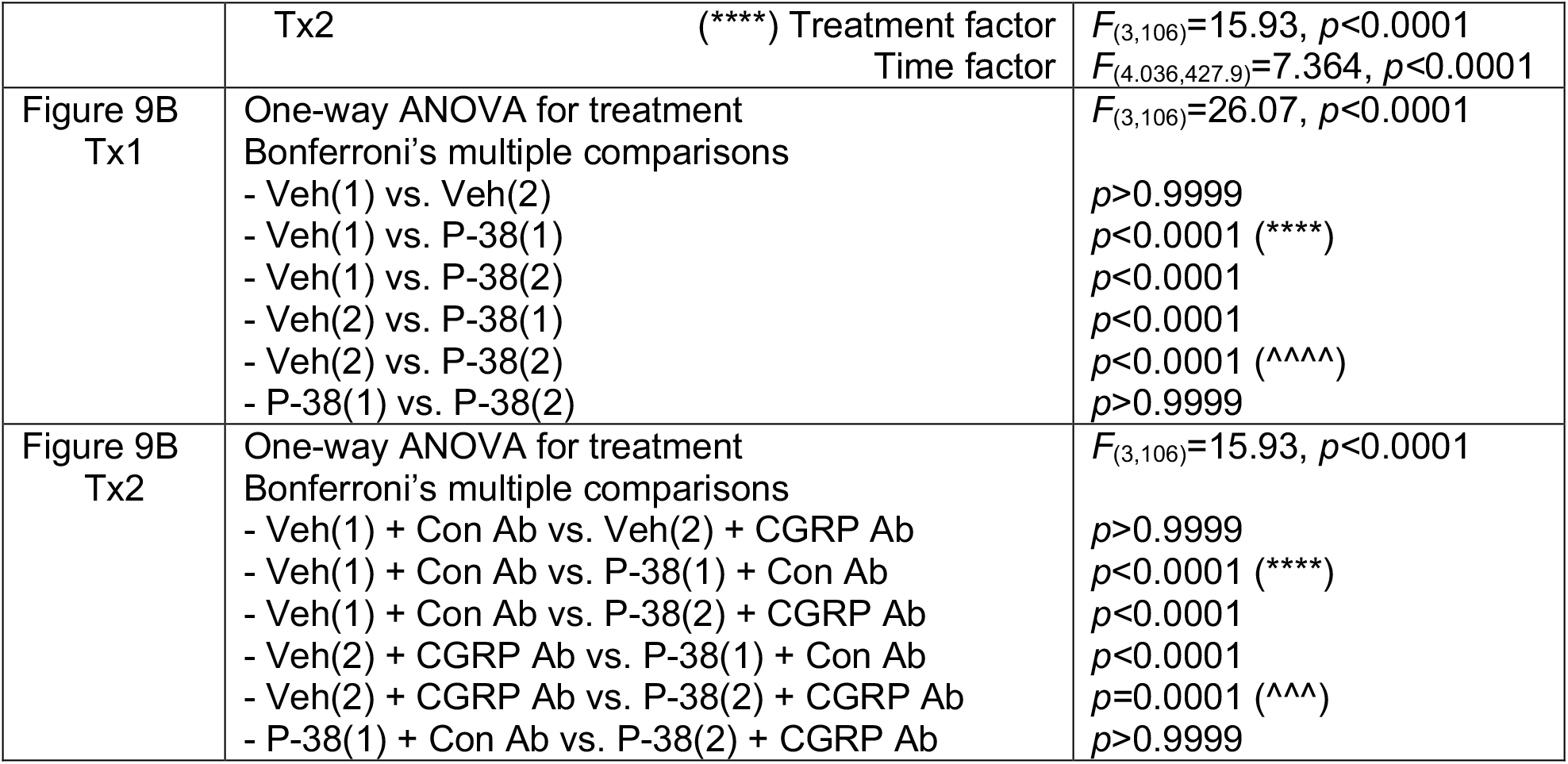
Statistical analyses. Analyses are described for each figure.

## Results

### PACAP-38 induced light aversion in CD-1 mice

We have previously shown that peripheral administration of CGRP causes light aversion in CD-1 mice (Mason et al., 2017). Here we investigated whether PACAP-38 can elicit light aversion similar to CGRP. CD-1 mice were given vehicle, PACAP-38, or CGRP in a single intraperitoneal injection and tested 30 min post injection. When we looked at light aversion as a function of time over the 30 min testing period there was a significant treatment effect (Fig. 1A). PACAP induced light aversion comparable to CGRP. Both PACAP and CGRP treated mice spent significantly less time in light compared to vehicle treated mice (Fig.1B left panel). On average, the vehicle treated mice spent 100 s in the light per 5 min interval compared with 55 s for PACAP-treated mice and 34 sec for CGRP-treated animals (Fig. 1B left panel). Since both male and female mice were tested we looked to see if there was any difference between sexes (Fig. 1B right panel). There is a trend towards males spending less time in light but it is not significantly different from females. Power analysis predicts that we would need n=100 mice of each sex to see a significant difference. Interestingly, with CGRP, although not statistically significant, we see an opposite trend as females spent less time in the light, as previously reported in Mason et al., 2017. The paradigm of two pre-exposures to the chamber to reduce the exploratory drive was originally developed with C57BL/6J mice (Kaiser et al., 2012). Our previous data suggested that CD-1 mice are more sensitive to CGRP than C57BL/6J mice (Mason et al., 2017), so we asked if only one pre-exposure was sufficient to detect PACAP and CGRP-induced light aversion in CD-1 mice. After only one pre-exposure to the chamber, mice treated with PACAP or CGRP spent significantly less time in light compared to vehicle treated mice (Fig. 1C). All subsequent studies used just a single pre-exposure.

**Figure 1.**
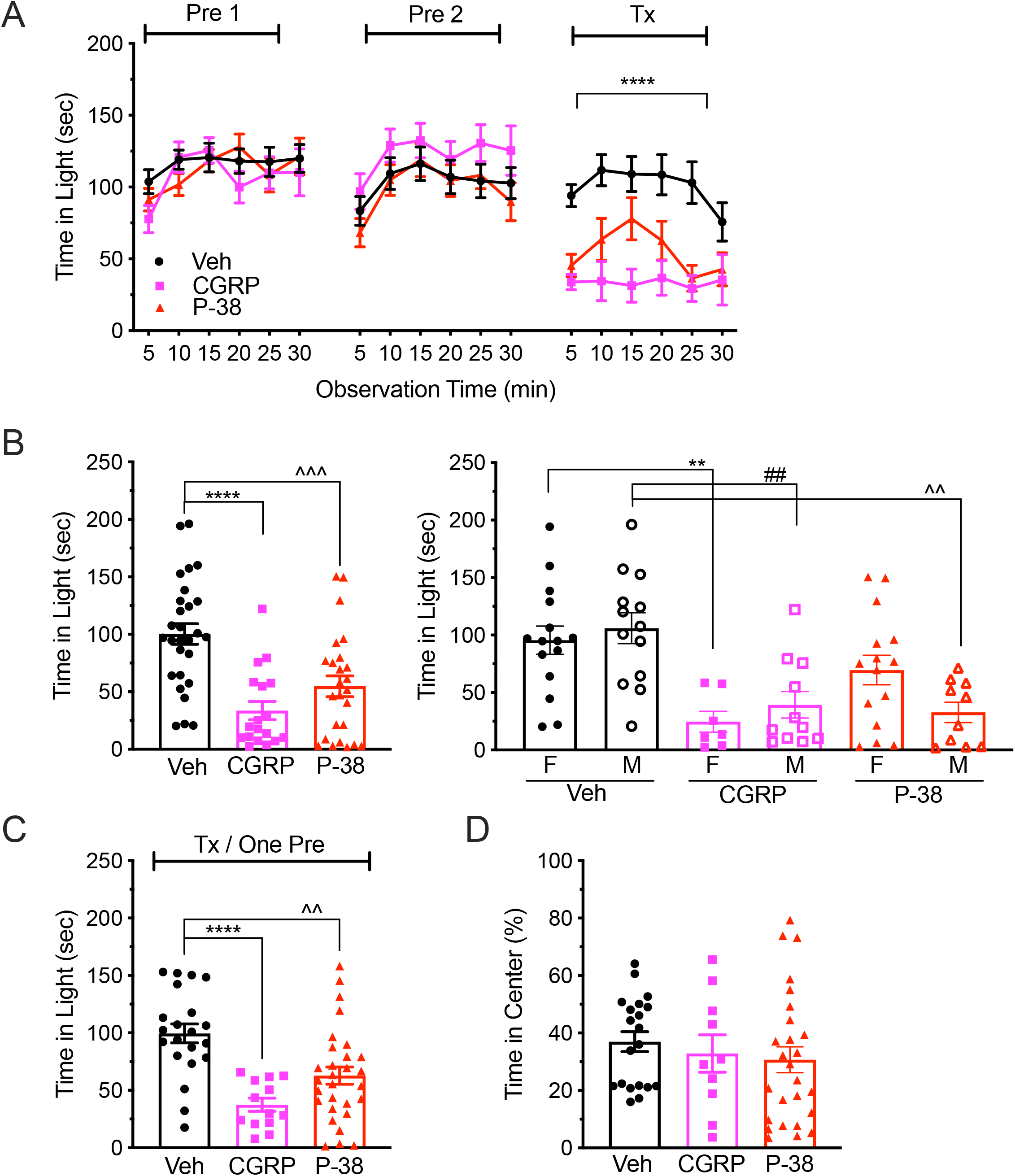
PACAP-38 induces light aversion in CD-1 mice. **A)** After two baseline pre-exposures in the light-dark box (Pre1, Pre2), mice were treated (Tx) with vehicle (Veh, n=28), PACAP-38 (P-38, 0.3 mg/kg, n=25) or CGRP (0.1 mg/kg, n=18). The mean (± SEM) time spent in the light zone every 5 min over a 30 min period is shown for each pre-exposure and after treatment. **B)** Left panel: Data from individual mice are shown with the mean (± SEM) time in light per 5 min interval calculated from the entire 30 min testing period from panel (A). Right panel: Data from left panel separated as male (M) and female (F). **C)** Effect of treatment with vehicle (n=21), PACAP-38 (0.3 mg/kg, n=29) or CGRP (0.1 mg/kg, n=13) after only one baseline pre-exposure (Tx/One Pre). Data are shown for individual mice with the mean (± SEM) time in light per 5 min interval from the entire 30 min testing period. **D)** Open field behavior of mice injected with vehicle (n=20), PACAP-38 (0.3 mg/kg, n=26) or CGRP (0.1 mg/kg, n=10). Data are shown for individual mice with the mean time (percent ± SEM) in the center zone. Statistics are described in Table 1.

To test whether the light aversion was due to an anxiety-like response, we tested mice in a light-independent open field assay. As with the light aversion assay, mice were tested 30 min after injection of PACAP-38, CGRP or vehicle. There was no significant difference in the time the mice spent in the center of the open field between vehicle and PACAP-38 or CGRP treated animals (Fig. 1D).

### PACAP-38 reduces motility in the dark zone

To see if PACAP has the same effect on motility as CGRP (Kaiser et al., 2012; Mason et al., 2017), we looked at resting time, rearing (vertical beam brakes), and transitions. Motility was measured at the same time from the same mice as during the light aversion experiment shown in Fig. 1A. PACAP-38 increased resting in the dark zone, but not in the light, similar to CGRP (Fig. 2A). Rearing behavior was also significantly decreased by PACAP-38 in the dark zone, but not the light zone (Fig. 2B). CGRP also decreased rearing, but the decrease was significant in both the light and dark zones, as previously reported for C57BL/6J mice (Mason et al., 2017). The number of transitions between the light and dark zones was significantly decreased in both PACAP-38 and CGRP treated mice compared to vehicle (Fig. 2C).

**Figure 2.**
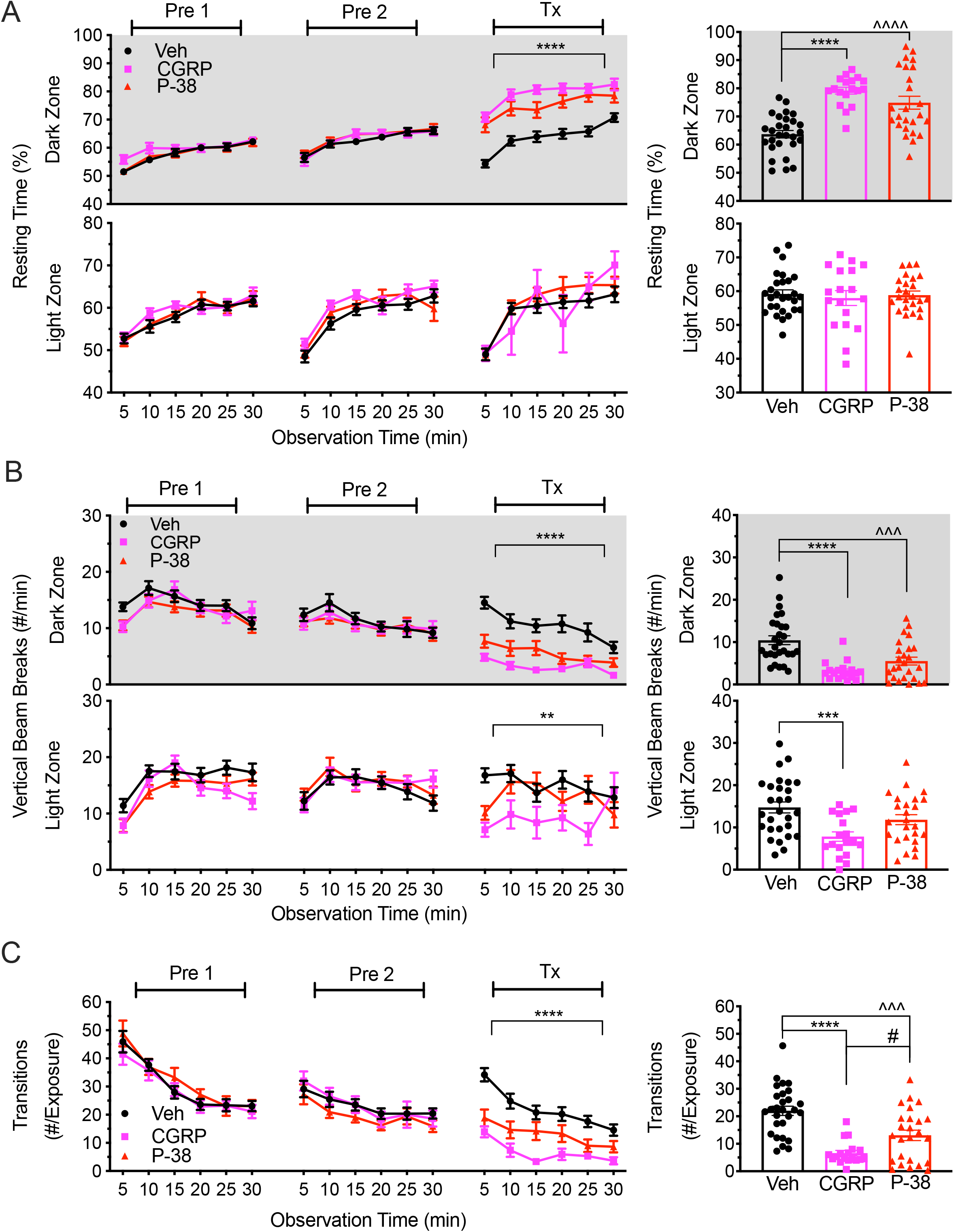
PACAP-38 reduces motility. Motility data were collected at the same time as light aversion data from the same mice shown in Figure 1A, B. After two baseline pre-exposures (Pre1, Pre2), mice were treated (Tx) with vehicle (Veh, n=28), PACAP-38 (P-38, 0.3 mg/kg, n=25) or CGRP (0.1 mg/kg, n=18). **A)** Resting time in light and dark zones. Left panel: percent (± SEM) time spent resting in each zone every 5 min over the 30 min testing period. Right panel: data for individual mice from treatment day shown as the mean percent time (± SEM) spent resting in each zone per 5 min interval. **B)** Rearing in light and dark zones. Left panel: mean (± SEM) number of vertical beam breaks in each zone every 5 min over the 30 min testing period. Right panel: data for individual mice from treatment day shown as mean (± SEM) number of vertical beam breaks in each zone per 5 min interval. **C)** Transitions between the light and dark zones. Left panel: mean (± SEM) number of transitions every 5 min over the 30 min testing period. Right panel: data for individual mice from treatment day shown as mean (± SEM) number of transitions per 5 min interval. Statistics are described in Table 1.

### Two subpopulations identified in CD-1 mice: PACAP-38 responders and nonresponders

In all experiments, we noticed that about one third of CD-1 mice treated with PACAP-38 did not appear to have a light aversive response. This was not seen with parallel treatments with CGRP or in previous CGRP studies (Mason et al., 2017). To explore the possibility that there might be two subpopulations in this outbred strain, we tested all the mice again after a second injection of PACAP (Fig. 3A). Responders were defined as mice with time in light one standard deviation below the group’s baseline average. Mice with time in light above that were considered nonresponders. The two phenotypes were equally represented in both sexes (Fig. 3B). Of the mice that were light-aversive after the first injection of PACAP, 89% were also light-aversive after the second injection of PACAP (Fig. 3C). Likewise, the mice that were not light aversive after the first injection, 85% were still not light aversive after the second injection. For comparison, using the same criteria for the vehicle treatments, 20% would have been classified as “responders after the 1^st^ vehicle treatmend and 30% after the 2^nd^ treatment.

**Figure 3.**
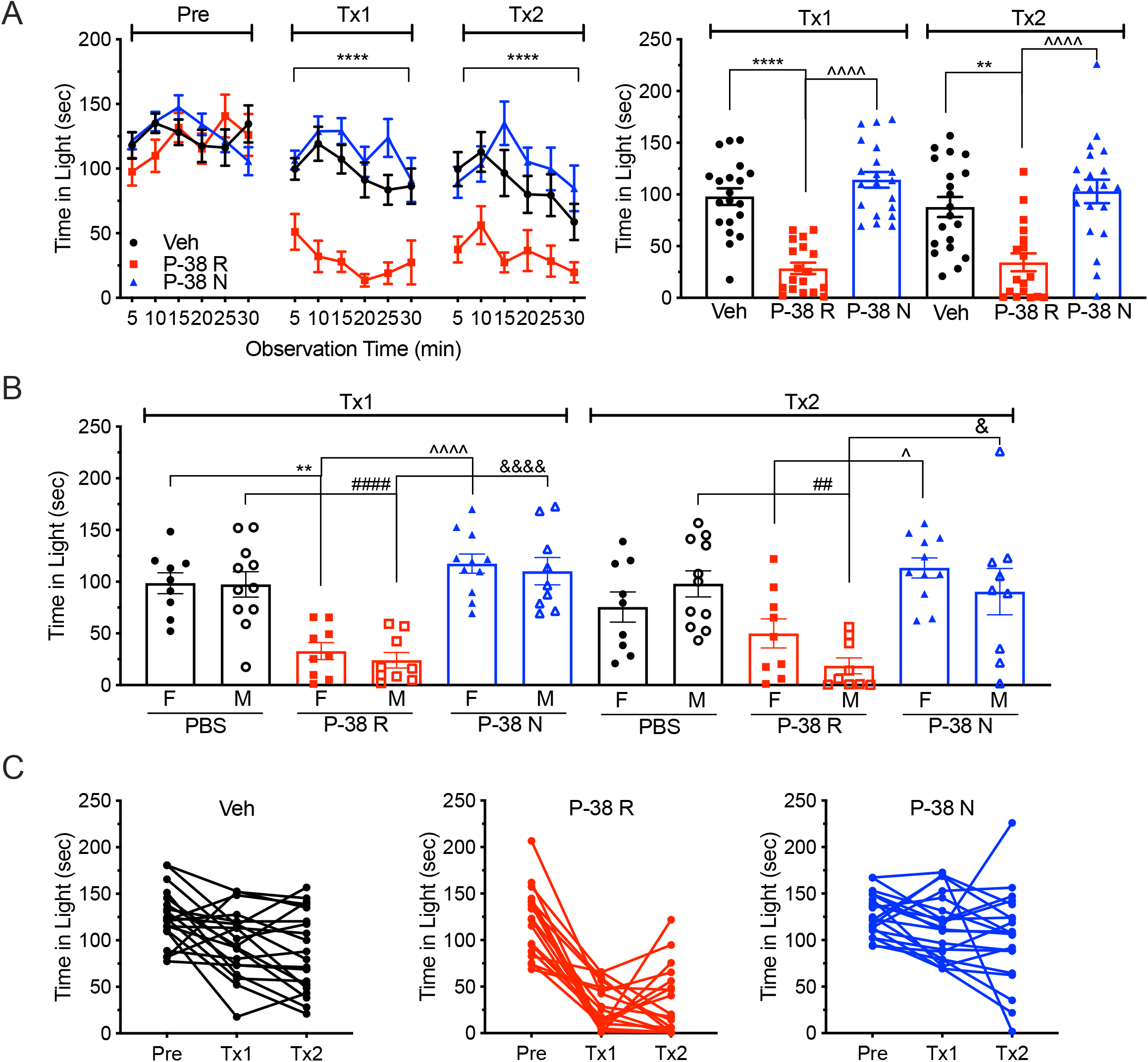
PACAP-38 responder and nonresponder subpopulations. **A)** After one baseline pre-exposure (Pre), mice were treated with either vehicle (Veh, n=20) or PACAP (P-38, 0.3 mg/kg, n=38) in the light-dark box (Tx1). After 3 days the same mice were treated and tested again (Tx2). PACAP treated mice are separated into responders (P-38 R, n=18) and nonresponders (P-38 N, n=20). Left panel: Mean ± SEM time spent in the light compartment every 5 min over a 30 min period is shown for each test day. Right panel: Data for individual mice from each treatment day shown as the mean time (± SEM) in light per 5 min interval. **B)** Data from (A) right panel separated by sex (male, M; female, F). **C)** Data from (A) right panel showing responses of individual mice to vehicle (Veh) or PACAP for the responder (P-38 R) and nonresponder (P-38 N) populations during the pre-exposure (Pre), first treatment (Tx1), and second treatment (Tx2). Statistics are described in Table 1.

In addition to light aversion, we also looked at the motility parameters of resting, rearing, and transitions in the responder and nonresponder populations. The responder population had increased resting in the dark only (Fig. 3-1A), as observed without separating the two groups (Fig. 2A). There was also decreased rearing in the dark for the responders, which was also now significantly decreased in the light (Fig. 3-1B). Finally, the responder population had decreased transitions (Fig. 3-1C). In contrast, there was no change in any of these parameters in nonresponders after PACAP-38 treatment (Fig. 3-1A-C). These properties were seen in both the first and second treatments. Thus, both light aversion and the associated decrease in motility were stable in the two populations.

### Responder and nonresponder behaviors are an inheritable trait

To further test the differences between the two subpopulations, we bred the responder and nonresponder mice to see if the trait was inheritable. The progeny (F1) were tested for PACAP responses in the light aversion assay twice to confirm their phenotypes. Crosses between responders and responders (RxR) and crosses between responders and nonresponders (RxN) yielded 88% and 73%, respectively, of the progeny responding to PACAP-38 (Fig. 4A, B). In contrast, only 24% of the progeny of crosses of nonresponders with nonresponders (NxN) responded to PACAP-38 (Fig. 4A, B) (Table 4-1). The percentage of mice that responded to PACAP-38 was about the same between males and females (RxR: 90% females and 87% males, RxN: 72% females, 74% males). Of relevance to interpretations of this experiment, 20-33% of mice treated with vehicle had low time in light that would be considered responders by our criteria (Table 4-1). We believe that this represents the inherent variability in the phenotype. In addition, PACAP reduced motility of the progeny from responder crosses (both RxR and RxN), but not in the nonresponder NxN crosses (Fig. 4-1). This was seen as both increased resting in the dark zone and decreased transitions between the light and dark zones (rearing behavior could not be measured due to a technical error). These results establish that the responder and nonresponder phenotypes are hereditary based on the high association between mating pair type and responder status (Pearson chi-square test, p-value = 1.48e-09).

**Figure 4.**
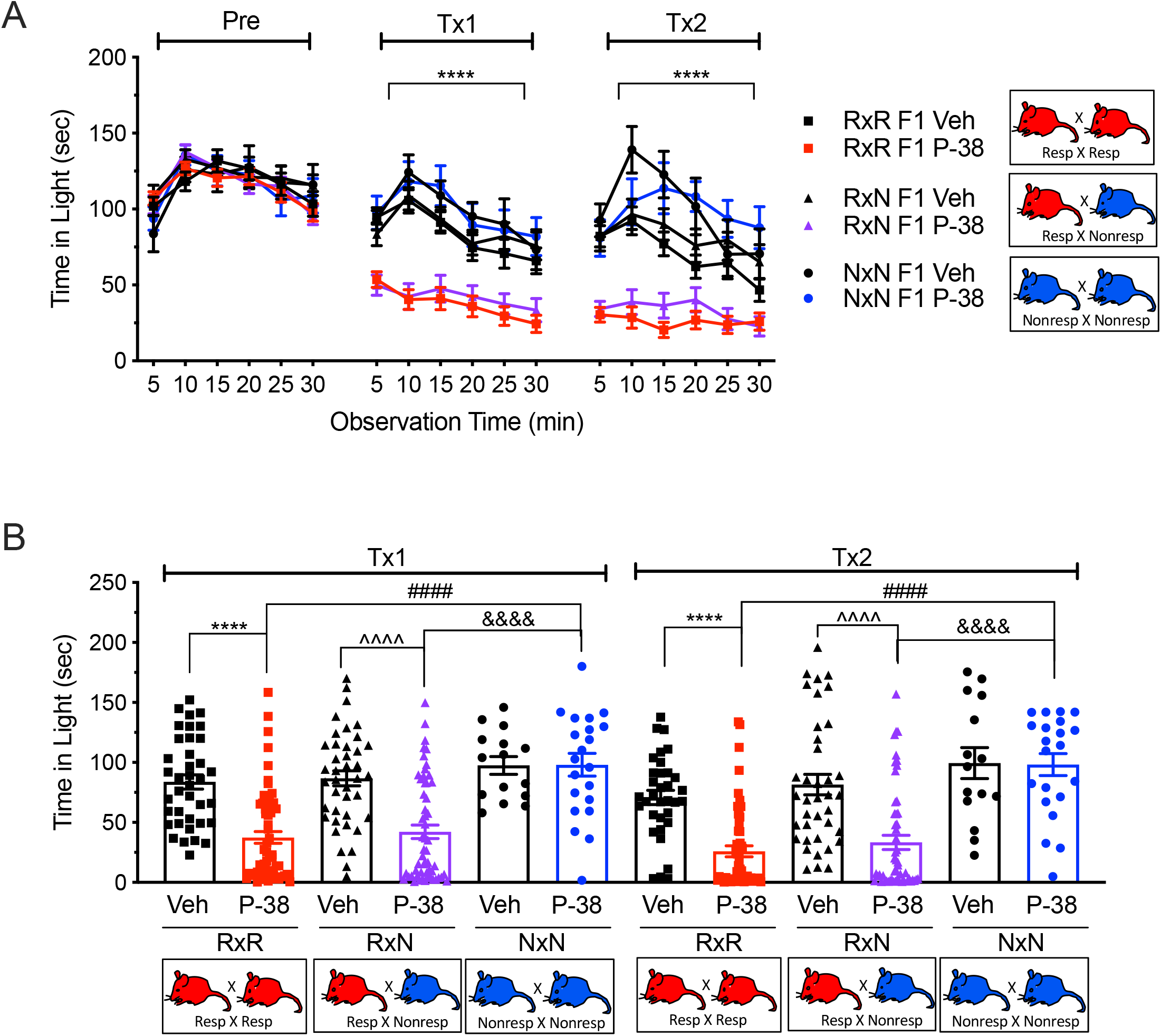
Light aversion in F1 responders and nonresponders. **A)** F1 progeny of crosses between responder and responder mice (RxR), responder and nonresponder mice (RxN), and nonresponder and nonresponder mice (NxN) were given one pre-exposure to the testing chamber (Pre), then treated (Tx1) with vehicle (Veh) (RxR, n=38; RxN, n=40; NxN, n=15), or PACAP-38 (P-38, 0.3 mg/kg) (RxR, n=59; RxN, n=56; NxN n=21). After 3 days the same mice were treated and tested again (Tx2). The mean (± SEM) time spent in the light zone every 5 min over a 30 min period is shown for each test day. **B)** Data from individual mice are shown with the mean (± SEM) time in light per 5 min interval over the entire 30 min testing period from panel (A). Statistics are described in Table 1.

The high prevalence of responders when at least one of the parental mice was a responder (both RxR and RxN crosses) suggested a dominant inheritance pattern. However, a likelihood ratio test revealed that there was a difference between the crosses (p = 0.0306), which meant that inheritance of a segregating allele with a dominant effect was not statistically supported by the data. Thus, the crosses indicate that responder status is hereditary, but apparently not by a simple dominance mechanism.

### Differences in gene expression and genotypes between responder and nonresponder mice

To evaluate potential gene expression differences between responder and nonresponder mice, we focused on trigeminal ganglia. While the best tissue to analyze is not known, we reasond that trigeminal ganglia are a good starting point since they are activated during photophobia and migraine (Moulton et al., 2009; Noseda and Burstein, 2013). RNA sequences (RNA-seq) were analyzed from 6 PACAP-38 responder and 6 nonresponder mice, with an equal number of both sexes. The mice were the F1 progeny from 2 independent crosses of responder mice and 2 independent crosses of nonresponder mice. Phenotypes of both parental and F1 progeny were confirmed by two sequential PACAP-38 treatments (0.3 mg/kg), 2 days apart, with testing at 30 min post-injection. Hierarchial clustering of the RNA-seq data in a heat map revealed that PACAP responders and nonresponders formed two distinct groups (Fig. 5A). Thus, the groups could be distinguished not only by phenotype, but also based on differential gene expression in trigeminal ganglia.

**Figure 5.**
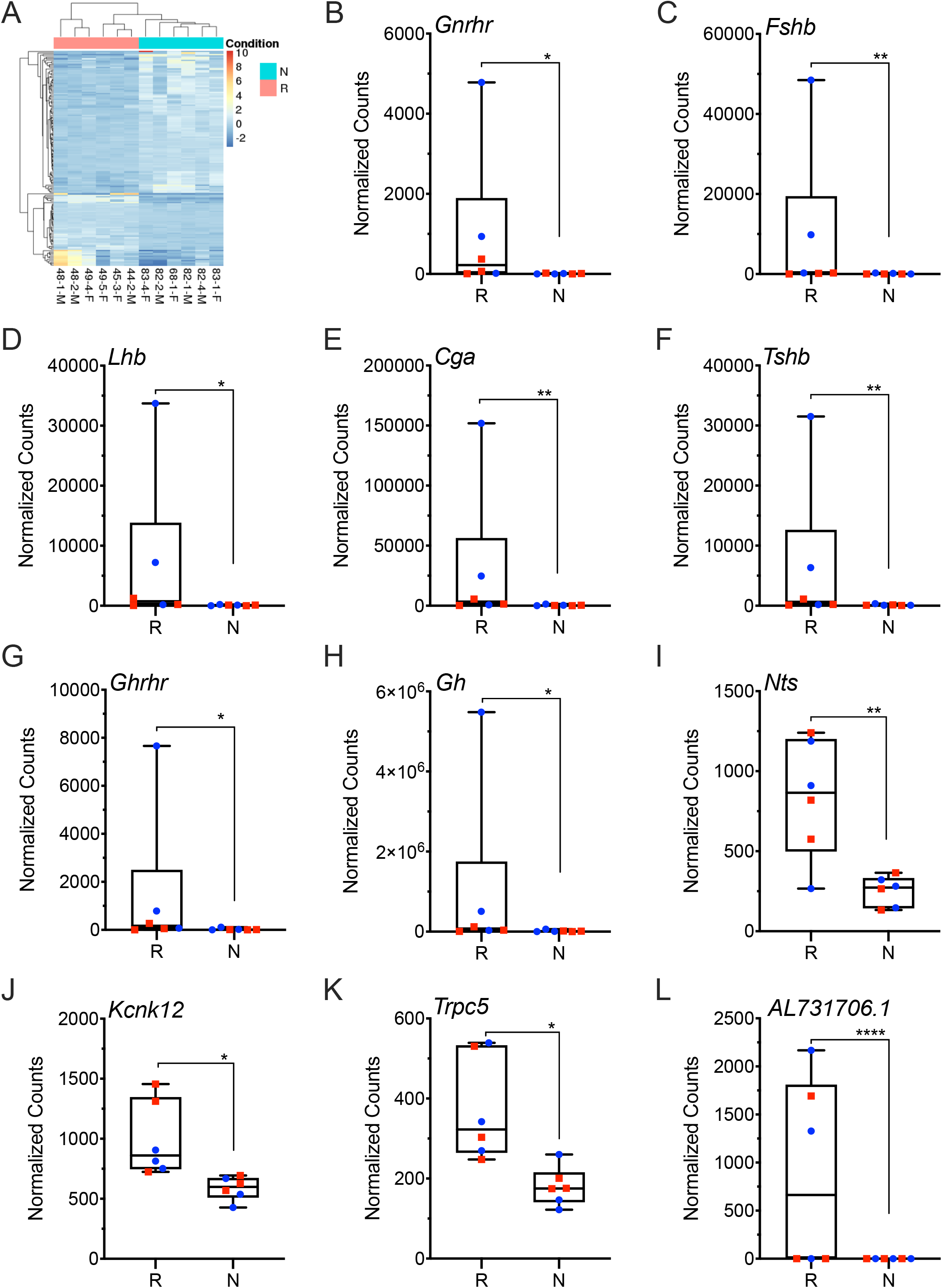
Differential gene expression between responder and nonresponder mice. Gene level counts were normalized to size factors calculated by DESeq2 to correct for library size. Normalized counts were compared between responders and nonresponders using Wald test for significance testing. Male and female mice in each cohort are indicated by blue and red symbols, respectively. (A) Heat map reflects gene expression differences between responder (R, orange) and nonresponder (N, teal) mice on the X axis from the list of differentially expressed genes in Table 5-1 that are grouped on the Y axis. Mouse number and sex, male (M) and female (F), are indicated. Mouse numbers with the same first numbers (e.g. 48) indicate mice from the same litter. (B) *Gnrhr*, (C) *Fshb*, (D) *Lhb*, (E) *Cga*, (F) *Tshb*, (G) *Ghrhr*, (H) *Gh*, (I) *Nts*, (J) *Kcnk12*, (K) *Trpc5* and (L) *AL731706.1*. Statistics are described in Table 1.

A total of 148 genes had significantly different expression between the two cohorts (Table 5-1). Genes more highly expressed in responders are designated with a positive log2 fold change (FC) value, while those expressed at a lower level in responders have a negative log2 FC value. While both male and female mice were responders and nonresponders, because there was a trend for greater responses in male mice, the RNA data were analyzed with sexes together and separately (Tables 5-2, 5-3). When both sexes were analyzed together, of the 148 differentially expressed genes, 50 were more highly expressed in responder mice. One striking finding was that responder mice have greater expression (4-6 log2 FC) of pituitary hormone and receptor RNAs. Specifically, the gonadotropin releasing hormone receptor (*Gnrhr*) (Fig. 5B) and the gonadotropin alpha and beta subunits: follicle stimulating hormone beta (*Fshb*), lutenizing hormone beta (*Lhb*), alpha glycoprotein hormones subunit (*Cga*) (Fig. 5C-E). In addition, RNAs for two other pituitary hormones, thyroid stimulating hormone beta (*Tshb*) and growth hormone (*Gh*), and growth hormone releasing hormone receptor (*Ghrhr*) were similarly elevated (Fig. 5F-H). Analyzing the data by sex revealed that all these genes were predominantly elevated in two out of three male, but not any female, responders (Fig. 5B-H).

Additional genes of interest with greater expression in responder mice include neurotensin (*Nts*) (1.7 log2 FC), which may be involved in migraine (Fig. 5I) (Theoharides et al., 2005), potassium two pore domain channel subfamily K member 12 *(Kcnk12)* (0.7 log2 FC), which has connections with pain and inflammation (Fig. 5J) (Marsh et al., 2012), and *Trpc5* (1 log2 FC), which can sense mechanical stimulation (Shen et al., 2015) Fig. 5K). The gene with the highest relative expression (26 log2 FC) was *AL731706.1*, which encodes a protein with unknown function predicted to have 50 amino acids and a transmembrane helix (https://www.uniprot.org/uniprot/A0A2I3BRA8#function) (Fig. 5L).

While not significantly different when analyzed as combined sexes, there were other genes of interest with significant differences between responders and nonresponders when analyzed within just males (Table 5-2) or females (Table 5-3). In male responder mice the dopamine receptor 2 (*Drd2*), prolactin (*Prl*), galanin receptor 3 (*Galr3*), and glycine receptor alpha 1 subunit (*Glra1*) were significantly higher (Fig. 5-1). In female responder mice, the *Rest*, *Trpc2*, gamma-aminobutyric acid type A receptor subunit beta 2 (*Gabrb2*) RNAs were higher, while *Kcnk5*, *Trpm8*, *Mlf1*, *Calcrl* RNAs were lower (Fig. 5-1). Of note, the transcriptional regulator *Rest* was not detectable in female nonresponders. The *Rest*, *Kcnk5*, and *Trpm8* genes are of particular interest since they were identified in a genome-wide migraine association study (Gormley et al., 2016) and *Calcrl* encodes the G protein-coupled receptor subunit of the canonical CGRP receptor (Aiyar et al., 1996).

A pathway analysis revealed that genes more highly expressed in responder mice were over-represented in pathways involved in G protein-coupled receptors and peptide hormones (Table 5-4). The pathways involving hormone ligand-binding receptors and glycoprotein hormones were exemplified by expression of *Cga*, *Tshb*, *Gnrhr*, *Lhb*, *Fshb*. These highly expressed genes were also over-represented in a number of pathways involved in pain modulation. These include the neuroactive ligand-receptor interaction (false discovery rate; FDR < 0.001) (Deng et al., 2018a; Jeong et al., 2018), signaling by G protein-coupled receptors (FDR < 0.001) (Stone and Molliver, 2009; Cottrell, 2019) and prolactin signaling pathway (FDR < 0.05) (Avona et al., 2019).

Similarly, a gene ontology analysis that shows connections between genes and their biological and cellular functions also revealed over-representation of genes in biological processes related to hormone signaling, including peptide hormone processing (FDR < 0.001), G protein-coupled receptor signaling pathway (FDR < 0.01) and regulation of signaling receptor activity (FDR < 0.01) (Table 5-5). These genes included *Cga*, *Tshb*, *Lhb*, *Fshb*, and *Nts*. Analysis of the gene ontology by sex also revealed peptide hormone processing, hormone activity, and pituitary gonadotropin complex in the male, but not the female responder cohort.

In contrast to genes more highly expressed in responder mice, the lower expressed genes were less informative. There were 98 genes with lower expression in responder mice (Table 5-1). In general, these genes were over-represented in pathways such as platelet activation, signaling and aggregation (FDR < 0.05), which is consistent with other studies showing PACAP being an inhibitor of platelet activation (Freson et al., 2004; Eneman et al., 2015) (Table 5-6). Other pathways include PI3K-Akt signaling pathway (FDR <0.05) and arachidonic acid metabolism (FDR < 0.05), which have been suggested to be involved in migraine (Puig-Parellada et al., 1993; Liu et al., 2017). Gene ontology revealed a number of biological functions involved in bone mineralization and ossification (Table 5-7). Genes involved in these biological functions include *BMP-7*, which is known to be negatively regulated by PACAP (Drahushuk et al., 2002).

In addition to gene expression, we also investigated sequence differences between responder and nonresponder mice. Genetic variance was expected since the CD-1 strain is outbred and the responders and nonresponders came from 2 litters each. Using FaST-LMM, which estimates and accounts for genetic relatedness, we identified 162 single nucleotide polymorphic (SNP) sites that had distinctly different genotypes between responder and nonresponder mice but were the same for all 6 mice (male and female) in each of the two groups. None of the other SNPs reached genome-wide significance (Fig. 5-2). The 162 SNPs were classified as 17 missense variations, 61 synonymous variations, 14 5’ UTR variations, 66 3’ UTR variations, 3 upstream gene variations, and 1 intron variation based on SNPEff analysis (Cingolani et al., 2012) (Table 5-8). Among the SNPs, three were in genes identified as differentially expressed between responder and nonresponder mice: in the coding region of *Atg10* and the untranslated region of *Kcnk12* and *Pmp2*. None of the variants were in PACAP receptor genes. Pathway analysis did not reveal any pathways that were significantly over-represented with genes impacted by missense variation. Similarly, no biological, cellular or molecular processes was over-represented by these genes.

### Pre-treatment with anti-PACAP monoclonal antibodies attenuates light aversion induced by both PACAP-38 and PACAP-27

We tested whether anti-PACAP monoclonal antibodies would be able to attenuate PACAP-induced light aversion using the same sequential treatment paradigm previously used with CGRP antibodies (Mason et al., 2017). For all antibody experiments, mice were prescreened for their response to PACAP and only the responder mice were further analyzed. After the first PACAP injection (Tx1: pre Ab), mice that responded to PACAP-38 spent significantly less time in light compared to vehicle treated mice (Fig. 6A, B). The mice that responded to PACAP-38 in the first treatment were then given either control isotype or anti-PACAP antibody. One day after antibody injection mice were injected with PACAP-38 and tested for light aversion (Tx2: post Ab). Pre-treatment with anti-PACAP antibody fully attenuated the effect of PACAP-38 (Fig. 6A, B). Mice treated with anti-PACAP antibody prior to PACAP-38 were indistinguishable from those treated with vehicle with either control or PACAP antibody, while those treated with the control antibody prior to PACAP-38 spent significantly less time in light compared to mice treated with anti-PACAP antibody or vehicle plus control or anti-PACAP antibody (Fig. 6B). Along with light aversion, antibody pre-treatment also blocked the effect the PACAP-38 had on resting, rearing behavior, and transitions (Fig. 6-1).

**Figure 6.**
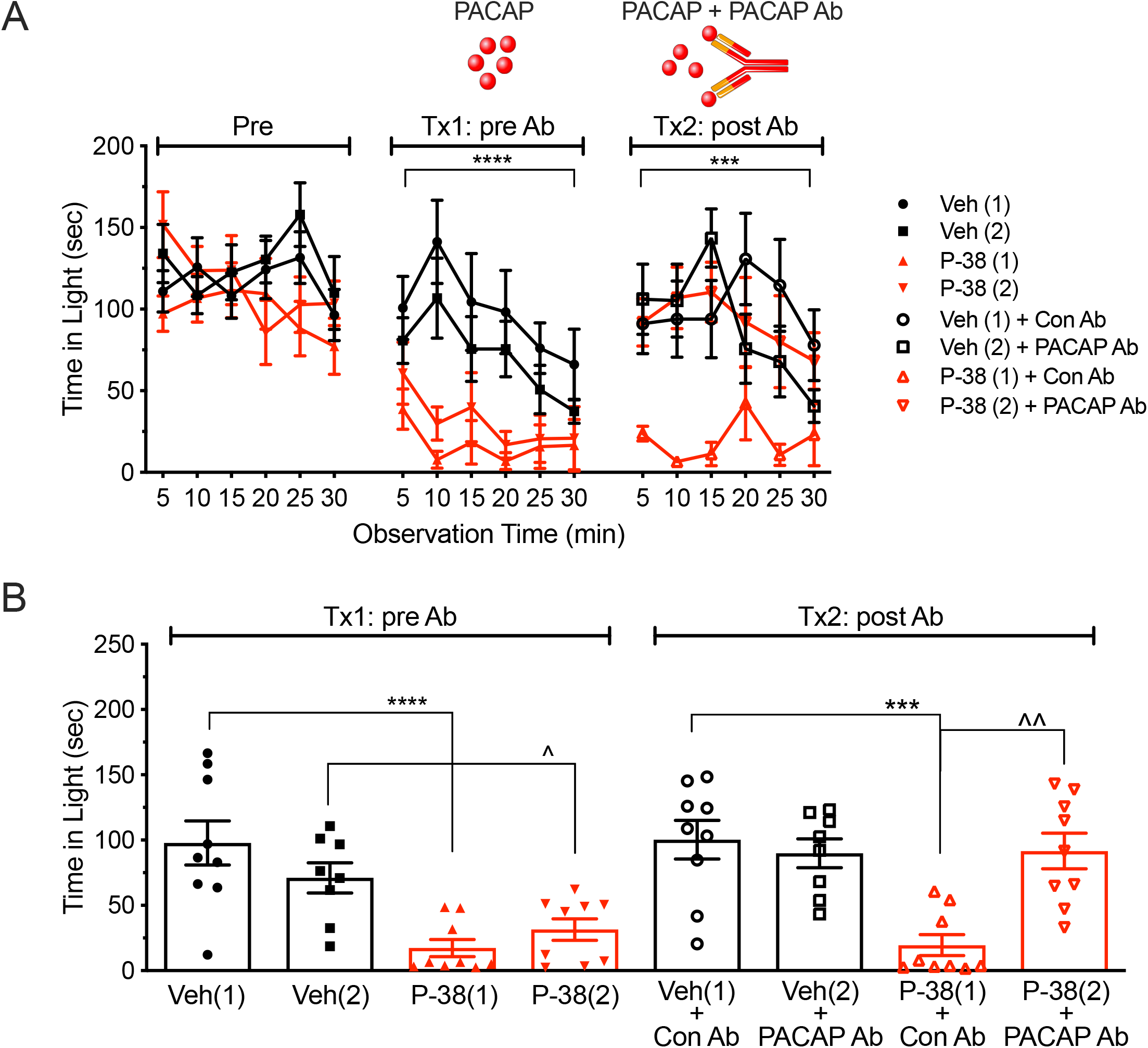
Pretreatment with anti-PACAP antibody inhibits PACAP-38 induced light aversion. **A)** Mice were divided into 4 groups, two of which would eventually get vehicle and two that would get PACAP-38. After a baseline pre-exposure (Pre), but before antibody treatments, mice were treated (Tx1: pre Ab) with vehicle (Veh(1), n=9; Veh(2), n=9) or PACAP-38 (0.3 mg/kg) (P-38(1), n=9; P-38(2), n=9). Three days after Tx1 mice were given an injection of control antibody (Con Ab) or anti-PACAP antibody (PACAP Ab), then 24 h later, mice were treated with vehicle or PACAP again (Tx2: 24 h post Ab). Mean (± SEM) time spent in the light compartment every 5 min over a 30 min period is shown. **B)** Data for individual mice from each treatment day are shown as the mean time (± SEM) in light per 5 min interval. Statistics are described in Table 1.

Since *in vitro* tests had shown that the PACAP antibody binds both PACAP-38 and PACAP-27 (Moldovan Loomis et al., 2019), we asked if the antibody could also block PACAP-27 actions. This question was relevant because PACAP-38 and PACAP-27 have the same receptor binding affinities and both can induce migraine in patients (Nilsson et al., 1994; Schytz et al., 2009; Hirabayashi et al., 2018; Ghanizada et al., 2020). The first step was to test the ability of PACAP-27 to induce light-aversion in CD-1 mice. The mice were first treated with PACAP-38 (Tx1: P-38) as a control to confirm the response rate, prior to treatment with PACAP-27 (Tx2: P-27) (0.2 mg/kg, equimolar to 0.3 mg/kg PACAP-38). When tested at 30 min after injection, the mice responded to PACAP-38, but not PACAP-27 (Fig. 7A). There was also no effect of PACAP-27 on motility under this condition (Fig. 7-1). Given that the two PACAP isoforms may have different stabilities (Bourgault et al., 2008), we tested them immediately after injection. Under these conditions, both PACAP-27 and PACAP-38 caused significant light aversion compared to vehicle, but PACAP-27 was significant only in the first 15 min post-injection (Fig. 7B). As with PACAP-38, there was a corresponding decrease in motility during this period (Fig. 7-1). Likewise, there was a trend towards the male mice spending less time in the dark but it was not significant.

**Figure 7.**
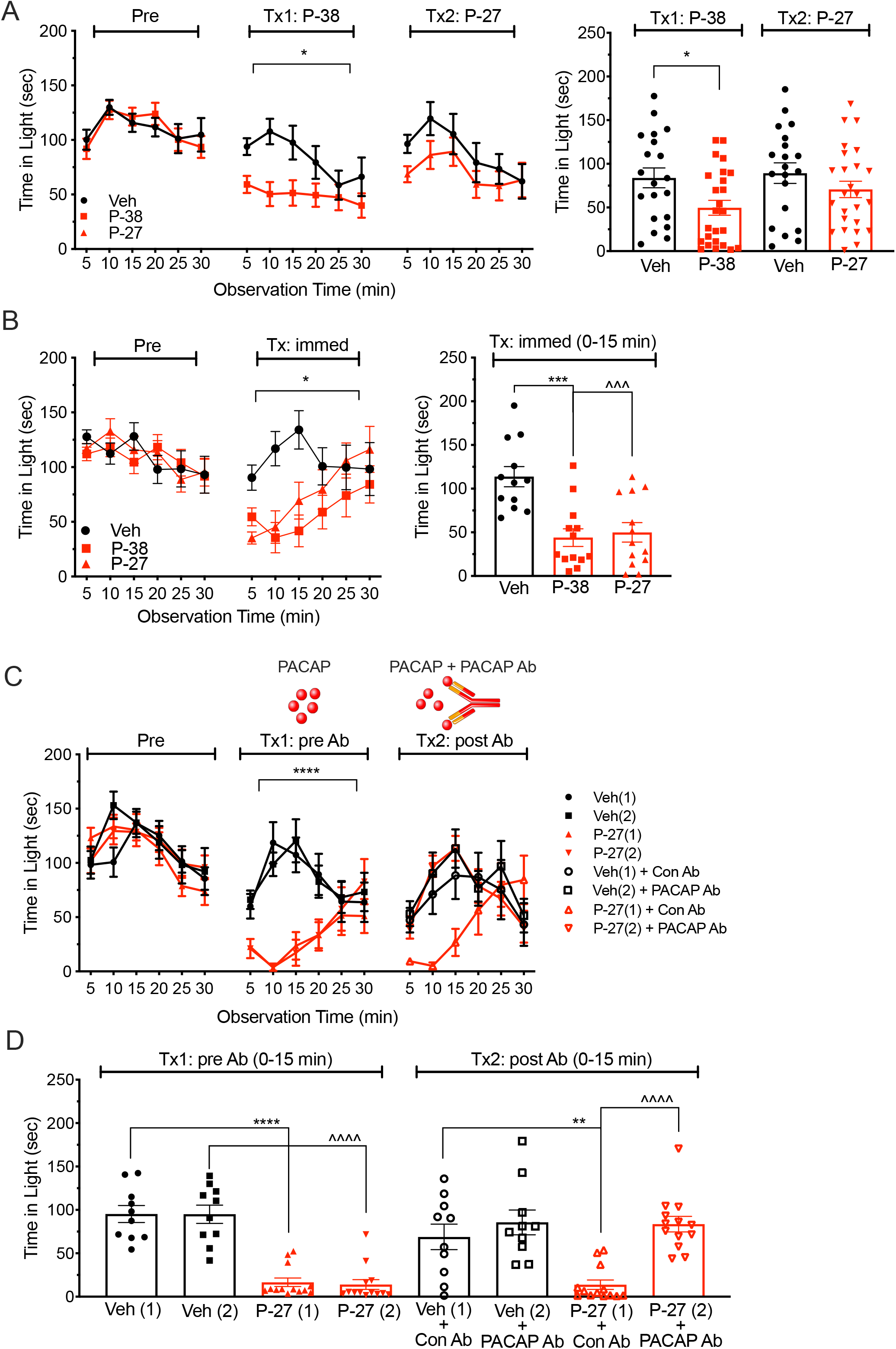
Immediate injection of PACAP-27 induces light aversion that can be blocked by anti-PACAP antibody. **A)** After a baseline pre-exposure (Pre), mice were treated (Tx1: P-38) with vehicle (Veh, n=20) or PACAP-38 (P-38, 0.3 mg/kg, n=25) and assessed for light aversion 30 min post injection. Three days later the same mice were treated (Tx2: P-27) with vehicle or PACAP-27 (P-27, 0.2 mg/kg) and assessed for light aversion 30 min post injection. Left panel: Mean (± SEM) time spent in the light every 5 min over a 30 min period is shown for each test day. Right panel: Data for individual mice from treatment days shown as the mean (± SEM) in light per 5 min interval. **B)** After a pre-exposure (Pre), mice were treated and tested immediately (Tx: immed) after injection of vehicle (n=12), P-38 (0.3 mg/kg, n=14), or P-27 (0.2 mg/kg, n=13). Left panel: Mean (± SEM) time spent in the light compartment every 5 min over the 30 min test period. Right panel: Data for individual mice from the first 15 min of treatment shown as the mean (± SEM) in light per 5 min interval. **C)** Mice were divided into 4 groups, two of which would eventually get vehicle and two that would get PACAP-27. After a baseline pre-exposure (Pre), but before antibody treatments, mice were treated (Tx1: pre Ab) with vehicle (Veh(1), n=10; Veh(2), n=10) or PACAP-27 (0.2 mg/kg) (P-27(1), n=13; P-27(2), n=13). Three days after Tx1 mice were given an injection of control antibody (Con Ab) or anti-PACAP antibody (PACAP Ab), then 24 h later, mice were treated with vehicle or PACAP again (Tx2: post Ab). Mean (± SEM) time spent in the light compartment every 5 min over a 30 min period is shown. **D)** Data for individual mice from the first 15 min of treatment before antibody (Tx1: pre Ab (0-15 min)) and after antibody (Tx2: post Ab (0-15 min)) shown as the mean ± SEM in light per 5 min interval. Statistics are described in Table 1.

Hence, following antibody pretreatments, we tested mice immediately after PACAP-27 injection. As with the PACAP-38 paradigm, we first treated the mice with PACAP-27 to identify the responder population (Fig. 7C, D). Subsequent pre-treatment with anti-PACAP antibody blocked the effect of PACAP-27 (Fig. 7C, D). As a control, mice treated with PACAP-27 and control antibody spent significantly less time in light compared to mice treated with PACAP-27 plus anti-PACAP antibody or vehicle plus control antibody (Fig. 7C, D). Antibody pre-treatment also blocked the effect of PACAP-27 on resting, rearing, and transitions (Fig. 7-2).

### Anti-PACAP antibodies do not inhibit CGRP-induced light aversion, nor do anti-CGRP antibodies inhibit PACAP-induced light aversion

PACAP and CGRP have similar biological activities (Kaiser and Russo, 2013), and PACAP has been reported to elicit CGRP release (Jansen-Olesen et al., 2014), which raises the question whether the two peptides might act in series to trigger migraine. To address this relationship, anti-PACAP and anti-CGRP antibodies were tested for their ability to cross-inhibit light aversion induced by PACAP or CGRP. We first tested whether anti-PACAP antibody could inhibit CGRP-induced light aversion. Mice were treated with CGRP, which caused significant light aversion compared to vehicle treated mice (Fig. 8A, B). The day prior to the 2^nd^ CGRP treatment, mice were given either control antibody or anti-PACAP antibody. Treatment of mice with anti-PACAP antibody, did not ameliorate light aversion to CGRP compared to vehicle or control antibody groups (Fig. 8A, B) and were not significantly different from each other. Thus, pre-treatment with anti-PACAP antibody was not able to inhibit the effect of CGRP. Consistent with the light aversion, the PACAP antibody was not able to inhibit CGRP effects on motility (Fig. 8-1).

**Figure 8.**
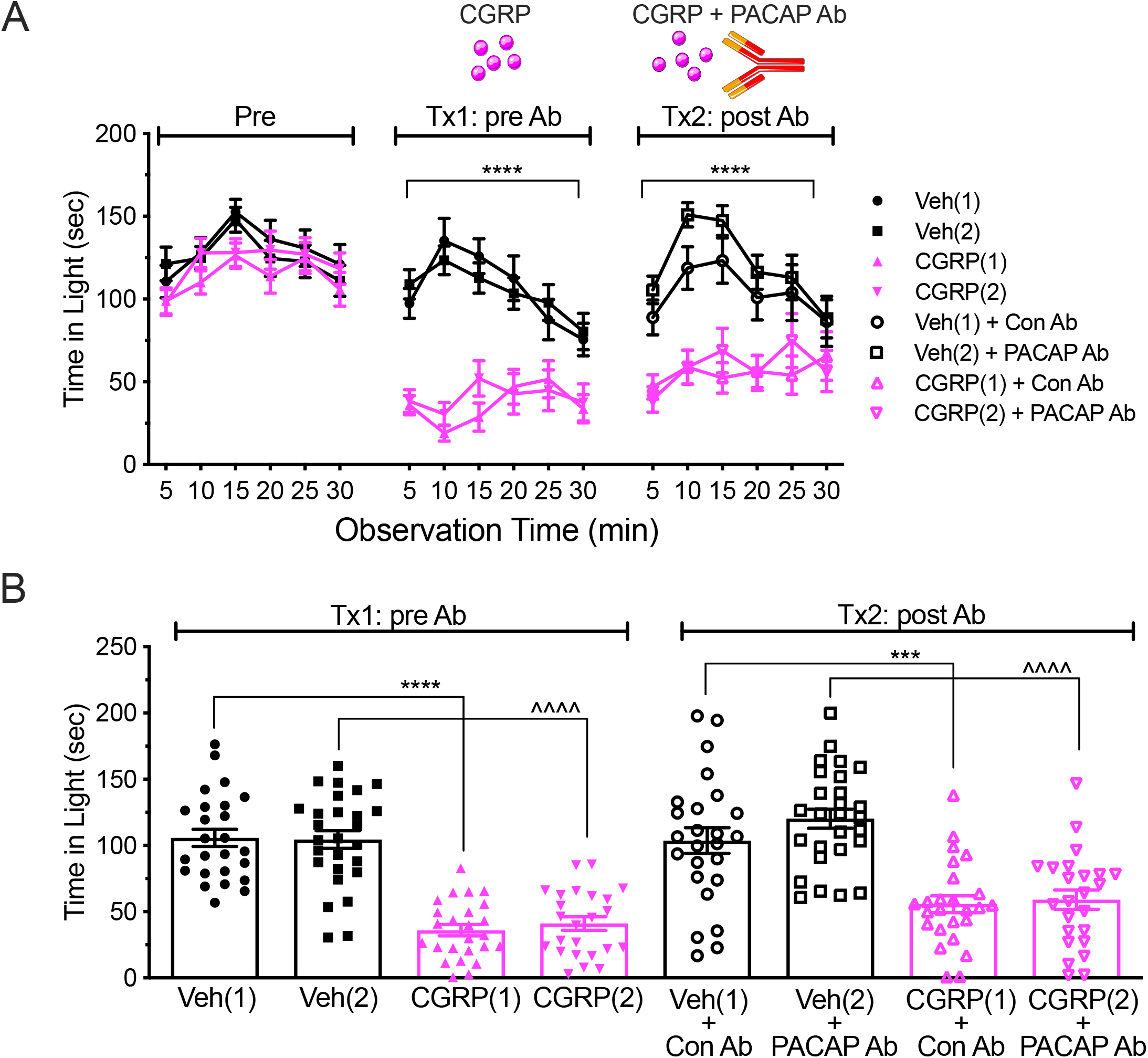
Anti-PACAP antibody does not inhibit CGRP-induce light aversion. **A)** Mice were divided into 4 groups, two of which would eventually get vehicle and two that would get CGRP. After a baseline pre-exposure (Pre), but before antibody treatments, mice were treated (Tx1: pre Ab) with vehicle (Veh(1), n=25; Veh(2), n=28) or CGRP (0.1 mg/kg) (CGRP(1), n=25; CGRP(2), n=24). Three days after Tx1 mice were given an injection of control antibody (Con Ab) or anti-PACAP antibody (PACAP Ab), then 24 h later, mice were treated with vehicle or CGRP again (Tx2: post Ab). The mean (± SEM) time spent in the light compartment every 5 min over a 30 min period is shown. **B)** Data for individual mice from each treatment day are shown as the mean time (± SEM) in light per 5 min interval. Statistics are described in Table 1.

Finally, we tested whether an anti-CGRP antibody could inhibit PACAP-induced light aversion. Mice were first treated with PACAP-38 and both cohorts of responders spent significantly less time in the light compared to the vehicle groups (Fig. 9A, B). The mice that responded to PACAP-38 in the first treatment were then given either the control isotype or anti-CGRP antibody. One day after antibody injection mice were injected with PACAP-38 again and assessed for light aversion. Pre-treatment with anti-CGRP antibody was not able to inhibit the effect of PACAP-38 (Fig. 9A, B). Anti-CGRP antibody was also not able to rescue the effect PACAP had on motility (Fig. 9-1).

**Figure 9.**
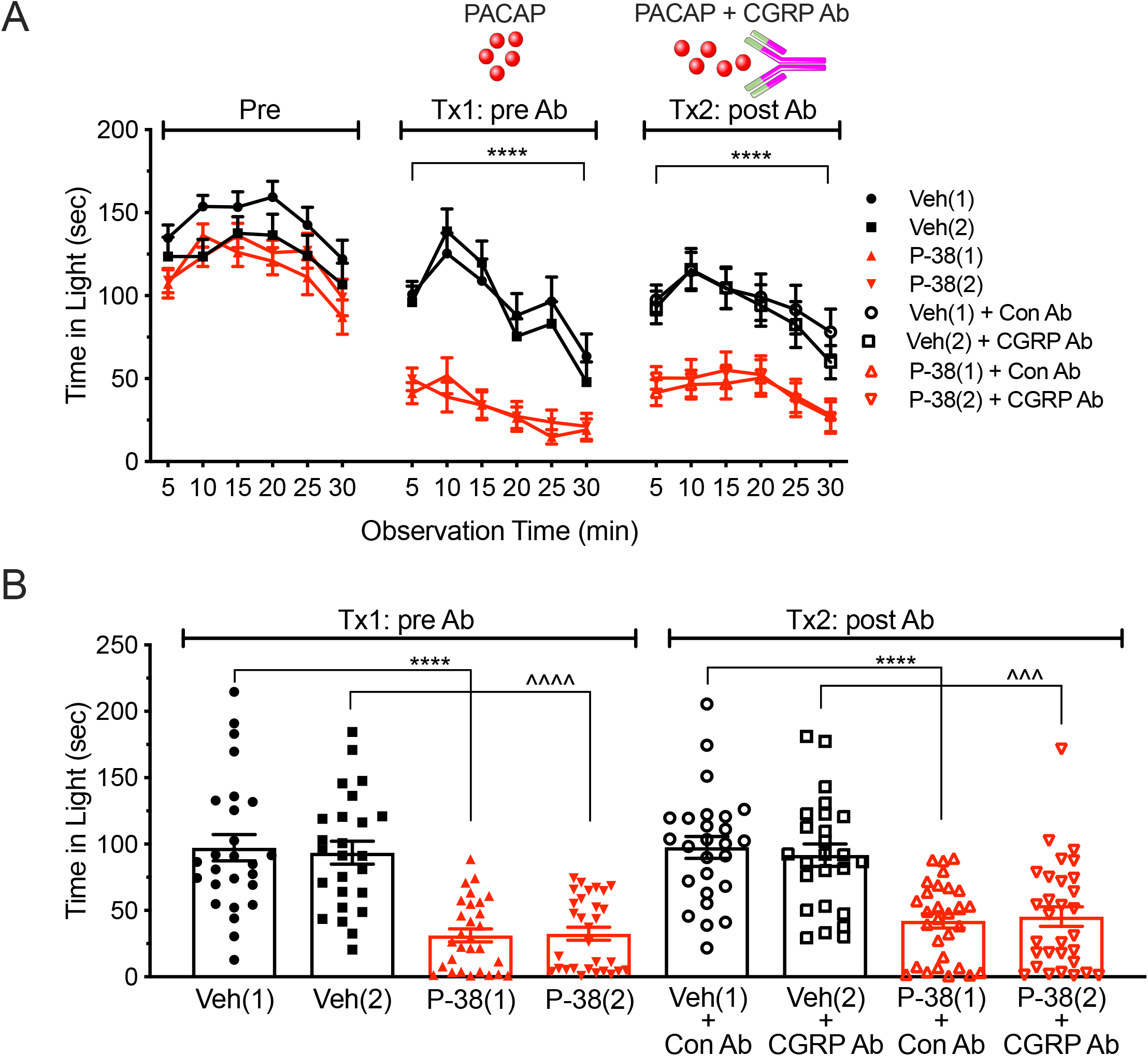
Anti-CGRP antibody does not inhibit PACAP-induced light aversion. **A)** Mice were divided into 4 groups, two of which would eventually get vehicle and two that would get PACAP. After a baseline pre-exposure (Pre), but before antibody treatments, mice were treated (Tx1: pre Ab) with vehicle (Veh(1), n=26; Veh(2), n=25) or PACAP-38 (0.3 mg/kg) (P-38(1), n=29; P-38(2), n=30). Three days after Tx1 mice were given an injection of control antibody (Con Ab) or anti-CGRP antibody (CGRP Ab), then 24 h later, mice were treated with vehicle or PACAP again (Tx2: post Ab). Mean (± SEM) time spent in the light compartment every 5 min over a 30 min period is shown. **B)** Data for individual mice from each treatment day are shown as the mean time (± SEM) in light per 5 min interval. Statistics are described in Table 1.

## Discussion

In this study we report that both PACAP-27 and PACAP-38 can induce light aversion in CD-1 mice. PACAP is predicted to play a role in migraine given it can induce migraine-like headaches (Schytz et al., 2009; Ghanizada et al., 2020), plasma levels are elevated during migraine (Tuka et al., 2013), and sumatriptan reduces circulating PACAP coincident with headache relief (Zagami et al., 2014). PACAP-induced light aversion in mice is consistent with the ability of PACAP to induce photophobia in people (Schytz et al., 2009) and with a previous study by Helyes and colleagues who reported PACAP-38 elicited light aversion in mice on a CD-1 background (Markovics et al., 2012). While they did not report populations of responders and nonresponders, this may be explained by different sources of mice and/or the possibility of genetic homogeneity in their transgenic CD-1 line. In comparison to CGRP, the PACAP-evoked light aversion and reduced motility in mice is similar to the effects of CGRP in this and a previous study (Mason et al., 2017). In particular, the mice rested more, but only in the dark zone, which is reminiscent of migraine patients preferring to rest in the dark.

A strength of this study was that we used CD-1 mice, which are a genetically diverse, outbred strain of mice. This strategy revealed distinct groups of PACAP responders and nonresponders that may be predictive of variability expected in humans (Aldinger et al., 2009). The responder and nonresponder phenotypes were stable and inheritable. While the inheritance pattern appeared to be dominant, this was not supported statistically, which suggests multigenic inheritance or incomplete penetrance. To understand the genetic basis underlying the responder phenotype, we did an RNA-seq analysis of trigeminal ganglia, which revealed a number of candidate genes, including pituitary hormones, receptors, and ion channels.

Perhaps most intriguing among the candidate genes is elevation of pituitary hormones and receptor RNAs in the trigeminal ganglia of male responder mice. RNAs for the gonadotropin hormones FSH and LH, along with the receptor that controls their release (GnRHR) were elevated in a subset of males. These proteins are most often viewed in the context of their regulation of gonadal hormones, however expression of gonadotropin releasing hormone has been reported in trigeminal ganglia neurons believed to be involved in nociception and chemosensory avoidance (Umatani and Oka, 2019). Increased expression of GnRHR in responder mice could thus enhance a local positive feedback loop. RNAs for other pituitary hormones (GH, TSH, prolactin) and receptors (GHRHR, Drd2) were also elevated only in responder males. The human dopamine receptor *DRD2* gene was identified in a migraine association study (Deng et al., 2018b), and dopamine has long been implicated in migraine (Goadsby et al., 2017). Likewise, the elevated expression of prolactin is reminiscent of the role of prolactin and its receptor in sex-specific pain responses (Dussor et al., 2018; Patil et al., 2019). The regulation may also be bidirectional since a study with male rats found that PACAP increased GnRH release (Li et al., 1996). How these hormones, as well as other changes in RNA levels observed between the responder and nonresponder populations may facilitate PACAP actions remains to be seen.

In contrast to the pituitary hormones and receptors, the neuropeptide *neurotensin* and *Trpc5* and *Kcnk12* ion channel RNAs were elevated in both male and female PACAP responder mice. The trigeminal ganglia expresses both neurotensin and its receptor (Manteniotis et al., 2013), which opens the possibility of autocrine and paracrine regulation within the trigeminal microenvironment. Furthermore, neurotensin can cause mast cell degranulation (Theoharides et al., 2005), vasodilation (Uddman and Edvinsson, 1989), cytokine production in neutrophils and macrophages (Lemaire, 1988; Li et al., 1996), and either reduce or enhance pain responses (Dobner, 2006; Boules et al., 2013). These actions are consistent with mechanisms that influence a pro-inflammatory milieu implicated in migraine pathogenesis. Likewise, Trpc5 can sense mechanical stimulation (Shen et al., 2015), which might possibly contribute to sensing of vascular tone in the dura or cutaneous stimuli. Increased expression of the two pore domain potassium channel *Kcnk12* in PACAP responder mice is intriguing given that a related family member, *KCNK18* (TRESK), has been linked to migraine with aura (Lafreniere et al., 2010). While *Kcnk12* is known to be expressed in the trigeminal ganglia (Manteniotis et al., 2013), little is known about its function, although it is regulated by intracellular trafficking (Bichet et al., 2015) and is elevated after inflammation in the dorsal root ganglia (Marsh et al., 2012). Other RNAs were elevated in the responder and nonresponder populations in both sexes or in only males or females. While their roles are not established, four of the genes (*Rest*, *Kcnk5*, *Trpm8*, *Mlf1*) are among the 38 loci associated with migraine in a large genome-wide association study (Gormley et al., 2016). Finally, SNPs were identified in the *Kcnk12* and *Pmp2* (myelin protein 2) untranslated regions and coding region of *Atg10* (E2-like enzyme involved in autophagy). Future studies understanding the role of these SNPs and the differentially expressed genes may prove to be important in trigeminal pain states.

When compared to PACAP-38, injection of PACAP-27 only caused transient light aversion. Whether PACAP-27 is less effective than PACAP-38 in inducing migraine in humans has not been directly tested, although PACAP-27 caused migraine-like attacks in only 55% of patients (Ghanizada et al., 2020), while PACAP-38 was effective in 91% of patients (Schytz et al., 2009). Because there are conflicting reports on the relative stability of the two PACAP isoforms (Bourgault et al., 2008), we cannot rule out a pharmacokinetic explanation for the transient PACAP-27 response in mice. However, another possible mechanism for the short-lived light aversion observed after PACAP-27 is that PACAP-38 causes greater dilation of the middle meningeal artery than PACAP-27 (Bhatt et al., 2014). Interestingly, PACAP-38-induced dilation of the artery was abolished in the absence of mast cells, whereas this had no effect on arterial changes from PACAP-27 infusion, suggesting that the sustained effects of PACAP-38 might be due to an inflammatory cascade of events. In fact, PACAP-38, and not PACAP-27, has been reported to degranulate mast cells via the orphan MrgB3 receptor (Pedersen et al., 2019). Studies exploring the role of MrgB3 may give insights to how PACAP-38 evokes symptoms of migraine in mice and people.

We found a trend, albeit not statistically significant, that PACAP had a greater effect in male compared to female mice. The reason for this trend is not known, but might reflect difference in actions on neutrophils. PACAP-38 can enhance inflammatory and chemotactic markers in neutrophils via activation of classical PACAP receptors (Kinhult et al., 2002; Harfi et al., 2004) and neutrophils are reportedly higher in males compared to females during acute inflammation (Kay et al., 2015). Neutrophil recruitment and activation may be relevant given reports of increased neutrophil to lymphocyte ratio during migraine attacks (Karabulut et al., 2016).

The inability of the anti-CGRP and anti-PACAP antibodies to inhibit light aversion induced by the other peptide suggests that these peptides can act independently of each other in the periphery. The possibility of sequential or dependent pathways was raised by the similar properties (Kaiser and Russo, 2013), co-expression in ~20% of trigeminal ganglia neurons (Eftekhari et al., 2015), and most notably PACAP-38 causing CGRP release in the trigeminal nucleus caudalis (although not from the dura or ganglia) (Jansen-Olesen et al., 2014). Against that hypothesis, a recent clinical study did not detect increased CGRP levels after PACAP-38 infusion (Guo et al., 2017). Hence, we propose that in the periphery CGRP and PACAP act by distinct, parallel paths that may converge downstream of their receptors on a common target.

The efficacy of the humanized anti-PACAP antibody to block PACAP-induced light aversion in mice supports the potential use of this antibody to attenuate or prevent migraine in patients. An alternative approach to ligand-blocking antibodies is to target PACAP receptors (VPAC1, VPAC2, PAC1) (Rubio-Beltran et al., 2018). Initial focus has been on the PAC1 receptor in part because it is highly selective to PACAP while the other PACAP receptors are equally activated by VIP (Rubio-Beltran et al., 2018) and VIP failed to induce migraine in people (Hansen et al., 2006). However, a role for VPAC receptors in migraine pathology should not be discarded. We have shown that, similar to our results when testing PACAP-27, VIP can also induce light aversive behavior in mice if measured immediately after administration, consistent with its shorter half-life compared to PACAP (Mason et al., 2020). This suggests that the VPAC receptors are capable of inducing migraine-like phenotypes in mice, which has recently been validated by prolonged VIP infusion causing delayed headache in people (Pellesi et al., 2020). Alternatively, it is possible that PACAP-38 involvement in migraine may be independent of the VPAC or PAC1 receptors. PACAP-38 can act in the trigeminal nucleus caudalis via a PAC1-independent mechanism (Jansen-Olesen et al., 2014) and the orphan receptor Mrgb3 can mediate PACAP actions on mast cells (Pedersen et al., 2019). Future studies targeting PACAP and its receptors in migraine are certainly warranted.

## Acknowledgments

This work was supported by the National Institutes of Health (NS098825 to BNM, NS075599 to AFR) and a grant from Alder Biopharmaceuticals. Additional support was provided by NIH/NEI Center Support Grant P30EY025580 (University of Iowa). The contents do not represent the views of VA or the United States Government. We thank Michael Anderson and Kai Wang for help interpreting the inheritance pattern and Johannes Ledolter for help with statistical analysis.

## Figure Legends

**Figure 3-1. PACAP reduces motility only in responder mice**. Motility data were collected at the same time as light aversion data from the same mice shown in Figure 3. **A)** Resting time in light and dark zones. Data for individual mice from treatment day shown as mean percent time (± SEM) spent resting in each zone per 5 min interval. **B)** Rearing in light and dark zones. Data for individual mice from treatment day shown as mean (± SEM) number of vertical beam breaks in each zone per 5 min interval. **C)** Transitions between light and dark zones. Data for individual mice from treatment day shown as mean (± SEM) number of transitions per 5 min interval. Statistics are described in Table 1-1.

**Figure 4-1. Motility of F1 responders and nonresponders**. Motility data were collected at the same time as light aversion data from the same mice shown in Figure 4. **A)** Resting time in light and dark zones. Data for individual mice from treatment days shown as mean percent time (± SEM) resting in each zone per 5 min interval. **B)** Transitions between light and dark zones. Data for individual mice from treatment days shown as the mean (± SEM) number of transitions per 5 min interval. Statistics are described in Table 1-1.

**Figure 5-1. Differential gene expression between responder and nonresponder mice that was significant in male or female, but not combined sexes**. Gene level counts were normalized to size factors calculated by DESeq2 to correct for library size. Normalized counts were compared between responders and nonresponders using Wald test for significance testing. Male and female mice in each cohort are indicated by blue circles and red squares symbols, respectively. (A) dopamine receptor D2 (*Drd2)*, (B) prolactin (*Prl*), (C) galanin receptor 3 (*Galr3)*, (D) glycine receptor alpha subunit (*Glra1*), (E) *Rest*, (F) *Trpc2*, (G) *Gabrb2*, (H) *Kcnk5*, (I) *Trpm8*, (J) *Mlf1*, (K) *Calcrl*. The sex with a statistical difference is indicated as (m) for males and (f) for females. Statistics are described in Table 1-1.

**Figure 5-2. Manhattan plot of genome-wide association analysis for SNPs associated with responsiveness to PACAP**. Red line indicates genome-wide significant threshold which was calculated using the Bonferroni method. The 162 SNPs that had an assigned p-value of “NA” (not available) by the FaST-LMM program due to distinct genotype differences between responders and nonresponders were replaced with a low p-value of 1.175e-38 to reflect the deterministic separation of genotypes and their phenotypes as described in Table 5-8.

**Figure 6-1. Pre-treatment with anti-PACAP antibody blocks PACAP-38 effects on motility**. Motility data were collected at the same time as light aversion data from the same mice shown in Figure 6. **A)** Resting time in light and dark zones. Data for individual mice from treatment days shown as mean percent time (± SEM) spent resting per 5 min interval. **B)** Rearing in light and dark zones. Data for individual mice from treatment days shown as mean (± SEM) number of vertical beam breaks per 5 min interval. **C)** Transitions between the light and dark zones. Data for individual mice from treatment days shown as mean (± SEM) number of transitions per 5 min interval. Statistics are described in Table 1-1.

**Figure 7-1. PACAP-27 reduces motility but only when assessed after immediate injection**. Motility data were collected at the same time as light aversion data from the same mice shown in Figure 7A, B. Panels A, C, E are from Figure 7A, measured at 30 min post-injection. Panels B, D, F are from Figure 7B, measured immediately after injection and showing only the first 15 min. **A, B)** Resting time in light and dark zones. Data for individual mice from treatment days shown as mean percent time (± SEM) spent resting per 5 min interval. **C, D)** Rearing in light and dark zones. Data for individual mice from treatment days shown as mean (± SEM) number of vertical beam breaks per 5 min interval. **E, F)** Transitions between the light and dark zones. Data for individual mice from treatment days shown as mean (± SEM) number of transitions per 5 min interval. Statistics are described in Table 1-1.

**Figure 7-2. Pre-treatment with anti-PACAP antibody blocks the effect PACAP-27 has on motility**. Motility data were collected at the same time as light aversion data from the same mice shown in Figure 7C and showing only the first 15 min of data, as shown in Figure 7D. **A)** Resting time in light and dark zones. Data for individual mice from treatment days shown as mean percent time (± SEM) spent resting per 5 min interval. **B)** Rearing in light and dark zones. Data for individual mice from treatment days shown as mean (± SEM) number of vertical beam breaks per 5 min interval. **C)** Transitions between the light and dark zones. Data for individual mice from treatment days shown as mean (± SEM) number of transitions per 5 min interval. Statistics are described in Table 1-1.

**Figure 8-1. Pre-treatment with anti-PACAP antibody does not block the effect CGRP has on motility**. Motility data were collected at the same time as light aversion data from the same mice shown in Figure 8. **A)** Resting time in light and dark zones. Data for individual mice from treatment days shown as mean percent time (± SEM) spent resting per 5 min interval. **B)** Rearing in light and dark zones. Data for individual mice from treatment days shown as mean (± SEM) number of vertical beam breaks per 5 min interval. **C)** Transitions between the light and dark zones. Data for individual mice from treatment days shown as mean (± SEM) number of transitions per 5 min interval. Statistics are described in Table 1-1.

**Figure 9-1. Pre-treatment with anti-CGRP antibody does not block the effect PACAP has on motility**. Motility data were collected at the same time as light aversion data from the same mice shown in Figure 9. **A)** Resting time in light and dark zones. Data for individual mice from treatment days shown as mean percent time (± SEM) spent resting per 5 min interval. **B)** Rearing in light and dark zones. Data for individual mice from treatment days shown as mean (± SEM) number of vertical beam breaks per 5 min interval. **C)** Transitions between the light and dark zones. Data for individual mice from treatment days shown as mean (± SEM) number of transitions per 5 min interval. Statistics are described in Table 1-1.

## Table Legends

**Table 1-1. Statistical analyses for extended data**. Analyses are described for each figure with extended data not in the main manuscript. FDR values are assigned NA (not available) when genes do not pass independent filtering criteria performed by DESeq2 as described in the DESeq2 vignette (http://bioconductor.org/packages/release/bioc/vignettes/DESeq2/inst/doc/DESeq2.html).

**Table 4-1. F1 progeny responder frequency**. The number and percentage of progeny from the indicated crosses that responded following PACAP treatments are listed. For comparison, the same criteria were used to assign responder and nonresponder labels was applied following vehicle treatments.

**Table 5-1. Relative gene expression in responder and nonresponder mice**. Trigeminal ganglia from 6 responder and 6 nonresponder mice were assessed for differential gene expression using Wald test in DESeq2. Correction for multiple testing (i.e.padj) was performed using the false discovery rate method. Relative change in gene expression is expressed as log2FC where a positive value denotes a higher gene expression in responder mice. In contrast, a negative log2FC denotes a lower gene expression in responder mice. Table is sorted according to log2FC. As described in the DESeq2 manual, baseMean is the mean of the normalized count values divided by the size factors of all samples in the dataset, lfcSE is log fold change standard error, stat is wald statistic and padj is adjusted p-value.

**Table 5-2. Relative expression in only male responder and nonresponder mice**. Trigeminal ganglia from 3 male responder and 3 male nonresponder mice were assessed for differential gene expression using the Wald test in DESeq2. Correction for multiple testing (i.e.padj) was performed using the false discovery rate method. Relative change in gene expression is expressed as log2FC where a positive value denotes a higher gene expression in responder mice. In contrast, a negative log2FC denotes a lower gene expression in responder mice. Data are organized in the same way as Table 5-1. Table is sorted according to log2FC. As described in the DESeq2 manual, baseMean is the mean of the normalized count values divided by the size factors of all samples in the dataset, lfcSE is log fold change standard error, stat is wald statistic and padj is adjusted p-value.

**Table 5-3. Relative expression in only female responder and nonresponder mice**. Trigeminal ganglia from 3 female responder and 3 female nonresponder mice were assessed for differential gene expression using the Wald test in DESeq2. Correction for multiple testing (i.e.padj) was performed using the false discovery rate method. Relative change in gene expression is expressed as log2FC where a positive value denotes a higher gene expression in responder mice. In contrast, a negative log2FC denotes a lower gene expression in responder mice. Data are organized in the same way as Table 5-1. Table is sorted according to log2FC. As described in the DESeq2 manual, baseMean is the mean of the normalized count values divided by the size factors of all samples in the dataset, lfcSE is log fold change standard error, stat is wald statistic and padj is adjusted p-value.

**Table 5-4. Pathway analysis of responder mice**. Genes that were more highly expressed in responder mice were used for pathway analysis. Correction for multiple testing was performed using the false discovery rate method. Table is ordered according to the false discovery rate (FDR). Source – database, external_id – pathway ID from the respective database, members_input_overlap – differentially expressed genes that overlap with set members in the pathway, members_input_overlap_geneids – Entrez IDs of genes that overlap with set members in the patway, effective size – number of set members in the pathway.

**Table 5-5. Gene ontology of responder mice**. Genes that were more highly expressed in responder mice were used for gene ontology analysis. Correction for multiple testing was performed using the false discovery rate method. Table is ordered according to the false discovery rate (FDR). Source – database, term_goid – Gene ontology term ID, term_category – biological processes (b), molecular function (m) and cellular component (c), term_level – gene ontology term level, members_input_overlap – differentially expressed genes that overlap with set members in the pathway, members_input_overlap_geneids – Entrez IDs of differentially expressed genes that overlap with set members in the patway, effective size – number of set members in the pathway.

**Table 5-6. Pathway analysis of nonresponder mice**. Genes that were more highly expressed in nonresponder mice were used for pathway analysis. Correction for multiple testing was performed using the false discovery rate method. Table is ordered according to the false discovery rate (FDR). Source – database, external_id – pathway ID from the respective database, members_input_overlap – differentially expressed genes that overlap with set members in the pathway, members_input_overlap_geneids – Entrez IDs of genes that overlap with set members in the patway, effective size – number of set members in the pathway.

**Table 5-7. Gene ontology of nonresponder mice**. Genes that were more highly expressed in nonresponder mice were used for gene ontology analysis. Correction for multiple testing was performed using the false discovery rate method. Table is ordered according to the false discovery rate (FDR). Source – database, term_goid – Gene ontology term ID, term_category – biological processes (b), molecular function (m) and cellular component (c), term_level – gene ontology term level, members_input_overlap – differentially expressed genes that overlap with set members in the pathway, members_input_overlap_geneids – Entrez IDs of differentially expressed genes that overlap with set members in the patway, effective size – number of set members in the pathway.

**Table 5-8. Gene SNPs in responder and nonresponder mice**. GWAS analysis was performed using FaST-LMM. The putative effect of the SNPs was annotated using SnpEff. P-values were assigned as “NA” (not available) by FaST-LMM due to distinct genotype differences between responders and nonresponders. This is due to distinct differences in genotypes between responders and nonresponders and is indicative of an extremely low p-value. Table is ordered according to chromosome location and base position.

