## Supplemental Figures and Tables for "PACAP induces light aversion in mice by an inheritable mechanism independent of CGRP"

Fig 3-1

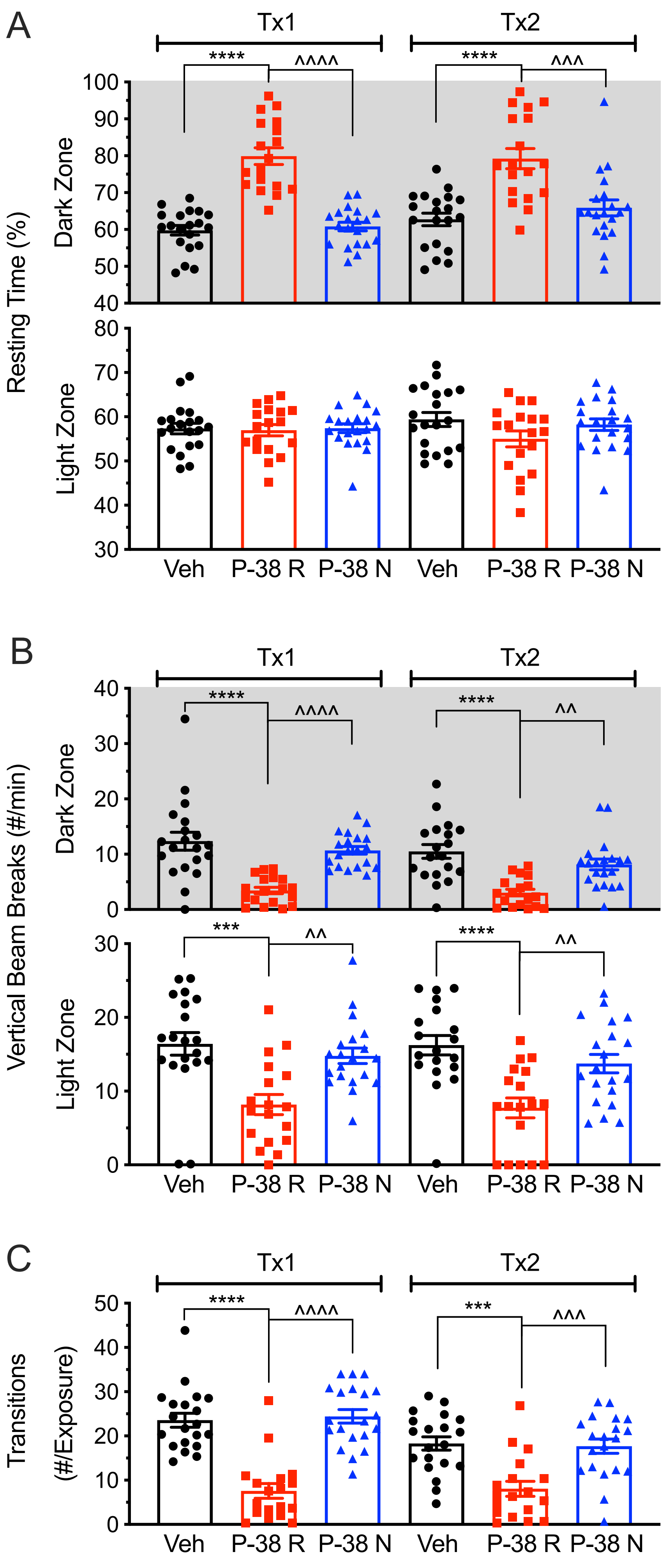

A
Tx1
Tx2
****
^^^^
****
^^^
Resting Time (%)
Dark Zone
Light Zone
100
90
80
70
60
50
40
80
70
60
50
40
30
Veh
P-38 R
P-38 N
Veh
P-38 R
P-38 N
B
Tx1
Tx2
****
^^^^
****
^^
***
^^
****
^^
Vertical Beam Breaks (#/min)
Dark Zone
Light Zone
40
30
20
10
0
30
20
10
0
Veh
P-38 R
P-38 N
Veh
P-38 R
P-38 N
C
Tx1
Tx2
****
^^^^
***
^^^
Transitions
(#/Exposure)
50
40
30
20
10
0
Veh
P-38 R
P-38 N
Veh
P-38 R
P-38 N

Fig 4-1

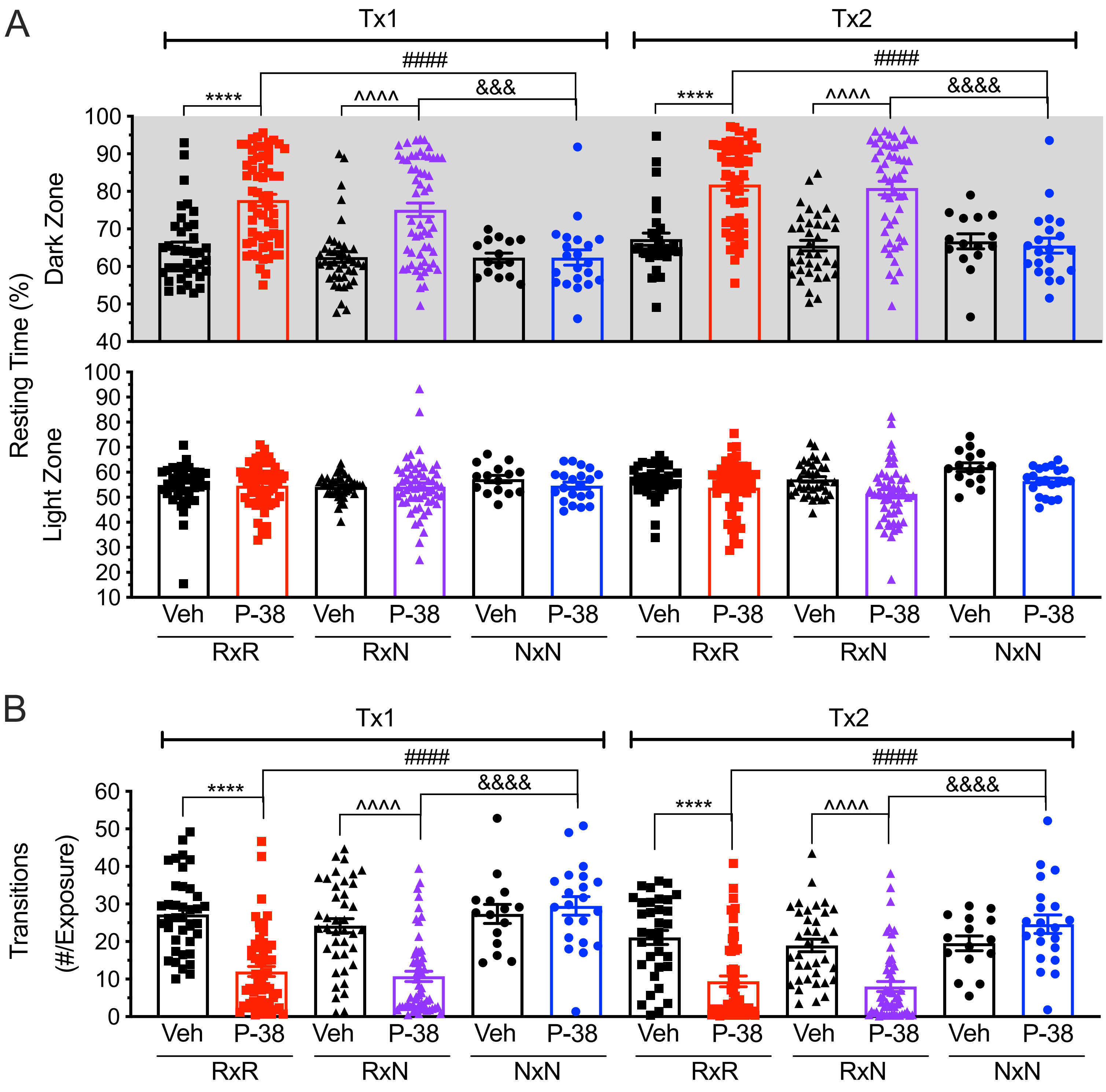
A
Tx1
Tx2
####
****
^^^^
&&&
&&&&
Resting Time (%)
Dark Zone
Light Zone
100
90
80
70
60
50
40
30
20
10
Veh
P-38
RxR
RxN
NxN
B
Transitions
(#/Exposure)
0

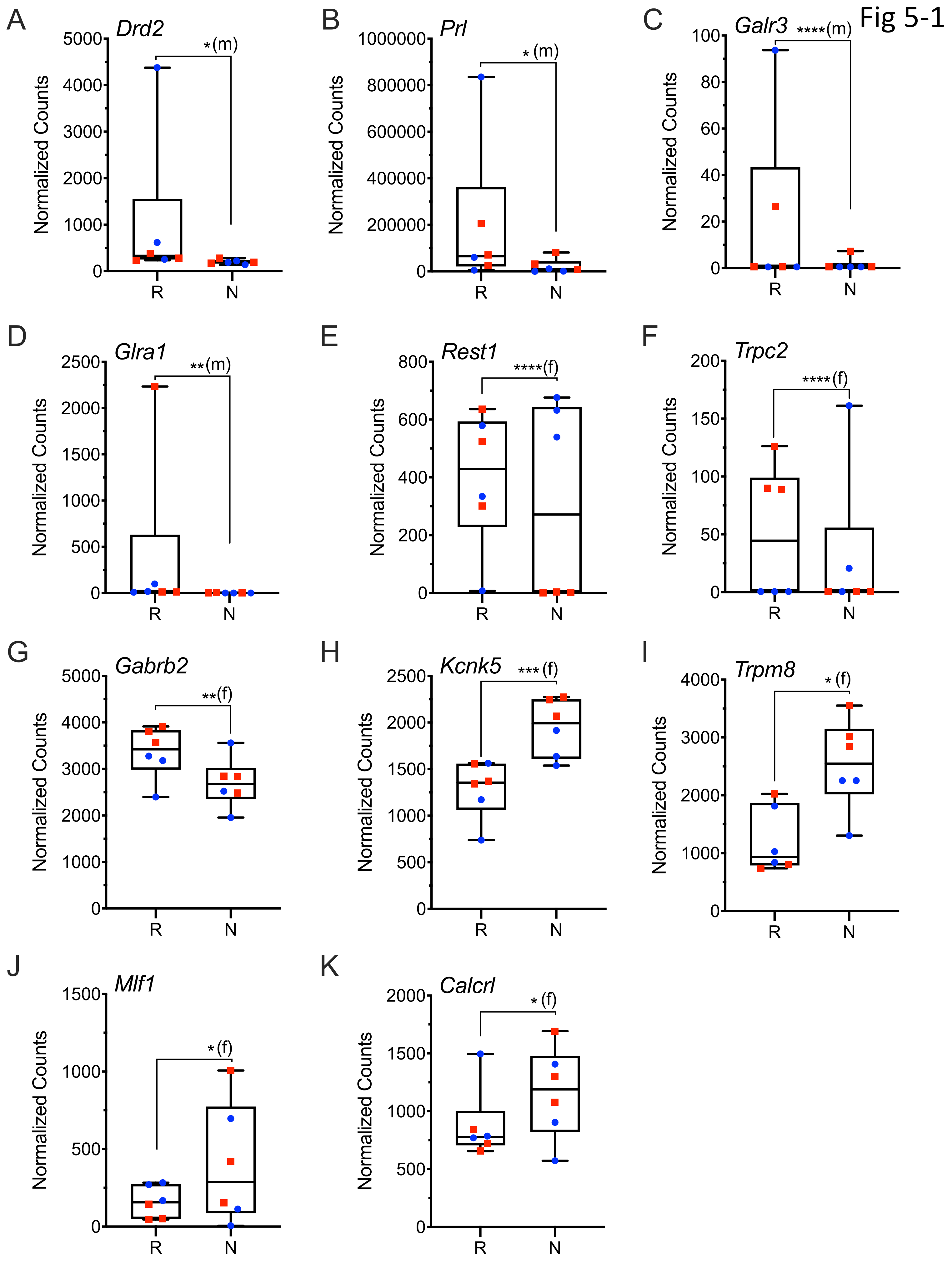

Fig 5-1
A
Drd2
*(m)
B
Prl
*(m)
C
Galr3
****(m)
D
Glra1
**(m)
E
Rest1
****(f)
F
Trpc2
****(f)
G
Gabrb2
**(f)
H
Kcnk5
***(f)
I
Trpm8
*(f)
J
Mlf1
*(f)
K
Calcrl
*(f)
Normalized Counts
R
N

Fig 5-2

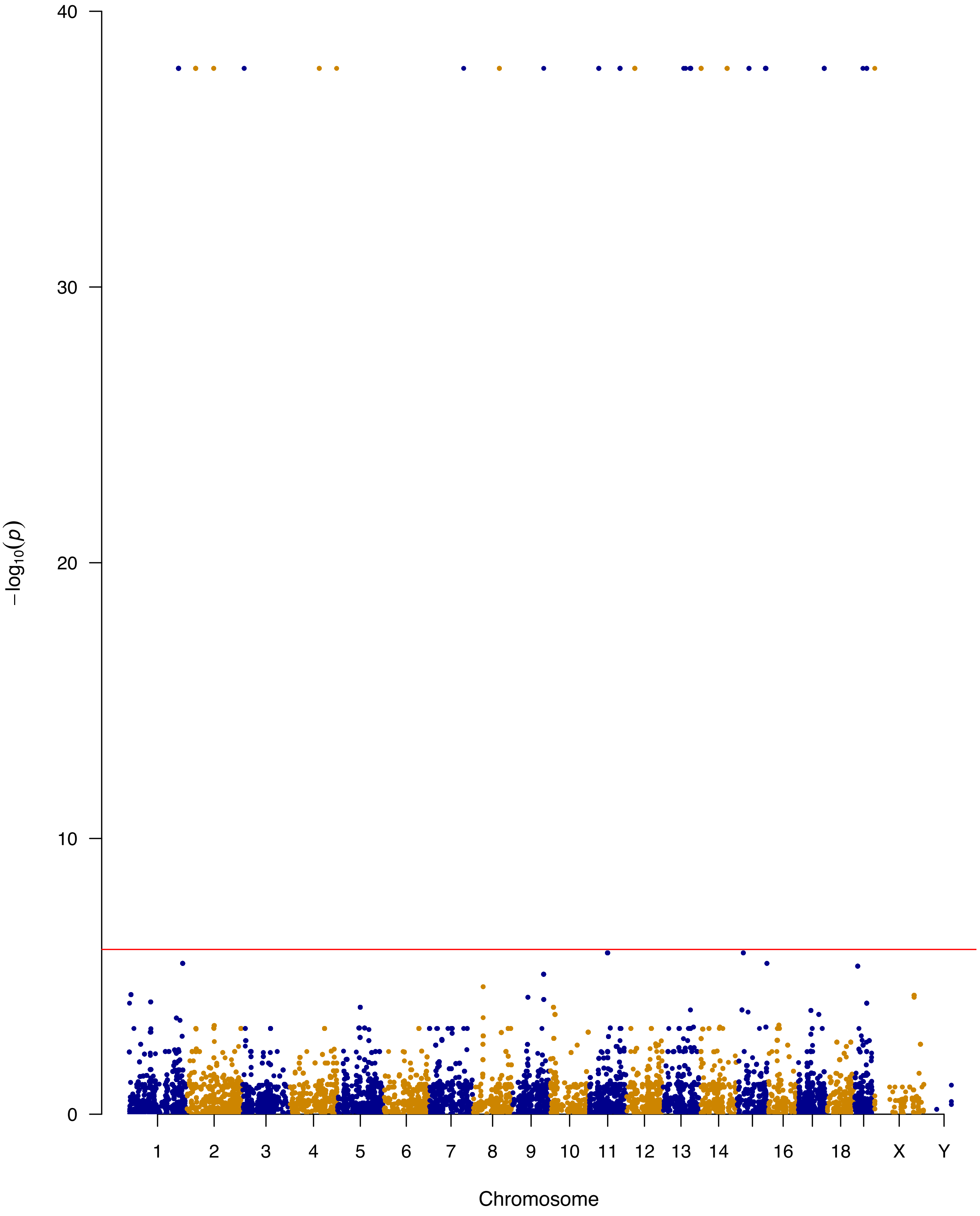

40
30
20
10
0
$-\log_{10}(p)$
1
2
3
4
5
6
7
8
9
10
11
12
13
14
16
18
X
Y
Chromosome

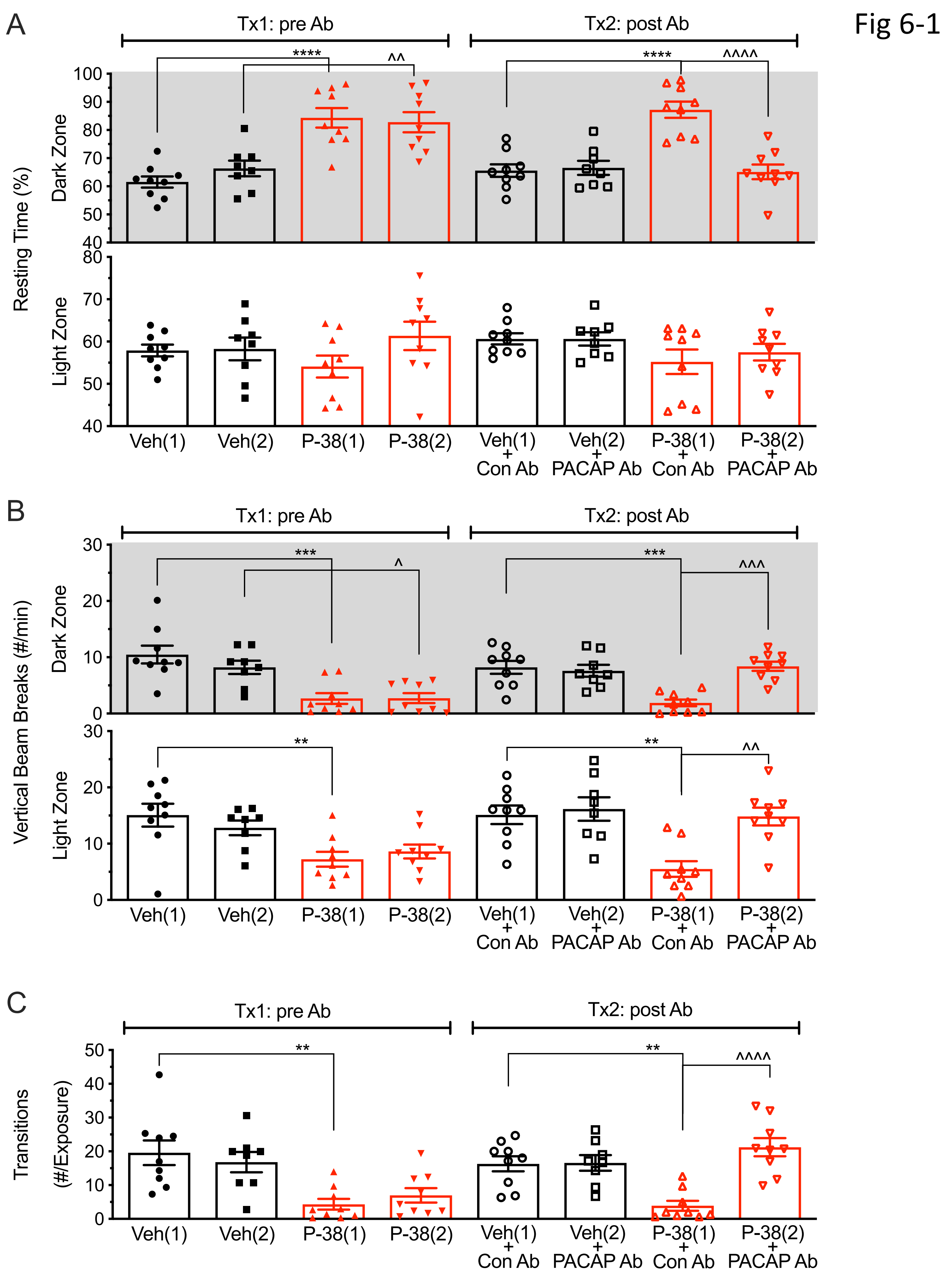
Fig 6-1
A
Tx1: pre Ab
Tx2: post Ab
Resting Time (%)
Dark Zone
Light Zone
****
^^
****
^^^^
Veh(1)
Veh(2)
P-38(1)
P-38(2)
Veh(1) + Con Ab
Veh(2) + PACAP Ab
P-38(1) + Con Ab
P-38(2) + PACAP Ab
B
Tx1: pre Ab
Tx2: post Ab
Vertical Beam Breaks (#/min)
Dark Zone
Light Zone
***
^
***
^^^
**
**
^^
Veh(1)
Veh(2)
P-38(1)
P-38(2)
Veh(1) + Con Ab
Veh(2) + PACAP Ab
P-38(1) + Con Ab
P-38(2) + PACAP Ab
C
Tx1: pre Ab
Tx2: post Ab
Transitions (#/Exposure)
**
**
^^^^
Veh(1)
Veh(2)
P-38(1)
P-38(2)
Veh(1) + Con Ab
Veh(2) + PACAP Ab
P-38(1) + Con Ab
P-38(2) + PACAP Ab

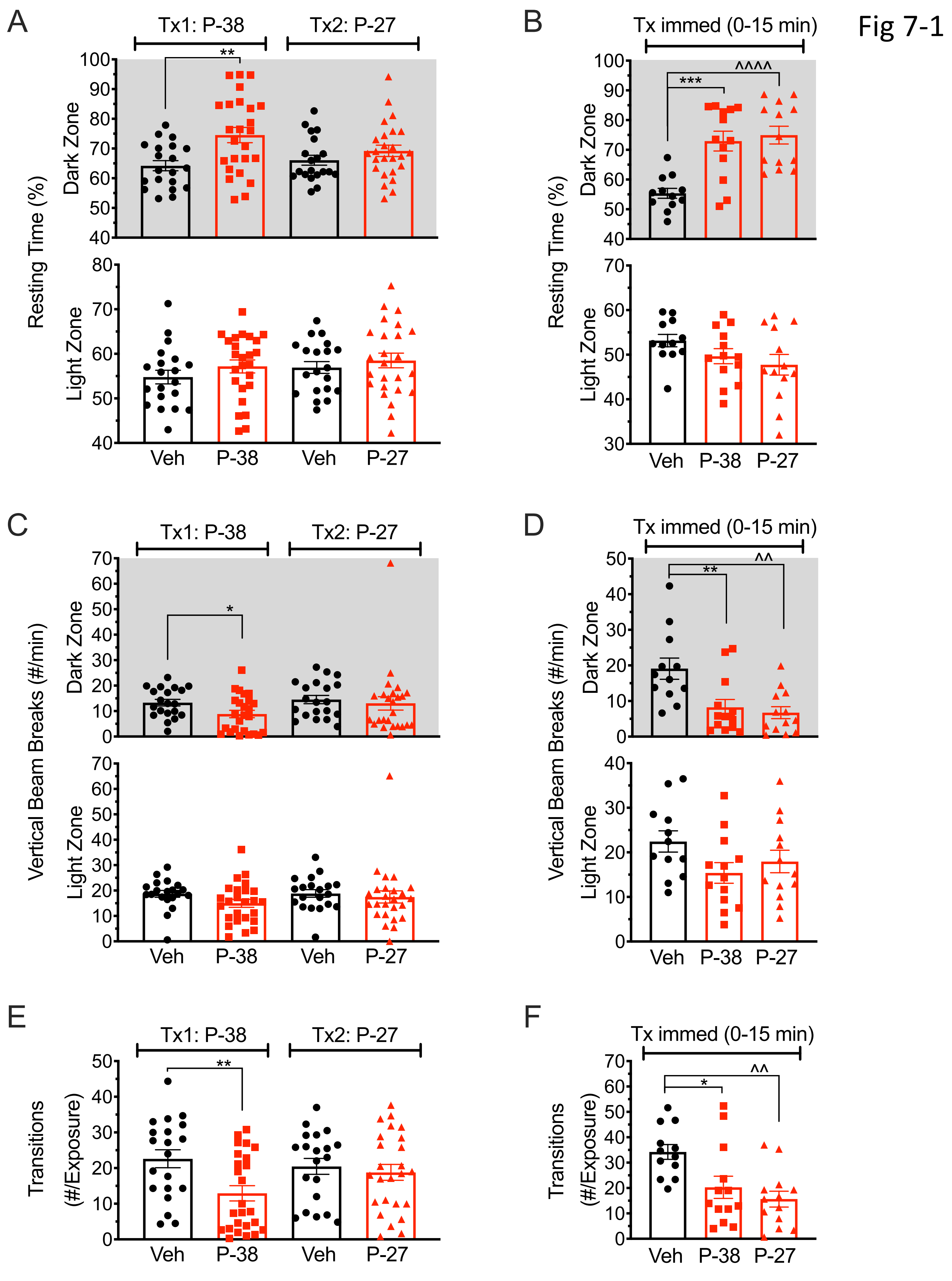

Fig 7-1
A
Tx1: P-38
Tx2: P-27
**
Resting Time (%)
Dark Zone
Light Zone
100
90
80
70
60
50
40
80
70
60
50
40
Veh
P-38
Veh
P-27
B
Tx immed (0-15 min)
***
^^^^
Resting Time (%)
Dark Zone
Light Zone
100
90
80
70
60
50
40
70
60
50
40
30
Veh
P-38
P-27
C
Tx1: P-38
Tx2: P-27
*
Vertical Beam Breaks (#/min)
Dark Zone
Light Zone
70
60
50
40
30
20
10
0
Veh
P-38
Veh
P-27
D
Tx immed (0-15 min)
**
^^
Vertical Beam Breaks (#/min)
Dark Zone
Light Zone
50
40
30
20
10
0
40
30
20
10
0
Veh
P-38
P-27
E
Tx1: P-38
Tx2: P-27
**
Transitions
(#/Exposure)
50
40
30
20
10
0
Veh
P-38
Veh
P-27
F
Tx immed (0-15 min)
*
^^
Transitions
(#/Exposure)
70
60
50
40
30
20
10
0
Veh
P-38
P-27

Fig 7-2

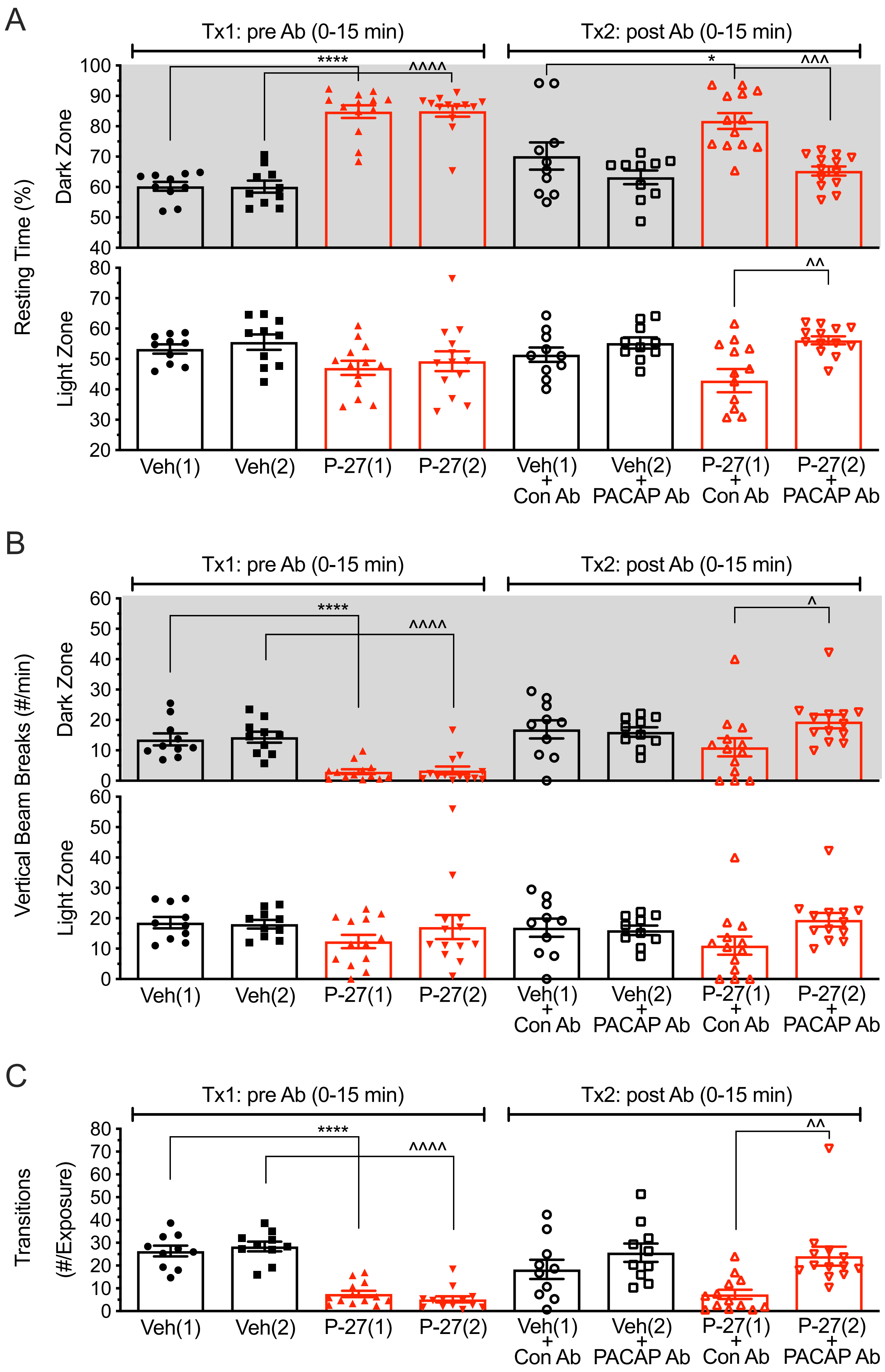
A
Tx1: pre Ab (0-15 min)
Tx2: post Ab (0-15 min)
Resting Time (%)
Dark Zone
Light Zone
****
^^^^
*
^^^
^^
Veh(1)
Veh(2)
P-27(1)
P-27(2)
Veh(1) + Con Ab
Veh(2) + PACAP Ab
P-27(1) + Con Ab
P-27(2) + PACAP Ab
B
Tx1: pre Ab (0-15 min)
Tx2: post Ab (0-15 min)
Vertical Beam Breaks (#/min)
Dark Zone
Light Zone
****
^^^^
^
Veh(1)
Veh(2)
P-27(1)
P-27(2)
Veh(1) + Con Ab
Veh(2) + PACAP Ab
P-27(1) + Con Ab
P-27(2) + PACAP Ab
C
Tx1: pre Ab (0-15 min)
Tx2: post Ab (0-15 min)
Transitions (#/Exposure)
****
^^^^
^^
Veh(1)
Veh(2)
P-27(1)
P-27(2)
Veh(1) + Con Ab
Veh(2) + PACAP Ab
P-27(1) + Con Ab
P-27(2) + PACAP Ab

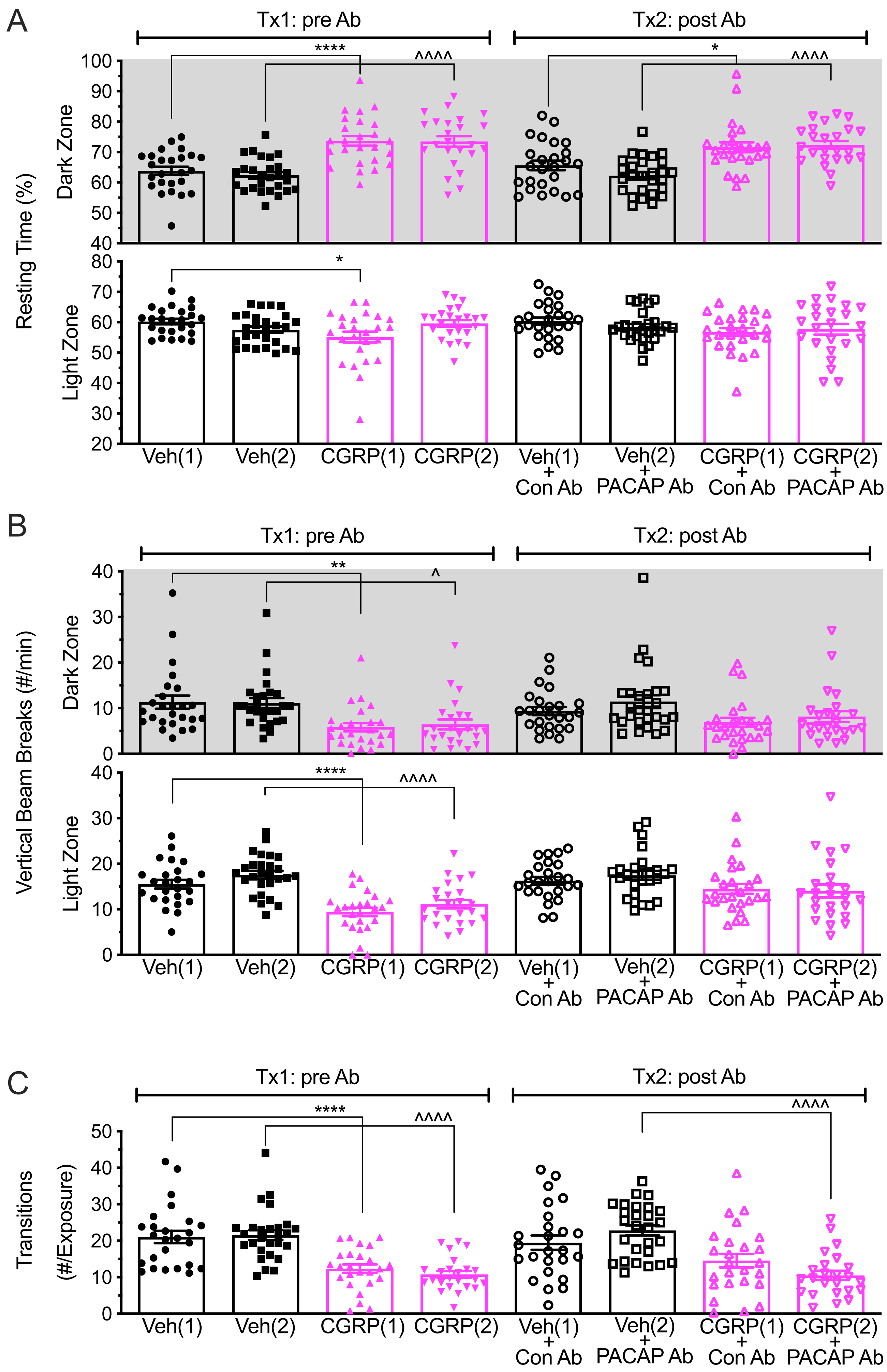

A
Tx1: pre Ab
Tx2: post Ab
Resting Time (%)
Dark Zone
Light Zone
****
^^^^
*
^^^^
*
B
Tx1: pre Ab
Tx2: post Ab
Vertical Beam Breaks (#/min)
Dark Zone
Light Zone
**
^
****
^^^^
C
Tx1: pre Ab
Tx2: post Ab
Transitions
(#/Exposure)
****
^^^^
^^^^
Veh(1)
Veh(2)
CGRP(1)
CGRP(2)
Veh(1) + Con Ab
Veh(2) + PACAP Ab
CGRP(1) + Con Ab
CGRP(2) + PACAP Ab

Fig 8-1

Fig 9-1

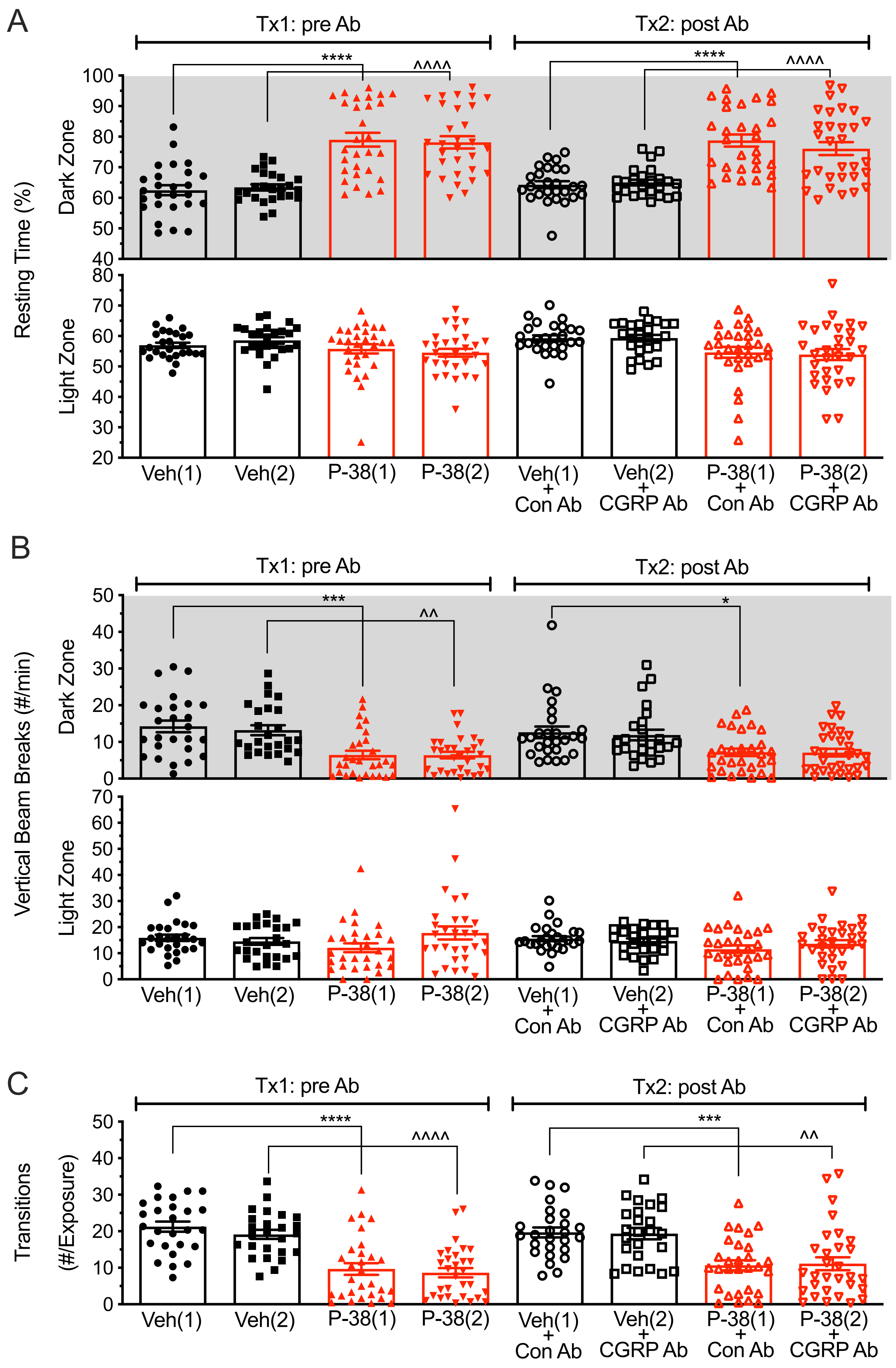
A
Tx1: pre Ab
Tx2: post Ab
****
^^^^
Resting Time (%)
Dark Zone
Light Zone
Veh(1)
Veh(2)
P-38(1)
P-38(2)
Veh(1) + Con Ab
Veh(2) + CGRP Ab
P-38(1) + Con Ab
P-38(2) + CGRP Ab
B
***
^^
*
Vertical Beam Breaks (#/min)
C
^^^^
^^
Transitions (#/Exposure)

| Figure # | Analysis | Statistics (symbol on Figure) |
|---|---|---|
| Figure 3-1A<br>Tx1<br>Dark Zone | One-way ANOVA for treatment<br>Bonferroni's multiple comparisons<br>- Veh vs. P-38 R<br>- Veh vs. P-38 N<br>- P-38 R vs. P-38 N | $F_{(2,55)}$=48.87, $p$<0.0001<br><br>$p$<0.0001 (****)<br>$p$>0.9999<br>$p$<0.0001 (^^^^) |
| Figure 3-1A<br>Tx2<br>Dark Zone | One-way ANOVA for treatment<br>Bonferroni's multiple comparisons<br>- Veh vs. P-38 R<br>- Veh vs. P-38 N<br>- P-38 R vs. P-38 N | $F_{(2,55)}$=15.40, $p$<0.0001<br><br>$p$<0.0001 (****)<br>$p$=0.9180<br>$p$=0.0002 (^^^) |
| Figure 3-1A<br>Tx1<br>Light Zone | One-way ANOVA for treatment<br>Bonferroni's multiple comparisons<br>- Veh vs. P-38 R<br>- Veh vs. P-38 N<br>- P-38 R vs. P-38 N | $F_{(2,55)}$=0.04101, $p$=.9599<br><br>$p$>0.9999<br>$p$>0.9999<br>$p$>0.9999 |
| Figure 3-1A<br>Tx2<br>Light zone | One-way ANOVA for treatment<br>Bonferroni's multiple comparisons<br>- Veh vs. P-38 R<br>- Veh vs. P-38 N<br>- P-38 R vs. P-38 N | $F_{(2,55)}$=2.068, $p$=0.1361<br><br>$p$=0.1593<br>$p$>0.9999<br>$p$=0.4600 |
| Figure 3-1B<br>Tx1<br>Dark Zone | One-way ANOVA for treatment<br>Bonferroni's multiple comparisons<br>- Veh vs. P-38 R<br>- Veh vs. P-38 N<br>- P-38 R vs. P-38 N | $F_{(2,55)}$=17.75, $p$<0.0001<br><br>$p$<0.0001 (****)<br>$p$=0.8611<br>$p$<0.0001 (^^^^) |
| Figure 3-1B<br>Tx2<br>Dark Zone | One-way ANOVA for treatment<br>Bonferroni's multiple comparisons<br>- Veh vs. P-38 R<br>- Veh vs. P-38 N<br>- P-38 R vs. P-38 N | $F_{(2,55)}$=14.31, $p$<0.0001<br><br>$p$<0.0001 (****)<br>$p$=0.2923<br>$p$=0.0018 (^^) |
| Figure 3-1B<br>Tx1<br>Light Zone | One-way ANOVA for treatment<br>Bonferroni's multiple comparisons<br>- Veh vs. P-38 R<br>- Veh vs. P-38 N<br>- P-38 R vs. P-38 N | $F_{(2,55)}$=10.36, $p$=0.0002<br><br>$P$=0.0002 (***)<br>$p$>0.9999<br>$p$=0.0030 (^^) |
| Figure 3-1B<br>Tx2<br>Light Zone | One-way ANOVA for treatment<br>Bonferroni's multiple comparisons<br>- Veh vs. P-38 R<br>- Veh vs. P-38 N<br>- P-38 R vs. P-38 N | $F_{(2,55)}$=10.92, $p$=0.0001<br><br>$p$<0.0001 (****)<br>$p$=0.5229<br>$p$=0.0059 (^^) |
| Figure 3-1C<br>Tx1 | One-way ANOVA for treatment<br>Bonferroni's multiple comparisons<br>- Veh vs. P-38 R<br>- Veh vs. P-38 N<br>- P-38 R vs. P-38 N | $F_{(2,55)}$=34.25, $p$<0.0001<br><br>$p$<0.0001 (****)<br>$p$>0.9999<br>$p$<0.0001 (^^^^) |
| Figure 3-1C<br>Tx2 | One-way ANOVA for treatment<br>Bonferroni's multiple comparisons<br>- Veh vs. P-38 R | $F_{(2,55)}$=12.51, $p$<0.0001<br><br>$p$=0.0001 (***) |

| | | |
|---|---|---|
| | - Veh vs. P-38 N<br>- P-38 R vs. P-38 N | $p>0.9999$<br>$p=0.0003$ (^^^) |
| Figure 4-1A<br>Tx1<br>Dark Zone | One-way ANOVA for treatment<br>Bonferroni's multiple comparisons<br>- RxR Veh vs. P-38<br>- RxN Veh vs. P-38<br>- NxN Veh vs. P-38<br>- RxR P-38 vs. NxN P-38<br>- RxN P-38 vs. NxN P-38 | $F_{(5,223)}=17.03$, $p<0.0001$<br><br>$p<0.0001$ (****)<br>$p<0.0001$ (^^^^)<br>$p>0.9999$<br>$p<0.0001$ (####)<br>$p=0.0001$ (&&&) |
| Figure 4-1A<br>Tx2<br>Dark Zone | One-way ANOVA for treatment<br>Bonferroni's multiple comparisons<br>- RxR Veh vs. P-38<br>- RxN Veh vs. P-38<br>- NxN Veh vs. P-38<br>- RxR P-38 vs. NxN P-38<br>- RxN P-38 vs. NxN P-38 | $F_{(5,206)}=21.93$, $p<0.0001$<br><br>$p<0.0001$ (****)<br>$p<0.0001$ (^^^^)<br>$p>0.9999$<br>$p<0.0001$ (####)<br>$p<0.0001$ (&&&&) |
| Figure 4-1A<br>Tx1<br>Light Zone | One-way ANOVA for treatment<br>Bonferroni's multiple comparisons<br>- RxR Veh vs. P-38<br>- RxN Veh vs. P-38<br>- NxN Veh vs. P-38<br>- RxR P-38 vs. NxN P-38<br>- RxN P-38 vs. NxN P-38 | $F_{(5,223)}=0.3552$, $p=0.8786$<br><br>$p>0.9999$<br>$p>0.9999$<br>$p>0.9999$<br>$p>0.9999$<br>$p>0.9999$ |
| Figure 4-1A<br>Tx2<br>Light Zone | One-way ANOVA for treatment<br>Bonferroni's multiple comparisons<br>- RxR Veh vs. P-38<br>- RxN Veh vs. P-38<br>- NxN Veh vs. P-38<br>- RxR P-38 vs. NxN P-38<br>- RxN P-38 vs. NxN P-38 | $F_{(5,206)}=4.257$, $p=0.0011$<br><br>$p>0.9999$<br>$p=0.0566$<br>$p>0.9999$<br>$p>0.9999$<br>$p=0.4903$ |
| Figure 4-1B<br>Tx1 | One-way ANOVA for treatment<br>Bonferroni's multiple comparisons<br>- RxR Veh vs. P-38<br>- RxN Veh vs. P-38<br>- NxN Veh vs. P-38<br>- RxR P-38 vs. NxN P-38<br>- RxN P-38 vs. NxN P-38 | $F_{(5,223)}=24.31$, $p<0.0001$<br><br>$p<0.0001$ (****)<br>$p<0.0001$ (^^^^)<br>$p>0.9999$<br>$p<0.0001$ (####)<br>$p<0.0001$ (&&&&) |
| Figure 4-1B<br>Tx2 | One-way ANOVA for treatment<br>Bonferroni's multiple comparisons<br>- RxR Veh vs. P-38<br>- RxN Veh vs. P-38<br>- NxN Veh vs. P-38<br>- RxR P-38 vs. NxN P-38<br>- RxN P-38 vs. NxN P-38 | $F_{(5,206)}=15.94$, $p<0.0001$<br><br>$p<0.0001$ (****)<br>$p<0.0001$ (^^^^)<br>$p>0.9999$<br>$p<0.0001$ (####)<br>$p<0.0001$ (&&&&) |
| Figure 5-1A | Wald test<br>Total<br>Male<br>Female | <br>FDR=0.1482<br>FDR=0.0275 (*)<br>FDR=0.5861 |
| Figure 5-1B | Wald test<br>Total | <br>FDR=0.1237 |

| | | |
|---|---|---|
| | Male<br>Female | FDR=0.0364 (*)<br>FDR=0.6718 |
| Figure 5-1C | Wald test<br>Total<br>Male<br>Female | <br>FDR=NA<br>FDR<0.0001 (****)<br>FDR=NA |
| Figure 5-1D | Wald test<br>Total<br>Male<br>Female | <br>FDR=NA<br>FDR=0.0081 (**)<br>FDR=NA |
| Figure 5-1E | Wald test<br>Total<br>Male<br>Female | <br>FDR=0.2587<br>FDR=0.9999<br>FDR<0.0001 (****) |
| Figure 5-1F | Wald test<br>Total<br>Male<br>Female | <br>FDR=0.9821<br>FDR=0.0869<br>FDR<0.0001 (****) |
| Figure 5-1G | Wald test<br>Total<br>Male<br>Female | <br>FDR=0.5267<br>FDR=0.9999<br>FDR=0.0071 (**) |
| Figure 5-1H | Wald test<br>Total<br>Male<br>Female | <br>FDR=0.0674<br>FDR=0.7625<br>FDR=0.0002 (***) |
| Figure 5-1I | Wald test<br>Total<br>Male<br>Female | <br>FDR=0.0910<br>FDR=0.7495<br>FDR=0.0331 (*) |
| Figure 5-1J | Wald test<br>Total<br>Male<br>Female | <br>FDR=0.5418<br>FDR=0.9999<br>FDR=0.0188 (*) |
| Figure 5-1K | Wald test<br>Total<br>Male<br>Female | <br>FDR=0.7071<br>FDR=0.9999<br>FDR=0.0163 (*) |
| Figure 6-1A<br>Tx1<br>Dark Zone | One-way ANOVA for treatment<br>Bonferroni's multiple comparisons<br>- Veh(1) vs. Veh(2)<br>- Veh(1) vs. P-38(1)<br>- Veh(1) vs. P-38(2)<br>- Veh(2) vs. P-38(1)<br>- Veh(2) vs. P-38(2)<br>- P-38(1) vs. P-38(2) | $F_{(3,31)}$=14.56, $p$<0.0001<br><br>$p$>0.9999<br>$p$<0.0001 (****)<br>$p$=0.0001<br>$p$=0.0016<br>$p$=0.0042 (^^)<br>$p$>0.9999 |
| Figure 6-1A<br>Tx2<br>Dark Zone | One-way ANOVA for treatment<br>Bonferroni's multiple comparisons<br>- Veh(1) + Con Ab vs. Veh(2) + PACAP Ab<br>- Veh(1) + Con Ab vs. P-38(1) + Con Ab | $F_{(3,31)}$=17.82, $p$<0.0001<br><br>$p$>0.9999<br>$p$<0.0001 (****) |

| | | |
|---|---|---|
| | - Veh(1) + Con Ab vs. P-38(2) + PACAP Ab<br>- Veh(2) + PACAP Ab vs. P-38(1) + Con Ab<br>- Veh(2) + PACAP Ab vs. P-38(2) + PACAP Ab<br>- P-38(1) + Con Ab vs. P-38(2) + PACAP Ab | $p>0.9999$<br>$p<0.0001$<br>$p>0.9999$<br>$p<0.0001$ (^^^^) |
| Figure 6-1A<br>Tx1<br>Light Zone | One-way ANOVA for treatment<br>Bonferroni's multiple comparisons<br>- Veh(1) vs. Veh(2)<br>- Veh(1) vs. P-38(1)<br>- Veh(1) vs. P-38(2)<br>- Veh(2) vs. P-38(1)<br>- Veh(2) vs. P-38(2)<br>- P-38(1) vs. P-38(2) | $F_{(3,31)}=1.337$, $p=0.2802$<br><br>$p>0.9999$<br>$p>0.9999$<br>$p>0.9999$<br>$p>0.9999$<br>$p>0.9999$<br>$p=0.3287$ |
| Figure 6-1A<br>Tx2<br>Light Zone | One-way ANOVA for treatment<br>Bonferroni's multiple comparisons<br>- Veh(1) + Con Ab vs. Veh(2) + PACAP Ab<br>- Veh(1) + Con Ab vs. P-38(1) + Con Ab<br>- Veh(1) + Con Ab vs. P-38(2) + PACAP Ab<br>- Veh(2) + PACAP Ab vs. P-38(1) + Con Ab<br>- Veh(2) + PACAP Ab vs. P-38(2) + PACAP Ab<br>- P-38(1) + Con Ab vs. P-38(2) + PACAP Ab | $F_{(3,31)}=1.648$, $p=0.1985$<br><br>$p>0.9999$<br>$p=0.4122$<br>$p>0.9999$<br>$p=0.4627$<br>$p>0.9999$<br>$p>0.9999$ |
| Figure 6-1B<br>Tx1<br>Dark Zone | One-way ANOVA for treatment<br>Bonferroni's multiple comparisons<br>- Veh(1) vs. Veh(2)<br>- Veh(1) vs. P-38(1)<br>- Veh(1) vs. P-38(2)<br>- Veh(2) vs. P-38(1)<br>- Veh(2) vs. P-38(2)<br>- P-38(1) vs. P-38(2) | $F_{(3,31)}=11.38$, $p<0.0001$<br><br>$p>0.9999$<br>$p=0.0003$ (***)<br>$p=0.0003$<br>$p=0.0161$<br>$p=0.0176$ (^)<br>$p>0.9999$ |
| Figure 6-1B<br>Tx2<br>Dark Zone | One-way ANOVA for treatment<br>Bonferroni's multiple comparisons<br>- Veh(1) + Con Ab vs. Veh(2) + PACAP Ab<br>- Veh(1) + Con Ab vs. P-38(1) + Con Ab<br>- Veh(1) + Con Ab vs. P-38(2) + PACAP Ab<br>- Veh(2) + PACAP Ab vs. P-38(1) + Con Ab<br>- Veh(2) + PACAP Ab vs. P-38(2) + PACAP Ab<br>- P-38(1) + Con Ab vs. P-38(2) + PACAP Ab | $F_{(3,31)}=11.65$, $p<0.0001$<br><br>$p>0.9999$<br>$p=0.0002$ (***)<br>$p>0.9999$<br>$p=0.0009$<br>$p>0.9999$<br>$p=0.0001$ (^^^) |
| Figure 6-1B<br>Tx1<br>Light Zone | One-way ANOVA for treatment<br>Bonferroni's multiple comparisons<br>- Veh(1) vs. Veh(2)<br>- Veh(1) vs. P-38(1)<br>- Veh(1) vs. P-38(2)<br>- Veh(2) vs. P-38(1)<br>- Veh(2) vs. P-38(2)<br>- P-38(1) vs. P-38(2) | $F_{(3,31)}=5.827$, $p=0.0028$<br><br>$p>0.9999$<br>$p=0.0051$ (**)<br>$p=0.0283$<br>$p=0.0953$<br>$p=0.3852$<br>$p>0.9999$ |
| Figure 6-1B<br>Tx2<br>Light Zone | One-way ANOVA for treatment<br>Bonferroni's multiple comparisons<br>- Veh(1) + Con Ab vs. Veh(2) + PACAP Ab<br>- Veh(1) + Con Ab vs. P-38(1) + Con Ab<br>- Veh(1) + Con Ab vs. P-38(2) + PACAP Ab<br>- Veh(2) + PACAP Ab vs. P-38(1) + Con Ab | $F_{(3,31)}=8.961$, $p=0.0002$<br><br>$p>0.9999$<br>$p=0.0015$ (**)<br>$p>0.9999$<br>$p=0.0006$ |

| | | |
|---|---|---|
| | - Veh(2) + PACAP Ab vs. P-38(2) + PACAP Ab<br>- P-38(1) + Con Ab vs. P-38(2) + PACAP Ab | *p*>0.9999<br>*p*=0.0022 (^^) |
| Figure 6-1C<br>Tx1 | One-way ANOVA for treatment<br>Bonferroni's multiple comparisons<br>- Veh(1) vs. Veh(2)<br>- Veh(1) vs. P-38(1)<br>- Veh(1) vs. P-38(2)<br>- Veh(2) vs. P-38(1)<br>- Veh(2) vs. P-38(2)<br>- P-38(1) vs. P-38(2) | $F_{(3,31)}$=7.674, *p*=0.0006<br><br>*p*>0.9999<br>*p*=0.0018 (**)<br>*p*=0.0124<br>*p*=0.0177<br>*p*=0.0970<br>*p*>0.9999 |
| Figure 6-1C<br>Tx2 | One-way ANOVA for treatment<br>Bonferroni's multiple comparisons<br>- Veh(1) + Con Ab vs. Veh(2) + PACAP Ab<br>- Veh(1) + Con Ab vs. P-38(1) + Con Ab<br>- Veh(1) + Con Ab vs. P-38(2) + PACAP Ab<br>- Veh(2) + PACAP Ab vs. P-38(1) + Con Ab<br>- Veh(2) + PACAP Ab vs. P-38(2) + PACAP Ab<br>- P-38(1) + Con Ab vs. P-38(2) + PACAP Ab | $F_{(3,31)}$=11.45, *p*<0.0001<br><br>*p*>0.9999<br>*p*=0.0021 (**)<br>*p*=0.7406<br>*p*=0.0024<br>*p*=0.9388<br>*p*<0.0001 (^^^^) |
| Figure 7-1A<br>Tx1<br>Dark Zone | Unpaired t-test, Two-tailed<br>- Veh vs. P-38 | t=3.182, df=43<br>*p*=0.0027 (**) |
| Figure 7-1A<br>Tx2<br>Dark Zone | Unpaired t-test, Two-tailed<br>- Veh vs. P-27 | t=1.211, df=43<br>*p*=0.2325 |
| Figure 7-1A<br>Tx1<br>Light Zone | Unpaired t-test, Two-tailed<br>- Veh vs. P-38 | t=1.118, df=43<br>*p*=0.2698 |
| Figure 7-1A<br>Tx2<br>Light Zone | Unpaired t-test, Two-tailed<br>- Veh vs. P-27 | t=0.7126, df=43<br>*p*=0.4800 |
| Figure 7-1B<br>Tx1<br>Dark Zone | Unpaired t-test, Two-tailed<br>- Veh vs. P-38 | t=2.267, df=43<br>*p*=0.0285 (*) |
| Figure 7-1B<br>Tx2<br>Dark Zone | Unpaired t-test, Two-tailed<br>- Veh vs. P-27 | t=0.4520, df=43<br>*p*=0.6536 |
| Figure 7-1B<br>Tx1<br>Light Zone | Unpaired t-test, Two-tailed<br>- Veh vs. P-38 | t=1.736, df=43<br>*p*=0.0897 |
| Figure 7-1B<br>Tx2<br>Light Zone | Unpaired t-test, Two-tailed<br>- Veh vs. P-27 | t=0.4561, df=43<br>*p*=0.6506 |
| Figure 7-1C<br>Tx1 | Unpaired t-test, Two-tailed<br>- Veh vs. P-38 | t=2.949, df=43<br>*p*=0.0051 (**) |
| Figure 7-1C<br>Tx2 | Unpaired t-test, Two-tailed<br>- Veh vs. P-27 | t=0.5254, df=43<br>*p*=0.6013 |
| Figure 7-1D<br>Dark Zone | One-way ANOVA for treatment<br>Bonferroni's multiple comparisons<br>- Veh vs. P-38<br>- Veh vs. P-27 | $F_{(2, 35)}$=14.27, *p*<0.0001<br><br>*p*=0.0003 (***)<br>*p*<0.0001 (^^^^) |

| | | |
|---|---|---|
| | - P-38 vs. P-27 | *p*>0.9999 |
| Figure 7-1D<br>Light Zone | One-way ANOVA for treatment<br>Bonferroni's multiple comparisons<br>- Veh vs. P-38<br>- Veh vs. P-27<br>- P-38 vs. P-27 | $F_{(2, 35)}$=2.103, *p*=0.1372<br><br>*p*=0.5950<br>*p*=0.1504<br>*p*>0.9999 |
| Figure 7-1E<br>Dark Zone | One-way ANOVA for treatment<br>Bonferroni's multiple comparisons<br>- Veh vs. P-38<br>- Veh vs. P-27<br>- P-38 vs. P-27 | $F_{(2, 35)}$=8.247, *p*=0.0012<br><br>*p*=0.0068 (**)<br>*p*=0.0020 (^^)<br>*p*>0.9999 |
| Figure 7-1E<br>Light Zone | One-way ANOVA for treatment<br>Bonferroni's multiple comparisons<br>- Veh vs. P-38<br>- Veh vs. P-27<br>- P-38 vs. P-27 | $F_{(2, 35)}$=2.156, *p*=0.1309<br><br>*p*=0.1414<br>*p*=0.5959<br>*p*>0.9999 |
| Figure 7-1F | One-way ANOVA for treatment<br>Bonferroni's multiple comparisons<br>- Veh vs. P-38<br>- Veh vs. P-27<br>- P-38 vs. P-27 | $F_{(2, 35)}$=7.174, *p*=0.0024<br><br>*p*=0.0282 (*)<br>*p*=0.0025 (^^)<br>*p*>0.9999 |
| Figure 7-2A<br>Tx1<br>Dark Zone | One-way ANOVA for treatment<br>Bonferroni's multiple comparisons<br>- Veh(1) vs. Veh(2)<br>- Veh(1) vs. P-27(1)<br>- Veh(1) vs. P-27(2)<br>- Veh(2) vs. P-27(1)<br>- Veh(2) vs. P-27(2)<br>- P-27(1) vs. P-27(2) | $F_{(3,42)}$=55.59, *p*<0.0001<br><br>*p*>0.9999<br>*p*<0.0001 (****)<br>*p*<0.0001<br>*p*<0.0001<br>*p*<0.0001 (^^^^)<br>*p*>0.9999 |
| Figure 7-2A<br>Tx2<br>Dark Zone | One-way ANOVA for treatment<br>Bonferroni's multiple comparisons<br>- Veh(1) + Con Ab vs. Veh(2) + PACAP Ab<br>- Veh(1) + Con Ab vs. P-27(1) + Con Ab<br>- Veh(1) + Con Ab vs. P-27(2) + PACAP Ab<br>- Veh(2) + PACAP Ab vs. P-27(1) + Con Ab<br>- Veh(2) + PACAP Ab vs. P-27(2) + PACAP Ab<br>- P-27(1) + Con Ab vs. P-27(2) + PACAP Ab | $F_{(3,42)}$=9.644, *p*<0.0001<br><br>*P*=0.6127<br>*p*=0.0322 (*)<br>*p*>0.9999<br>*p*=0.0002<br>*p*>0.9999<br>*p*=0.0003 (^^^) |
| Figure 7-2A<br>Tx1<br>Light Zone | One-way ANOVA for treatment<br>Bonferroni's multiple comparisons<br>- Veh(1) vs. Veh(2)<br>- Veh(1) vs. P-27(1)<br>- Veh(1) vs. P-27(2)<br>- Veh(2) vs. P-27(1)<br>- Veh(2) vs. P-27(2)<br>- P-27(1) vs. P-27(2) | $F_{(3,42)}$=2.129, *p*=0.1109<br><br>*p*>0.9999<br>*p*=0.6087<br>*p*>0.9999<br>*p*=0.1657<br>*p*=0.5917<br>*p*>0.9999 |
| Figure 7-2A<br>Tx2<br>Light Zone | One-way ANOVA for treatment<br>Bonferroni's multiple comparisons<br>- Veh(1) + Con Ab vs. Veh(2) + PACAP Ab<br>- Veh(1) + Con Ab vs. P-27(1) + Con Ab<br>- Veh(1) + Con Ab vs. P-27(2) + PACAP Ab | $F_{(3,42)}$=5.760, *p*=0.0022<br><br>*P*>0.9999<br>*p*=0.1706<br>*p*>0.9999 |

| | | |
|---|---|---|
| | - Veh(2) + PACAP Ab vs. P-27(1) + Con Ab<br>- Veh(2) + PACAP Ab vs. P-27(2) + PACAP Ab<br>- P-27(1) + Con Ab vs. P-27(2) + PACAP Ab | *p*=0.0121<br>*p*>0.9999<br>*p*=0.0029 (^^) |
| Figure 7-2B<br>Tx1<br>Dark Zone | One-way ANOVA for treatment<br>Bonferroni's multiple comparisons<br>- Veh(1) vs. Veh(2)<br>- Veh(1) vs. P-27(1)<br>- Veh(1) vs. P-27(2)<br>- Veh(2) vs. P-27(1)<br>- Veh(2) vs. P-27(2)<br>- P-27(1) vs. P-27(2) | $F_{(3,42)}$=18.44, *p*<0.0001<br><br>*p*>0.9999<br>*p*<0.0001 (****)<br>*p*<0.0001<br>*p*<0.0001<br>*p*<0.0001 (^^^^)<br>*p*>0.9999 |
| Figure 7-2B<br>Tx2<br>Dark Zone | One-way ANOVA for treatment<br>Bonferroni's multiple comparisons<br>- Veh(1) + Con Ab vs. Veh(2) + PACAP Ab<br>- Veh(1) + Con Ab vs. P-27(1) + Con Ab<br>- Veh(1) + Con Ab vs. P-27(2) + PACAP Ab<br>- Veh(2) + PACAP Ab vs. P-27(1) + Con Ab<br>- Veh(2) + PACAP Ab vs. P-27(2) + PACAP Ab<br>- P-27(1) + Con Ab vs. P-27(2) + PACAP Ab | $F_{(3,42)}$=4.411, *p*=0.0087<br><br>*p*>0.9999<br>*p*=0.1975<br>*p*>0.9999<br>*p*=0.0392<br>*p*>0.9999<br>*p*=0.0120 (^) |
| Figure 7-2B<br>Tx1<br>Light Zone | One-way ANOVA for treatment<br>Bonferroni's multiple comparisons<br>- Veh(1) vs. Veh(2)<br>- Veh(1) vs. P-27(1)<br>- Veh(1) vs. P-27(2)<br>- Veh(2) vs. P-27(1)<br>- Veh(2) vs. P-27(2)<br>- P-27(1) vs. P-27(2) | $F_{(3,42)}$=1.118, *p*=0.3528<br><br>*p*>0.9999<br>*p*=0.7291<br>*p*>0.9999<br>*p*=0.9235<br>*p*>0.9999<br>*p*>0.9999 |
| Figure 7-2B<br>Tx2<br>Light Zone | One-way ANOVA for treatment<br>Bonferroni's multiple comparisons<br>- Veh(1) + Con Ab vs. Veh(2) + PACAP Ab<br>- Veh(1) + Con Ab vs. P-27(1) + Con Ab<br>- Veh(1) + Con Ab vs. P-27(2) + PACAP Ab<br>- Veh(2) + PACAP Ab vs. P-27(1) + Con Ab<br>- Veh(2) + PACAP Ab vs. P-27(2) + PACAP Ab<br>- P-27(1) + Con Ab vs. P-27(2) + PACAP Ab | $F_{(3,42)}$=4.411, *p*=0.0087<br><br>*p*>0.9999<br>*p*=0.6923<br>*p*>0.9999<br>*p*>0.9999<br>*p*>0.9999<br>*p*=0.1031 |
| Figure 7-2C<br>Tx1 | One-way ANOVA for treatment<br>Bonferroni's multiple comparisons<br>- Veh(1) vs. Veh(2)<br>- Veh(1) vs. P-27(1)<br>- Veh(1) vs. P-27(2)<br>- Veh(2) vs. P-27(1)<br>- Veh(2) vs. P-27(2)<br>- P-27(1) vs. P-27(2) | $F_{(3,42)}$=47.68, *p*<0.0001<br><br>*p*>0.9999<br>*p*<0.0001 (****)<br>*p*<0.0001<br>*p*<0.0001<br>*p*<0.0001 (^^^^)<br>*p*>0.9999 |
| Figure 7-2C<br>Tx2 | One-way ANOVA for treatment<br>Bonferroni's multiple comparisons<br>- Veh(1) + Con Ab vs. Veh(2) + PACAP Ab<br>- Veh(1) + Con Ab vs. P-27(1) + Con Ab<br>- Veh(1) + Con Ab vs. P-27(2) + PACAP Ab<br>- Veh(2) + PACAP Ab vs. P-27(1) + Con Ab<br>- Veh(2) + PACAP Ab vs. P-27(2) + PACAP Ab | $F_{(3,42)}$=5.515, *p*=0.0028<br><br>*p*>0.9999<br>*p*=0.2489<br>*p*>0.9999<br>*p*=0.0064<br>*p*>0.9999 |

| | | |
|---|---|---|
| | - P-27(1) + Con Ab vs. P-27(2) + PACAP Ab | $p$=0.0078 (^^) |
| Figure 8-1A<br>Tx1<br>Dark Zone | One-way ANOVA for treatment<br>Bonferroni's multiple comparisons<br>- Veh(1) vs. Veh(2)<br>- Veh(1) vs. CGRP(1)<br>- Veh(1) vs. CGRP(2)<br>- Veh(2) vs. CGRP(1)<br>- Veh(2) vs. CGRP(2)<br>- CGRP(1) vs. CGRP(2) | $F_{(3,97)}$=17.99, $p$<0.0001<br><br>$p$>0.9999<br>$p$<0.0001 (****)<br>$p$<0.0001<br>$p$<0.0001<br>$p$<0.0001 (^^^^)<br>$p$>0.9999 |
| Figure 8-1A<br>Tx2<br>Dark Zone | One-way ANOVA for treatment<br>Bonferroni's multiple comparisons<br>- Veh(1) + Con Ab vs. Veh(2) + PACAP Ab<br>- Veh(1) + Con Ab vs. CGRP(1) + Con Ab<br>- Veh(1) + Con Ab vs. CGRP(2) + PACAP Ab<br>- Veh(2) + PACAP Ab vs. CGRP(1) + Con Ab<br>- Veh(2) + PACAP Ab vs. CGRP(2) + PACAP Ab<br>- CGRP(1) + Con Ab vs. CGRP(2) + PACAP Ab | $F_{(3,97)}$=11.86, $p$<0.0001<br><br>$p$=0.5545<br>$p$=0.0225 (*)<br>$p$=0.0080<br>$p$<0.0001<br>$p$<0.0001 (^^^^)<br>$p$>0.9999 |
| Figure 8-1A<br>Tx1<br>Light zone | One-way ANOVA for treatment<br>Bonferroni's multiple comparisons<br>- Veh(1) vs. Veh(2)<br>- Veh(1) vs. CGRP(1)<br>- Veh(1) vs. CGRP(2)<br>- Veh(2) vs. CGRP(1)<br>- Veh(2) vs. CGRP(2)<br>- CGRP(1) vs. CGRP(2) | $F_{(3,97)}$=3.523, $p$=0.0179<br><br>$p$=0.7007<br>$p$=0.0250 (*)<br>$p$>0.9999<br>$p$=0.9747<br>$p$>0.9999<br>$p$=0.0712 |
| Figure 8-1A<br>Tx2<br>Light Zone | One-way ANOVA for treatment<br>Bonferroni's multiple comparisons<br>- Veh(1) + Con Ab vs. Veh(2) + PACAP Ab<br>- Veh(1) + Con Ab vs. CGRP(1) + Con Ab<br>- Veh(1) + Con Ab vs. CGRP(2) + PACAP Ab<br>- Veh(2) + PACAP Ab vs. CGRP(1) + Con Ab<br>- Veh(2) + PACAP Ab vs. CGRP(2) + PACAP Ab<br>- CGRP(1) + Con Ab vs. CGRP(2) + PACAP Ab | $F_{(3,97)}$=1.353, $p$=0.2617<br><br>$p$>0.9999<br>$p$=0.3334<br>$p$=0.9406<br>$p$>0.9999<br>$p$>0.9999<br>$p$>0.9999 |
| Figure 8-1B<br>Tx1<br>Dark Zone | One-way ANOVA for treatment<br>Bonferroni's multiple comparisons<br>- Veh(1) vs. Veh(2)<br>- Veh(1) vs. CGRP(1)<br>- Veh(1) vs. CGRP(2)<br>- Veh(2) vs. CGRP(1)<br>- Veh(2) vs. CGRP(2)<br>- CGRP(1) vs. CGRP(2) | $F_{(3,97)}$=6.697, $p$=0.0004<br><br>$p$>0.9999<br>$p$=0.0063 (**)<br>$p$=0.0221<br>$p$=0.0072<br>$p$=0.0255 (^)<br>$p$>0.9999 |
| Figure 8-1B<br>Tx2<br>Dark Zone | One-way ANOVA for treatment<br>Bonferroni's multiple comparisons<br>- Veh(1) vs. Veh(2)<br>- Veh(1) vs. CGRP(1)<br>- Veh(1) vs. CGRP(2)<br>- Veh(2) vs. CGRP(1)<br>- Veh(2) vs. CGRP(2)<br>- CGRP(1) vs. CGRP(2) | $F_{(3,97)}$=2.885, $p$=0.0396<br><br>$p$>0.9999<br>$p$=0.8235<br>$p$>0.9999<br>$p$=0.0351<br>$p$=0.2769<br>$p$>0.9999 |
| Figure 8-1B | One-way ANOVA for treatment | $F_{(3,97)}$=17.63, $p$<0.0001 |

| | | |
|---|---|---|
| Tx1<br>Light Zone | Bonferroni's multiple comparisons<br>- Veh(1) vs. Veh(2)<br>- Veh(1) vs. CGRP(1)<br>- Veh(1) vs. CGRP(2)<br>- Veh(2) vs. CGRP(1)<br>- Veh(2) vs. CGRP(2)<br>- CGRP(1) vs. CGRP(2) | <br>$p$=0.6507<br>$p$<0.0001 (****)<br>$p$=0.0066<br>$p$<0.0001<br>$p$<0.0001 (^^^^)<br>$p$>0.9999 |
| Figure 8-1B<br>Tx2<br>Light Zone | One-way ANOVA for treatment<br>Bonferroni's multiple comparisons<br>- Veh(1) vs. Veh(2)<br>- Veh(1) vs. CGRP(1)<br>- Veh(1) vs. CGRP(2)<br>- Veh(2) vs. CGRP(1)<br>- Veh(2) vs. CGRP(2)<br>- CGRP(1) vs. CGRP(2) | $F_{(3,97)}$=2.230, $p$=0.0896<br><br>$p$>0.9999<br>$p$>0.9999<br>$p$=0.9096<br>$p$=0.2982<br>$p$=0.1530<br>$p$>0.9999 |
| Figure 8-1C<br>Tx1 | One-way ANOVA for treatment<br>Bonferroni's multiple comparisons<br>- Veh(1) vs. Veh(2)<br>- Veh(1) vs. CGRP(1)<br>- Veh(1) vs. CGRP(2)<br>- Veh(2) vs. CGRP(1)<br>- Veh(2) vs. CGRP(2)<br>- CGRP(1) vs. CGRP(2) | $F_{(3,97)}$=18.14, $p$<0.0001<br><br>$p$>0.9999<br>$p$<0.0001 (****)<br>$p$<0.0001<br>$p$<0.0001<br>$p$<0.0001 (^^^^)<br>$p$>0.9999 |
| Figure 8-1C<br>Tx2 | One-way ANOVA for treatment<br>Bonferroni's multiple comparisons<br>- Veh(1) + Con Ab vs. Veh(2) + PACAP Ab<br>- Veh(1) + Con Ab vs. CGRP(1) + Con Ab<br>- Veh(1) + Con Ab vs. CGRP(2) + PACAP Ab<br>- Veh(2) + PACAP Ab vs. CGRP(1) + Con Ab<br>- Veh(2) + PACAP Ab vs. CGRP(2) + PACAP Ab<br>- CGRP(1) + Con Ab vs. CGRP(2) + PACAP Ab | $F_{(3,97)}$=10.81, $p$<0.0001<br><br>$p$=0.8976<br>$p$=0.2275<br>$p$=0.0016<br>$p$=0.0031<br>$p$<0.0001 (^^^^)<br>$p$=0.5542 |
| Figure 9-1A<br>Tx1<br>Dark Zone | One-way ANOVA for treatment<br>Bonferroni's multiple comparisons<br>- Veh(1) vs. Veh(2)<br>- Veh(1) vs. P-38(1)<br>- Veh(1) vs. P-38(2)<br>- Veh(2) vs. P-38(1)<br>- Veh(2) vs. P-38(2)<br>- P-38(1) vs. P-38(2) | $F_{(3,106)}$=23.49, $p$<0.0001<br><br>$p$>0.9999<br>$p$<0.0001 (****)<br>$p$<0.0001<br>$p$<0.0001<br>$p$<0.0001 (^^^^)<br>$p$>0.9999 |
| Figure 9-1A<br>Tx2<br>Dark Zone | One-way ANOVA for treatment<br>Bonferroni's multiple comparisons<br>- Veh(1) + Con Ab vs. Veh(2) + CGRP Ab<br>- Veh(1) + Con Ab vs. P-38(1) + Con Ab<br>- Veh(1) + Con Ab vs. P-38(2) + CGRP Ab<br>- Veh(2) + CGRP Ab vs. P-38(1) + Con Ab<br>- Veh(2) + CGRP Ab vs. P-38(2) + CGRP Ab<br>- P-38(1) + Con Ab vs. P-38(2) + CGRP Ab | $F_{(3,106)}$=19.32, $p$<0.0001<br><br>$p$>0.9999<br>$p$<0.0001 (****)<br>$p$<0.0001<br>$p$<0.0001<br>$p$<0.0001 (^^^^)<br>$p$>0.9999 |
| Figure 9-1A<br>Tx1<br>Light Zone | One-way ANOVA for treatment<br>Bonferroni's multiple comparisons<br>- Veh(1) vs. Veh(2) | $F_{(3,106)}$=1.839, $p$=0.1446<br><br>$p$>0.9999 |

| | | |
|---|---|---|
| | - Veh(1) vs. P-38(1)<br>- Veh(1) vs. P-38(2)<br>- Veh(2) vs. P-38(1)<br>- Veh(2) vs. P-38(2)<br>- P-38(1) vs. P-38(2) | $p>0.9999$<br>$p>0.9999$<br>$p=0.7705$<br>$p=0.1564$<br>$p>0.9999$ |
| Figure 9-1A<br>Tx2<br>Light Zone | One-way ANOVA for treatment<br>Bonferroni's multiple comparisons<br>- Veh(1) + Con Ab vs. Veh(2) + CGRP Ab<br>- Veh(1) + Con Ab vs. P-38(1) + Con Ab<br>- Veh(1) + Con Ab vs. P-38(2) + CGRP Ab<br>- Veh(2) + CGRP Ab vs. P-38(1) + Con Ab<br>- Veh(2) + CGRP Ab vs. P-38(2) + CGRP Ab<br>- P-38(1) + Con Ab vs. P-38(2) + CGRP Ab | $F_{(3,106)}=3.659$, $p=0.0148$<br><br>$p>0.9999$<br>$p=0.2039$<br>$p=0.0827$<br>$p=0.2018$<br>$p=0.0826$<br>$p>0.9999$ |
| Figure 9-1B<br>Tx1<br>Dark Zone | One-way ANOVA for treatment<br>Bonferroni's multiple comparisons<br>- Veh(1) vs. Veh(2)<br>- Veh(1) vs. P-38(1)<br>- Veh(1) vs. P-38(2)<br>- Veh(2) vs. P-38(1)<br>- Veh(2) vs. P-38(2)<br>- P-38(1) vs. P-38(2) | $F_{(3,106)}=11.41$, $p<0.0001$<br><br>$p>0.9999$<br>$p=0.0002$ (***)<br>$p=0.0001$<br>$p=0.0016$<br>$p=0.0013$ (^^)<br>$p>0.9999$ |
| Figure 9-1B<br>Tx2<br>Dark Zone | One-way ANOVA for treatment<br>Bonferroni's multiple comparisons<br>- Veh(1) + Con Ab vs. Veh(2) + CGRP Ab<br>- Veh(1) + Con Ab vs. P-38(1) + Con Ab<br>- Veh(1) + Con Ab vs. P-38(2) + CGRP Ab<br>- Veh(2) + CGRP Ab vs. P-38(1) + Con Ab<br>- Veh(2) + CGRP Ab vs. P-38(2) + CGRP Ab<br>- P-38(1) + Con Ab vs. P-38(2) + CGRP Ab | $F_{(3,106)}=5.527$, $p=0.0015$<br><br>$p>0.9999$<br>$p=0.0180$ (*)<br>$p=0.0141$<br>$p=0.0619$<br>$p=0.0503$<br>$p>0.9999$ |
| Figure 9-1B<br>Tx1<br>Light Zone | One-way ANOVA for treatment<br>Bonferroni's multiple comparisons<br>- Veh(1) vs. Veh(2)<br>- Veh(1) vs. P-38(1)<br>- Veh(1) vs. P-38(2)<br>- Veh(2) vs. P-38(1)<br>- Veh(2) vs. P-38(2)<br>- P-38(1) vs. P-38(2) | $F_{(3,106)}=1.793$, $p=0.1531$<br><br>$p>0.9999$<br>$p=0.8878$<br>$p>0.9999$<br>$p>0.9999$<br>$p>0.9999$<br>$p=0.1555$ |
| Figure 9-1B<br>Tx2<br>Light Zone | One-way ANOVA for treatment<br>Bonferroni's multiple comparisons<br>- Veh(1) + Con Ab vs. Veh(2) + CGRP Ab<br>- Veh(1) + Con Ab vs. P-38(1) + Con Ab<br>- Veh(1) + Con Ab vs. P-38(2) + CGRP Ab<br>- Veh(2) + CGRP Ab vs. P-38(1) + Con Ab<br>- Veh(2) + CGRP Ab vs. P-38(2) + CGRP Ab<br>- P-38(1) + Con Ab vs. P-38(2) + CGRP Ab | $F_{(3,106)}=1.947$, $p=0.1265$<br><br>$p>0.9999$<br>$p=0.1467$<br>$p>0.9999$<br>$p=0.4941$<br>$p>0.9999$<br>$p>0.9999$ |
| Figure 9-1C<br>Tx1 | One-way ANOVA for treatment<br>Bonferroni's multiple comparisons<br>- Veh(1) vs. Veh(2)<br>- Veh(1) vs. P-38(1)<br>- Veh(1) vs. P-38(2) | $F_{(3,106)}=20.99$, $p<0.0001$<br><br>$p>0.9999$<br>$p<0.0001$ (****)<br>$p<0.0001$ |

| | | |
|---|---|---|
| | - Veh(2) vs. P-38(1)<br>- Veh(2) vs. P-38(2)<br>- P-38(1) vs. P-38(2) | $p<0.0001$<br>$p<0.0001$ (^^^^)<br>$p>0.9999$ |
| Figure 9-1C<br>Tx2 | One-way ANOVA for treatment<br>Bonferroni's multiple comparisons<br>- Veh(1) + Con Ab vs. Veh(2) + CGRP Ab<br>- Veh(1) + Con Ab vs. P-38(1) + Con Ab<br>- Veh(1) + Con Ab vs. P-38(2) + CGRP Ab<br>- Veh(2) + CGRP Ab vs. P-38(1) + Con Ab<br>- Veh(2) + CGRP Ab vs. P-38(2) + CGRP Ab<br>- P-38(1) + Con Ab vs. P-38(2) + CGRP Ab | $F_{(3,106)}=10.44$, $p<0.0001$<br><br>$p>0.9999$<br>$p=0.0004$ (***)<br>$p=0.0008$<br>$p=0.0009$<br>$p=0.0016$ (^^)<br>$p>0.9999$ |

Table 4-1. Percerent of responders in F1 progeny.

| F1 Crosses | Responder x Responder | Responder x Nonresponder | Nonresponder x Nonresponder |
|---|---|---|---|
| # of breeding pairs | 4 | 3 | 3 |
| # of mice tested with PACAP | 59 (29 F, 30 M) | 56 (25 F, 31 M) | 21 (13 F, 8 M) |
| # of responders | 52 (26 F, 26 M) | 41 (18 F, 23 M) | 5 (5 F, 0 M) |
| # of non-responders | 7 (3 F, 4 M) | 15 (7 F, 8 M) | 16 (8 F, 8 M) |
| % of responders | **88.1** | **73.2** | **23.8** |
| # of mice tested with Vehicle | 38 (20 F, 18 M) | 40 (19 F, 21 M) | 15 (8 F, 7 M) |
| # of responders | 12 (6 F, 6 M) | 13 (6 F, 7 M) | 3 (1 F, 2 M) |
| # of non-responders | 26 (14 F, 12 M) | 27 (13 F, 14 M) | 12 (7 F, 5 M) |
| % of responders | **31.6** | **32.5** | **20.0** |

Table 5-1. Relative gene expression in responder and nonresponder mice.

| Gene | baseMean | log2FC | lfcSE | stat | pvalue | padj | chr | start | end |
|---|---|---|---|---|---|---|---|---|---|
| AL731706.1 | 432.182 | 26.193 | 3.003 | 8.722 | 2.75E-18 | 2.09E-14 | 2 | 136881271 | 136891389 |
| Rpl31-ps21 | 9.816 | 6.693 | 1.615 | 4.144 | 3.41E-05 | 9.97E-03 | 5 | 21119149 | 21119522 |
| Gnrhr | 519.128 | 6.149 | 1.503 | 4.090 | 4.31E-05 | 1.17E-02 | 5 | 86180754 | 86197901 |
| Fshb | 4958.379 | 5.307 | 1.257 | 4.223 | 2.41E-05 | 8.46E-03 | 2 | 107056140 | 107059656 |
| Cga | 15581.178 | 4.939 | 1.167 | 4.232 | 2.32E-05 | 8.28E-03 | 4 | 34893779 | 34907370 |
| Gm48684 | 11.523 | 4.909 | 1.309 | 3.750 | 1.77E-04 | 3.30E-02 | 13 | 107034284 | 107035217 |
| Ghrhr | 750.265 | 4.893 | 1.280 | 3.822 | 1.32E-04 | 2.65E-02 | 6 | 55376295 | 55388530 |
| Gm45623 | 11.070 | 4.839 | 1.306 | 3.705 | 2.12E-04 | 3.74E-02 | 13 | 56752371 | 56753827 |
| Lhb | 3605.568 | 4.725 | 1.178 | 4.011 | 6.05E-05 | 1.55E-02 | 7 | 45420820 | 45421897 |
| Dlk1 | 5653.682 | 4.594 | 1.178 | 3.899 | 9.65E-05 | 2.08E-02 | 12 | 109452823 | 109463336 |
| Gm14165 | 83.599 | 4.509 | 0.652 | 6.916 | 4.64E-12 | 1.51E-08 | 2 | 152353615 | 152354566 |
| Gm6166 | 469.092 | 4.439 | 0.310 | 14.305 | 2.04E-46 | 4.66E-42 | 9 | 57483963 | 57484371 |
| Gh | 524571.303 | 4.364 | 1.181 | 3.695 | 2.20E-04 | 3.87E-02 | 11 | 106300271 | 106301865 |
| Npm3-ps1 | 6.662 | 4.341 | 1.090 | 3.984 | 6.79E-05 | 1.65E-02 | 6 | 85076192 | 85076719 |
| Tshb | 3355.026 | 4.304 | 1.038 | 4.148 | 3.35E-05 | 9.94E-03 | 3 | 102775465 | 102782718 |
| Gm8730 | 1224.168 | 3.698 | 0.456 | 8.112 | 5.00E-16 | 2.33E-12 | 8 | 102864845 | 102865798 |
| Sstr3 | 306.322 | 3.687 | 0.863 | 4.270 | 1.95E-05 | 7.08E-03 | 15 | 78537008 | 78544685 |
| Lhx3 | 395.410 | 3.619 | 0.870 | 4.161 | 3.17E-05 | 9.50E-03 | 2 | 26200212 | 26208289 |
| Lmbr1 | 569.738 | 3.478 | 0.647 | 5.372 | 7.78E-08 | 1.00E-04 | 5 | 29229802 | 29378390 |
| Gm3636 | 32.211 | 3.470 | 0.789 | 4.400 | 1.08E-05 | 4.67E-03 | 14 | 6735685 | 6742332 |
| Tgfbr3l | 1174.427 | 2.747 | 0.756 | 3.632 | 2.81E-04 | 4.58E-02 | 8 | 4248214 | 4251423 |
| Got2-ps1 | 393.306 | 2.516 | 0.445 | 5.651 | 1.59E-08 | 2.60E-05 | 5 | 138364260 | 138365552 |
| Hdac2 | 1191.805 | 2.413 | 0.597 | 4.044 | 5.26E-05 | 1.38E-02 | 10 | 36974544 | 37001889 |
| Znrd1as | 91.907 | 2.246 | 0.597 | 3.763 | 1.68E-04 | 3.17E-02 | 17 | 36958592 | 36965625 |
| Nts | 542.442 | 1.725 | 0.383 | 4.505 | 6.64E-06 | 3.23E-03 | 10 | 102481756 | 102490486 |
| AI506816 | 334.963 | 1.606 | 0.347 | 4.632 | 3.63E-06 | 1.84E-03 | 5 | 23698296 | 23712667 |
| Tmsb15l | 324.655 | 1.438 | 0.274 | 5.248 | 1.54E-07 | 1.76E-04 | X | 136954988 | 136976869 |
| Ndufs5 | 2062.789 | 1.408 | 0.197 | 7.134 | 9.73E-13 | 3.70E-09 | 4 | 123712710 | 123718202 |
| Rcan3 | 399.242 | 1.365 | 0.229 | 5.953 | 2.63E-09 | 5.00E-06 | 4 | 135412308 | 135433853 |
| Ctdsp2-ps | 249.138 | 1.304 | 0.302 | 4.314 | 1.60E-05 | 6.31E-03 | 10 | 130412499 | 130413309 |
| Trpc5 | 275.709 | 1.050 | 0.278 | 3.778 | 1.58E-04 | 3.03E-02 | X | 144381671 | 144688180 |
| C130046K22Rik | 110.888 | 1.047 | 0.256 | 4.094 | 4.24E-05 | 1.17E-02 | 11 | 103697724 | 103722832 |
| Phlda2 | 87.700 | 1.032 | 0.280 | 3.688 | 2.26E-04 | 3.93E-02 | 7 | 143501545 | 143503150 |
| Zdhhc24 | 1621.796 | 0.936 | 0.225 | 4.164 | 3.12E-05 | 9.50E-03 | 19 | 4878668 | 4885397 |
| Tmsb15b1 | 129.821 | 0.899 | 0.241 | 3.733 | 1.89E-04 | 3.41E-02 | X | 136974022 | 136976874 |
| Gm20186 | 60.808 | 0.893 | 0.245 | 3.647 | 2.65E-04 | 4.38E-02 | 6 | 18845910 | 18848207 |
| B230110C06Rik | 195.470 | 0.876 | 0.195 | 4.493 | 7.01E-06 | 3.26E-03 | 14 | 15437623 | 15452445 |
| Kcnk12 | 790.427 | 0.743 | 0.196 | 3.785 | 1.54E-04 | 2.99E-02 | 17 | 87745801 | 87797994 |
| Gm10874 | 1039.685 | 0.665 | 0.121 | 5.482 | 4.21E-08 | 6.40E-05 | 5 | 138363719 | 138388287 |
| Zbtb5 | 1158.101 | 0.647 | 0.164 | 3.949 | 7.84E-05 | 1.79E-02 | 4 | 44991242 | 45012412 |
| Rab33b | 2893.657 | 0.614 | 0.155 | 3.960 | 7.48E-05 | 1.76E-02 | 3 | 51483920 | 51496232 |

| | | | | | | | | | |
|---|---|---|---|---|---|---|---|---|---|
| Agap1 | 2589.232 | 0.584 | 0.136 | 4.299 | 1.72E-05 | 6.51E-03 | 1 | 89454806 | 89897617 |
| Il31ra | 993.461 | 0.545 | 0.151 | 3.602 | 3.16E-04 | 4.91E-02 | 13 | 112519898 | 112594360 |
| Klhl32 | 916.748 | 0.529 | 0.146 | 3.621 | 2.93E-04 | 4.70E-02 | 4 | 24612554 | 24851124 |
| Patj | 6755.194 | 0.491 | 0.124 | 3.953 | 7.73E-05 | 1.79E-02 | 4 | 98395785 | 98719603 |
| Ubb-ps | 21929.180 | 0.467 | 0.109 | 4.296 | 1.74E-05 | 6.51E-03 | 14 | 46084028 | 46085000 |
| Gaa | 23705.667 | 0.449 | 0.100 | 4.508 | 6.56E-06 | 3.23E-03 | 11 | 119267887 | 119285454 |
| Mcf2l | 6905.208 | 0.330 | 0.091 | 3.612 | 3.04E-04 | 4.79E-02 | 8 | 12873806 | 13020905 |
| Polr2b | 7295.129 | 0.324 | 0.064 | 5.026 | 5.02E-07 | 4.09E-04 | 5 | 77310147 | 77349324 |
| Atg10 | 1175.204 | 0.319 | 0.087 | 3.659 | 2.54E-04 | 4.23E-02 | 13 | 90935356 | 91223968 |
| Fkbp10 | 1111.745 | -0.502 | 0.137 | -3.664 | 2.48E-04 | 4.20E-02 | 11 | 100415697 | 100424824 |
| Marcks | 8426.932 | -0.518 | 0.126 | -4.121 | 3.77E-05 | 1.09E-02 | 10 | 37133375 | 37138920 |
| Josd2 | 2339.114 | -0.550 | 0.125 | -4.413 | 1.02E-05 | 4.48E-03 | 7 | 44467980 | 44471662 |
| Cdo1 | 1572.498 | -0.552 | 0.144 | -3.842 | 1.22E-04 | 2.49E-02 | 18 | 46713193 | 46728395 |
| Eva1b | 786.985 | -0.559 | 0.108 | -5.166 | 2.39E-07 | 2.27E-04 | 4 | 126147744 | 126149875 |
| Gpx8 | 2064.449 | -0.582 | 0.148 | -3.924 | 8.70E-05 | 1.89E-02 | 13 | 113042753 | 113046410 |
| Fxyd3 | 6172.465 | -0.597 | 0.160 | -3.738 | 1.85E-04 | 3.41E-02 | 7 | 31068172 | 31076704 |
| Enpp1 | 2987.450 | -0.634 | 0.128 | -4.955 | 7.25E-07 | 5.52E-04 | 10 | 24637914 | 24712159 |
| Myl9 | 3722.178 | -0.700 | 0.192 | -3.645 | 2.67E-04 | 4.39E-02 | 2 | 156775420 | 156781658 |
| Angptl2 | 3149.080 | -0.735 | 0.184 | -3.983 | 6.80E-05 | 1.65E-02 | 2 | 33216069 | 33247717 |
| Bmp7 | 3560.010 | -0.750 | 0.202 | -3.721 | 1.98E-04 | 3.53E-02 | 2 | 172868012 | 172940321 |
| Lpar4 | 246.524 | -0.762 | 0.159 | -4.787 | 1.70E-06 | 1.04E-03 | X | 106920625 | 106933900 |
| Dcdc2a | 348.339 | -0.764 | 0.209 | -3.658 | 2.54E-04 | 4.23E-02 | 13 | 25056004 | 25210706 |
| Gldn | 6382.100 | -0.768 | 0.212 | -3.618 | 2.97E-04 | 4.71E-02 | 9 | 54286486 | 54341786 |
| Col9a2 | 1434.476 | -0.773 | 0.178 | -4.336 | 1.45E-05 | 5.80E-03 | 4 | 121039385 | 121055322 |
| Car14 | 164.190 | -0.781 | 0.196 | -3.985 | 6.74E-05 | 1.65E-02 | 3 | 95897768 | 95904691 |
| Rflna | 136.591 | -0.789 | 0.219 | -3.606 | 3.11E-04 | 4.86E-02 | 5 | 125003221 | 125012547 |
| Colca2 | 66.926 | -0.811 | 0.220 | -3.680 | 2.33E-04 | 3.98E-02 | 9 | 51270258 | 51279651 |
| Acot1 | 983.839 | -0.823 | 0.153 | -5.369 | 7.90E-08 | 1.00E-04 | 12 | 84009490 | 84018371 |
| Itm2a | 11680.832 | -0.859 | 0.164 | -5.230 | 1.69E-07 | 1.81E-04 | X | 107397099 | 107403376 |
| Gm9780 | 273.575 | -0.905 | 0.168 | -5.390 | 7.04E-08 | 1.00E-04 | 14 | 26027282 | 26042711 |
| Pla2g4a | 1104.112 | -0.958 | 0.249 | -3.847 | 1.19E-04 | 2.48E-02 | 1 | 149829618 | 149961290 |
| Fndc1 | 1007.392 | -1.011 | 0.242 | -4.174 | 3.00E-05 | 9.50E-03 | 17 | 7738569 | 7827302 |
| Ssc5d | 394.539 | -1.036 | 0.214 | -4.832 | 1.35E-06 | 9.05E-04 | 7 | 4925785 | 4944826 |
| Aebp1 | 5509.341 | -1.045 | 0.254 | -4.114 | 3.89E-05 | 1.11E-02 | 11 | 5861947 | 5872088 |
| Dipk2b | 148.434 | -1.060 | 0.269 | -3.935 | 8.33E-05 | 1.83E-02 | X | 18414881 | 18461397 |
| Aoc3 | 318.697 | -1.072 | 0.272 | -3.941 | 8.12E-05 | 1.80E-02 | 11 | 101330605 | 101341938 |
| Plac9b | 224.046 | -1.078 | 0.158 | -6.815 | 9.39E-12 | 2.68E-08 | 14 | 26167405 | 26182480 |
| Gpr4 | 251.441 | -1.120 | 0.234 | -4.794 | 1.63E-06 | 1.04E-03 | 7 | 19212538 | 19224174 |
| Gkn3 | 145.709 | -1.179 | 0.254 | -4.645 | 3.40E-06 | 1.76E-03 | 6 | 87383256 | 87388935 |
| Rpl15-ps6 | 1745.827 | -1.180 | 0.328 | -3.599 | 3.19E-04 | 4.92E-02 | 15 | 52477617 | 52478228 |
| Ptgis | 883.395 | -1.186 | 0.270 | -4.386 | 1.15E-05 | 4.87E-03 | 2 | 167191805 | 167240604 |
| Tmem119 | 644.499 | -1.200 | 0.312 | -3.849 | 1.18E-04 | 2.48E-02 | 5 | 113793729 | 113800516 |
| Psmc3ip | 181.160 | -1.235 | 0.326 | -3.783 | 1.55E-04 | 2.99E-02 | 11 | 101091823 | 101095436 |
| Epha3 | 496.061 | -1.242 | 0.238 | -5.212 | 1.87E-07 | 1.85E-04 | 16 | 63543534 | 63864175 |

| | | | | | | | | | |
|---|---|---|---|---|---|---|---|---|---|
| Il3ra | 248.176 | -1.278 | 0.330 | -3.877 | 1.06E-04 | 2.26E-02 | 14 | 14346275 | 14356011 |
| Cdh3 | 115.939 | -1.289 | 0.254 | -5.079 | 3.80E-07 | 3.34E-04 | 8 | 106510891 | 106557297 |
| Gxylt2 | 618.378 | -1.289 | 0.345 | -3.732 | 1.90E-04 | 3.41E-02 | 6 | 100704734 | 100810913 |
| H2-DMb1 | 888.510 | -1.336 | 0.358 | -3.736 | 1.87E-04 | 3.41E-02 | 17 | 34153072 | 34160230 |
| Ccn2 | 83.261 | -1.348 | 0.342 | -3.941 | 8.11E-05 | 1.80E-02 | 10 | 24595442 | 24598683 |
| Mall | 67.643 | -1.364 | 0.327 | -4.167 | 3.08E-05 | 9.50E-03 | 2 | 127704386 | 127729932 |
| Ppic | 1058.164 | -1.378 | 0.349 | -3.944 | 8.00E-05 | 1.80E-02 | 18 | 53406332 | 53418115 |
| Gm6477 | 136.179 | -1.425 | 0.270 | -5.284 | 1.26E-07 | 1.52E-04 | 10 | 39198538 | 39199301 |
| Omd | 708.961 | -1.506 | 0.378 | -3.987 | 6.69E-05 | 1.65E-02 | 13 | 49582462 | 49592822 |
| Slc13a4 | 2172.741 | -1.559 | 0.371 | -4.198 | 2.69E-05 | 9.17E-03 | 6 | 35267957 | 35308131 |
| Fam180a | 321.602 | -1.565 | 0.311 | -5.033 | 4.84E-07 | 4.09E-04 | 6 | 35312668 | 35326141 |
| Nupr1 | 798.900 | -1.574 | 0.383 | -4.108 | 3.99E-05 | 1.11E-02 | 7 | 126623249 | 126630861 |
| Sp7 | 386.792 | -1.575 | 0.376 | -4.192 | 2.77E-05 | 9.21E-03 | 15 | 102356606 | 102367182 |
| Alx4 | 488.115 | -1.601 | 0.404 | -3.963 | 7.41E-05 | 1.76E-02 | 2 | 93642384 | 93681339 |
| Ypel3 | 2733.371 | -1.626 | 0.343 | -4.733 | 2.21E-06 | 1.25E-03 | 7 | 126776955 | 126780514 |
| Gm38101 | 37.246 | -1.629 | 0.330 | -4.939 | 7.87E-07 | 5.79E-04 | 1 | 176276213 | 176279769 |
| Pmp2 | 1026.497 | -1.685 | 0.346 | -4.877 | 1.08E-06 | 7.69E-04 | 3 | 10179851 | 10183929 |
| Kcmf1 | 1426.414 | -1.720 | 0.445 | -3.860 | 1.13E-04 | 2.39E-02 | 6 | 72841114 | 72899979 |
| 9130024F11Rik | 59.795 | -1.720 | 0.394 | -4.368 | 1.25E-05 | 5.20E-03 | 1 | 56971412 | 57050084 |
| Wt1 | 151.949 | -1.746 | 0.462 | -3.776 | 1.59E-04 | 3.03E-02 | 2 | 105126529 | 105173616 |
| Col12a1 | 335.044 | -1.750 | 0.434 | -4.031 | 5.56E-05 | 1.44E-02 | 9 | 79598991 | 79718831 |
| Osr1 | 384.639 | -1.750 | 0.455 | -3.842 | 1.22E-04 | 2.49E-02 | 12 | 9570116 | 9581500 |
| Gm37412 | 93.917 | -1.789 | 0.300 | -5.970 | 2.37E-09 | 4.91E-06 | 1 | 14788645 | 14792965 |
| Clec3b | 389.071 | -1.817 | 0.460 | -3.949 | 7.85E-05 | 1.79E-02 | 9 | 123150946 | 123157432 |
| Car3 | 1151.775 | -1.852 | 0.354 | -5.225 | 1.74E-07 | 1.81E-04 | 3 | 14863512 | 14872523 |
| Fmod | 4296.396 | -1.868 | 0.390 | -4.783 | 1.73E-06 | 1.04E-03 | 1 | 134037254 | 134048277 |
| Dpt | 283.898 | -1.952 | 0.479 | -4.077 | 4.56E-05 | 1.22E-02 | 1 | 164796644 | 164824266 |
| Nphs2 | 194.627 | -2.037 | 0.429 | -4.744 | 2.10E-06 | 1.23E-03 | 1 | 156310727 | 156328035 |
| 9630028B13Rik | 92.398 | -2.064 | 0.481 | -4.290 | 1.79E-05 | 6.58E-03 | 1 | 185429355 | 185441819 |
| Col1a1 | 39251.325 | -2.085 | 0.566 | -3.686 | 2.28E-04 | 3.94E-02 | 11 | 94936224 | 94953042 |
| Trim5 | 74.290 | -2.119 | 0.508 | -4.172 | 3.02E-05 | 9.50E-03 | 7 | 104263386 | 104288094 |
| Fn1 | 2558.613 | -2.178 | 0.602 | -3.620 | 2.94E-04 | 4.70E-02 | 1 | 71585520 | 71653200 |
| Tbx15 | 1143.178 | -2.275 | 0.628 | -3.624 | 2.90E-04 | 4.69E-02 | 3 | 99240381 | 99354259 |
| Fcgbp | 38.396 | -2.325 | 0.587 | -3.960 | 7.50E-05 | 1.76E-02 | 7 | 28071236 | 28120862 |
| Cthrc1 | 319.329 | -2.343 | 0.250 | -9.354 | 8.46E-21 | 9.65E-17 | 15 | 39076932 | 39087121 |
| Asns | 2057.806 | -2.359 | 0.547 | -4.309 | 1.64E-05 | 6.33E-03 | 6 | 7675169 | 7693254 |
| Il13ra2 | 59.292 | -2.369 | 0.644 | -3.679 | 2.34E-04 | 3.98E-02 | X | 147383476 | 147429192 |
| Barx1 | 125.692 | -2.458 | 0.553 | -4.446 | 8.74E-06 | 3.99E-03 | 13 | 48662998 | 48666507 |
| Postn | 3786.294 | -2.460 | 0.547 | -4.495 | 6.96E-06 | 3.26E-03 | 3 | 54361109 | 54391037 |
| Dnah14 | 123.325 | -2.460 | 0.483 | -5.096 | 3.47E-07 | 3.17E-04 | 1 | 181576559 | 181815774 |
| Fabp4 | 1066.377 | -2.472 | 0.590 | -4.190 | 2.79E-05 | 9.21E-03 | 3 | 10204088 | 10208576 |
| Alx3 | 78.374 | -2.681 | 0.461 | -5.819 | 5.93E-09 | 1.04E-05 | 3 | 107595031 | 107605776 |
| Tnni1 | 183.021 | -2.839 | 0.569 | -4.987 | 6.14E-07 | 4.83E-04 | 1 | 135779434 | 135810989 |
| Lpp | 116.505 | -3.072 | 0.811 | -3.785 | 1.54E-04 | 2.99E-02 | 16 | 24393507 | 24992578 |

| | | | | | | | | | |
|---|---|---|---|---|---|---|---|---|---|
| Slc2a4 | 211.636 | -3.097 | 0.742 | -4.173 | 3.01E-05 | 9.50E-03 | 11 | 69942539 | 69948188 |
| Bglap2 | 665.091 | -3.605 | 0.814 | -4.426 | 9.59E-06 | 4.29E-03 | 3 | 88377736 | 88378699 |
| Cidec | 58.634 | -3.803 | 0.903 | -4.209 | 2.56E-05 | 8.87E-03 | 6 | 113424634 | 113435760 |
| Oscar | 29.629 | -3.965 | 1.058 | -3.748 | 1.78E-04 | 3.30E-02 | 7 | 3609813 | 3616157 |
| Aspn | 44.454 | -3.972 | 0.845 | -4.699 | 2.61E-06 | 1.42E-03 | 13 | 49544443 | 49567565 |
| Dlx5 | 71.096 | -4.039 | 0.834 | -4.840 | 1.30E-06 | 8.98E-04 | 6 | 6877805 | 6882085 |
| Wnt10b | 37.167 | -4.069 | 1.004 | -4.051 | 5.11E-05 | 1.36E-02 | 15 | 98770712 | 98778150 |
| E130008D07Rik | 60.103 | -4.208 | 1.112 | -3.783 | 1.55E-04 | 2.99E-02 | 17 | 43146041 | 43158291 |
| Emx2 | 26.672 | -4.395 | 0.676 | -6.502 | 7.94E-11 | 2.01E-07 | 19 | 59458372 | 59465357 |
| Pck1 | 145.305 | -4.482 | 1.027 | -4.363 | 1.28E-05 | 5.23E-03 | 2 | 173153048 | 173159273 |
| Vav1 | 97.741 | -4.548 | 1.107 | -4.110 | 3.95E-05 | 1.11E-02 | 17 | 57279100 | 57328031 |
| Hivep2 | 762.606 | -4.651 | 0.742 | -6.270 | 3.62E-10 | 8.25E-07 | 10 | 13966075 | 14151374 |
| 1700085C21Rik | 35.918 | -5.144 | 1.070 | -4.807 | 1.53E-06 | 9.99E-04 | 12 | 82932520 | 82939155 |
| Cfd | 118.141 | -5.370 | 1.151 | -4.664 | 3.09E-06 | 1.64E-03 | 10 | 79890853 | 79892655 |
| mt-Ts2 | 34.393 | -5.697 | 1.204 | -4.731 | 2.24E-06 | 1.25E-03 | MT | 11613 | 11671 |
| Fezf2 | 15.198 | -5.956 | 1.556 | -3.828 | 1.29E-04 | 2.61E-02 | 14 | 12342094 | 12348189 |
| Gm45472 | 17.112 | -6.042 | 1.510 | -4.001 | 6.30E-05 | 1.60E-02 | 8 | 92853653 | 92856334 |
| C130093G08Rik | 8.617 | -6.418 | 1.542 | -4.162 | 3.16E-05 | 9.50E-03 | 6 | 23251133 | 23253358 |
| Ugt2a1 | 66.037 | -24.801 | 3.058 | -8.109 | 5.10E-16 | 2.33E-12 | 5 | 87459490 | 87490871 |

Table 5-2. Relative expression in responder and nonresponder male mice only.

| Gene | baseMean | log2FC | lfcSE | stat | pvalue | padj | chr | start | end |
|---|---|---|---|---|---|---|---|---|---|
| Galr3 | 14.124 | 20.871 | 3.648 | 5.721 | 1.06E-08 | 9.01E-06 | 15 | 79041885 | 79043558 |
| Pcdhb1 | 141.221 | 10.824 | 2.570 | 4.212 | 2.54E-05 | 6.36E-03 | 18 | 37264938 | 37267525 |
| Nr0b1 | 52.614 | 9.399 | 1.939 | 4.848 | 1.25E-06 | 6.11E-04 | X | 86191764 | 86195947 |
| Bfsp1 | 39.151 | 8.974 | 1.860 | 4.825 | 1.40E-06 | 6.72E-04 | 2 | 143826528 | 143863173 |
| Lrp8os2 | 36.348 | 8.865 | 2.416 | 3.668 | 2.44E-04 | 2.85E-02 | 4 | 107806033 | 107823022 |
| Smco2 | 34.401 | 8.786 | 2.504 | 3.508 | 4.51E-04 | 4.23E-02 | 6 | 146850104 | 146871406 |
| Mmp20 | 30.770 | 8.625 | 2.419 | 3.565 | 3.63E-04 | 3.74E-02 | 9 | 7628231 | 7674979 |
| Gnrhr | 868.887 | 8.565 | 2.113 | 4.053 | 5.06E-05 | 1.02E-02 | 5 | 86180754 | 86197901 |
| Gm14618 | 242.617 | 8.508 | 2.278 | 3.735 | 1.88E-04 | 2.35E-02 | 2 | 174842305 | 174853357 |
| Adam28 | 23.579 | 8.240 | 1.914 | 4.306 | 1.66E-05 | 4.65E-03 | 14 | 68606027 | 68655842 |
| Glra1 | 18.079 | 7.856 | 1.905 | 4.125 | 3.71E-05 | 8.11E-03 | 11 | 55514238 | 55608198 |
| Rax | 17.637 | 7.820 | 1.810 | 4.320 | 1.56E-05 | 4.40E-03 | 18 | 65928277 | 65939787 |
| Gm14617 | 372.632 | 7.691 | 1.974 | 3.896 | 9.76E-05 | 1.57E-02 | 2 | 174835964 | 174841174 |
| 5730420D15Rik | 14.963 | 7.584 | 2.053 | 3.694 | 2.20E-04 | 2.64E-02 | 10 | 95417375 | 95428640 |
| Fshb | 8942.662 | 6.996 | 1.752 | 3.993 | 6.52E-05 | 1.21E-02 | 2 | 107056140 | 107059656 |
| Lhb | 6282.795 | 6.902 | 1.721 | 4.012 | 6.03E-05 | 1.15E-02 | 7 | 45420820 | 45421897 |
| Olfr283 | 9.268 | 6.894 | 1.913 | 3.603 | 3.14E-04 | 3.43E-02 | 15 | 98377699 | 98380946 |
| 5730403I07Rik | 226.948 | 6.844 | 1.461 | 4.685 | 2.80E-06 | 1.13E-03 | 9 | 77366412 | 77399418 |
| Dlk1 | 9688.871 | 6.719 | 1.633 | 4.113 | 3.90E-05 | 8.39E-03 | 12 | 109452823 | 109463336 |
| Cga | 26998.015 | 6.659 | 1.771 | 3.759 | 1.70E-04 | 2.21E-02 | 4 | 34893779 | 34907370 |
| Gm14685 | 53.896 | 6.576 | 1.739 | 3.782 | 1.55E-04 | 2.09E-02 | X | 73117047 | 73129262 |
| Irs4 | 1003.694 | 6.554 | 1.647 | 3.979 | 6.92E-05 | 1.23E-02 | X | 141710998 | 141725263 |
| Slc5a8 | 99.488 | 6.536 | 1.487 | 4.395 | 1.11E-05 | 3.31E-03 | 10 | 88885992 | 88929515 |
| Gm13691 | 13.102 | 6.419 | 1.668 | 3.848 | 1.19E-04 | 1.80E-02 | 2 | 83990553 | 83993045 |
| Gm13697 | 13.102 | 6.419 | 1.668 | 3.848 | 1.19E-04 | 1.80E-02 | 2 | 83984743 | 83987235 |
| Gm13694 | 13.102 | 6.419 | 1.668 | 3.848 | 1.19E-04 | 1.80E-02 | 2 | 83978933 | 83981425 |
| Gm13696 | 13.102 | 6.419 | 1.668 | 3.848 | 1.19E-04 | 1.80E-02 | 2 | 83973123 | 83975615 |
| Gm13698 | 12.927 | 6.400 | 1.672 | 3.828 | 1.29E-04 | 1.84E-02 | 2 | 83955693 | 83958185 |
| Gm13693 | 12.927 | 6.400 | 1.672 | 3.828 | 1.29E-04 | 1.84E-02 | 2 | 83961503 | 83963995 |
| Gm13695 | 12.927 | 6.400 | 1.672 | 3.828 | 1.29E-04 | 1.84E-02 | 2 | 83967313 | 83969805 |
| Gh | 923606.332 | 6.383 | 1.696 | 3.764 | 1.67E-04 | 2.18E-02 | 11 | 106300271 | 106301865 |
| Tshb | 4940.895 | 6.269 | 1.569 | 3.996 | 6.46E-05 | 1.21E-02 | 3 | 102775465 | 102782718 |
| Pitx1 | 1643.969 | 6.229 | 1.576 | 3.953 | 7.73E-05 | 1.32E-02 | 13 | 55825051 | 55836192 |
| Upk3a | 86.408 | 6.175 | 1.159 | 5.330 | 9.83E-08 | 6.02E-05 | 15 | 85017141 | 85022547 |
| Prl | 138564.338 | 6.115 | 1.708 | 3.579 | 3.44E-04 | 3.64E-02 | 13 | 27057570 | 27065205 |
| DXBay18 | 50.138 | 6.085 | 1.545 | 3.939 | 8.19E-05 | 1.39E-02 | X | 73137232 | 73149450 |
| Pitx2 | 282.215 | 6.001 | 1.296 | 4.629 | 3.68E-06 | 1.38E-03 | 3 | 129199878 | 129219591 |
| Ahrr | 15.911 | 5.996 | 1.735 | 3.457 | 5.47E-04 | 4.79E-02 | 13 | 74211118 | 74292331 |
| Ghrhr | 1244.091 | 5.971 | 1.723 | 3.466 | 5.29E-04 | 4.72E-02 | 6 | 55376295 | 55388530 |
| Six6 | 1415.464 | 5.927 | 1.512 | 3.920 | 8.84E-05 | 1.43E-02 | 12 | 72939892 | 72944899 |
| Epcam | 4719.791 | 5.796 | 1.561 | 3.712 | 2.06E-04 | 2.55E-02 | 17 | 87635979 | 87651129 |

| | | | | | | | | | |
|---|---|---|---|---|---|---|---|---|---|
| Tmem184a | 913.208 | 5.764 | 1.141 | 5.050 | 4.41E-07 | 2.37E-04 | 5 | 139802485 | 139819917 |
| 1810064F22Rik | 95.327 | 5.738 | 1.183 | 4.851 | 1.23E-06 | 6.11E-04 | 9 | 22196963 | 22213860 |
| Lhx3 | 700.554 | 5.721 | 1.128 | 5.073 | 3.91E-07 | 2.16E-04 | 2 | 26200212 | 26208289 |
| Elmod1 | 71.263 | 5.582 | 1.424 | 3.920 | 8.84E-05 | 1.43E-02 | 9 | 53911457 | 53975301 |
| Gm49542 | 89.225 | 5.379 | 1.398 | 3.846 | 1.20E-04 | 1.80E-02 | 14 | 79898467 | 79907230 |
| Rnf183 | 62.569 | 5.263 | 1.521 | 3.460 | 5.41E-04 | 4.77E-02 | 4 | 62427540 | 62435252 |
| Cldn4 | 156.484 | 5.248 | 1.493 | 3.514 | 4.42E-04 | 4.20E-02 | 5 | 134945119 | 134946934 |
| Thrb | 208.518 | 5.187 | 1.234 | 4.204 | 2.62E-05 | 6.43E-03 | 14 | 17660261 | 18038090 |
| Asic4 | 733.509 | 4.874 | 1.100 | 4.431 | 9.38E-06 | 2.96E-03 | 1 | 75450436 | 75474343 |
| Cdcp1 | 373.043 | 4.746 | 1.118 | 4.244 | 2.20E-05 | 6.06E-03 | 9 | 123170824 | 123216038 |
| C5ar2 | 34.532 | 4.742 | 0.865 | 5.484 | 4.16E-08 | 2.96E-05 | 7 | 16234585 | 16244154 |
| Speer4e | 79.618 | 4.735 | 1.350 | 3.507 | 4.54E-04 | 4.23E-02 | 5 | 14933221 | 14938429 |
| Sstr3 | 488.670 | 4.724 | 1.126 | 4.196 | 2.72E-05 | 6.59E-03 | 15 | 78537008 | 78544685 |
| Ascl1 | 1064.295 | 4.664 | 1.057 | 4.414 | 1.01E-05 | 3.15E-03 | 10 | 87490819 | 87493660 |
| Gm27196 | 98.181 | 4.609 | 1.336 | 3.450 | 5.60E-04 | 4.85E-02 | 2 | 26206702 | 26209283 |
| C2cd4a | 82.535 | 4.570 | 1.259 | 3.629 | 2.84E-04 | 3.22E-02 | 9 | 67830489 | 67832330 |
| Esrp1 | 659.522 | 4.516 | 1.117 | 4.043 | 5.29E-05 | 1.04E-02 | 4 | 11331933 | 11386783 |
| Rmst | 72.586 | 4.491 | 1.173 | 3.829 | 1.29E-04 | 1.84E-02 | 10 | 92071037 | 92165170 |
| Tgfbr3l | 1943.727 | 4.336 | 1.093 | 3.968 | 7.25E-05 | 1.26E-02 | 8 | 4248214 | 4251423 |
| Gm6166 | 406.630 | 4.324 | 0.421 | 10.270 | 9.63E-25 | 7.09E-21 | 9 | 57483963 | 57484371 |
| Cldn9 | 809.077 | 4.309 | 1.020 | 4.223 | 2.41E-05 | 6.11E-03 | 17 | 23682584 | 23684026 |
| Cpa2 | 161.470 | 4.297 | 1.194 | 3.599 | 3.19E-04 | 3.47E-02 | 6 | 30541582 | 30564476 |
| Prkch | 84.658 | 4.289 | 1.234 | 3.474 | 5.12E-04 | 4.61E-02 | 12 | 73584796 | 73778185 |
| 5330417C22Rik | 7427.302 | 4.268 | 1.010 | 4.225 | 2.39E-05 | 6.11E-03 | 3 | 108455694 | 108536536 |
| Gm14165 | 72.447 | 4.246 | 0.562 | 7.549 | 4.38E-14 | 1.21E-10 | 2 | 152353615 | 152354566 |
| Krt18 | 537.637 | 4.216 | 1.125 | 3.748 | 1.78E-04 | 2.28E-02 | 15 | 102028180 | 102032027 |
| Sec14l4 | 241.279 | 4.148 | 1.106 | 3.750 | 1.77E-04 | 2.27E-02 | 11 | 4031462 | 4048024 |
| Zfp985 | 73.675 | 4.146 | 0.952 | 4.355 | 1.33E-05 | 3.86E-03 | 4 | 147553277 | 147585198 |
| Gm10354 | 73.525 | 4.080 | 1.152 | 3.543 | 3.96E-04 | 3.95E-02 | 5 | 14974113 | 14978935 |
| Oacyl | 925.271 | 4.001 | 1.069 | 3.742 | 1.83E-04 | 2.30E-02 | 18 | 65698268 | 65751601 |
| Caps2 | 107.861 | 3.963 | 1.098 | 3.608 | 3.08E-04 | 3.38E-02 | 10 | 112163621 | 112216555 |
| 4932438H23Rik | 100.819 | 3.959 | 1.121 | 3.532 | 4.12E-04 | 4.04E-02 | 16 | 91053935 | 91095122 |
| Eppk1 | 210.669 | 3.951 | 1.112 | 3.552 | 3.82E-04 | 3.90E-02 | 15 | 76101481 | 76120195 |
| Drd4 | 45.090 | 3.781 | 1.023 | 3.697 | 2.18E-04 | 2.64E-02 | 7 | 141292006 | 141296464 |
| Lrrc10b | 673.300 | 3.780 | 0.986 | 3.835 | 1.26E-04 | 1.84E-02 | 19 | 10455371 | 10457473 |
| Tekt1 | 146.530 | 3.763 | 1.062 | 3.543 | 3.95E-04 | 3.95E-02 | 11 | 72344722 | 72362442 |
| Zfp125 | 198.246 | 3.628 | 1.002 | 3.622 | 2.92E-04 | 3.27E-02 | 12 | 20877084 | 20921262 |
| Ubxn10 | 347.602 | 3.577 | 1.017 | 3.517 | 4.37E-04 | 4.18E-02 | 4 | 138709837 | 138737167 |
| Nnat | 12493.422 | 3.536 | 0.846 | 4.182 | 2.89E-05 | 6.86E-03 | 2 | 157560078 | 157562522 |
| Ccdc187 | 902.238 | 3.514 | 0.938 | 3.745 | 1.80E-04 | 2.29E-02 | 2 | 26243469 | 26294557 |
| Lmbr1 | 708.664 | 3.465 | 0.967 | 3.584 | 3.38E-04 | 3.63E-02 | 5 | 29229802 | 29378390 |
| Gnb3 | 813.250 | 3.447 | 0.983 | 3.506 | 4.54E-04 | 4.23E-02 | 6 | 124834240 | 124840275 |
| Rtl1 | 306.419 | 3.407 | 0.942 | 3.617 | 2.99E-04 | 3.31E-02 | 12 | 109589193 | 109600330 |
| Gm8730 | 1076.073 | 3.333 | 0.282 | 11.806 | 3.64E-32 | 4.01E-28 | 8 | 102864845 | 102865798 |

| | | | | | | | | | |
|---|---|---|---|---|---|---|---|---|---|
| Drd2 | 879.063 | 3.253 | 0.884 | 3.680 | 2.33E-04 | 2.75E-02 | 9 | 49340627 | 49408177 |
| Pcdhgb1 | 1809.397 | 3.250 | 0.778 | 4.175 | 2.98E-05 | 6.98E-03 | 18 | 37680233 | 37841870 |
| Cul7 | 629.793 | 3.154 | 0.872 | 3.615 | 3.00E-04 | 3.31E-02 | 17 | 46650337 | 46664364 |
| Cwc22 | 5923.327 | 3.109 | 0.420 | 7.396 | 1.40E-13 | 3.09E-10 | 2 | 77881159 | 77946375 |
| Exph5 | 260.051 | 3.072 | 0.791 | 3.883 | 1.03E-04 | 1.62E-02 | 9 | 53301670 | 53377514 |
| Scg2 | 45948.307 | 3.009 | 0.796 | 3.782 | 1.56E-04 | 2.09E-02 | 1 | 79434669 | 79440120 |
| Gm49322 | 851.102 | 2.909 | 0.815 | 3.568 | 3.59E-04 | 3.74E-02 | 10 | 80606021 | 80615784 |
| Gm49380 | 87.885 | 2.827 | 0.816 | 3.465 | 5.31E-04 | 4.72E-02 | 9 | 44066327 | 44113500 |
| Stk33 | 248.446 | 2.804 | 0.745 | 3.764 | 1.67E-04 | 2.18E-02 | 7 | 109279223 | 109439081 |
| Fgf14 | 628.896 | 2.753 | 0.745 | 3.695 | 2.20E-04 | 2.64E-02 | 14 | 123977907 | 124677127 |
| Got2-ps1 | 456.086 | 2.578 | 0.321 | 8.028 | 9.89E-16 | 3.12E-12 | 5 | 138364260 | 138365552 |
| Spint1 | 472.905 | 2.518 | 0.535 | 4.705 | 2.54E-06 | 1.06E-03 | 2 | 119237362 | 119249527 |
| Gm48898 | 42.997 | 2.439 | 0.620 | 3.936 | 8.28E-05 | 1.39E-02 | 12 | 8291024 | 8293694 |
| Gm8399 | 1367.095 | 2.391 | 0.623 | 3.836 | 1.25E-04 | 1.84E-02 | 13 | 81064684 | 81065800 |
| Gm47980 | 26.721 | 2.326 | 0.652 | 3.567 | 3.61E-04 | 3.74E-02 | 12 | 108960912 | 108963832 |
| Capza2 | 748.903 | 2.244 | 0.408 | 5.505 | 3.70E-08 | 2.82E-05 | 6 | 17636234 | 17666972 |
| Tmem221 | 87.266 | 2.230 | 0.629 | 3.543 | 3.95E-04 | 3.95E-02 | 8 | 71554238 | 71558871 |
| Rlf | 224.656 | 2.183 | 0.398 | 5.485 | 4.13E-08 | 2.96E-05 | 4 | 121145373 | 121215084 |
| Resp18 | 247.024 | 2.150 | 0.360 | 5.970 | 2.37E-09 | 2.24E-06 | 1 | 75272199 | 75278415 |
| Tdrd5 | 68.524 | 2.089 | 0.578 | 3.616 | 2.99E-04 | 3.31E-02 | 1 | 156255296 | 156303664 |
| AI506816 | 323.166 | 2.086 | 0.509 | 4.102 | 4.10E-05 | 8.69E-03 | 5 | 23698296 | 23712667 |
| Fgfr1op2 | 2290.243 | 2.054 | 0.368 | 5.585 | 2.33E-08 | 1.84E-05 | 6 | 146577203 | 146599198 |
| Tmsb15l | 330.379 | 2.003 | 0.329 | 6.092 | 1.12E-09 | 1.17E-06 | X | 136954988 | 136976869 |
| 4930550C14Rik | 87.566 | 1.977 | 0.510 | 3.877 | 1.06E-04 | 1.65E-02 | 9 | 53402325 | 53434402 |
| Gm5909 | 251.218 | 1.950 | 0.365 | 5.346 | 9.02E-08 | 5.68E-05 | 8 | 80110806 | 80112086 |
| Pkib | 246.811 | 1.831 | 0.351 | 5.222 | 1.77E-07 | 1.03E-04 | 10 | 57631981 | 57741112 |
| Pfdn6 | 314.178 | 1.828 | 0.323 | 5.664 | 1.48E-08 | 1.21E-05 | 17 | 33938821 | 33940343 |
| Rcor2 | 262.090 | 1.723 | 0.279 | 6.166 | 6.99E-10 | 7.71E-07 | 19 | 7267325 | 7275225 |
| Ctdsp2-ps | 225.481 | 1.686 | 0.418 | 4.034 | 5.49E-05 | 1.06E-02 | 10 | 130412499 | 130413309 |
| Ccdc122 | 215.528 | 1.624 | 0.464 | 3.503 | 4.61E-04 | 4.25E-02 | 14 | 77036772 | 77112257 |
| Nudt12 | 473.144 | 1.521 | 0.425 | 3.580 | 3.44E-04 | 3.64E-02 | 17 | 58999618 | 59013372 |
| C130046K22Rik | 86.930 | 1.514 | 0.418 | 3.625 | 2.89E-04 | 3.26E-02 | 11 | 103697724 | 103722832 |
| Kif2a | 2688.395 | 1.346 | 0.271 | 4.967 | 6.80E-07 | 3.57E-04 | 13 | 106958996 | 107022126 |
| Ntrk3 | 1783.751 | 1.308 | 0.369 | 3.543 | 3.95E-04 | 3.95E-02 | 7 | 78175959 | 78738012 |
| Sez6l | 406.655 | 1.268 | 0.337 | 3.767 | 1.65E-04 | 2.18E-02 | 5 | 112419151 | 112577185 |
| Syt9 | 606.193 | 1.221 | 0.314 | 3.883 | 1.03E-04 | 1.62E-02 | 7 | 107370728 | 107548656 |
| Ndufs5 | 1542.708 | 1.220 | 0.307 | 3.979 | 6.93E-05 | 1.23E-02 | 4 | 123712710 | 123718202 |
| B230110C06Rik | 171.102 | 1.182 | 0.339 | 3.487 | 4.88E-04 | 4.45E-02 | 14 | 15437623 | 15452445 |
| Znf41-ps | 216.610 | 1.110 | 0.316 | 3.511 | 4.46E-04 | 4.22E-02 | 4 | 145825237 | 145829942 |
| Gm5182 | 829.344 | 1.092 | 0.295 | 3.700 | 2.16E-04 | 2.64E-02 | 12 | 10152142 | 10158233 |
| Bud23 | 445.315 | 1.062 | 0.281 | 3.775 | 1.60E-04 | 2.14E-02 | 5 | 135052957 | 135064959 |
| Dync2h1 | 3643.570 | 0.966 | 0.254 | 3.796 | 1.47E-04 | 2.04E-02 | 9 | 6928503 | 7184446 |
| Cds2 | 1781.997 | 0.931 | 0.215 | 4.323 | 1.54E-05 | 4.40E-03 | 2 | 132263148 | 132312050 |
| Qdpr | 1970.364 | 0.905 | 0.219 | 4.134 | 3.57E-05 | 7.96E-03 | 5 | 45434021 | 45450236 |

| | | | | | | | | | |
|---|---|---|---|---|---|---|---|---|---|
| Acsl3 | 4794.627 | 0.832 | 0.232 | 3.583 | 3.39E-04 | 3.63E-02 | 1 | 78657825 | 78707743 |
| Ankrd49 | 672.537 | 0.793 | 0.229 | 3.460 | 5.40E-04 | 4.77E-02 | 9 | 14779618 | 14784856 |
| Specc1l | 8165.141 | 0.714 | 0.178 | 4.007 | 6.16E-05 | 1.16E-02 | 10 | 75212073 | 75312743 |
| Lrp4 | 1923.883 | -0.693 | 0.199 | -3.481 | 5.00E-04 | 4.54E-02 | 2 | 91457511 | 91513779 |
| Josd2 | 2115.387 | -0.756 | 0.215 | -3.524 | 4.25E-04 | 4.09E-02 | 7 | 44467980 | 44471662 |
| Tro | 4755.908 | -0.959 | 0.272 | -3.529 | 4.18E-04 | 4.08E-02 | X | 150645304 | 150657583 |
| Limch1 | 8393.864 | -1.080 | 0.227 | -4.750 | 2.03E-06 | 8.97E-04 | 5 | 66745827 | 67057158 |
| Redrum | 523.589 | -1.138 | 0.308 | -3.697 | 2.18E-04 | 2.64E-02 | 18 | 54422295 | 54453294 |
| Map4k2 | 2618.324 | -1.140 | 0.261 | -4.365 | 1.27E-05 | 3.74E-03 | 19 | 6341135 | 6355615 |
| 9630028B13Rik | 77.721 | -1.336 | 0.385 | -3.469 | 5.22E-04 | 4.68E-02 | 1 | 185429355 | 185441819 |
| Hnrnpul1 | 2593.643 | -1.345 | 0.318 | -4.234 | 2.29E-05 | 6.11E-03 | 7 | 25721165 | 25754757 |
| Selenoi | 3771.249 | -1.356 | 0.191 | -7.100 | 1.24E-12 | 2.29E-09 | 5 | 30232581 | 30272427 |
| Ildr2 | 1123.894 | -1.356 | 0.301 | -4.506 | 6.59E-06 | 2.14E-03 | 1 | 166254139 | 166316823 |
| Nsmce2 | 407.820 | -1.406 | 0.306 | -4.600 | 4.23E-06 | 1.56E-03 | 15 | 59374247 | 59601684 |
| Eya3 | 1903.107 | -1.406 | 0.344 | -4.087 | 4.37E-05 | 9.10E-03 | 4 | 132638987 | 132724765 |
| Dgat1 | 296.433 | -1.423 | 0.407 | -3.492 | 4.80E-04 | 4.40E-02 | 15 | 76502015 | 76511953 |
| Nphs2 | 142.273 | -1.428 | 0.377 | -3.788 | 1.52E-04 | 2.07E-02 | 1 | 156310727 | 156328035 |
| Col11a2 | 1788.933 | -1.450 | 0.396 | -3.661 | 2.51E-04 | 2.90E-02 | 17 | 34039437 | 34066685 |
| Me3 | 731.630 | -1.450 | 0.308 | -4.711 | 2.47E-06 | 1.05E-03 | 7 | 89632392 | 89854359 |
| Cacnb1 | 594.078 | -1.451 | 0.348 | -4.172 | 3.02E-05 | 7.02E-03 | 11 | 98001508 | 98023034 |
| Cd300lg | 197.796 | -1.456 | 0.412 | -3.532 | 4.12E-04 | 4.04E-02 | 11 | 102041509 | 102055620 |
| E230016M11Rik | 77.657 | -1.480 | 0.394 | -3.758 | 1.71E-04 | 2.21E-02 | 6 | 67036599 | 67080654 |
| Slc30a10 | 237.397 | -1.503 | 0.424 | -3.547 | 3.89E-04 | 3.95E-02 | 1 | 185454848 | 185468762 |
| Acat2 | 4657.038 | -1.547 | 0.424 | -3.650 | 2.62E-04 | 3.01E-02 | 17 | 12942890 | 12960747 |
| Pmp2 | 1007.019 | -1.567 | 0.439 | -3.566 | 3.62E-04 | 3.74E-02 | 3 | 10179851 | 10183929 |
| Vxn | 166.596 | -1.571 | 0.422 | -3.721 | 1.98E-04 | 2.47E-02 | 1 | 9601199 | 9627143 |
| Efemp2 | 481.798 | -1.596 | 0.394 | -4.046 | 5.20E-05 | 1.03E-02 | 19 | 5473954 | 5482517 |
| Il3ra | 269.719 | -1.673 | 0.352 | -4.750 | 2.03E-06 | 8.97E-04 | 14 | 14346275 | 14356011 |
| Rpl15-ps6 | 1755.732 | -1.705 | 0.246 | -6.916 | 4.65E-12 | 7.58E-09 | 15 | 52477617 | 52478228 |
| Mkrn3 | 68.533 | -1.714 | 0.437 | -3.924 | 8.70E-05 | 1.43E-02 | 7 | 62417593 | 62420139 |
| Aldh1a7 | 480.332 | -1.720 | 0.425 | -4.052 | 5.08E-05 | 1.02E-02 | 19 | 20692953 | 20727562 |
| Psmc3ip | 223.196 | -1.761 | 0.420 | -4.191 | 2.78E-05 | 6.66E-03 | 11 | 101091823 | 101095436 |
| Vma21 | 3278.865 | -1.767 | 0.466 | -3.795 | 1.48E-04 | 2.04E-02 | X | 71815924 | 71839757 |
| Ubl3 | 7324.551 | -1.782 | 0.436 | -4.084 | 4.42E-05 | 9.12E-03 | 5 | 148504635 | 148552789 |
| Commd8 | 986.760 | -1.827 | 0.432 | -4.230 | 2.34E-05 | 6.11E-03 | 5 | 72156575 | 72168189 |
| Rpl15-ps2 | 567.979 | -1.929 | 0.267 | -7.231 | 4.81E-13 | 9.65E-10 | 9 | 72060937 | 72061549 |
| Car3 | 1154.644 | -1.972 | 0.447 | -4.408 | 1.04E-05 | 3.20E-03 | 3 | 14863512 | 14872523 |
| Sema4g | 881.774 | -2.015 | 0.373 | -5.403 | 6.54E-08 | 4.38E-05 | 19 | 44989101 | 45003397 |
| Ibsp | 2800.520 | -2.046 | 0.592 | -3.458 | 5.44E-04 | 4.78E-02 | 5 | 104299171 | 104311469 |
| Prss41 | 150.501 | -2.051 | 0.353 | -5.812 | 6.16E-09 | 5.44E-06 | 17 | 23836785 | 23844172 |
| Kcmf1 | 1675.111 | -2.146 | 0.475 | -4.516 | 6.31E-06 | 2.08E-03 | 6 | 72841114 | 72899979 |
| Polrmt | 460.600 | -2.223 | 0.373 | -5.966 | 2.44E-09 | 2.24E-06 | 10 | 79736123 | 79746581 |
| Ypel3 | 3151.294 | -2.227 | 0.255 | -8.718 | 2.82E-18 | 1.56E-14 | 7 | 126776955 | 126780514 |
| Tacc3 | 520.417 | -2.283 | 0.572 | -3.991 | 6.57E-05 | 1.21E-02 | 5 | 33658128 | 33678995 |

| | | | | | | | | | |
|---|---|---|---|---|---|---|---|---|---|
| Astn1 | 1366.731 | -2.289 | 0.620 | -3.690 | 2.24E-04 | 2.67E-02 | 1 | 158362273 | 158691781 |
| Itgb3 | 5208.198 | -2.393 | 0.685 | -3.495 | 4.74E-04 | 4.36E-02 | 11 | 104608000 | 104670476 |
| Clu | 2846.333 | -2.516 | 0.374 | -6.727 | 1.74E-11 | 2.47E-08 | 14 | 65968483 | 65981547 |
| Slc7a1 | 348.038 | -2.581 | 0.709 | -3.639 | 2.74E-04 | 3.13E-02 | 5 | 148327410 | 148399904 |
| Trim27 | 474.448 | -2.625 | 0.576 | -4.557 | 5.18E-06 | 1.78E-03 | 13 | 21179445 | 21194724 |
| Cthrc1 | 267.537 | -2.758 | 0.326 | -8.454 | 2.81E-17 | 1.03E-13 | 15 | 39076932 | 39087121 |
| Tnc | 2178.044 | -2.781 | 0.766 | -3.632 | 2.82E-04 | 3.20E-02 | 4 | 63959785 | 64047015 |
| Gm8189 | 65.918 | -2.860 | 0.830 | -3.446 | 5.69E-04 | 4.90E-02 | 19 | 8473313 | 8541798 |
| Fabp4 | 1365.493 | -2.898 | 0.659 | -4.395 | 1.11E-05 | 3.31E-03 | 3 | 10204088 | 10208576 |
| Bst1 | 286.639 | -2.963 | 0.828 | -3.581 | 3.43E-04 | 3.64E-02 | 5 | 43818885 | 43843986 |
| Nrros | 415.335 | -2.981 | 0.845 | -3.526 | 4.21E-04 | 4.08E-02 | 16 | 32142785 | 32165594 |
| Arhgap4 | 875.332 | -3.014 | 0.820 | -3.674 | 2.39E-04 | 2.80E-02 | X | 73891442 | 73921870 |
| Dnah14 | 106.989 | -3.059 | 0.571 | -5.354 | 8.59E-08 | 5.57E-05 | 1 | 181576559 | 181815774 |
| Col1a1 | 38009.926 | -3.067 | 0.774 | -3.960 | 7.49E-05 | 1.29E-02 | 11 | 94936224 | 94953042 |
| Fn1 | 1648.001 | -3.197 | 0.715 | -4.473 | 7.71E-06 | 2.47E-03 | 1 | 71585520 | 71653200 |
| Slco4c1 | 296.634 | -3.204 | 0.900 | -3.561 | 3.69E-04 | 3.79E-02 | 1 | 96816270 | 96872171 |
| Eme1 | 104.972 | -3.270 | 0.932 | -3.510 | 4.48E-04 | 4.22E-02 | 11 | 94644996 | 94653964 |
| Pold1 | 215.930 | -3.322 | 0.865 | -3.839 | 1.24E-04 | 1.83E-02 | 7 | 44532746 | 44548849 |
| Ckap2l | 625.621 | -3.338 | 0.943 | -3.541 | 3.99E-04 | 3.97E-02 | 2 | 129268210 | 129297212 |
| Zdhhc9 | 1105.501 | -3.394 | 0.398 | -8.524 | 1.54E-17 | 6.79E-14 | X | 48171969 | 48208878 |
| Acp5 | 1959.898 | -3.410 | 0.877 | -3.890 | 1.00E-04 | 1.60E-02 | 9 | 22126727 | 22135711 |
| Ncaph | 425.221 | -3.443 | 0.973 | -3.539 | 4.01E-04 | 3.97E-02 | 2 | 127103809 | 127133954 |
| Ptprv | 332.045 | -3.564 | 0.896 | -3.978 | 6.96E-05 | 1.23E-02 | 1 | 135108497 | 135132594 |
| Gp1ba | 484.597 | -3.573 | 0.869 | -4.112 | 3.92E-05 | 8.39E-03 | 11 | 70639122 | 70642036 |
| Postn | 5636.422 | -3.589 | 0.690 | -5.203 | 1.96E-07 | 1.11E-04 | 3 | 54361109 | 54391037 |
| Nuf2 | 397.997 | -3.593 | 1.021 | -3.519 | 4.33E-04 | 4.15E-02 | 1 | 169497934 | 169531464 |
| Mybl2 | 509.354 | -3.604 | 0.947 | -3.806 | 1.41E-04 | 1.99E-02 | 2 | 163054687 | 163084688 |
| Gm47015 | 472.273 | -3.606 | 0.751 | -4.804 | 1.55E-06 | 7.29E-04 | 10 | 79412962 | 79417905 |
| Gm28035 | 1306.621 | -3.613 | 0.935 | -3.862 | 1.12E-04 | 1.75E-02 | 2 | 30240567 | 30282148 |
| Adipoq | 216.032 | -3.624 | 0.605 | -5.989 | 2.11E-09 | 2.12E-06 | 16 | 23146536 | 23158028 |
| Mmp13 | 2231.660 | -3.797 | 1.084 | -3.503 | 4.59E-04 | 4.25E-02 | 9 | 7272514 | 7283331 |
| Cphx1 | 14.153 | -3.835 | 1.111 | -3.451 | 5.58E-04 | 4.85E-02 | 14 | 25938304 | 25956356 |
| Tnni1 | 177.797 | -3.871 | 0.783 | -4.947 | 7.56E-07 | 3.88E-04 | 1 | 135779434 | 135810989 |
| Ppp1r9b | 3278.625 | -3.941 | 0.862 | -4.571 | 4.85E-06 | 1.75E-03 | 11 | 94991035 | 95006899 |
| Rad21 | 33.906 | -3.970 | 1.001 | -3.967 | 7.28E-05 | 1.26E-02 | 15 | 51962240 | 51991747 |
| Tubb1 | 869.510 | -4.059 | 1.151 | -3.526 | 4.22E-04 | 4.08E-02 | 2 | 174450695 | 174457882 |
| Podnl1 | 136.668 | -4.212 | 1.011 | -4.168 | 3.07E-05 | 7.05E-03 | 8 | 84125989 | 84132527 |
| F5 | 841.340 | -4.218 | 1.176 | -3.587 | 3.35E-04 | 3.62E-02 | 1 | 164151838 | 164220277 |
| Nox4 | 68.501 | -4.361 | 0.931 | -4.685 | 2.81E-06 | 1.13E-03 | 7 | 87246096 | 87398710 |
| Arhgap27os3 | 48.632 | -4.362 | 1.256 | -3.474 | 5.12E-04 | 4.61E-02 | 11 | 103344753 | 103350124 |
| Mrgprb4 | 155.872 | -4.366 | 1.074 | -4.065 | 4.80E-05 | 9.81E-03 | 7 | 48198070 | 48199288 |
| Gp1bb | 417.295 | -4.429 | 1.208 | -3.666 | 2.46E-04 | 2.86E-02 | 16 | 18620317 | 18622403 |
| Hcfc2 | 466.427 | -4.471 | 0.364 | -12.286 | 1.08E-34 | 2.38E-30 | 10 | 82696160 | 82742428 |
| Pygl | 586.556 | -4.647 | 1.018 | -4.565 | 5.00E-06 | 1.77E-03 | 12 | 70190811 | 70231488 |

| | | | | | | | | | |
|---|---|---|---|---|---|---|---|---|---|
| Mirt1 | 202.548 | -4.698 | 0.988 | -4.755 | 1.98E-06 | 8.97E-04 | 19 | 53443230 | 53464796 |
| Gm13237 | 61.697 | -4.731 | 1.283 | -3.688 | 2.26E-04 | 2.68E-02 | 4 | 146564453 | 146565087 |
| Slc40a1 | 96.800 | -4.811 | 0.919 | -5.235 | 1.65E-07 | 9.81E-05 | 1 | 45908068 | 45926523 |
| Igsf23 | 22.242 | -4.861 | 1.315 | -3.696 | 2.19E-04 | 2.64E-02 | 7 | 19937305 | 19950756 |
| Gm34680 | 59.793 | -4.955 | 1.243 | -3.987 | 6.68E-05 | 1.21E-02 | 16 | 32550211 | 32553708 |
| Rpl32l | 113.798 | -5.061 | 0.732 | -6.911 | 4.81E-12 | 7.58E-09 | 10 | 20208490 | 20208888 |
| Ramac | 190.990 | -5.069 | 1.421 | -3.566 | 3.62E-04 | 3.74E-02 | 7 | 81762925 | 81769491 |
| Bglap2 | 625.550 | -5.191 | 0.951 | -5.460 | 4.77E-08 | 3.29E-05 | 3 | 88377736 | 88378699 |
| Selp | 268.929 | -5.279 | 1.341 | -3.936 | 8.29E-05 | 1.39E-02 | 1 | 164115264 | 164150026 |
| Gm5734 | 32.688 | -5.302 | 1.121 | -4.729 | 2.26E-06 | 9.77E-04 | 7 | 47915322 | 47916234 |
| 1700085C21Rik | 35.025 | -5.344 | 1.358 | -3.934 | 8.36E-05 | 1.39E-02 | 12 | 82932520 | 82939155 |
| Gm37811 | 61.467 | -5.499 | 1.432 | -3.839 | 1.23E-04 | 1.83E-02 | 2 | 13009886 | 13012486 |
| Gm14719 | 47.999 | -5.573 | 0.829 | -6.722 | 1.79E-11 | 2.47E-08 | X | 49660960 | 49661299 |
| Tmprss3 | 99.930 | -5.639 | 1.486 | -3.794 | 1.48E-04 | 2.04E-02 | 17 | 31179265 | 31198977 |
| Gm10143 | 255.337 | -5.716 | 1.415 | -4.040 | 5.36E-05 | 1.05E-02 | 19 | 10198368 | 10202384 |
| Ifi214 | 54.755 | -5.836 | 1.401 | -4.166 | 3.10E-05 | 7.06E-03 | 1 | 173519717 | 173535957 |
| Mir142b | 38.464 | -5.950 | 1.408 | -4.226 | 2.37E-05 | 6.11E-03 | 11 | 87756841 | 87756955 |
| Dlx5 | 57.579 | -6.038 | 1.424 | -4.240 | 2.23E-05 | 6.08E-03 | 6 | 6877805 | 6882085 |
| Vav1 | 196.416 | -6.045 | 1.460 | -4.142 | 3.45E-05 | 7.77E-03 | 17 | 57279100 | 57328031 |
| D130058E05Rik | 39.715 | -6.129 | 1.348 | -4.547 | 5.44E-06 | 1.85E-03 | 1 | 89930234 | 89933290 |
| Gm10320 | 124.432 | -6.184 | 1.470 | -4.206 | 2.59E-05 | 6.43E-03 | 13 | 98489348 | 98492001 |
| Hand2 | 63.978 | -6.212 | 1.635 | -3.799 | 1.45E-04 | 2.04E-02 | 8 | 57320983 | 57324633 |
| mt-Ts2 | 51.323 | -6.425 | 1.572 | -4.087 | 4.37E-05 | 9.10E-03 | MT | 11613 | 11671 |
| Uhrf1 | 705.449 | -6.464 | 0.976 | -6.623 | 3.53E-11 | 4.32E-08 | 17 | 56303321 | 56323486 |
| Gm44689 | 11.833 | -6.767 | 1.697 | -3.988 | 6.65E-05 | 1.21E-02 | 7 | 71681775 | 71688447 |
| Gm8885 | 12.419 | -6.837 | 1.656 | -4.129 | 3.64E-05 | 8.03E-03 | 1 | 72087857 | 72088454 |
| Gm27553 | 13.019 | -6.905 | 1.634 | -4.225 | 2.39E-05 | 6.11E-03 | 5 | 100421968 | 100422041 |
| Ighv1-14 | 13.609 | -6.973 | 1.851 | -3.766 | 1.66E-04 | 2.18E-02 | 12 | 114646572 | 114647056 |
| Cyp2j13 | 16.944 | -7.286 | 1.597 | -4.563 | 5.05E-06 | 1.77E-03 | 4 | 96027534 | 96077546 |
| Loxl4 | 16.996 | -7.289 | 1.613 | -4.519 | 6.23E-06 | 2.08E-03 | 19 | 42593982 | 42612813 |
| Emx2 | 18.309 | -7.400 | 1.594 | -4.643 | 3.44E-06 | 1.31E-03 | 19 | 59458372 | 59465357 |
| Igkv3-2 | 21.935 | -7.661 | 2.020 | -3.793 | 1.49E-04 | 2.04E-02 | 6 | 70698449 | 70699067 |
| Gm8113 | 23.743 | -7.775 | 1.664 | -4.672 | 2.98E-06 | 1.17E-03 | 14 | 43925390 | 43933411 |
| Oscar | 31.643 | -8.190 | 1.764 | -4.643 | 3.44E-06 | 1.31E-03 | 7 | 3609813 | 3616157 |
| Gm20649 | 34.547 | -8.313 | 2.406 | -3.455 | 5.51E-04 | 4.80E-02 | 9 | 74848463 | 74891476 |
| Gm47308 | 248.862 | -8.673 | 2.154 | -4.027 | 5.66E-05 | 1.09E-02 | 9 | 64726624 | 64726730 |
| 4930405J17Rik | 74.795 | -9.427 | 1.420 | -6.640 | 3.14E-11 | 4.08E-08 | 10 | 20199240 | 20201838 |
| Fcrla | 86.095 | -9.632 | 1.537 | -6.266 | 3.70E-10 | 4.30E-07 | 1 | 170917576 | 170927583 |
| Heph | 106.281 | -9.936 | 1.321 | -7.521 | 5.43E-14 | 1.33E-10 | X | 96455359 | 96574485 |

Table 5-3. Relative gene expression in female responder and nonresponder mice.

| Gene | baseMean | log2FC | lfcSE | stat | pvalue | padj | chr | start | end |
|---|---|---|---|---|---|---|---|---|---|
| Elovl2 | 247.137 | 11.132 | 3.112 | 3.577 | 3.47E-04 | 2.16E-02 | 13 | 41182381 | 41220405 |
| Usp13 | 139.753 | 10.312 | 1.619 | 6.369 | 1.90E-10 | 1.04E-07 | 3 | 32817546 | 32938071 |
| Onecut2 | 359.243 | 9.824 | 1.300 | 7.560 | 4.04E-14 | 3.68E-11 | 18 | 64340020 | 64400987 |
| Casp8 | 93.601 | 9.731 | 1.831 | 5.316 | 1.06E-07 | 3.12E-05 | 1 | 58795374 | 58847503 |
| Nod2 | 63.710 | 9.178 | 1.369 | 6.705 | 2.01E-11 | 1.32E-08 | 8 | 88647315 | 88688474 |
| Trpc2 | 55.658 | 8.980 | 1.303 | 6.891 | 5.54E-12 | 3.92E-09 | 7 | 102083116 | 102096396 |
| Igkv3-7 | 42.949 | 8.608 | 1.985 | 4.335 | 1.45E-05 | 2.40E-03 | 6 | 70607437 | 70608036 |
| Rest | 268.798 | 8.415 | 0.850 | 9.904 | 3.99E-23 | 1.27E-19 | 5 | 77265491 | 77286432 |
| Psg16 | 35.653 | 8.339 | 2.500 | 3.336 | 8.50E-04 | 3.90E-02 | 7 | 17074040 | 17133450 |
| Gm28030 | 28.633 | 8.023 | 1.387 | 5.783 | 7.33E-09 | 2.98E-06 | 5 | 100429101 | 100429238 |
| Igkv16-104 | 28.364 | 8.007 | 1.952 | 4.103 | 4.08E-05 | 5.16E-03 | 6 | 68425575 | 68426072 |
| Kif28 | 28.341 | 8.007 | 1.378 | 5.809 | 6.28E-09 | 2.67E-06 | 1 | 179695297 | 179745271 |
| Igkv6-20 | 26.283 | 7.899 | 1.870 | 4.224 | 2.40E-05 | 3.48E-03 | 6 | 70335841 | 70336507 |
| Med27 | 249.615 | 7.796 | 2.137 | 3.647 | 2.65E-04 | 1.85E-02 | 2 | 29346819 | 29524793 |
| Afg1l | 22.738 | 7.688 | 1.985 | 3.874 | 1.07E-04 | 1.02E-02 | 10 | 42312585 | 42478565 |
| Rad51c | 70.807 | 6.519 | 1.006 | 6.479 | 9.22E-11 | 5.34E-08 | 11 | 87376645 | 87404954 |
| Ighg2c | 3181.920 | 6.223 | 1.736 | 3.585 | 3.37E-04 | 2.14E-02 | 12 | 113285325 | 113288932 |
| Gm27640 | 40.189 | 6.046 | 1.597 | 3.786 | 1.53E-04 | 1.31E-02 | 11 | 3647179 | 3647325 |
| Ighv14-3 | 46.807 | 5.807 | 1.380 | 4.208 | 2.57E-05 | 3.59E-03 | 12 | 114059845 | 114060321 |
| Gm27671 | 33.640 | 5.784 | 1.619 | 3.572 | 3.55E-04 | 2.19E-02 | 16 | 14128566 | 14128631 |
| Fbn1 | 620.106 | 5.623 | 1.518 | 3.704 | 2.13E-04 | 1.62E-02 | 2 | 125300594 | 125507993 |
| Snhg3 | 29.369 | 5.608 | 1.709 | 3.281 | 1.04E-03 | 4.32E-02 | 4 | 132348014 | 132353686 |
| Igkv4-53 | 47.916 | 5.552 | 1.552 | 3.578 | 3.47E-04 | 2.16E-02 | 6 | 69648824 | 69649363 |
| Irf5 | 67.400 | 5.362 | 0.960 | 5.583 | 2.37E-08 | 8.71E-06 | 6 | 29526625 | 29541871 |
| Sgsh | 25.833 | 5.284 | 1.587 | 3.330 | 8.69E-04 | 3.94E-02 | 11 | 119343425 | 119355536 |
| Gm14165 | 96.484 | 4.780 | 1.142 | 4.184 | 2.86E-05 | 3.89E-03 | 2 | 152353615 | 152354566 |
| Gm6166 | 541.470 | 4.564 | 0.483 | 9.456 | 3.20E-21 | 8.74E-18 | 9 | 57483963 | 57484371 |
| Prpf8 | 880.014 | 4.231 | 1.235 | 3.426 | 6.13E-04 | 3.09E-02 | 11 | 75486816 | 75509449 |
| Gm8730 | 1393.835 | 4.073 | 0.751 | 5.427 | 5.74E-08 | 1.92E-05 | 8 | 102864845 | 102865798 |
| Tor3a | 138.063 | 3.529 | 0.759 | 4.652 | 3.29E-06 | 6.76E-04 | 1 | 156653617 | 156674356 |
| Lmbr1 | 441.717 | 3.508 | 0.730 | 4.808 | 1.53E-06 | 3.64E-04 | 5 | 29229802 | 29378390 |
| Hcfc2 | 217.767 | 3.275 | 1.018 | 3.219 | 1.29E-03 | 4.98E-02 | 10 | 82696160 | 82742428 |
| Gm38182 | 125.299 | 3.214 | 0.952 | 3.374 | 7.40E-04 | 3.52E-02 | 18 | 37823879 | 37840382 |
| Slc26a10 | 348.497 | 3.175 | 0.858 | 3.702 | 2.14E-04 | 1.63E-02 | 10 | 127171393 | 127180645 |
| Mpp2 | 98.605 | 3.036 | 0.784 | 3.875 | 1.07E-04 | 1.02E-02 | 11 | 102057015 | 102088515 |
| Znrd1as | 148.697 | 2.818 | 0.869 | 3.243 | 1.18E-03 | 4.68E-02 | 17 | 36958592 | 36965625 |
| Eps8 | 118.658 | 2.735 | 0.807 | 3.389 | 7.02E-04 | 3.38E-02 | 6 | 137477245 | 137654876 |
| Slc7a1 | 520.639 | 2.676 | 0.728 | 3.676 | 2.37E-04 | 1.72E-02 | 5 | 148327410 | 148399904 |
| Hdac2 | 1621.736 | 2.664 | 0.694 | 3.842 | 1.22E-04 | 1.12E-02 | 10 | 36974544 | 37001889 |
| Rnf115 | 186.725 | 2.629 | 0.749 | 3.511 | 4.47E-04 | 2.55E-02 | 3 | 96727664 | 96791638 |
| Get1 | 241.905 | 2.614 | 0.330 | 7.925 | 2.29E-15 | 2.92E-12 | 16 | 96145407 | 96157852 |
| Hmgcr | 2072.432 | 2.521 | 0.175 | 14.431 | 3.32E-47 | 2.11E-43 | 13 | 96648967 | 96670936 |
| Hace1 | 595.103 | 2.322 | 0.423 | 5.490 | 4.01E-08 | 1.37E-05 | 10 | 45577829 | 45712345 |
| Clu | 5335.855 | 2.278 | 0.482 | 4.722 | 2.34E-06 | 5.20E-04 | 14 | 65968483 | 65981547 |
| Csgalnact2 | 90.134 | 2.202 | 0.480 | 4.590 | 4.43E-06 | 8.64E-04 | 6 | 118107452 | 118139140 |
| Arhgap27 | 871.655 | 2.154 | 0.499 | 4.313 | 1.61E-05 | 2.52E-03 | 11 | 103331497 | 103363692 |
| A230020J21Rik | 211.441 | 2.121 | 0.264 | 8.042 | 8.85E-16 | 1.21E-12 | 1 | 191025350 | 191029781 |

| | | | | | | | | | |
|---|---|---|---|---|---|---|---|---|---|
| Rnf146 | 3773.304 | 1.988 | 0.162 | 12.302 | 8.85E-35 | 3.38E-31 | 10 | 29344176 | 29362442 |
| Coro7 | 420.593 | 1.969 | 0.347 | 5.670 | 1.42E-08 | 5.44E-06 | 16 | 4626133 | 4679777 |
| Ccl8 | 129.424 | 1.949 | 0.599 | 3.253 | 1.14E-03 | 4.61E-02 | 11 | 82115185 | 82116799 |
| Mras | 423.016 | 1.879 | 0.490 | 3.830 | 1.28E-04 | 1.16E-02 | 9 | 99385420 | 99437381 |
| Tubb2a-ps2 | 88.295 | 1.878 | 0.569 | 3.299 | 9.72E-04 | 4.19E-02 | 12 | 11881325 | 11882663 |
| Rcan3 | 405.521 | 1.818 | 0.310 | 5.856 | 4.74E-09 | 2.16E-06 | 4 | 135412308 | 135433853 |
| Tfg | 1062.828 | 1.799 | 0.203 | 8.872 | 7.20E-19 | 1.38E-15 | 16 | 56690332 | 56717450 |
| Nts | 622.797 | 1.794 | 0.457 | 3.922 | 8.76E-05 | 8.76E-03 | 10 | 102481756 | 102490486 |
| 1700101I11Rik | 2256.164 | 1.787 | 0.423 | 4.222 | 2.43E-05 | 3.49E-03 | 6 | 129531950 | 129533817 |
| Gpx1 | 7626.209 | 1.751 | 0.490 | 3.573 | 3.53E-04 | 2.18E-02 | 9 | 108338903 | 108340343 |
| Erdr1 | 360.341 | 1.730 | 0.473 | 3.660 | 2.52E-04 | 1.79E-02 | X | 170009659 | 170019281 |
| Smad5 | 444.338 | 1.702 | 0.403 | 4.225 | 2.38E-05 | 3.48E-03 | 13 | 56703010 | 56742377 |
| Zfp667 | 408.009 | 1.679 | 0.504 | 3.333 | 8.58E-04 | 3.92E-02 | 7 | 6286579 | 6307883 |
| Gm47870 | 88.702 | 1.675 | 0.422 | 3.970 | 7.18E-05 | 7.60E-03 | 12 | 65074990 | 65076315 |
| Gm5855 | 169.734 | 1.657 | 0.449 | 3.687 | 2.27E-04 | 1.69E-02 | 3 | 130929366 | 130930670 |
| Actr3 | 11880.609 | 1.617 | 0.412 | 3.926 | 8.65E-05 | 8.69E-03 | 1 | 125392905 | 125435727 |
| Ndufs5 | 2678.461 | 1.603 | 0.264 | 6.074 | 1.25E-09 | 6.45E-07 | 4 | 123712710 | 123718202 |
| Gm10434 | 219.142 | 1.579 | 0.310 | 5.090 | 3.58E-07 | 9.90E-05 | 11 | 59208246 | 59208720 |
| Sigmar1 | 1237.442 | 1.525 | 0.454 | 3.356 | 7.90E-04 | 3.70E-02 | 4 | 41738493 | 41756157 |
| Slc35e2 | 1632.311 | 1.501 | 0.415 | 3.621 | 2.94E-04 | 1.97E-02 | 4 | 155601416 | 155623340 |
| H2-K2 | 133.851 | 1.500 | 0.436 | 3.438 | 5.86E-04 | 2.99E-02 | 17 | 33974659 | 33978827 |
| Bbc3 | 97.765 | 1.453 | 0.427 | 3.404 | 6.64E-04 | 3.26E-02 | 7 | 16308393 | 16318205 |
| Vps52 | 286.188 | 1.453 | 0.323 | 4.494 | 6.97E-06 | 1.26E-03 | 17 | 33955812 | 33967035 |
| Ech1 | 449.798 | 1.397 | 0.406 | 3.439 | 5.83E-04 | 2.99E-02 | 7 | 28825217 | 28832247 |
| Hmox2 | 1883.659 | 1.215 | 0.225 | 5.402 | 6.59E-08 | 2.13E-05 | 16 | 4726361 | 4766742 |
| Rb1cc1 | 1014.625 | 1.207 | 0.312 | 3.866 | 1.11E-04 | 1.04E-02 | 1 | 6206197 | 6276648 |
| Gm6767 | 512.109 | 1.197 | 0.324 | 3.698 | 2.18E-04 | 1.64E-02 | 16 | 51362452 | 51363832 |
| Ift122 | 1206.828 | 1.170 | 0.353 | 3.310 | 9.32E-04 | 4.11E-02 | 6 | 115853470 | 115926699 |
| 8430429K09Rik | 361.518 | 1.162 | 0.251 | 4.624 | 3.77E-06 | 7.50E-04 | 11 | 3452365 | 3479831 |
| Zdhhc24 | 1988.467 | 1.161 | 0.199 | 5.827 | 5.64E-09 | 2.51E-06 | 19 | 4878668 | 4885397 |
| Tifa | 970.816 | 1.155 | 0.194 | 5.940 | 2.85E-09 | 1.36E-06 | 3 | 127789805 | 127832164 |
| Cul2 | 6312.943 | 1.108 | 0.199 | 5.555 | 2.77E-08 | 9.81E-06 | 18 | 3382988 | 3436377 |
| Usp29 | 274.462 | 1.093 | 0.260 | 4.212 | 2.53E-05 | 3.55E-03 | 7 | 6730578 | 6967219 |
| Grn | 381.984 | 1.089 | 0.325 | 3.351 | 8.05E-04 | 3.75E-02 | 11 | 102430315 | 102437048 |
| Wdr1 | 1112.718 | 1.083 | 0.260 | 4.160 | 3.18E-05 | 4.14E-03 | 5 | 38526813 | 38563221 |
| Kcnj14 | 422.243 | 1.072 | 0.282 | 3.796 | 1.47E-04 | 1.28E-02 | 7 | 45816460 | 45824782 |
| Mir124a-1hg | 803.432 | 1.060 | 0.317 | 3.344 | 8.26E-04 | 3.83E-02 | 14 | 64587331 | 64593961 |
| Tpi-rs11 | 137.343 | 1.057 | 0.327 | 3.233 | 1.23E-03 | 4.79E-02 | 6 | 127183409 | 127183718 |
| Gm7292 | 25878.825 | 1.037 | 0.111 | 9.367 | 7.47E-21 | 1.78E-17 | 6 | 117716342 | 117718868 |
| Gm6210 | 2806.075 | 1.014 | 0.300 | 3.378 | 7.30E-04 | 3.48E-02 | 6 | 75296118 | 75298059 |
| Gins2 | 603.283 | 0.998 | 0.304 | 3.281 | 1.03E-03 | 4.32E-02 | 8 | 120578633 | 120589304 |
| Calm1 | 8377.209 | 0.985 | 0.272 | 3.618 | 2.97E-04 | 1.97E-02 | 12 | 100199435 | 100209814 |
| Gm14418 | 484.982 | 0.960 | 0.206 | 4.652 | 3.29E-06 | 6.76E-04 | 2 | 177387124 | 177398320 |
| B3gnt5 | 1015.943 | 0.941 | 0.236 | 3.994 | 6.50E-05 | 7.18E-03 | 16 | 19760208 | 19772753 |
| Nrxn2 | 10571.144 | 0.929 | 0.282 | 3.296 | 9.80E-04 | 4.20E-02 | 19 | 6418731 | 6544169 |
| Adat2 | 571.227 | 0.922 | 0.221 | 4.173 | 3.01E-05 | 3.94E-03 | 10 | 13552894 | 13563376 |
| Ccdc115 | 2113.977 | 0.918 | 0.172 | 5.324 | 1.01E-07 | 3.03E-05 | 1 | 34436670 | 34439672 |
| Atp1a3 | 51580.466 | 0.892 | 0.267 | 3.339 | 8.40E-04 | 3.87E-02 | 7 | 24978167 | 25005958 |
| 1700069B07Rik | 279.678 | 0.866 | 0.251 | 3.447 | 5.67E-04 | 2.95E-02 | 7 | 121707075 | 121708013 |
| Gm16580 | 169.031 | 0.853 | 0.256 | 3.329 | 8.72E-04 | 3.95E-02 | 17 | 26269337 | 26270384 |

| | | | | | | | | | |
|---|---|---|---|---|---|---|---|---|---|
| Gm14322 | 323.373 | 0.843 | 0.236 | 3.566 | 3.63E-04 | 2.23E-02 | 2 | 177759288 | 177770472 |
| Cacna2d2 | 4019.906 | 0.843 | 0.207 | 4.076 | 4.58E-05 | 5.53E-03 | 9 | 107399612 | 107529343 |
| Lgi1 | 3075.494 | 0.843 | 0.196 | 4.290 | 1.78E-05 | 2.73E-03 | 19 | 38264536 | 38312214 |
| Ift140 | 2152.867 | 0.838 | 0.211 | 3.965 | 7.35E-05 | 7.67E-03 | 17 | 25016085 | 25099495 |
| Fndc3a | 1056.625 | 0.834 | 0.217 | 3.845 | 1.20E-04 | 1.11E-02 | 14 | 72537946 | 72710003 |
| Pop4 | 2044.410 | 0.816 | 0.248 | 3.290 | 1.00E-03 | 4.23E-02 | 7 | 38261996 | 38271423 |
| Gm49521 | 828.728 | 0.775 | 0.202 | 3.838 | 1.24E-04 | 1.13E-02 | 16 | 16349203 | 16359011 |
| Ubr4 | 36533.736 | 0.772 | 0.207 | 3.721 | 1.98E-04 | 1.55E-02 | 4 | 139352609 | 139489588 |
| Gm13446 | 579.996 | 0.764 | 0.222 | 3.442 | 5.78E-04 | 2.98E-02 | 2 | 35549538 | 35558702 |
| Rab40c | 1629.128 | 0.745 | 0.214 | 3.490 | 4.84E-04 | 2.66E-02 | 17 | 25882114 | 25919727 |
| Atp2b2 | 21748.824 | 0.741 | 0.222 | 3.333 | 8.60E-04 | 3.92E-02 | 6 | 113743831 | 114042613 |
| Khk | 2114.778 | 0.735 | 0.146 | 5.049 | 4.44E-07 | 1.19E-04 | 5 | 30921431 | 30931248 |
| Aurkaip1 | 4322.033 | 0.728 | 0.205 | 3.557 | 3.75E-04 | 2.28E-02 | 4 | 155831272 | 155833130 |
| Rasa3 | 4595.411 | 0.713 | 0.202 | 3.523 | 4.26E-04 | 2.49E-02 | 8 | 13566948 | 13677603 |
| Dynll2 | 84945.759 | 0.702 | 0.104 | 6.758 | 1.40E-11 | 9.56E-09 | 11 | 87979525 | 87987533 |
| Gm49678 | 731.790 | 0.687 | 0.211 | 3.261 | 1.11E-03 | 4.52E-02 | 16 | 75388544 | 75447293 |
| 9230112E08Rik | 938.644 | 0.684 | 0.197 | 3.469 | 5.23E-04 | 2.80E-02 | 2 | 180983924 | 180987275 |
| Homer1 | 1746.839 | 0.681 | 0.148 | 4.613 | 3.97E-06 | 7.83E-04 | 13 | 93299635 | 93405129 |
| Crtac1 | 3518.775 | 0.671 | 0.182 | 3.678 | 2.35E-04 | 1.72E-02 | 19 | 42268287 | 42431785 |
| Hs6st2 | 403.423 | 0.666 | 0.205 | 3.245 | 1.17E-03 | 4.67E-02 | X | 51387212 | 51681856 |
| Gm26673 | 1180.214 | 0.661 | 0.191 | 3.460 | 5.40E-04 | 2.86E-02 | 6 | 126718684 | 126741601 |
| Acads | 306.271 | 0.661 | 0.200 | 3.305 | 9.50E-04 | 4.16E-02 | 5 | 115110299 | 115119346 |
| Gm10874 | 1110.393 | 0.659 | 0.188 | 3.497 | 4.70E-04 | 2.62E-02 | 5 | 138363719 | 138388287 |
| Tmem108 | 1329.386 | 0.656 | 0.189 | 3.477 | 5.07E-04 | 2.76E-02 | 9 | 103482947 | 103761837 |
| Fam131a | 1010.742 | 0.656 | 0.165 | 3.970 | 7.20E-05 | 7.60E-03 | 16 | 20693241 | 20703048 |
| Klhl25 | 1389.281 | 0.649 | 0.187 | 3.465 | 5.29E-04 | 2.83E-02 | 7 | 75848310 | 75874131 |
| A730098A19Rik | 303.135 | 0.641 | 0.192 | 3.341 | 8.34E-04 | 3.86E-02 | 8 | 125897914 | 125908144 |
| Ndufb8 | 3723.919 | 0.641 | 0.181 | 3.547 | 3.90E-04 | 2.34E-02 | 19 | 44548572 | 44555440 |
| Gm5526 | 279.157 | 0.634 | 0.184 | 3.453 | 5.54E-04 | 2.91E-02 | 1 | 45857332 | 45857754 |
| Ssx2ip | 5537.597 | 0.630 | 0.181 | 3.478 | 5.05E-04 | 2.76E-02 | 3 | 146404642 | 146440144 |
| Tbc1d24 | 2057.574 | 0.629 | 0.145 | 4.331 | 1.48E-05 | 2.40E-03 | 17 | 24175431 | 24205562 |
| Dhx30 | 7783.575 | 0.627 | 0.179 | 3.502 | 4.62E-04 | 2.62E-02 | 9 | 110084320 | 110117830 |
| Cyfip2 | 1372.471 | 0.622 | 0.160 | 3.886 | 1.02E-04 | 9.90E-03 | 11 | 46193850 | 46312859 |
| Pex26 | 617.962 | 0.619 | 0.163 | 3.792 | 1.50E-04 | 1.28E-02 | 6 | 121183667 | 121198837 |
| Tmem59 | 2324.430 | 0.600 | 0.181 | 3.309 | 9.36E-04 | 4.12E-02 | 4 | 107178399 | 107200996 |
| Tppp3 | 8841.233 | 0.590 | 0.181 | 3.251 | 1.15E-03 | 4.61E-02 | 8 | 105467493 | 105471526 |
| Ubb-ps | 25506.554 | 0.580 | 0.154 | 3.771 | 1.63E-04 | 1.35E-02 | 14 | 46084028 | 46085000 |
| Klhl32 | 1021.189 | 0.572 | 0.139 | 4.102 | 4.10E-05 | 5.16E-03 | 4 | 24612554 | 24851124 |
| Fry | 12803.414 | 0.568 | 0.159 | 3.574 | 3.51E-04 | 2.18E-02 | 5 | 150118645 | 150497753 |
| Zfp618 | 1095.692 | 0.546 | 0.156 | 3.499 | 4.67E-04 | 2.62E-02 | 4 | 62965573 | 63139708 |
| Htr1a | 752.461 | 0.539 | 0.155 | 3.484 | 4.94E-04 | 2.70E-02 | 13 | 105443639 | 105448122 |
| Acot8 | 605.879 | 0.538 | 0.145 | 3.712 | 2.06E-04 | 1.60E-02 | 2 | 164792765 | 164804882 |
| Pde2a | 7311.832 | 0.537 | 0.106 | 5.061 | 4.16E-07 | 1.14E-04 | 7 | 101421691 | 101512827 |
| Arhgef40 | 3152.685 | 0.534 | 0.156 | 3.412 | 6.44E-04 | 3.20E-02 | 14 | 51984719 | 52006251 |
| Mcc | 1762.429 | 0.525 | 0.136 | 3.872 | 1.08E-04 | 1.02E-02 | 18 | 44425060 | 44812182 |
| Gm14295 | 1418.137 | 0.525 | 0.160 | 3.278 | 1.04E-03 | 4.35E-02 | 2 | 176798612 | 176811223 |
| Unc13a | 4205.278 | 0.509 | 0.156 | 3.274 | 1.06E-03 | 4.39E-02 | 8 | 71624417 | 71671757 |
| Syne1 | 22228.636 | 0.509 | 0.109 | 4.693 | 2.69E-06 | 5.84E-04 | 10 | 5020917 | 5551482 |
| Runx1t1 | 1625.843 | 0.507 | 0.146 | 3.464 | 5.33E-04 | 2.84E-02 | 4 | 13743436 | 13893649 |
| Lancl3 | 2200.908 | 0.501 | 0.101 | 4.981 | 6.32E-07 | 1.63E-04 | X | 9199902 | 9268085 |

| | | | | | | | | | |
|---|---|---|---|---|---|---|---|---|---|
| Stac2 | 9246.559 | 0.499 | 0.154 | 3.245 | 1.17E-03 | 4.67E-02 | 11 | 98036623 | 98053462 |
| Calb2 | 9988.351 | 0.495 | 0.118 | 4.178 | 2.94E-05 | 3.90E-03 | 8 | 110137502 | 110168210 |
| Cntnap1 | 21779.151 | 0.487 | 0.121 | 4.010 | 6.06E-05 | 6.94E-03 | 11 | 101170523 | 101190724 |
| Ints1 | 8249.660 | 0.485 | 0.130 | 3.726 | 1.95E-04 | 1.53E-02 | 5 | 139751282 | 139775674 |
| Scrt1 | 20956.013 | 0.484 | 0.148 | 3.283 | 1.03E-03 | 4.30E-02 | 15 | 76516203 | 76522499 |
| Gm15459 | 53256.563 | 0.481 | 0.137 | 3.511 | 4.46E-04 | 2.55E-02 | 5 | 5781615 | 5783555 |
| Med12l | 4445.759 | 0.473 | 0.108 | 4.368 | 1.25E-05 | 2.08E-03 | 3 | 59005825 | 59318682 |
| Gabrb2 | 3541.671 | 0.471 | 0.118 | 4.000 | 6.33E-05 | 7.08E-03 | 11 | 42419757 | 42629028 |
| Zfp27 | 973.362 | 0.470 | 0.125 | 3.770 | 1.63E-04 | 1.35E-02 | 7 | 29893333 | 29906572 |
| Plxna4 | 13555.333 | 0.470 | 0.123 | 3.820 | 1.33E-04 | 1.18E-02 | 6 | 32144268 | 32588192 |
| Nrg1 | 10588.054 | 0.466 | 0.133 | 3.498 | 4.68E-04 | 2.62E-02 | 8 | 31814551 | 32884797 |
| Coro6 | 3205.404 | 0.465 | 0.140 | 3.311 | 9.28E-04 | 4.11E-02 | 11 | 77462411 | 77470484 |
| Rasl10b | 2926.630 | 0.462 | 0.123 | 3.763 | 1.68E-04 | 1.38E-02 | 11 | 83409137 | 83421039 |
| Mycbp2 | 37650.401 | 0.447 | 0.127 | 3.528 | 4.19E-04 | 2.45E-02 | 14 | 103113411 | 103346814 |
| Ptprn2 | 8360.082 | 0.445 | 0.124 | 3.578 | 3.46E-04 | 2.16E-02 | 12 | 116485720 | 117276849 |
| Rpl3 | 23883.404 | 0.436 | 0.078 | 5.599 | 2.16E-08 | 8.09E-06 | 15 | 80077791 | 80091868 |
| Card10 | 4407.016 | 0.427 | 0.114 | 3.751 | 1.76E-04 | 1.42E-02 | 15 | 78775138 | 78803042 |
| Gatb | 1537.457 | 0.425 | 0.120 | 3.549 | 3.87E-04 | 2.34E-02 | 3 | 85574119 | 85655622 |
| Rrp12 | 4324.850 | 0.416 | 0.126 | 3.317 | 9.09E-04 | 4.05E-02 | 19 | 41862851 | 41896173 |
| Sorcs3 | 1640.991 | 0.414 | 0.099 | 4.175 | 2.97E-05 | 3.92E-03 | 19 | 48206025 | 48805505 |
| Arid1b | 7071.578 | 0.409 | 0.117 | 3.491 | 4.82E-04 | 2.65E-02 | 17 | 4994332 | 5347656 |
| Rabepk | 2060.696 | 0.402 | 0.109 | 3.704 | 2.12E-04 | 1.62E-02 | 2 | 34777556 | 34799912 |
| Mapk8ip3 | 47684.218 | 0.384 | 0.109 | 3.507 | 4.54E-04 | 2.58E-02 | 17 | 24892153 | 24936977 |
| Epb41l1 | 13350.316 | 0.382 | 0.117 | 3.255 | 1.14E-03 | 4.60E-02 | 2 | 156420909 | 156543214 |
| Ntng2 | 6220.017 | 0.381 | 0.115 | 3.299 | 9.69E-04 | 4.19E-02 | 2 | 29194541 | 29253005 |
| Tubg1 | 6508.358 | 0.379 | 0.100 | 3.794 | 1.48E-04 | 1.28E-02 | 11 | 101119938 | 101126419 |
| Pde4d | 4943.745 | 0.370 | 0.091 | 4.076 | 4.57E-05 | 5.53E-03 | 13 | 108449948 | 109953461 |
| Wdr4 | 2083.313 | 0.368 | 0.102 | 3.625 | 2.89E-04 | 1.95E-02 | 17 | 31494322 | 31519980 |
| Ighmbp2 | 2192.111 | 0.368 | 0.106 | 3.454 | 5.52E-04 | 2.90E-02 | 19 | 3259076 | 3283017 |
| Abcg4 | 5126.537 | 0.362 | 0.107 | 3.388 | 7.05E-04 | 3.39E-02 | 9 | 44273188 | 44288615 |
| Tecpr1 | 10317.958 | 0.362 | 0.081 | 4.469 | 7.84E-06 | 1.40E-03 | 5 | 144194442 | 144223615 |
| C2cd2l | 6107.831 | 0.358 | 0.083 | 4.314 | 1.60E-05 | 2.52E-03 | 9 | 44309237 | 44320285 |
| B4galt5 | 7439.787 | 0.355 | 0.093 | 3.821 | 1.33E-04 | 1.18E-02 | 2 | 167298444 | 167349183 |
| Etl4 | 11167.581 | 0.348 | 0.106 | 3.294 | 9.88E-04 | 4.20E-02 | 2 | 19909780 | 20810713 |
| Vwa8 | 10184.174 | 0.337 | 0.081 | 4.179 | 2.93E-05 | 3.90E-03 | 14 | 78849052 | 79202310 |
| Eml2 | 27753.184 | 0.333 | 0.083 | 4.001 | 6.29E-05 | 7.08E-03 | 7 | 19176421 | 19206482 |
| Sh3glb2 | 15314.377 | 0.325 | 0.099 | 3.294 | 9.87E-04 | 4.20E-02 | 2 | 30344809 | 30359337 |
| Emc10 | 18786.916 | 0.318 | 0.090 | 3.545 | 3.92E-04 | 2.34E-02 | 7 | 44489937 | 44496529 |
| Ank3 | 23503.539 | 0.316 | 0.095 | 3.316 | 9.12E-04 | 4.05E-02 | 10 | 69398773 | 70027438 |
| Dync1i1 | 15185.507 | 0.312 | 0.088 | 3.542 | 3.97E-04 | 2.36E-02 | 6 | 5725639 | 6028039 |
| Arfgef3 | 25018.191 | 0.299 | 0.075 | 3.980 | 6.88E-05 | 7.47E-03 | 10 | 18581839 | 18743949 |
| Mybbp1a | 6091.501 | 0.291 | 0.082 | 3.535 | 4.08E-04 | 2.41E-02 | 11 | 72441355 | 72451768 |
| Rabac1 | 21897.438 | 0.270 | 0.070 | 3.881 | 1.04E-04 | 1.00E-02 | 7 | 24969752 | 24972754 |
| Rab8b | 6775.767 | -0.243 | 0.074 | -3.275 | 1.06E-03 | 4.38E-02 | 9 | 66843664 | 66919687 |
| Syap1 | 4635.998 | -0.301 | 0.091 | -3.296 | 9.80E-04 | 4.20E-02 | X | 162857057 | 162888447 |
| Tmed7 | 13366.743 | -0.304 | 0.083 | -3.680 | 2.33E-04 | 1.71E-02 | 18 | 46560235 | 46597535 |
| Zfp91 | 19486.733 | -0.313 | 0.089 | -3.531 | 4.15E-04 | 2.44E-02 | 19 | 12767020 | 12796126 |
| Rnf130 | 8918.034 | -0.325 | 0.079 | -4.086 | 4.39E-05 | 5.41E-03 | 11 | 50025346 | 50125719 |
| Sav1 | 2158.170 | -0.328 | 0.101 | -3.238 | 1.20E-03 | 4.75E-02 | 12 | 69965012 | 69987002 |
| Stag2 | 11089.378 | -0.329 | 0.093 | -3.515 | 4.39E-04 | 2.53E-02 | X | 42149317 | 42277185 |

| | | | | | | | | | |
|---|---|---|---|---|---|---|---|---|---|
| Smim10l1 | 11805.207 | -0.345 | 0.097 | -3.554 | 3.79E-04 | 2.30E-02 | 6 | 133104909 | 133110899 |
| Plin3 | 6211.303 | -0.347 | 0.104 | -3.338 | 8.45E-04 | 3.89E-02 | 17 | 56277476 | 56292873 |
| Cnot6 | 5177.474 | -0.349 | 0.089 | -3.926 | 8.65E-05 | 8.69E-03 | 11 | 49671503 | 49712723 |
| Hdgfl3 | 10175.483 | -0.350 | 0.096 | -3.655 | 2.57E-04 | 1.81E-02 | 7 | 81881251 | 81934473 |
| Selenop | 85517.620 | -0.361 | 0.094 | -3.846 | 1.20E-04 | 1.11E-02 | 15 | 3268547 | 3280508 |
| Poglut3 | 1842.450 | -0.362 | 0.104 | -3.492 | 4.80E-04 | 2.65E-02 | 9 | 53384025 | 53401867 |
| Mfsd14a | 7826.397 | -0.372 | 0.091 | -4.099 | 4.16E-05 | 5.19E-03 | 3 | 116631164 | 116662677 |
| Marveld1 | 3934.392 | -0.373 | 0.111 | -3.355 | 7.92E-04 | 3.70E-02 | 19 | 42147400 | 42151703 |
| Stambp | 3011.263 | -0.373 | 0.089 | -4.186 | 2.84E-05 | 3.89E-03 | 6 | 83543211 | 83572729 |
| Irs1 | 2617.073 | -0.378 | 0.116 | -3.269 | 1.08E-03 | 4.44E-02 | 1 | 82233101 | 82291416 |
| Phf10 | 2690.296 | -0.380 | 0.112 | -3.406 | 6.60E-04 | 3.25E-02 | 17 | 14945009 | 14961273 |
| Prrg1 | 2026.853 | -0.385 | 0.112 | -3.443 | 5.76E-04 | 2.98E-02 | X | 78449613 | 78583896 |
| Zfp748 | 1493.433 | -0.391 | 0.100 | -3.897 | 9.76E-05 | 9.51E-03 | 13 | 67538641 | 67553830 |
| Tbc1d12 | 2628.989 | -0.394 | 0.109 | -3.615 | 3.00E-04 | 1.97E-02 | 19 | 38836579 | 38919923 |
| Commd6 | 2591.426 | -0.395 | 0.108 | -3.655 | 2.57E-04 | 1.81E-02 | 14 | 101632981 | 101640686 |
| Ss18 | 3180.225 | -0.397 | 0.122 | -3.245 | 1.17E-03 | 4.67E-02 | 18 | 14624198 | 14682914 |
| Manea | 6037.772 | -0.398 | 0.089 | -4.448 | 8.68E-06 | 1.51E-03 | 4 | 26324506 | 26346891 |
| Tbp | 1777.345 | -0.399 | 0.115 | -3.455 | 5.51E-04 | 2.90E-02 | 17 | 15499888 | 15528379 |
| Morf4l1-ps1 | 5001.575 | -0.404 | 0.116 | -3.496 | 4.73E-04 | 2.62E-02 | 16 | 24529394 | 24530362 |
| Smarcad1 | 2350.473 | -0.407 | 0.120 | -3.394 | 6.88E-04 | 3.36E-02 | 6 | 65042583 | 65116061 |
| Slco3a1 | 3256.285 | -0.415 | 0.115 | -3.614 | 3.02E-04 | 1.97E-02 | 7 | 74275419 | 74554780 |
| Sesn1 | 7329.236 | -0.425 | 0.088 | -4.839 | 1.30E-06 | 3.19E-04 | 10 | 41809935 | 41908424 |
| Elk4 | 812.164 | -0.428 | 0.131 | -3.276 | 1.05E-03 | 4.38E-02 | 1 | 132007607 | 132032612 |
| Utp18 | 1591.244 | -0.433 | 0.110 | -3.949 | 7.84E-05 | 8.05E-03 | 11 | 93859243 | 93885766 |
| Zdhhc20 | 4269.377 | -0.443 | 0.128 | -3.476 | 5.09E-04 | 2.76E-02 | 14 | 57832703 | 57890276 |
| Arfip1 | 3785.586 | -0.444 | 0.095 | -4.690 | 2.73E-06 | 5.87E-04 | 3 | 84496093 | 84582625 |
| Pon2 | 5309.433 | -0.449 | 0.106 | -4.232 | 2.31E-05 | 3.43E-03 | 6 | 5264147 | 5298455 |
| Lrrc28 | 2378.456 | -0.456 | 0.094 | -4.872 | 1.10E-06 | 2.74E-04 | 7 | 67513410 | 67645268 |
| Nxt2 | 4424.131 | -0.458 | 0.112 | -4.094 | 4.25E-05 | 5.27E-03 | X | 142226770 | 142239692 |
| Hopx | 11928.296 | -0.458 | 0.140 | -3.263 | 1.10E-03 | 4.52E-02 | 5 | 77086988 | 77115121 |
| Zfp759 | 783.727 | -0.459 | 0.125 | -3.661 | 2.51E-04 | 1.79E-02 | 13 | 67121660 | 67141787 |
| Micu2 | 4228.205 | -0.465 | 0.134 | -3.474 | 5.13E-04 | 2.76E-02 | 14 | 57916261 | 57999262 |
| Creg1 | 5464.297 | -0.474 | 0.135 | -3.507 | 4.53E-04 | 2.58E-02 | 1 | 165763746 | 165775308 |
| Sh3bgrl | 14076.026 | -0.475 | 0.128 | -3.725 | 1.95E-04 | 1.53E-02 | X | 109095365 | 109197873 |
| Trim13 | 1517.562 | -0.481 | 0.132 | -3.641 | 2.72E-04 | 1.88E-02 | 14 | 61598247 | 61605946 |
| Ak3 | 11220.716 | -0.485 | 0.130 | -3.743 | 1.82E-04 | 1.45E-02 | 19 | 29020833 | 29047961 |
| Mbtd1 | 1548.433 | -0.488 | 0.148 | -3.305 | 9.51E-04 | 4.16E-02 | 11 | 93885852 | 93946985 |
| Septin10 | 1540.905 | -0.489 | 0.118 | -4.155 | 3.25E-05 | 4.19E-03 | 10 | 59141627 | 59221847 |
| Qk | 55419.877 | -0.490 | 0.142 | -3.448 | 5.66E-04 | 2.95E-02 | 17 | 10202601 | 10319854 |
| Septin7 | 27122.942 | -0.495 | 0.145 | -3.405 | 6.61E-04 | 3.25E-02 | 9 | 25252439 | 25308571 |
| Tmem47 | 10011.181 | -0.497 | 0.151 | -3.302 | 9.61E-04 | 4.18E-02 | X | 81070698 | 81097872 |
| Fbxl4 | 2754.262 | -0.497 | 0.144 | -3.445 | 5.71E-04 | 2.97E-02 | 4 | 22357543 | 22434091 |
| Zfp958 | 1029.766 | -0.499 | 0.136 | -3.656 | 2.56E-04 | 1.81E-02 | 8 | 4613167 | 4630231 |
| Ggh | 1760.560 | -0.499 | 0.142 | -3.513 | 4.43E-04 | 2.55E-02 | 4 | 20042052 | 20066750 |
| Rras2 | 1540.858 | -0.504 | 0.105 | -4.800 | 1.59E-06 | 3.74E-04 | 7 | 114046782 | 114117781 |
| Gm14698 | 597.300 | -0.509 | 0.156 | -3.272 | 1.07E-03 | 4.40E-02 | X | 68821093 | 68825318 |
| Tnip2 | 671.986 | -0.510 | 0.143 | -3.564 | 3.65E-04 | 2.24E-02 | 5 | 34496087 | 34513991 |
| Enpp1 | 3590.361 | -0.518 | 0.156 | -3.328 | 8.76E-04 | 3.96E-02 | 10 | 24637914 | 24712159 |
| Ap1s2 | 7173.354 | -0.519 | 0.123 | -4.218 | 2.46E-05 | 3.49E-03 | X | 163909017 | 163933666 |
| Stau2 | 14329.861 | -0.522 | 0.130 | -4.012 | 6.03E-05 | 6.94E-03 | 1 | 16228674 | 16520112 |

| | | | | | | | | | |
|---|---|---|---|---|---|---|---|---|---|
| Vamp3 | 4539.658 | -0.526 | 0.088 | -5.978 | 2.26E-09 | 1.13E-06 | 4 | 151047300 | 151057963 |
| Septin2 | 10692.508 | -0.537 | 0.119 | -4.506 | 6.60E-06 | 1.20E-03 | 1 | 93478993 | 93509733 |
| Ackr3 | 954.974 | -0.539 | 0.163 | -3.313 | 9.24E-04 | 4.10E-02 | 1 | 90203980 | 90216751 |
| Snapc1 | 3181.484 | -0.542 | 0.115 | -4.716 | 2.40E-06 | 5.27E-04 | 12 | 73964481 | 73988966 |
| Rps2 | 3138.048 | -0.542 | 0.142 | -3.825 | 1.31E-04 | 1.18E-02 | 17 | 24718116 | 24721929 |
| Plp1 | 113872.779 | -0.545 | 0.161 | -3.393 | 6.92E-04 | 3.36E-02 | X | 136822671 | 136839733 |
| Zfp619 | 612.862 | -0.549 | 0.162 | -3.393 | 6.91E-04 | 3.36E-02 | 7 | 39517766 | 39540420 |
| Car13 | 2411.399 | -0.555 | 0.165 | -3.371 | 7.49E-04 | 3.55E-02 | 3 | 14641727 | 14663002 |
| Tia1 | 1258.383 | -0.560 | 0.160 | -3.496 | 4.73E-04 | 2.62E-02 | 6 | 86404219 | 86433405 |
| Arsk | 1258.874 | -0.567 | 0.145 | -3.921 | 8.81E-05 | 8.76E-03 | 13 | 76060422 | 76098660 |
| Carnmt1 | 1955.654 | -0.573 | 0.136 | -4.226 | 2.38E-05 | 3.48E-03 | 19 | 18670764 | 18707200 |
| Eva1b | 826.284 | -0.574 | 0.178 | -3.221 | 1.28E-03 | 4.96E-02 | 4 | 126147744 | 126149875 |
| Marcks | 8988.449 | -0.614 | 0.146 | -4.219 | 2.45E-05 | 3.49E-03 | 10 | 37133375 | 37138920 |
| Hmgcs1 | 59372.097 | -0.617 | 0.146 | -4.235 | 2.29E-05 | 3.42E-03 | 13 | 119690379 | 119708260 |
| Kcnk5 | 1989.662 | -0.623 | 0.127 | -4.921 | 8.62E-07 | 2.17E-04 | 14 | 20140057 | 20181809 |
| Wtip | 1180.543 | -0.624 | 0.157 | -3.973 | 7.11E-05 | 7.60E-03 | 7 | 34109543 | 34133268 |
| Ginm1 | 2071.129 | -0.626 | 0.183 | -3.413 | 6.43E-04 | 3.20E-02 | 10 | 7767947 | 7792824 |
| Cdo1 | 1885.301 | -0.626 | 0.170 | -3.681 | 2.32E-04 | 1.70E-02 | 18 | 46713193 | 46728395 |
| Zc3hav1l | 700.573 | -0.630 | 0.181 | -3.475 | 5.12E-04 | 2.76E-02 | 6 | 38287396 | 38299259 |
| Gpx8 | 2458.845 | -0.635 | 0.126 | -5.029 | 4.93E-07 | 1.29E-04 | 13 | 113042753 | 113046410 |
| Nudt19 | 2714.828 | -0.651 | 0.200 | -3.251 | 1.15E-03 | 4.61E-02 | 7 | 35547185 | 35556304 |
| Art3 | 5890.724 | -0.663 | 0.194 | -3.424 | 6.16E-04 | 3.10E-02 | 5 | 92331827 | 92414628 |
| Lhpp | 660.065 | -0.671 | 0.181 | -3.706 | 2.11E-04 | 1.62E-02 | 7 | 132610638 | 132706420 |
| Senp6 | 5074.547 | -0.672 | 0.167 | -4.026 | 5.68E-05 | 6.58E-03 | 9 | 80066903 | 80144953 |
| Rcn1 | 8259.555 | -0.676 | 0.149 | -4.546 | 5.47E-06 | 1.01E-03 | 2 | 105386291 | 105399319 |
| Chpt1 | 8124.610 | -0.676 | 0.142 | -4.762 | 1.91E-06 | 4.41E-04 | 10 | 88452745 | 88504073 |
| Cebpb | 1058.001 | -0.679 | 0.180 | -3.771 | 1.63E-04 | 1.35E-02 | 2 | 167688915 | 167690418 |
| Pard6g | 3364.108 | -0.681 | 0.181 | -3.771 | 1.62E-04 | 1.35E-02 | 18 | 80046890 | 80119639 |
| Dchs2 | 603.269 | -0.690 | 0.171 | -4.038 | 5.40E-05 | 6.33E-03 | 3 | 83127948 | 83357209 |
| Tmem64 | 10461.912 | -0.699 | 0.098 | -7.146 | 8.94E-13 | 7.12E-10 | 4 | 15265831 | 15286753 |
| Casp6 | 589.972 | -0.739 | 0.224 | -3.300 | 9.67E-04 | 4.19E-02 | 3 | 129901425 | 129914103 |
| Zkscan7 | 459.562 | -0.739 | 0.213 | -3.472 | 5.16E-04 | 2.77E-02 | 9 | 122885685 | 122898618 |
| Mlip | 2889.341 | -0.758 | 0.234 | -3.245 | 1.17E-03 | 4.67E-02 | 9 | 77102081 | 77352969 |
| Cdc14a | 1486.882 | -0.761 | 0.207 | -3.683 | 2.30E-04 | 1.70E-02 | 3 | 116272553 | 116428745 |
| Ogn | 47750.318 | -0.764 | 0.138 | -5.519 | 3.41E-08 | 1.19E-05 | 13 | 49608046 | 49624501 |
| Myl9 | 4075.688 | -0.767 | 0.174 | -4.421 | 9.84E-06 | 1.66E-03 | 2 | 156775420 | 156781658 |
| Iars2 | 375.994 | -0.776 | 0.212 | -3.660 | 2.52E-04 | 1.79E-02 | 1 | 185284726 | 185329396 |
| Olfml1 | 1587.347 | -0.786 | 0.237 | -3.320 | 9.00E-04 | 4.03E-02 | 7 | 107567446 | 107591094 |
| Rnf208 | 497.151 | -0.787 | 0.201 | -3.911 | 9.21E-05 | 9.11E-03 | 2 | 25242929 | 25244262 |
| Lox | 1921.122 | -0.789 | 0.226 | -3.498 | 4.68E-04 | 2.62E-02 | 18 | 52516067 | 52529867 |
| Wtap | 3960.743 | -0.798 | 0.221 | -3.614 | 3.01E-04 | 1.97E-02 | 17 | 12966796 | 12994169 |
| Mmp19 | 394.643 | -0.804 | 0.188 | -4.288 | 1.81E-05 | 2.74E-03 | 10 | 128790910 | 128800824 |
| Mpp6 | 1760.464 | -0.805 | 0.204 | -3.945 | 7.96E-05 | 8.12E-03 | 6 | 50110241 | 50198939 |
| Kmt5a | 5135.346 | -0.807 | 0.127 | -6.369 | 1.91E-10 | 1.04E-07 | 5 | 124439930 | 124462308 |
| Spin4 | 180.891 | -0.825 | 0.241 | -3.419 | 6.27E-04 | 3.14E-02 | X | 95022510 | 95026682 |
| Gm28438 | 17903.678 | -0.828 | 0.252 | -3.283 | 1.03E-03 | 4.30E-02 | 1 | 24612775 | 24613119 |
| Rrp1 | 6991.910 | -0.830 | 0.116 | -7.126 | 1.03E-12 | 7.88E-10 | 10 | 78400384 | 78413043 |
| Lpar4 | 264.738 | -0.834 | 0.253 | -3.293 | 9.90E-04 | 4.20E-02 | X | 106920625 | 106933900 |
| Cdc42ep5 | 293.524 | -0.835 | 0.248 | -3.364 | 7.67E-04 | 3.61E-02 | 7 | 4151260 | 4164860 |
| Mbp | 114115.243 | -0.835 | 0.230 | -3.635 | 2.78E-04 | 1.90E-02 | 18 | 82475146 | 82585637 |

| | | | | | | | | | |
|---|---|---|---|---|---|---|---|---|---|
| Pkdcc | 2565.059 | -0.840 | 0.255 | -3.295 | 9.84E-04 | 4.20E-02 | 17 | 83215292 | 83225070 |
| mt-Nd3 | 17209.139 | -0.845 | 0.250 | -3.378 | 7.30E-04 | 3.48E-02 | MT | 9459 | 9806 |
| B3galt2 | 2999.946 | -0.847 | 0.236 | -3.582 | 3.41E-04 | 2.15E-02 | 1 | 143640664 | 143654614 |
| Calcrl | 1084.732 | -0.873 | 0.236 | -3.700 | 2.15E-04 | 1.63E-02 | 2 | 84330626 | 84425411 |
| Ppp1r3b | 1174.310 | -0.873 | 0.221 | -3.945 | 7.99E-05 | 8.12E-03 | 8 | 35375739 | 35388139 |
| Cybrd1 | 1954.631 | -0.887 | 0.200 | -4.446 | 8.77E-06 | 1.51E-03 | 2 | 71117923 | 71142926 |
| Slc1a3 | 2716.709 | -0.900 | 0.231 | -3.904 | 9.47E-05 | 9.28E-03 | 15 | 8634124 | 8710764 |
| Ccn4 | 1260.820 | -0.912 | 0.251 | -3.627 | 2.86E-04 | 1.94E-02 | 15 | 66891320 | 66923201 |
| Efcab14 | 1472.717 | -0.931 | 0.196 | -4.755 | 1.99E-06 | 4.52E-04 | 4 | 115737744 | 115777327 |
| Nexn | 619.930 | -0.947 | 0.280 | -3.383 | 7.16E-04 | 3.43E-02 | 3 | 152236982 | 152266350 |
| Kcne4 | 285.147 | -0.948 | 0.288 | -3.288 | 1.01E-03 | 4.25E-02 | 1 | 78816758 | 78820028 |
| Fibin | 1235.136 | -0.953 | 0.294 | -3.244 | 1.18E-03 | 4.68E-02 | 2 | 110360917 | 110363183 |
| Fndc1 | 992.771 | -0.954 | 0.214 | -4.449 | 8.63E-06 | 1.51E-03 | 17 | 7738569 | 7827302 |
| Cmbl | 180.772 | -0.961 | 0.281 | -3.419 | 6.29E-04 | 3.14E-02 | 15 | 31565389 | 31590119 |
| Sncaip | 432.873 | -0.964 | 0.263 | -3.660 | 2.53E-04 | 1.79E-02 | 18 | 52767709 | 52915935 |
| Mycl | 387.763 | -0.970 | 0.185 | -5.241 | 1.59E-07 | 4.61E-05 | 4 | 122995652 | 123002485 |
| Ssc5d | 419.419 | -0.980 | 0.242 | -4.047 | 5.20E-05 | 6.13E-03 | 7 | 4925785 | 4944826 |
| Yipf2 | 943.637 | -0.981 | 0.171 | -5.753 | 8.76E-09 | 3.42E-06 | 9 | 21588682 | 21592828 |
| Acot1 | 1195.356 | -1.012 | 0.221 | -4.572 | 4.84E-06 | 9.34E-04 | 12 | 84009490 | 84018371 |
| Gm9780 | 353.262 | -1.033 | 0.198 | -5.220 | 1.79E-07 | 5.10E-05 | 14 | 26027282 | 26042711 |
| Vps29 | 9493.994 | -1.062 | 0.139 | -7.612 | 2.70E-14 | 2.71E-11 | 5 | 122354369 | 122364984 |
| Plac9b | 282.635 | -1.066 | 0.197 | -5.411 | 6.27E-08 | 2.07E-05 | 14 | 26167405 | 26182480 |
| Wdcp | 922.312 | -1.068 | 0.163 | -6.561 | 5.33E-11 | 3.29E-08 | 12 | 4843303 | 4860043 |
| Aoc3 | 373.604 | -1.079 | 0.284 | -3.799 | 1.46E-04 | 1.27E-02 | 11 | 101330605 | 101341938 |
| Itgb5 | 6419.606 | -1.080 | 0.315 | -3.428 | 6.08E-04 | 3.07E-02 | 16 | 33829665 | 33949338 |
| Dcdc2a | 451.648 | -1.093 | 0.291 | -3.751 | 1.76E-04 | 1.42E-02 | 13 | 25056004 | 25210706 |
| Atp11b | 3716.342 | -1.103 | 0.198 | -5.569 | 2.56E-08 | 9.25E-06 | 3 | 35754106 | 35856276 |
| Atp10b | 553.219 | -1.106 | 0.304 | -3.640 | 2.72E-04 | 1.88E-02 | 11 | 43149877 | 43262285 |
| Gm48529 | 714.115 | -1.114 | 0.316 | -3.521 | 4.30E-04 | 2.51E-02 | 9 | 45930112 | 45933738 |
| Itm2a | 13802.212 | -1.122 | 0.136 | -8.235 | 1.79E-16 | 2.64E-13 | X | 107397099 | 107403376 |
| Fiz1 | 1346.996 | -1.124 | 0.309 | -3.635 | 2.78E-04 | 1.90E-02 | 7 | 5007059 | 5014697 |
| Col4a5 | 779.455 | -1.132 | 0.338 | -3.344 | 8.26E-04 | 3.83E-02 | X | 141475385 | 141689234 |
| Clec14a | 1255.015 | -1.142 | 0.265 | -4.311 | 1.63E-05 | 2.53E-03 | 12 | 58264720 | 58269290 |
| Atg9a | 2592.171 | -1.149 | 0.136 | -8.438 | 3.23E-17 | 5.61E-14 | 1 | 75180860 | 75192196 |
| Tagln | 1528.879 | -1.172 | 0.247 | -4.745 | 2.09E-06 | 4.70E-04 | 9 | 45929619 | 45936058 |
| Hspd1 | 1455.359 | -1.178 | 0.156 | -7.571 | 3.69E-14 | 3.53E-11 | 1 | 55077835 | 55088243 |
| Resp18 | 237.149 | -1.186 | 0.358 | -3.317 | 9.09E-04 | 4.05E-02 | 1 | 75272199 | 75278415 |
| Gpr4 | 304.325 | -1.192 | 0.368 | -3.236 | 1.21E-03 | 4.76E-02 | 7 | 19212538 | 19224174 |
| Cfap97 | 2916.188 | -1.208 | 0.368 | -3.284 | 1.02E-03 | 4.30E-02 | 8 | 46151771 | 46195590 |
| Usp21 | 507.110 | -1.233 | 0.366 | -3.365 | 7.67E-04 | 3.61E-02 | 1 | 171281945 | 171287991 |
| Dbi | 3126.946 | -1.235 | 0.303 | -4.074 | 4.62E-05 | 5.55E-03 | 1 | 120113280 | 120121078 |
| Prrx2 | 314.621 | -1.244 | 0.288 | -4.316 | 1.59E-05 | 2.52E-03 | 2 | 30834972 | 30881251 |
| Aebp1 | 6742.215 | -1.264 | 0.339 | -3.730 | 1.92E-04 | 1.52E-02 | 11 | 5861947 | 5872088 |
| Ptgis | 949.197 | -1.270 | 0.320 | -3.971 | 7.15E-05 | 7.60E-03 | 2 | 167191805 | 167240604 |
| Colec12 | 3075.784 | -1.285 | 0.379 | -3.392 | 6.93E-04 | 3.36E-02 | 18 | 9707595 | 9882644 |
| Ppic | 1165.890 | -1.304 | 0.366 | -3.560 | 3.70E-04 | 2.26E-02 | 18 | 53406332 | 53418115 |
| Ankrd13c | 1555.324 | -1.305 | 0.393 | -3.319 | 9.04E-04 | 4.04E-02 | 3 | 157947239 | 158008034 |
| Clcf1 | 156.060 | -1.306 | 0.354 | -3.686 | 2.28E-04 | 1.69E-02 | 19 | 4214238 | 4223490 |
| Twist1 | 463.617 | -1.321 | 0.388 | -3.407 | 6.58E-04 | 3.25E-02 | 12 | 33957671 | 33959829 |
| Lratd2 | 749.669 | -1.331 | 0.404 | -3.296 | 9.82E-04 | 4.20E-02 | 15 | 60818994 | 60853778 |

| | | | | | | | | | |
|---|---|---|---|---|---|---|---|---|---|
| Epha3 | 630.188 | -1.337 | 0.365 | -3.668 | 2.44E-04 | 1.76E-02 | 16 | 63543534 | 63864175 |
| Aph1b | 697.983 | -1.341 | 0.228 | -5.881 | 4.09E-09 | 1.90E-06 | 9 | 66775202 | 66795490 |
| Gxylt2 | 720.913 | -1.341 | 0.271 | -4.946 | 7.58E-07 | 1.93E-04 | 6 | 100704734 | 100810913 |
| Cdh3 | 137.314 | -1.343 | 0.411 | -3.265 | 1.09E-03 | 4.48E-02 | 8 | 106510891 | 106557297 |
| Sp7 | 409.905 | -1.343 | 0.372 | -3.616 | 2.99E-04 | 1.97E-02 | 15 | 102356606 | 102367182 |
| Gm6477 | 120.155 | -1.345 | 0.347 | -3.873 | 1.08E-04 | 1.02E-02 | 10 | 39198538 | 39199301 |
| Gkn3 | 208.476 | -1.395 | 0.300 | -4.650 | 3.33E-06 | 6.76E-04 | 6 | 87383256 | 87388935 |
| Trpm8 | 2391.152 | -1.398 | 0.411 | -3.399 | 6.75E-04 | 3.31E-02 | 1 | 88277661 | 88389293 |
| Spsb2 | 616.507 | -1.400 | 0.277 | -5.047 | 4.50E-07 | 1.19E-04 | 6 | 124808661 | 124810619 |
| Dipk2b | 184.817 | -1.427 | 0.346 | -4.126 | 3.69E-05 | 4.70E-03 | X | 18414881 | 18461397 |
| Acta2 | 3376.040 | -1.431 | 0.314 | -4.561 | 5.09E-06 | 9.53E-04 | 19 | 34241090 | 34255590 |
| Mfap4 | 1216.979 | -1.448 | 0.422 | -3.430 | 6.03E-04 | 3.06E-02 | 11 | 61485431 | 61488900 |
| Snx13 | 1752.541 | -1.449 | 0.329 | -4.408 | 1.04E-05 | 1.75E-03 | 12 | 35047186 | 35147479 |
| Pter | 746.819 | -1.481 | 0.205 | -7.206 | 5.75E-13 | 4.78E-10 | 2 | 12924041 | 13003455 |
| Pdgfra | 2593.599 | -1.497 | 0.313 | -4.786 | 1.70E-06 | 3.97E-04 | 5 | 75152292 | 75198215 |
| Six2 | 501.353 | -1.501 | 0.418 | -3.588 | 3.33E-04 | 2.14E-02 | 17 | 85684277 | 85688274 |
| Gm48840 | 556.350 | -1.506 | 0.360 | -4.184 | 2.87E-05 | 3.89E-03 | 9 | 45051424 | 45055344 |
| Zfp185 | 311.456 | -1.510 | 0.280 | -5.396 | 6.82E-08 | 2.17E-05 | X | 72987339 | 73031543 |
| Mrps9 | 930.775 | -1.517 | 0.429 | -3.533 | 4.10E-04 | 2.42E-02 | 1 | 42851233 | 42905683 |
| Omd | 898.458 | -1.520 | 0.407 | -3.735 | 1.88E-04 | 1.50E-02 | 13 | 49582462 | 49592822 |
| Sesn3 | 6110.573 | -1.574 | 0.439 | -3.584 | 3.38E-04 | 2.14E-02 | 9 | 14275067 | 14333101 |
| Ifitm1 | 559.223 | -1.598 | 0.433 | -3.694 | 2.21E-04 | 1.66E-02 | 7 | 140967221 | 140969825 |
| Tmem252 | 233.824 | -1.599 | 0.445 | -3.591 | 3.29E-04 | 2.13E-02 | 19 | 24674008 | 24682233 |
| Crabp2 | 695.422 | -1.609 | 0.492 | -3.272 | 1.07E-03 | 4.40E-02 | 3 | 87948666 | 87953376 |
| Dapl1 | 122.968 | -1.612 | 0.483 | -3.335 | 8.52E-04 | 3.90E-02 | 2 | 59484653 | 59505020 |
| Alx4 | 676.720 | -1.628 | 0.450 | -3.616 | 2.99E-04 | 1.97E-02 | 2 | 93642384 | 93681339 |
| Gm37412 | 99.308 | -1.650 | 0.353 | -4.678 | 2.90E-06 | 6.13E-04 | 1 | 14788645 | 14792965 |
| Mall | 84.660 | -1.656 | 0.482 | -3.438 | 5.87E-04 | 2.99E-02 | 2 | 127704386 | 127729932 |
| H19 | 1487.676 | -1.716 | 0.376 | -4.561 | 5.08E-06 | 9.53E-04 | 7 | 142575529 | 142578143 |
| Car3 | 1133.281 | -1.723 | 0.522 | -3.302 | 9.61E-04 | 4.18E-02 | 3 | 14863512 | 14872523 |
| Tnmd | 105.846 | -1.737 | 0.537 | -3.236 | 1.21E-03 | 4.76E-02 | X | 133850980 | 133865577 |
| Myh11 | 1885.585 | -1.737 | 0.372 | -4.676 | 2.92E-06 | 6.13E-04 | 16 | 14194535 | 14291372 |
| Gm7903 | 45.073 | -1.756 | 0.477 | -3.682 | 2.32E-04 | 1.70E-02 | X | 135433835 | 135443830 |
| Gm5128 | 46.335 | -1.784 | 0.476 | -3.746 | 1.79E-04 | 1.45E-02 | X | 135373284 | 135383393 |
| Pmp2 | 1038.696 | -1.794 | 0.513 | -3.496 | 4.72E-04 | 2.62E-02 | 3 | 10179851 | 10183929 |
| Mn1 | 542.600 | -1.840 | 0.461 | -3.995 | 6.47E-05 | 7.18E-03 | 5 | 111417362 | 111457033 |
| Slc2a12 | 585.859 | -1.846 | 0.554 | -3.332 | 8.62E-04 | 3.92E-02 | 10 | 22645011 | 22704285 |
| Fam180a | 411.920 | -1.848 | 0.427 | -4.328 | 1.50E-05 | 2.42E-03 | 6 | 35312668 | 35326141 |
| Gm38101 | 44.507 | -1.863 | 0.529 | -3.518 | 4.35E-04 | 2.52E-02 | 1 | 176276213 | 176279769 |
| Lyrm7 | 618.377 | -1.883 | 0.284 | -6.620 | 3.58E-11 | 2.28E-08 | 11 | 54826866 | 54860916 |
| Lmod1 | 357.419 | -1.943 | 0.509 | -3.821 | 1.33E-04 | 1.18E-02 | 1 | 135324807 | 135368065 |
| Cthrc1 | 378.839 | -1.947 | 0.362 | -5.383 | 7.31E-08 | 2.29E-05 | 15 | 39076932 | 39087121 |
| H2-DMb1 | 1321.442 | -1.992 | 0.272 | -7.324 | 2.40E-13 | 2.09E-10 | 17 | 34153072 | 34160230 |
| Aldh1a2 | 2592.534 | -1.997 | 0.570 | -3.501 | 4.63E-04 | 2.62E-02 | 9 | 71215789 | 71296243 |
| Slc13a4 | 2732.676 | -2.012 | 0.536 | -3.754 | 1.74E-04 | 1.42E-02 | 6 | 35267957 | 35308131 |
| Clec3b | 443.503 | -2.020 | 0.625 | -3.230 | 1.24E-03 | 4.83E-02 | 9 | 123150946 | 123157432 |
| Dhx16 | 305.439 | -2.111 | 0.255 | -8.289 | 1.14E-16 | 1.82E-13 | 17 | 35879819 | 35892670 |
| Fmod | 5804.661 | -2.168 | 0.547 | -3.964 | 7.36E-05 | 7.67E-03 | 1 | 134037254 | 134048277 |
| Hmcn2 | 71.818 | -2.236 | 0.689 | -3.245 | 1.18E-03 | 4.67E-02 | 2 | 31314415 | 31460738 |
| Ptgds | 45874.845 | -2.242 | 0.605 | -3.705 | 2.12E-04 | 1.62E-02 | 2 | 25466709 | 25470046 |

| | | | | | | | | | |
|---|---|---|---|---|---|---|---|---|---|
| Nupr1 | 997.233 | -2.275 | 0.348 | -6.541 | 6.10E-11 | 3.64E-08 | 7 | 126623249 | 126630861 |
| Arhgap29 | 868.227 | -2.307 | 0.505 | -4.564 | 5.02E-06 | 9.53E-04 | 3 | 121952541 | 122016753 |
| Hcar1 | 317.193 | -2.307 | 0.614 | -3.759 | 1.70E-04 | 1.40E-02 | 5 | 123876736 | 123880020 |
| C430049B03Rik | 31.695 | -2.322 | 0.679 | -3.420 | 6.26E-04 | 3.14E-02 | X | 53055046 | 53057190 |
| Ppp1r3a | 907.597 | -2.331 | 0.693 | -3.362 | 7.72E-04 | 3.63E-02 | 6 | 14713977 | 14755274 |
| Coch | 289.411 | -2.354 | 0.589 | -4.000 | 6.34E-05 | 7.08E-03 | 12 | 51593341 | 51605771 |
| Rangap1 | 2424.781 | -2.379 | 0.657 | -3.620 | 2.94E-04 | 1.97E-02 | 15 | 81704248 | 81745530 |
| Slc22a6 | 1045.899 | -2.383 | 0.693 | -3.439 | 5.83E-04 | 2.99E-02 | 19 | 8618039 | 8628299 |
| Xntrpc | 115.066 | -2.393 | 0.342 | -7.005 | 2.47E-12 | 1.82E-09 | 7 | 102065713 | 102096864 |
| Dlx5 | 86.064 | -2.469 | 0.683 | -3.618 | 2.97E-04 | 1.97E-02 | 6 | 6877805 | 6882085 |
| Asns | 3335.367 | -2.505 | 0.656 | -3.819 | 1.34E-04 | 1.18E-02 | 6 | 7675169 | 7693254 |
| Trim63 | 134.284 | -2.508 | 0.700 | -3.585 | 3.36E-04 | 2.14E-02 | 4 | 134315120 | 134329629 |
| Osr1 | 464.090 | -2.519 | 0.592 | -4.256 | 2.08E-05 | 3.12E-03 | 12 | 9570116 | 9581500 |
| Gm15484 | 23.810 | -2.525 | 0.718 | -3.518 | 4.35E-04 | 2.52E-02 | 5 | 110118022 | 110118849 |
| Pcdhgb1 | 2956.361 | -2.547 | 0.718 | -3.547 | 3.90E-04 | 2.34E-02 | 18 | 37680233 | 37841870 |
| Barx1 | 169.073 | -2.558 | 0.635 | -4.026 | 5.67E-05 | 6.58E-03 | 13 | 48662998 | 48666507 |
| Tbx15 | 1321.641 | -2.564 | 0.710 | -3.611 | 3.05E-04 | 1.99E-02 | 3 | 99240381 | 99354259 |
| Slc22a8 | 1178.304 | -2.571 | 0.687 | -3.743 | 1.82E-04 | 1.45E-02 | 19 | 8591254 | 8611835 |
| Myh2 | 2226.548 | -2.601 | 0.600 | -4.332 | 1.48E-05 | 2.40E-03 | 11 | 67171027 | 67197517 |
| Nphs2 | 256.207 | -2.633 | 0.689 | -3.821 | 1.33E-04 | 1.18E-02 | 1 | 156310727 | 156328035 |
| Vnn1 | 240.398 | -2.652 | 0.702 | -3.777 | 1.59E-04 | 1.35E-02 | 10 | 23894688 | 23905343 |
| Cul4a | 1078.443 | -2.685 | 0.602 | -4.461 | 8.15E-06 | 1.44E-03 | 8 | 13105621 | 13147940 |
| Mlf1 | 333.827 | -2.713 | 0.745 | -3.640 | 2.72E-04 | 1.88E-02 | 3 | 67374097 | 67400003 |
| AA986860 | 415.781 | -2.760 | 0.304 | -9.069 | 1.20E-19 | 2.54E-16 | 1 | 130731976 | 130744622 |
| Cnn1 | 222.655 | -2.765 | 0.572 | -4.835 | 1.33E-06 | 3.23E-04 | 9 | 22099216 | 22109630 |
| 9630028B13Rik | 109.156 | -2.786 | 0.811 | -3.436 | 5.90E-04 | 3.00E-02 | 1 | 185429355 | 185441819 |
| Trim5 | 78.820 | -2.814 | 0.366 | -7.689 | 1.48E-14 | 1.57E-11 | 7 | 104263386 | 104288094 |
| Slc6a20b | 137.265 | -2.866 | 0.760 | -3.770 | 1.63E-04 | 1.35E-02 | 9 | 123590800 | 123632565 |
| Mxd4 | 1329.783 | -2.910 | 0.765 | -3.805 | 1.42E-04 | 1.24E-02 | 5 | 34173883 | 34187720 |
| Ranbp3l | 873.469 | -2.922 | 0.730 | -4.005 | 6.21E-05 | 7.07E-03 | 15 | 8967949 | 9067335 |
| Tpm2 | 7722.136 | -2.926 | 0.659 | -4.437 | 9.12E-06 | 1.56E-03 | 4 | 43514711 | 43523765 |
| Trim72 | 247.359 | -2.930 | 0.890 | -3.293 | 9.91E-04 | 4.20E-02 | 7 | 128003949 | 128011033 |
| Agpat1 | 878.924 | -2.931 | 0.718 | -4.079 | 4.52E-05 | 5.53E-03 | 17 | 34604262 | 34613449 |
| Prss53 | 304.067 | -2.933 | 0.907 | -3.235 | 1.21E-03 | 4.76E-02 | 7 | 127885841 | 127890970 |
| Myom3 | 336.131 | -2.957 | 0.550 | -5.373 | 7.73E-08 | 2.38E-05 | 4 | 135759715 | 135815564 |
| 2310065F04Rik | 365.279 | -3.067 | 0.883 | -3.474 | 5.12E-04 | 2.76E-02 | 11 | 67112461 | 67120080 |
| Stac3 | 234.763 | -3.117 | 0.958 | -3.252 | 1.14E-03 | 4.61E-02 | 10 | 127501686 | 127508823 |
| Efhb | 36.990 | -3.130 | 0.677 | -4.624 | 3.76E-06 | 7.50E-04 | 17 | 53398889 | 53463321 |
| 1700016K19Rik | 65.950 | -3.131 | 0.810 | -3.864 | 1.12E-04 | 1.04E-02 | 11 | 75999912 | 76003569 |
| Slc5a9 | 31.577 | -3.158 | 0.931 | -3.392 | 6.95E-04 | 3.36E-02 | 4 | 111875375 | 111902918 |
| Fcgbp | 62.489 | -3.181 | 0.799 | -3.983 | 6.81E-05 | 7.43E-03 | 7 | 28071236 | 28120862 |
| Sypl2 | 497.486 | -3.298 | 0.914 | -3.608 | 3.08E-04 | 2.00E-02 | 3 | 108211472 | 108226648 |
| Gm1966 | 20.093 | -3.300 | 0.888 | -3.715 | 2.03E-04 | 1.59E-02 | 7 | 106596743 | 106604035 |
| Emx2 | 36.583 | -3.327 | 0.734 | -4.532 | 5.84E-06 | 1.07E-03 | 19 | 59458372 | 59465357 |
| Txlnb | 1181.822 | -3.345 | 0.811 | -4.126 | 3.68E-05 | 4.70E-03 | 10 | 17796226 | 17845665 |
| Muc1 | 246.676 | -3.350 | 0.969 | -3.458 | 5.45E-04 | 2.88E-02 | 3 | 89229057 | 89233381 |
| Alx3 | 111.642 | -3.461 | 0.602 | -5.753 | 8.75E-09 | 3.42E-06 | 3 | 107595031 | 107605776 |
| Cblc | 29.563 | -3.552 | 0.991 | -3.586 | 3.36E-04 | 2.14E-02 | 7 | 19778881 | 19796809 |
| Art1 | 468.705 | -3.562 | 0.939 | -3.794 | 1.48E-04 | 1.28E-02 | 7 | 102101743 | 102113933 |
| Gm5611 | 33.543 | -3.649 | 0.709 | -5.146 | 2.66E-07 | 7.48E-05 | 9 | 17030045 | 17030896 |

| | | | | | | | | | |
|---|---|---|---|---|---|---|---|---|---|
| Wdr64 | 40.037 | -3.655 | 0.613 | -5.961 | 2.50E-09 | 1.23E-06 | 1 | 175698593 | 175815734 |
| Atcayos | 146.770 | -3.722 | 0.915 | -4.068 | 4.73E-05 | 5.62E-03 | 10 | 81194609 | 81210877 |
| Gm9118 | 246.265 | -3.744 | 0.301 | -12.419 | 2.05E-35 | 9.80E-32 | 10 | 56497341 | 56498094 |
| Plet1 | 335.282 | -3.748 | 1.032 | -3.633 | 2.80E-04 | 1.90E-02 | 9 | 50494525 | 50505482 |
| Serpinb11 | 102.214 | -3.776 | 1.160 | -3.256 | 1.13E-03 | 4.60E-02 | 1 | 107361198 | 107380475 |
| Dusp13 | 130.580 | -3.796 | 1.072 | -3.540 | 4.00E-04 | 2.38E-02 | 14 | 21733394 | 21751181 |
| Lmod3 | 211.236 | -3.798 | 0.980 | -3.876 | 1.06E-04 | 1.02E-02 | 6 | 97238534 | 97252759 |
| Mb | 4497.486 | -3.806 | 0.935 | -4.071 | 4.67E-05 | 5.58E-03 | 15 | 77015487 | 77050670 |
| Gsta3 | 105.729 | -3.845 | 1.192 | -3.227 | 1.25E-03 | 4.87E-02 | 1 | 21240589 | 21265661 |
| Fitm1 | 243.816 | -3.872 | 1.122 | -3.450 | 5.61E-04 | 2.93E-02 | 14 | 55575617 | 55576954 |
| Acta1 | 13603.395 | -3.872 | 1.080 | -3.586 | 3.36E-04 | 2.14E-02 | 8 | 123891769 | 123894751 |
| Klhl31 | 248.438 | -3.951 | 1.114 | -3.546 | 3.92E-04 | 2.34E-02 | 9 | 77636500 | 77660127 |
| Lgr4 | 151.108 | -4.002 | 1.091 | -3.669 | 2.43E-04 | 1.76E-02 | 2 | 109917647 | 110014257 |
| Kcna7 | 226.156 | -4.003 | 1.045 | -3.831 | 1.28E-04 | 1.16E-02 | 7 | 45405653 | 45409763 |
| Wnt10b | 44.434 | -4.015 | 1.167 | -3.441 | 5.79E-04 | 2.98E-02 | 15 | 98770712 | 98778150 |
| Rwdd4a | 613.807 | -4.074 | 0.246 | -16.552 | 1.56E-61 | 1.49E-57 | 8 | 47533664 | 47552955 |
| Lrrc30 | 195.839 | -4.103 | 1.210 | -3.391 | 6.96E-04 | 3.36E-02 | 17 | 67630964 | 67632723 |
| Pcdhga2 | 4346.017 | -4.169 | 1.067 | -3.906 | 9.37E-05 | 9.23E-03 | 18 | 37668953 | 37841870 |
| Myf6 | 170.738 | -4.209 | 1.139 | -3.696 | 2.19E-04 | 1.65E-02 | 10 | 107492853 | 107494737 |
| Aspn | 38.314 | -4.212 | 1.277 | -3.298 | 9.73E-04 | 4.19E-02 | 13 | 49544443 | 49567565 |
| Ttn | 11247.891 | -4.301 | 1.301 | -3.305 | 9.49E-04 | 4.16E-02 | 2 | 76703980 | 76982547 |
| Erich2 | 82.022 | -4.347 | 1.337 | -3.252 | 1.15E-03 | 4.61E-02 | 2 | 70508819 | 70540884 |
| Aox2 | 130.415 | -4.348 | 1.091 | -3.985 | 6.76E-05 | 7.42E-03 | 1 | 58278326 | 58380259 |
| Spa17 | 88.875 | -4.418 | 0.564 | -7.836 | 4.66E-15 | 5.23E-12 | 9 | 37603290 | 37613722 |
| Stoml3 | 295.781 | -4.533 | 1.346 | -3.369 | 7.55E-04 | 3.57E-02 | 3 | 53488653 | 53508502 |
| Gm47908 | 24.818 | -4.665 | 1.284 | -3.633 | 2.80E-04 | 1.90E-02 | 10 | 33477115 | 33477472 |
| Yipf7 | 63.673 | -4.728 | 1.196 | -3.955 | 7.66E-05 | 7.91E-03 | 5 | 69516671 | 69542648 |
| Cfd | 100.331 | -4.993 | 1.193 | -4.185 | 2.85E-05 | 3.89E-03 | 10 | 79890853 | 79892655 |
| Myod1 | 30.329 | -5.226 | 1.620 | -3.225 | 1.26E-03 | 4.88E-02 | 7 | 46376474 | 46379099 |
| Pck1 | 225.052 | -5.242 | 1.323 | -3.964 | 7.38E-05 | 7.67E-03 | 2 | 173153048 | 173159273 |
| Scgb1c1 | 76.276 | -5.248 | 1.609 | -3.262 | 1.11E-03 | 4.52E-02 | 7 | 140845547 | 140846769 |
| Lpp | 218.915 | -5.261 | 0.907 | -5.803 | 6.53E-09 | 2.71E-06 | 16 | 24393507 | 24992578 |
| Megf10 | 78.449 | -5.554 | 1.547 | -3.589 | 3.32E-04 | 2.14E-02 | 18 | 57133090 | 57297467 |
| Mapk15 | 61.289 | -5.709 | 1.325 | -4.308 | 1.65E-05 | 2.54E-03 | 15 | 75993769 | 75999154 |
| Hivep2 | 1087.977 | -6.465 | 0.336 | -19.222 | 2.40E-82 | 4.59E-78 | 10 | 13966075 | 14151374 |
| Sult1b1 | 55.297 | -6.562 | 1.972 | -3.327 | 8.78E-04 | 3.96E-02 | 5 | 87513339 | 87538195 |
| E130008D07Rik | 57.670 | -6.848 | 1.178 | -5.815 | 6.08E-09 | 2.64E-06 | 17 | 43146041 | 43158291 |
| Mettl11b | 22.398 | -8.187 | 1.958 | -4.181 | 2.91E-05 | 3.90E-03 | 1 | 163702256 | 163725232 |
| Dnah6 | 24.714 | -8.329 | 2.576 | -3.234 | 1.22E-03 | 4.78E-02 | 6 | 73017606 | 73221651 |
| Vax2 | 26.358 | -8.422 | 2.122 | -3.969 | 7.20E-05 | 7.60E-03 | 6 | 83711264 | 83738313 |
| Ugt1a9 | 27.103 | -8.464 | 2.545 | -3.326 | 8.82E-04 | 3.96E-02 | 1 | 88070800 | 88218997 |
| Sftpd | 38.171 | -8.958 | 2.456 | -3.648 | 2.65E-04 | 1.85E-02 | 14 | 41172214 | 41185149 |
| Crybb1 | 45.678 | -9.216 | 2.443 | -3.772 | 1.62E-04 | 1.35E-02 | 5 | 112255815 | 112269585 |
| Abca8b | 46.657 | -9.246 | 1.737 | -5.324 | 1.01E-07 | 3.03E-05 | 11 | 109932190 | 109995845 |
| Lrrc71 | 47.408 | -9.269 | 2.589 | -3.581 | 3.42E-04 | 2.15E-02 | 3 | 87736923 | 87748625 |
| Bpifb3 | 82.548 | -10.071 | 1.609 | -6.261 | 3.83E-10 | 2.03E-07 | 2 | 153918230 | 153932996 |
| Gm20521 | 83.144 | -10.081 | 1.281 | -7.867 | 3.62E-15 | 4.32E-12 | 14 | 54883441 | 54898137 |

Table 5-4. Pathway analysis of responder mice

| p-value | FDR | pathway | source | external_id | members_input_overlap | members_input_overlap_geneids | effective_size |
|---|---|---|---|---|---|---|---|
| 1.09E-11 | 6.76E-10 | Hormone ligand-binding receptors | Reactome | R-HSA-375281 | CGA; TSHB; GNRHR; LHB; FSHB | 7252; 2798; 2488; 1081; 3972 | 13 |
| 3.02E-09 | 9.37E-08 | Glycoprotein hormones | Reactome | R-HSA-209822 | CGA; TSHB; LHB; FSHB | 7252; 2488; 1081; 3972 | 12 |
| 6.10E-09 | 1.26E-07 | Peptide hormone biosynthesis | Reactome | R-HSA-209952 | CGA; TSHB; LHB; FSHB | 7252; 2488; 1081; 3972 | 14 |
| 2.41E-07 | 3.73E-06 | Neuroactive ligand-receptor interaction - Homo sapiens (human) | KEGG | path:hsa04080 | GHRHR; LHB; FSHB; CGA; TSHB; SSTR3; GNRHR | 1081; 7252; 6753; 2692; 2798; 3972; 2488 | 277 |
| 5.21E-07 | 6.46E-06 | GPCR ligand binding | Reactome | R-HSA-500792 | GHRHR; LHB; FSHB; CGA; TSHB; NTS; SSTR3; GNRHR | 7252; 2692; 2798; 3972; 2488; 1081; 4922; 6753 | 465 |
| 7.86E-07 | 8.12E-06 | Class A/1 (Rhodopsin-like receptors) | Reactome | R-HSA-373076 | LHB; FSHB; CGA; TSHB; NTS; SSTR3; GNRHR | 2798; 3972; 1081; 7252; 4922; 6753; 2488 | 330 |
| 1.71E-06 | 1.51E-05 | Peptide hormone metabolism | Reactome | R-HSA-2980736 | CGA; TSHB; LHB; FSHB | 7252; 3972; 1081; 2488 | 53 |
| 1.63E-05 | 0.000126 | GnRH signaling pathway - Homo sapiens (human) | KEGG | path:hsa04912 | CGA; GNRHR; LHB; FSHB | 1081; 2798; 3972; 2488 | 93 |
| 4.09E-05 | 0.000282 | Mineralocorticoid biosynthesis | Reactome | R-HSA-193993 | CGA; LHB | 1081; 3972 | 6 |
| 6.41E-05 | 0.000397 | GPCR downstream signalling | Reactome | R-HSA-388396 | GHRHR; LHB; FSHB; CGA; TSHB; NTS; MCF2L; SSTR3; GNRHR | 7252; 23263; 2692; 6753; 1081; 2798; 4922; 3972; 2488 | 1171 |
| 7.38E-05 | 0.000416 | Ovarian steroidogenesis - Homo sapiens (human) | KEGG | path:hsa04913 | CGA; LHB; FSHB | 3972; 2488; 1081 | 49 |
| 9.67E-05 | 0.000445 | Signaling by GPCR | Reactome | R-HSA-372790 | GHRHR; LHB; FSHB; CGA; TSHB; NTS; MCF2L; SSTR3; GNRHR | 7252; 23263; 2692; 6753; 2798; 1081; 4922; 3972; 2488 | 1234 |
| 9.78E-05 | 0.000445 | FSH | NetPath | Pathway_FSH | CGA; FSHB | 2488; 1081 | 9 |
| 0.000122 | 0.000445 | Intracellular Signalling Through FSH Receptor and Follicle Stimulating Hormone | SMPDB | SMP00333 | CGA; FSHB | 1081; 2488 | 10 |
| 0.000122 | 0.000445 | Intracellular Signalling Through LHCGR Receptor and Luteinizing Hormone/Choriogonadotropin | SMPDB | SMP00338 | CGA; LHB | 1081; 3972 | 10 |
| 0.000122 | 0.000445 | Reactions specific to the complex N-glycan synthesis pathway | Reactome | R-HSA-975578 | CGA; LHB | 3972; 1081 | 10 |
| 0.000122 | 0.000445 | Thyroxine biosynthesis | Reactome | R-HSA-209968 | CGA; TSHB | 7252; 1081 | 10 |
| 0.000149 | 0.000514 | Androgen biosynthesis | Reactome | R-HSA-193048 | CGA; LHB | 1081; 3972 | 11 |
| 0.000392 | 0.001277 | Signal Transduction | Reactome | R-HSA-162582 | GHRHR; HDAC2; LHB; FSHB; CGA; TSHB; NTS; MCF2L; SSTR3; DLK1; POLR2B; GNRHR | 6753; 2488; 5431; 23263; 2692; 7252; 8788; 2798; 3972; 1081; 4922; 3066 | 2633 |
| 0.000412 | 0.001277 | Amine-derived hormones | Reactome | R-HSA-209776 | CGA; TSHB | 7252; 1081 | 18 |
| 0.000868 | 0.002293 | N-glycan antennae elongation in the medial/trans-Golgi | Reactome | R-HSA-975576 | CGA; LHB | 3972; 1081 | 26 |
| 0.000868 | 0.002293 | TSH | NetPath | Pathway_TSH | CGA; TSHB | 7252; 1081 | 26 |
| 0.000888 | 0.002293 | GPCR signaling-G alpha i | INOH | None | CGA; TSHB; LHB; FSHB | 3972; 1081; 7252; 2488 | 262 |
| 0.000888 | 0.002293 | GPCR signaling-pertussis toxin | INOH | None | CGA; TSHB; LHB; FSHB | 3972; 1081; 7252; 2488 | 262 |
| 0.000936 | 0.002322 | Follicle Stimulating Hormone (FSH) signaling pathway | Wikipathways | WP2035 | CGA; FSHB | 1081; 2488 | 27 |
| 0.000993 | 0.002322 | GPCR signaling-cholera toxin | INOH | None | CGA; TSHB; LHB; FSHB | 3972; 1081; 7252; 2488 | 270 |
| 0.001021 | 0.002322 | GPCR signaling-G alpha s Epac and ERK | INOH | None | CGA; TSHB; LHB; FSHB | 3972; 1081; 7252; 2488 | 272 |
| 0.001049 | 0.002322 | GPCR signaling-G alpha q | INOH | None | CGA; TSHB; LHB; FSHB | 1081; 7252; 3972; 2488 | 274 |
| 0.001214 | 0.002595 | GPCR signaling-G alpha s PKA and ERK | INOH | None | CGA; TSHB; LHB; FSHB | 3972; 2488; 1081; 7252 | 285 |
| 0.0014 | 0.002893 | Metabolism of steroid hormones | Reactome | R-HSA-196071 | CGA; LHB | 3972; 1081 | 33 |
| 0.00237 | 0.004721 | Steroid hormones | Reactome | R-HSA-209943 | CGA; LHB | 3972; 1081 | 43 |
| 0.002437 | 0.004721 | G alpha (s) signalling events | Reactome | R-HSA-418555 | CGA; TSHB; GHRHR; LHB; FSHB | 7252; 2692; 3972; 2488; 1081 | 580 |
| 0.003448 | 0.006478 | Autoimmune thyroid disease - Homo sapiens (human) | KEGG | path:hsa05320 | CGA; TSHB | 1081; 7252 | 52 |
| 0.003713 | 0.006771 | Regulation of lipolysis in adipocytes - Homo sapiens (human) | KEGG | path:hsa04923 | CGA; TSHB | 1081; 7252 | 54 |
| 0.004036 | 0.007149 | Huntington disease - Homo sapiens (human) | KEGG | path:hsa05016 | HDAC2; NDUFS5; POLR2B | 4725; 5431; 3066 | 193 |
| 0.005498 | 0.009469 | Human Thyroid Stimulating Hormone (TSH) signaling pathway | Wikipathways | WP2032 | CGA; TSHB | 1081; 7252 | 66 |
| 0.006165 | 0.01033 | Prolactin signaling pathway - Homo sapiens (human) | KEGG | path:hsa04917 | CGA; LHB | 3972; 1081 | 70 |
| 0.006688 | 0.010912 | Signaling by NOTCH1 | Reactome | R-HSA-1980143 | HDAC2; DLK1 | 8788; 3066 | 73 |
| 0.006867 | 0.010916 | Thyroid hormone synthesis - Homo sapiens (human) | KEGG | path:hsa04918 | CGA; TSHB | 1081; 7252 | 74 |
| 0.007048 | 0.010924 | Peptide GPCRs | Wikipathways | WP24 | SSTR3; GNRHR | 2798; 6753 | 75 |
| 0.007985 | 0.012074 | Glucocorticoid receptor regulatory network | PID | reg_gr_pathway | CGA; HDAC2 | 1081; 3066 | 80 |

Table 5-5. Gene ontology of responder mice

| p-value | FDR | term_goid | term_category | term_level | term_name | members_input_overlap | members_input_overlap_geneids | effective_size |
|---|---|---|---|---|---|---|---|---|
| 1.81E-07 | 1.69E-05 | GO:0016486 | b | 3 | peptide hormone processing | 1081; 2488; 3972; 7252 | CGA; TSHB; LHB; FSHB | 35 |
| 8.91E-07 | 1.69E-05 | GO:0005179 | m | 4 | hormone activity | 1081; 2488; 3972; 4922; 7252 | CGA; TSHB; NTS; LHB; FSHB | 124 |
| 2.04E-06 | 4.08E-05 | GO:0016914 | c | 3 | follicle-stimulating hormone complex | 1081; 2488 | CGA; FSHB | 2 |
| 2.04E-06 | 5.50E-05 | GO:0061696 | c | 2 | pituitary gonadotropin complex | 1081; 2488 | CGA; FSHB | 2 |
| 1.72E-05 | 0.000961 | GO:0042445 | b | 2 | hormone metabolic process | 1081; 2488; 2692; 3972; 7252 | CGA; TSHB; GHRHR; LHB; FSHB | 227 |
| 2.62E-05 | 0.00122 | GO:0009755 | b | 3 | hormone-mediated signaling pathway | 2488; 2692; 3972; 6753; 7252 | SSTR3; TSHB; GHRHR; LHB; FSHB | 248 |
| 9.57E-05 | 0.002965 | GO:0007186 | b | 3 | G protein-coupled receptor signaling pathway | 1081; 2488; 2692; 2798; 3972; 4922; 6753; 7252; 23263 | GHRHR; LHB; FSHB; CGA; TSHB; NTS; MCF2L; SSTR3; GNRHR | 1379 |
| 4.26E-05 | 0.003026 | GO:0006701 | b | 5 | progesterone biosynthetic process | 2488; 3972 | LHB; FSHB | 7 |
| 0.000138 | 0.003198 | GO:0010469 | b | 3 | regulation of signaling receptor activity | 1081; 2488; 3972; 4922; 7252; 7262 | LHB; FSHB; CGA; TSHB; NTS; PHLDA2 | 567 |
| 0.000615 | 0.005847 | GO:0048018 | m | 4 | receptor ligand activity | 1081; 2488; 3972; 4922; 7252 | CGA; TSHB; NTS; LHB; FSHB | 487 |
| 0.000442 | 0.007951 | GO:0005796 | c | 4 | Golgi lumen | 1081; 3972; 83452 | CGA; LHB; RAB33B | 103 |
| 0.000336 | 0.009406 | GO:0007165 | b | 2 | signal transduction | 1081; 2488; 2692; 2798; 3066; 3972; 4922; 5431; 6753; 7252; 7262; 8788; 10207; 11123; 133396; 83452; 83734; 23263 | GHRHR; HDAC2; LHB; IL31RA; FSHB; CGA; TSHB; NTS; RAB33B; MCF2L; ATG10; SSTR3; GNRHR; POLR2B; DLK1; PATJ; RCAN3; PHLDA2 | 6037 |
| 0.000546 | 0.010163 | GO:0071495 | b | 3 | cellular response to endogenous stimulus | 2488; 2692; 2798; 3066; 3972; 5431; 6753; 7252 | GHRHR; HDAC2; LHB; FSHB; TSHB; SSTR3; GNRHR; POLR2B | 1368 |
| 0.000871 | 0.013505 | GO:0010817 | b | 3 | regulation of hormone levels | 1081; 2488; 2692; 3972; 7252 | CGA; TSHB; GHRHR; LHB; FSHB | 526 |
| 0.000956 | 0.016253 | GO:0030545 | m | 2 | receptor regulator activity | 1081; 2488; 3972; 4922; 7252 | CGA; TSHB; NTS; LHB; FSHB | 537 |
| 0.000242 | 0.018034 | GO:0042448 | b | 4 | progesterone metabolic process | 2488; 3972 | LHB; FSHB | 16 |
| 0.000286 | 0.018034 | GO:0042127 | b | 4 | regulation of cell proliferation | 1081; 2488; 2692; 3066; 6753; 7224; 7262; 133396; 83452 | GHRHR; HDAC2; IL31RA; FSHB; CGA; RAB33B; SSTR3; TRPC5; PHLDA2 | 1592 |
| 0.000439 | 0.018425 | GO:0032870 | b | 4 | cellular response to hormone stimulus | 2488; 2692; 2798; 3972; 6753; 7252 | GHRHR; LHB; FSHB; TSHB; SSTR3; GNRHR | 703 |
| 0.000996 | 0.01859 | GO:0007154 | b | 2 | cell communication | 1081; 2488; 2692; 2798; 3066; 3972; 4922; 5431; 6753; 7252; 7262; 8788; 10207; 11123; 133396; 83452; 83734; 23263 | GHRHR; HDAC2; LHB; IL31RA; FSHB; CGA; TSHB; NTS; RAB33B; MCF2L; ATG10; SSTR3; GNRHR; POLR2B; DLK1; PATJ; RCAN3; PHLDA2 | 6538 |
| 0.001669 | 0.018737 | GO:0051716 | b | 2 | cellular response to stimulus | 1081; 2488; 2692; 2798; 3066; 3972; 4922; 5431; 6753; 7252; 7262; 8788; 9925; 10207; 11123; 133396; 83452; 83734; 23263 | GHRHR; HDAC2; LHB; ZBTB5; FSHB; CGA; TSHB; NTS; RAB33B; IL31RA; MCF2L; ATG10; SSTR3; GNRHR; POLR2B; DLK1; PATJ; RCAN3; PHLDA2 | 7448 |
| 0.001673 | 0.018737 | GO:0009719 | b | 2 | response to endogenous stimulus | 2488; 2692; 2798; 3066; 3972; 5431; 6753; 7252 | GHRHR; HDAC2; LHB; FSHB; TSHB; SSTR3; GNRHR; POLR2B | 1622 |
| 0.000648 | 0.020158 | GO:0006700 | b | 4 | C21-steroid hormone biosynthetic process | 2488; 3972 | LHB; FSHB | 26 |
| 0.0008 | 0.020158 | GO:0009966 | b | 4 | regulation of signal transduction | 1081; 2488; 2692; 3066; 3972; 4922; 7252; 7262; 8788; 11123; 133396; 23263 | GHRHR; HDAC2; LHB; IL31RA; FSHB; CGA; TSHB; NTS; MCF2L; DLK1; RCAN3; PHLDA2 | 3139 |
| 0.002291 | 0.029146 | GO:0023051 | b | 3 | regulation of signaling | 1081; 2488; 2692; 3066; 3972; 4922; 7252; 7262; 8788; 11123; 133396; 23263 | GHRHR; HDAC2; LHB; IL31RA; FSHB; CGA; TSHB; NTS; MCF2L; DLK1; RCAN3; PHLDA2 | 3526 |
| 0.002507 | 0.029146 | GO:0009725 | b | 3 | response to hormone | 2488; 2692; 2798; 3972; 6753; 7252 | GHRHR; LHB; FSHB; TSHB; SSTR3; GNRHR | 984 |
| 0.001461 | 0.03048 | GO:0008207 | b | 4 | C21-steroid hormone metabolic process | 2488; 3972 | LHB; FSHB | 39 |
| 0.001693 | 0.03048 | GO:0042181 | b | 4 | ketone biosynthetic process | 2488; 3972 | LHB; FSHB | 42 |
| 0.001211 | 0.031495 | GO:0016485 | b | 5 | protein processing | 1081; 2488; 3972; 7252 | CGA; TSHB; LHB; FSHB | 327 |
| 0.001744 | 0.031495 | GO:0008284 | b | 5 | positive regulation of cell proliferation | 1081; 2488; 2692; 3066; 7224; 133396 | GHRHR; HDAC2; IL31RA; FSHB; CGA; TRPC5 | 916 |
| 0.001774 | 0.031495 | GO:0021983 | b | 5 | pituitary gland development | 2692; 8022 | GHRHR; LHX3 | 43 |
| 0.002097 | 0.033035 | GO:0010646 | b | 4 | regulation of cell communication | 1081; 2488; 2692; 3066; 3972; 4922; 7252; 7262; 8788; 11123; 133396; 23263 | GHRHR; HDAC2; LHB; IL31RA; FSHB; CGA; TSHB; NTS; MCF2L; DLK1; RCAN3; PHLDA2 | 3491 |
| 0.003228 | 0.033358 | GO:0048608 | b | 3 | reproductive structure development | 2488; 3972; 7262; 8022 | LHX3; PHLDA2; LHB; FSHB | 428 |

| | | | | | | | | |
|---|---|---|---|---|---|---|---|---|
| 0.002462 | 0.034469 | GO:0051604 | b | 4 | protein maturation | 1081; 2488; 3972; 7252 | CGA; TSHB; LHB; FSHB | 397 |
| 0.002432 | 0.034529 | GO:0006694 | b | 5 | steroid biosynthetic process | 1081; 2488; 3972 | CGA; LHB; FSHB | 186 |
| 0.00331 | 0.040529 | GO:0061458 | b | 4 | reproductive system development | 2488; 3972; 7262; 8022 | LHX3; PHLDA2; LHB; FSHB | 431 |
| 0.003538 | 0.040529 | GO:0042733 | b | 4 | embryonic digit morphogenesis | 3066; 64327 | HDAC2; LMBR1 | 61 |
| 0.004893 | 0.044613 | GO:0005977 | b | 5 | glycogen metabolic process | 2548; 7262 | GAA; PHLDA2 | 72 |
| 0.005027 | 0.044613 | GO:0006073 | b | 5 | cellular glucan metabolic process | 2548; 7262 | GAA; PHLDA2 | 73 |
| 0.005027 | 0.044613 | GO:0044042 | b | 5 | glucan metabolic process | 2548; 7262 | GAA; PHLDA2 | 73 |
| 0.004893 | 0.045673 | GO:0043627 | b | 2 | response to estrogen | 2692; 7252 | TSHB; GHRHR | 72 |

Table 5-6. Pathway analysis of nonresponder mice

| p-value | FDR | pathway | source | external_id | members_input_overlap | members_input_overlap_geneids | effective_size |
|---|---|---|---|---|---|---|---|
| 3.15E-05 | 0.004726 | Transcriptional regulation of white adipocyte differentiation | Wikipathways | WP2751 | PCK1; FABP4; SLC2A4 | 6517; 2167; 5105 | 15 |
| 0.000173 | 0.008487 | Integrin | INOH | None | VAV1; COL9A2; COL12A1; FN1; COL1A1 | 1277; 1298; 2335; 7409; 1303 | 124 |
| 0.000224 | 0.008487 | Adipogenesis | Wikipathways | WP236 | PCK1; CFD; SLC2A4; WNT10B; PTGIS | 5740; 1675; 6517; 7480; 5105 | 131 |
| 0.000226 | 0.008487 | Extracellular matrix organization | Reactome | R-HSA-1474244 | BMP7; FN1; ASPN; COL9A2; FMOD; COL1A1; COL12A1 | 54829; 1277; 1298; 2335; 2331; 655; 1303 | 294 |
| 0.000841 | 0.025221 | Collagen chain trimerization | Reactome | R-HSA-8948216 | COL9A2; COL12A1; COL1A1 | 1277; 1303; 1298 | 44 |
| 0.001498 | 0.03727 | Focal adhesion - Homo sapiens (human) | KEGG | path:hsa04510 | VAV1; COL1A1; COL9A2; FN1; MYL9 | 10398; 2335; 7409; 1277; 1298 | 199 |
| 0.001787 | 0.03727 | ECM proteoglycans | Reactome | R-HSA-3000178 | ASPN; COL9A2; FMOD | 54829; 2331; 1298 | 57 |
| 0.002382 | 0.03727 | Arachidonic acid metabolism - Homo sapiens (human) | KEGG | path:hsa00590 | GPX8; PLA2G4A; PTGIS | 493869; 5321; 5740 | 63 |
| 0.002577 | 0.03727 | miR-509-3p alteration of YAP1-ECM axis | Wikipathways | WP3967 | FN1; COL1A1 | 2335; 1277 | 18 |
| 0.002873 | 0.03727 | Overview of nanoparticle effects | Wikipathways | WP3287 | CDH3; FN1 | 2335; 1001 | 19 |
| 0.002961 | 0.03727 | Collagen biosynthesis and modifying enzymes | Reactome | R-HSA-1650814 | COL9A2; COL12A1; COL1A1 | 1277; 1303; 1298 | 68 |
| 0.002982 | 0.03727 | PI3K-Akt Signaling Pathway | Wikipathways | WP4172 | PCK1; FN1; COL9A2; IL3RA; COL1A1; LPAR4 | 2846; 3563; 5105; 1277; 2335; 1298 | 339 |
| 0.00364 | 0.039628 | PI3K-Akt signaling pathway - Homo sapiens (human) | KEGG | path:hsa04151 | PCK1; FN1; COL9A2; IL3RA; COL1A1; LPAR4 | 2335; 2846; 5105; 1277; 1298; 3563 | 353 |
| 0.003908 | 0.039628 | Transcriptional regulation by RUNX2 | Reactome | R-HSA-8878166 | DLX5; SP7; COL1A1 | 1277; 121340; 1749 | 75 |
| 0.004205 | 0.039628 | Eicosanoid Synthesis | Wikipathways | WP167 | PLA2G4A; PTGIS | 5740; 5321 | 23 |
| 0.004574 | 0.039628 | RUNX2 regulates osteoblast differentiation | Reactome | R-HSA-8940973 | SP7; COL1A1 | 1277; 121340 | 24 |
| 0.004753 | 0.039628 | Platelet activation, signaling and aggregation | Reactome | R-HSA-76002 | CLEC3B; VAV1; PLA2G4A; CFD; FN1 | 7123; 2335; 5321; 1675; 7409 | 260 |
| 0.004959 | 0.039628 | VEGFR3 signaling in lymphatic endothelium | PID | lymphangiogenesis_pathway | FN1; COL1A1 | 1277; 2335 | 25 |
| 0.00502 | 0.039628 | ECM-receptor interaction - Homo sapiens (human) | KEGG | path:hsa04512 | COL9A2; FN1; COL1A1 | 1298; 2335; 1277 | 82 |
| 0.006198 | 0.043184 | Keratan sulfate biosynthesis | Reactome | R-HSA-2022854 | FMOD; OMD | 4958; 2331 | 28 |
| 0.006502 | 0.043184 | Protein digestion and absorption - Homo sapiens (human) | KEGG | path:hsa04974 | COL9A2; COL12A1; COL1A1 | 1277; 1298; 1303 | 90 |
| 0.006502 | 0.043184 | Fc gamma R-mediated phagocytosis - Homo sapiens (human) | KEGG | path:hsa04666 | VAV1; PLA2G4A; MARCKS | 5321; 7409; 4082 | 90 |
| 0.006704 | 0.043184 | Amino Acid metabolism | Wikipathways | WP3925 | AOC3; PCK1; ASNS | 8639; 5105; 440 | 91 |
| 0.006909 | 0.043184 | Collagen formation | Reactome | R-HSA-1474290 | COL9A2; COL12A1; COL1A1 | 1277; 1298; 1303 | 92 |
| 0.008045 | 0.045647 | RUNX2 regulates bone development | Reactome | R-HSA-8941326 | SP7; COL1A1 | 1277; 121340 | 32 |
| 0.008541 | 0.045647 | Inflammatory Response Pathway | Wikipathways | WP453 | FN1; COL1A1 | 1277; 2335 | 33 |
| 0.008541 | 0.045647 | Keratan sulfate/keratin metabolism | Reactome | R-HSA-1638074 | FMOD; OMD | 4958; 2331 | 33 |
| 0.00884 | 0.045647 | Focal Adhesion-PI3K-Akt-mTOR-signaling pathway | Wikipathways | WP3932 | SLC2A4; IL3RA; FN1; LPAR4; COL1A1 | 2846; 3563; 1277; 2335; 6517 | 302 |
| 0.00905 | 0.045647 | BMP signaling Dro | INOH | None | BMP7; WT1 | 655; 7490 | 34 |
| 0.009573 | 0.045647 | Role of Osx and miRNAs in tooth development | Wikipathways | WP3971 | BMP7; SP7 | 655; 121340 | 35 |
| 0.009573 | 0.045647 | Collagen degradation | Reactome | R-HSA-1442490 | COL9A2; COL12A1 | 1303; 1298 | 35 |
| 0.009738 | 0.045647 | Focal Adhesion | Wikipathways | WP306 | VAV1; COL1A1; FN1; MYL9 | 10398; 2335; 7409; 1277 | 198 |

Table 5-7. Gene ontology of nonresponder mice

| p-value | FDR | term_goid | term_category | term_level | term_name | members_input_overlap | members_input_overlap_geneids | effective_size |
|---|---|---|---|---|---|---|---|---|
| 1.55E-11 | 4.33E-10 | GO:0062023 | c | 3 | collagen-containing extracellular matrix | 655; 1277; 1298; 1303; 1805; 2331; 2335; 4958; 165; 7123; 10631; 115908; 284297; 54829; 23452 | BMP7; DPT; FN1; OMD; CLEC3B; ASPN; SSC5D; COL9A2; POSTN; AEBP1; ANGPTL2; FMOD; COL1A1; CTHRC1; COL12A1 | 374 |
| 6.62E-10 | 1.79E-08 | GO:0005201 | m | 2 | extracellular matrix structural constituent | 1277; 1298; 1303; 1805; 2331; 2335; 165; 10631; 115908; 54829 | DPT; FN1; ASPN; COL9A2; POSTN; AEBP1; FMOD; COL1A1; CTHRC1; COL12A1 | 159 |
| 9.60E-10 | 3.27E-08 | GO:0031012 | c | 2 | extracellular matrix | 655; 1277; 1298; 1303; 1805; 2331; 2335; 4958; 165; 7123; 10631; 115908; 284297; 54829; 23452 | BMP7; DPT; FN1; OMD; CLEC3B; ASPN; SSC5D; COL9A2; POSTN; AEBP1; ANGPTL2; FMOD; COL1A1; CTHRC1; COL12A1 | 503 |
| 3.75E-10 | 8.25E-08 | GO:0030282 | b | 5 | bone mineralization | 655; 4958; 5167; 7123; 7480; 338773; 144347; 54829; 130497 | BMP7; OMD; CLEC3B; WNT10B; OSR1; ENPP1; TMEM119; ASPN; RFLNA | 107 |
| 4.54E-10 | 1.47E-07 | GO:0031214 | b | 4 | biomineral tissue development | 655; 1277; 4958; 5167; 7123; 7480; 338773; 144347; 54829; 130497 | BMP7; OMD; CLEC3B; WNT10B; OSR1; ENPP1; TMEM119; COL1A1; ASPN; RFLNA | 153 |
| 2.16E-09 | 1.83E-07 | GO:0001503 | b | 2 | ossification | 655; 121340; 1277; 1749; 4958; 5167; 7123; 7480; 338773; 144347; 115908; 54829; 130497 | BMP7; OMD; CLEC3B; WNT10B; CTHRC1; OSR1; ENPP1; TMEM119; ASPN; COL1A1; DLX5; SP7; RFLNA | 372 |
| 1.83E-09 | 2.02E-07 | GO:0070167 | b | 5 | regulation of biomineral tissue development | 655; 4958; 5167; 7480; 338773; 144347; 54829; 130497 | BMP7; OMD; WNT10B; OSR1; ENPP1; TMEM119; ASPN; RFLNA | 87 |
| 3.38E-09 | 6.73E-07 | GO:0048513 | b | 3 | animal organ development | 257; 440; 655; 1001; 1036; 121340; 1277; 1303; 1749; 2018; 2042; 2335; 60529; 2828; 4958; 5105; 5167; 6913; 7123; 7135; 7409; 7480; 7490; 7827; 5740; 9452; 10398; 338773; 56033; 144347; 115908; 54829; 55079; 26471; 126014; 130497 | PCK1; EPHA3; CDO1; TMEM119; MYL9; FEZF2; ALX3; OSCAR; WNT10B; ALX4; TBX15; TNNI1; DLX5; CDH3; ASPN; WT1; RFLNA; NUPR1; OSR1; PTGIS; ENPP1; BARX1; NPHS2; SP7; BMP7; FN1; GPR4; VAV1; OMD; EMX2; ASNS; CLEC3B; ITM2A; COL1A1; CTHRC1; COL12A1 | 3442 |
| 1.10E-08 | 8.10E-07 | GO:0030500 | b | 5 | regulation of bone mineralization | 655; 4958; 5167; 7480; 338773; 144347; 130497 | BMP7; OMD; WNT10B; OSR1; ENPP1; TMEM119; RFLNA | 70 |
| 1.19E-08 | 1.18E-06 | GO:0009888 | b | 3 | tissue development | 655; 1001; 1277; 1303; 1749; 2042; 2335; 60529; 2828; 4958; 5105; 5167; 7123; 7135; 7480; 7490; 7827; 5740; 10631; 338773; 56033; 144347; 115908; 54829; 26471; 130497 | PCK1; EPHA3; TMEM119; CLEC3B; WNT10B; ALX4; TNNI1; DLX5; CDH3; ASPN; WT1; RFLNA; NUPR1; OSR1; PTGIS; ENPP1; POSTN; BARX1; NPHS2; BMP7; FN1; GPR4; OMD; COL1A1; CTHRC1; COL12A1 | 1951 |
| 3.22E-08 | 5.20E-06 | GO:0048705 | b | 4 | skeletal system morphogenesis | 257; 655; 1277; 1303; 1749; 60529; 6913; 7480; 144347; 130497 | BMP7; ALX3; WNT10B; OSR1; ALX4; TBX15; COL1A1; DLX5; RFLNA; COL12A1 | 238 |
| 1.07E-07 | 7.11E-06 | GO:0048731 | b | 3 | system development | 257; 440; 655; 1001; 1036; 121340; 1277; 1298; 1303; 1749; 2018; 2042; 2335; 60529; 2828; 4958; 5105; 5167; 6913; 7123; 7135; 7409; 7480; 7490; 7827; 5740; 9452; 10398; 10631; 338773; 56033; 144347; 7851; 342035; 115908; 54829; 55079; 26471; 126014; 130497 | PCK1; EPHA3; CDO1; TMEM119; MYL9; FEZF2; GLDN; ALX3; OSCAR; WNT10B; ALX4; MALL; TBX15; TNNI1; DLX5; CDH3; ASPN; WT1; RFLNA; NUPR1; COL9A2; OSR1; PTGIS; ENPP1; POSTN; BARX1; NPHS2; SP7; BMP7; FN1; GPR4; VAV1; OMD; EMX2; ASNS; CLEC3B; ITM2A; COL1A1; CTHRC1; COL12A1 | 4699 |
| 1.03E-07 | 1.11E-05 | GO:0001501 | b | 4 | skeletal system development | 257; 655; 1277; 1298; 1303; 1749; 60529; 6913; 7480; 10631; 338773; 144347; 130497 | BMP7; TMEM119; ALX3; WNT10B; COL9A2; OSR1; ALX4; POSTN; TBX15; COL1A1; DLX5; RFLNA; COL12A1 | 516 |
| 2.73E-07 | 1.16E-05 | GO:0007275 | b | 2 | multicellular organism development | 257; 440; 655; 1001; 1036; 121340; 1277; 1298; 1303; 1749; 2018; 2042; 2335; 60529; 2828; 4958; 5105; 5167; 6913; 7123; 7135; 7409; 7480; 7490; 7827; 5740; 9452; 10398; 10631; 338773; 56033; 144347; 7851; 342035; 115908; 284297; 54829; 55079; 23452; 26471; 126014; 130497 | PCK1; EPHA3; CDO1; TMEM119; MYL9; FEZF2; GLDN; ALX3; OSCAR; WNT10B; ALX4; MALL; TBX15; TNNI1; DLX5; CDH3; ASPN; WT1; SSC5D; RFLNA; NUPR1; COL9A2; OSR1; PTGIS; ENPP1; POSTN; BARX1; NPHS2; SP7; BMP7; FN1; GPR4; VAV1; OMD; EMX2; ASNS; CLEC3B; ITM2A; ANGPTL2; COL1A1; CTHRC1; COL12A1 | 5275 |
| 6.20E-07 | 5.01E-05 | GO:0030278 | b | 4 | regulation of ossification | 655; 4958; 5167; 7480; 338773; 144347; 115908; 130497 | BMP7; OMD; WNT10B; OSR1; ENPP1; TMEM119; CTHRC1; RFLNA | 183 |

| | | | | | | | | |
|---|---|---|---|---|---|---|---|---|
| 4.11E-06 | 6.98E-05 | GO:0005615 | c | 2 | extracellular space | 655; 1277; 1298; 1303; 1675; 1805; 2167; 2331; 2335; 3598; 4082; 4958; 165; 5105; 5167; 5349; 5375; 5480; 6517; 7123; 7480; 7827; 5740; 10631; 342035; 115908; 284297; 8857; 23452; 126014 | DPT; FXYD3; FABP4; PMP2; AEBP1; GLDN; CLEC3B; OSCAR; WNT10B; MARCKS; FMOD; SSC5D; COL9A2; PTGIS; ENPP1; POSTN; NPHS2; FCGBP; BMP7; IL13RA2; FN1; OMD; CFD; SLC2A4; PCK1; ANGPTL2; COL1A1; PPIC; CTHRC1; COL12A1 | 3339 |
| 1.71E-06 | 0.000101 | GO:0030199 | b | 4 | collagen fibril organization | 1277; 1303; 1805; 2331; 165 | DPT; AEBP1; FMOD; COL12A1; COL1A1 | 51 |
| 1.87E-06 | 0.000101 | GO:0072012 | b | 4 | glomerulus vasculature development | 655; 2828; 7490; 130497 | BMP7; WT1; OSR1; GPR4 | 23 |
| 3.77E-06 | 0.000107 | GO:0048856 | b | 2 | anatomical structure development | 257; 440; 655; 1001; 1036; 121340; 1277; 1298; 1303; 1749; 2018; 2042; 2335; 60529; 2828; 4958; 5105; 5167; 6913; 7123; 7135; 7409; 7480; 7490; 7827; 5740; 9452; 10398; 10631; 338773; 56033; 144347; 7851; 342035; 115908; 284297; 54829; 55079; 23452; 26471; 126014; 130497 | PCK1; EPHA3; CDO1; TMEM119; MYL9; FEZF2; GLDN; ALX3; OSCAR; WNT10B; ALX4; MALL; TBX15; TNNI1; DLX5; CDH3; ASPN; WT1; SSC5D; RFLNA; NUPR1; COL9A2; OSR1; PTGIS; ENPP1; POSTN; BARX1; NPHS2; SP7; BMP7; FN1; GPR4; VAV1; OMD; EMX2; ASNS; CLEC3B; ITM2A; ANGPTL2; COL1A1; CTHRC1; COL12A1 | 5773 |
| 2.66E-06 | 0.000146 | GO:0061437 | b | 5 | renal system vasculature development | 655; 2828; 7490; 130497 | BMP7; WT1; OSR1; GPR4 | 25 |
| 3.86E-06 | 0.000164 | GO:0032835 | b | 3 | glomerulus development | 655; 2828; 7490; 7827; 130497 | BMP7; WT1; NPHS2; OSR1; GPR4 | 60 |
| 4.11E-06 | 0.000164 | GO:0043062 | b | 3 | extracellular structure organization | 655; 1277; 1298; 1303; 1805; 2331; 2335; 165; 7490; 10631 | BMP7; WT1; DPT; FN1; COL9A2; POSTN; AEBP1; FMOD; COL1A1; COL12A1 | 404 |
| 6.95E-06 | 0.00023 | GO:0072203 | b | 3 | cell proliferation involved in metanephros development | 655; 7490; 130497 | BMP7; WT1; OSR1 | 10 |
| 2.54E-05 | 0.000288 | GO:0005581 | c | 2 | collagen trimer | 1277; 1298; 1303; 342035; 115908 | GLDN; COL9A2; CTHRC1; COL12A1; COL1A1 | 88 |
| 8.30E-06 | 0.000365 | GO:0071300 | b | 5 | cellular response to retinoic acid | 1277; 2042; 5105; 7480; 130497 | WNT10B; PCK1; EPHA3; OSR1; COL1A1 | 70 |
| 1.05E-05 | 0.000384 | GO:0048706 | b | 5 | embryonic skeletal system development | 257; 655; 1277; 60529; 6913; 130497 | BMP7; ALX3; OSR1; ALX4; TBX15; COL1A1 | 126 |
| 9.31E-06 | 0.000415 | GO:0030198 | b | 4 | extracellular matrix organization | 1277; 1298; 1303; 1805; 2331; 2335; 165; 7490; 10631 | WT1; DPT; FN1; COL9A2; POSTN; AEBP1; FMOD; COL1A1; COL12A1 | 349 |
| 1.03E-05 | 0.000415 | GO:0061448 | b | 4 | connective tissue development | 655; 1277; 1303; 2828; 7480; 7490; 144347; 130497 | BMP7; WT1; GPR4; WNT10B; OSR1; COL1A1; RFLNA; COL12A1 | 267 |
| 1.54E-05 | 0.000483 | GO:0072166 | b | 5 | posterior mesonephric tubule development | 7490; 130497 | WT1; OSR1 | 2 |
| 1.80E-05 | 0.000496 | GO:0045778 | b | 5 | positive regulation of ossification | 655; 7480; 338773; 115908; 130497 | BMP7; TMEM119; WNT10B; CTHRC1; OSR1 | 82 |
| 2.08E-05 | 0.000509 | GO:0072109 | b | 5 | glomerular mesangium development | 655; 2828; 7490 | BMP7; WT1; GPR4 | 14 |
| 1.54E-05 | 0.000553 | GO:0001649 | b | 4 | osteoblast differentiation | 655; 121340; 1277; 1749; 7480; 338773; 115908 | BMP7; WNT10B; SP7; TMEM119; COL1A1; DLX5; CTHRC1 | 203 |
| 2.70E-05 | 0.000593 | GO:0035850 | b | 5 | epithelial cell differentiation involved in kidney development | 2828; 7490; 7827; 130497 | WT1; NPHS2; OSR1; GPR4 | 44 |
| 3.22E-05 | 0.000641 | GO:0070169 | b | 5 | positive regulation of biomineral tissue development | 655; 7480; 338773; 130497 | BMP7; TMEM119; WNT10B; OSR1 | 46 |
| 3.50E-05 | 0.000641 | GO:0048704 | b | 5 | embryonic skeletal system morphogenesis | 257; 655; 60529; 6913; 130497 | ALX3; BMP7; OSR1; TBX15; ALX4 | 94 |
| 2.53E-05 | 0.000718 | GO:0009887 | b | 3 | animal organ morphogenesis | 257; 655; 1277; 1303; 1749; 60529; 6913; 7135; 7480; 7490; 144347; 115908; 54829; 130497 | BMP7; WT1; ALX3; WNT10B; TNNI1; CTHRC1; OSR1; ALX4; TBX15; COL1A1; DLX5; ASPN; RFLNA; COL12A1 | 979 |
| 4.62E-05 | 0.000782 | GO:0090185 | b | 5 | negative regulation of kidney development | 655; 7490; 130497 | BMP7; WT1; OSR1 | 18 |
| 5.47E-05 | 0.00086 | GO:0072074 | b | 5 | kidney mesenchyme development | 655; 7490; 130497 | BMP7; WT1; OSR1 | 19 |
| 3.19E-05 | 0.00103 | GO:0072224 | b | 4 | metanephric glomerulus development | 7490; 7827; 130497 | WT1; NPHS2; OSR1 | 16 |
| 9.18E-05 | 0.001346 | GO:0072131 | b | 5 | kidney mesenchyme morphogenesis | 655; 130497 | BMP7; OSR1 | 4 |

| | | | | | | | | |
|---|---|---|---|---|---|---|---|---|
| 6.42E-05 | 0.001364 | GO:0072111 | b | 2 | cell proliferation involved in kidney development | 655; 7490; 130497 | BMP7; WT1; OSR1 | 20 |
| 4.71E-05 | 0.001383 | GO:0051591 | b | 4 | response to cAMP | 1036; 1277; 5105; 7480; 7490 | WT1; WNT10B; PCK1; CDO1; COL1A1 | 100 |
| 6.10E-05 | 0.001617 | GO:0061005 | b | 4 | cell differentiation involved in kidney development | 2828; 7490; 7827; 130497 | WT1; NPHS2; OSR1; GPR4 | 54 |
| 6.51E-05 | 0.001617 | GO:0032526 | b | 4 | response to retinoic acid | 1277; 2042; 5105; 7480; 130497 | WNT10B; PCK1; EPHA3; OSR1; COL1A1 | 107 |
| 0.000136 | 0.001869 | GO:0030326 | b | 5 | embryonic limb morphogenesis | 257; 655; 1749; 60529; 130497 | ALX3; BMP7; ALX4; OSR1; DLX5 | 125 |
| 0.000104 | 0.002589 | GO:0070887 | b | 3 | cellular response to chemical stimulus | 440; 655; 1036; 1277; 1749; 2042; 2167; 2331; 2335; 493869; 3563; 3598; 5105; 5167; 5321; 6517; 7123; 7409; 7480; 7490; 5740; 8639; 10631; 85363; 54829; 130497 | AOC3; PCK1; EPHA3; CDO1; FABP4; CLEC3B; WNT10B; FMOD; DLX5; WT1; OSR1; PTGIS; ENPP1; POSTN; IL3RA; PLA2G4A; BMP7; IL13RA2; FN1; VAV1; SLC2A4; ASNS; COL1A1; GPX8; TRIM5; ASPN | 3141 |
| 0.000166 | 0.002796 | GO:0042221 | b | 2 | response to chemical | 440; 655; 1001; 1036; 1277; 1749; 2018; 2042; 2167; 2331; 2335; 493869; 3563; 3598; 5105; 5167; 5321; 6517; 7123; 7409; 7480; 7490; 5740; 8639; 10631; 10941; 284297; 85363; 54829; 55079; 26266; 26471; 130497 | AOC3; PCK1; EPHA3; CDO1; UGT2A1; FABP4; FEZF2; CLEC3B; WNT10B; FMOD; DLX5; CDH3; WT1; SSC5D; NUPR1; SLC13A4; OSR1; PTGIS; ENPP1; POSTN; IL3RA; PLA2G4A; BMP7; IL13RA2; FN1; VAV1; SLC2A4; EMX2; ASNS; COL1A1; GPX8; TRIM5; ASPN | 4644 |
| 0.000197 | 0.002796 | GO:0048869 | b | 2 | cellular developmental process | 655; 1001; 121340; 1277; 1303; 1749; 2018; 2042; 2167; 2335; 2828; 5105; 5167; 6517; 7409; 7480; 7490; 7827; 9452; 10398; 10631; 338773; 56033; 144347; 342035; 115908; 83719; 55079; 26471; 126014; 130497 | PCK1; EPHA3; TMEM119; FABP4; MYL9; YPEL3; GLDN; OSCAR; WNT10B; DLX5; CDH3; WT1; RFLNA; NUPR1; OSR1; SP7; POSTN; BARX1; NPHS2; ENPP1; BMP7; FEZF2; FN1; GPR4; VAV1; SLC2A4; EMX2; ITM2A; COL1A1; CTHRC1; COL12A1 | 4265 |
| 0.000129 | 0.002927 | GO:0048562 | b | 4 | embryonic organ morphogenesis | 257; 655; 1749; 60529; 6913; 115908; 130497 | BMP7; ALX3; OSR1; ALX4; TBX15; DLX5; CTHRC1 | 284 |
| 0.000136 | 0.002927 | GO:0035113 | b | 4 | embryonic appendage morphogenesis | 257; 655; 1749; 60529; 130497 | ALX3; BMP7; ALX4; OSR1; DLX5 | 125 |
| 0.000228 | 0.002953 | GO:0072007 | b | 5 | mesangial cell differentiation | 2828; 130497 | OSR1; GPR4 | 6 |
| 0.000153 | 0.00301 | GO:0072143 | b | 4 | mesangial cell development | 2828; 130497 | OSR1; GPR4 | 5 |
| 0.000163 | 0.00301 | GO:2000026 | b | 4 | regulation of multicellular organismal development | 655; 1001; 1277; 2042; 2335; 2828; 4958; 5167; 7480; 7490; 5740; 10398; 338773; 144347; 115908; 54829; 55079; 26471; 130497 | BMP7; WT1; NUPR1; FEZF2; FN1; GPR4; WNT10B; OMD; EPHA3; OSR1; PTGIS; ENPP1; TMEM119; CTHRC1; COL1A1; CDH3; ASPN; RFLNA; MYL9 | 1957 |
| 0.000168 | 0.00301 | GO:0051216 | b | 4 | cartilage development | 655; 1277; 1303; 7480; 144347; 130497 | BMP7; WNT10B; OSR1; COL1A1; RFLNA; COL12A1 | 207 |
| 0.000186 | 0.003164 | GO:0009952 | b | 4 | anterior/posterior pattern specification | 2018; 60529; 7490; 56033; 55079; 130497 | WT1; FEZF2; EMX2; OSR1; ALX4; BARX1 | 211 |
| 0.000228 | 0.003195 | GO:0005593 | c | 3 | FACIT collagen trimer | 1298; 1303 | COL9A2; COL12A1 | 6 |
| 0.000216 | 0.003486 | GO:0046683 | b | 4 | response to organophosphorus | 1036; 1277; 5105; 7480; 7490 | WT1; WNT10B; PCK1; CDO1; COL1A1 | 138 |
| 0.000318 | 0.003859 | GO:0007389 | b | 2 | pattern specification process | 257; 655; 2018; 60529; 7490; 56033; 55079; 130497 | BMP7; WT1; FEZF2; ALX3; EMX2; OSR1; ALX4; BARX1 | 436 |
| 0.000179 | 0.003906 | GO:0050793 | b | 3 | regulation of developmental process | 655; 1001; 121340; 1277; 2042; 2335; 2828; 4958; 5167; 7480; 7490; 5740; 10398; 10631; 338773; 144347; 115908; 83719; 54829; 55079; 26471; 130497 | EPHA3; TMEM119; MYL9; YPEL3; WNT10B; CDH3; ASPN; WT1; RFLNA; NUPR1; OSR1; PTGIS; SP7; POSTN; ENPP1; BMP7; FEZF2; FN1; GPR4; OMD; COL1A1; CTHRC1 | 2497 |
| 0.000202 | 0.003906 | GO:0072006 | b | 3 | nephron development | 655; 2828; 7490; 7827; 130497 | BMP7; WT1; NPHS2; OSR1; GPR4 | 136 |
| 0.000216 | 0.003906 | GO:0030154 | b | 3 | cell differentiation | 655; 1001; 121340; 1277; 1303; 1749; 2018; 2042; 2167; 2335; 2828; 5105; 5167; 6517; 7409; 7480; 7490; 7827; 9452; 10398; 10631; 338773; 56033; 144347; 342035; 115908; 55079; 26471; 126014; 130497 | PCK1; EPHA3; TMEM119; FABP4; MYL9; FEZF2; GLDN; OSCAR; WNT10B; DLX5; CDH3; WT1; RFLNA; NUPR1; OSR1; SP7; POSTN; BARX1; NPHS2; ENPP1; BMP7; FN1; GPR4; VAV1; SLC2A4; EMX2; ITM2A; COL1A1; CTHRC1; COL12A1 | 4079 |
| 0.000327 | 0.004003 | GO:0014074 | b | 5 | response to purine-containing compound | 1036; 1277; 5105; 7480; 7490 | WT1; WNT10B; PCK1; CDO1; COL1A1 | 151 |
| 0.000384 | 0.004445 | GO:0035137 | b | 5 | hindlimb morphogenesis | 257; 60529; 130497 | ALX3; OSR1; ALX4 | 36 |

| | | | | | | | | |
|---|---|---|---|---|---|---|---|---|
| 0.000424 | 0.004445 | GO:0072239 | b | 5 | metanephric glomerulus vasculature development | 7490; 130497 | WT1; OSR1 | 8 |
| 0.000424 | 0.004445 | GO:0001656 | b | 5 | metanephros development | 655; 7490; 7827; 130497 | BMP7; WT1; NPHS2; OSR1 | 89 |
| 0.000289 | 0.00445 | GO:0035108 | b | 4 | limb morphogenesis | 257; 655; 1749; 60529; 130497 | ALX3; BMP7; ALX4; OSR1; DLX5 | 147 |
| 0.000314 | 0.004476 | GO:0071310 | b | 4 | cellular response to organic substance | 440; 655; 1277; 1749; 2042; 2167; 2331; 2335; 3563; 3598; 5105; 5167; 6517; 7123; 7409; 7480; 7490; 5740; 10631; 85363; 54829; 130497 | PCK1; EPHA3; FABP4; CLEC3B; WNT10B; FMOD; DLX5; WT1; OSR1; PTGIS; ENPP1; POSTN; IL3RA; BMP7; IL13RA2; FN1; VAV1; SLC2A4; ASNS; COL1A1; TRIM5; ASPN | 2596 |
| 0.000319 | 0.004476 | GO:1904238 | b | 4 | pericyte cell differentiation | 2828; 130497 | OSR1; GPR4 | 7 |
| 0.000289 | 0.004798 | GO:0035107 | b | 3 | appendage morphogenesis | 257; 655; 1749; 60529; 130497 | ALX3; BMP7; ALX4; OSR1; DLX5 | 147 |
| 0.000565 | 0.005653 | GO:0035136 | b | 5 | forelimb morphogenesis | 257; 60529; 130497 | ALX3; OSR1; ALX4 | 41 |
| 0.000374 | 0.005718 | GO:0001101 | b | 3 | response to acid chemical | 440; 1036; 1277; 2042; 5105; 7480; 130497 | PCK1; WNT10B; EPHA3; ASNS; CDO1; COL1A1; OSR1 | 338 |
| 0.000544 | 0.005775 | GO:0044851 | b | 2 | hair cycle phase | 1001; 7480 | CDH3; WNT10B | 9 |
| 0.000415 | 0.0059 | GO:0048598 | b | 3 | embryonic morphogenesis | 257; 655; 1303; 1749; 2335; 60529; 6913; 115908; 130497 | BMP7; FN1; ALX3; OSR1; ALX4; TBX15; DLX5; CTHRC1; COL12A1 | 573 |
| 0.000675 | 0.006372 | GO:0009653 | b | 2 | anatomical structure morphogenesis | 257; 655; 1001; 1277; 1303; 1749; 2042; 2335; 60529; 2828; 6913; 7135; 7480; 7490; 5740; 10631; 144347; 115908; 54829; 55079; 130497 | BMP7; WT1; FEZF2; FN1; GPR4; ALX3; WNT10B; TNNI1; EPHA3; OSR1; PTGIS; ALX4; POSTN; CTHRC1; TBX15; COL1A1; DLX5; CDH3; ASPN; RFLNA; COL12A1 | 2554 |
| 0.000565 | 0.007608 | GO:0072210 | b | 4 | metanephric nephron development | 7490; 7827; 130497 | WT1; NPHS2; OSR1 | 41 |
| 0.000953 | 0.008101 | GO:0007155 | b | 2 | cell adhesion | 655; 1001; 1277; 1303; 1805; 2042; 2335; 4958; 7409; 8639; 10398; 10631; 342035; 4026 | BMP7; AOC3; DPT; FN1; OMD; VAV1; GLDN; EPHA3; LPP; POSTN; COL1A1; CDH3; COL12A1; MYL9 | 1387 |
| 0.000639 | 0.008107 | GO:0060485 | b | 4 | mesenchyme development | 655; 1277; 2042; 2335; 7490; 130497 | BMP7; WT1; FN1; EPHA3; OSR1; COL1A1 | 266 |
| 0.000676 | 0.008107 | GO:0060173 | b | 4 | limb development | 257; 655; 1749; 60529; 130497 | ALX3; BMP7; ALX4; OSR1; DLX5 | 177 |
| 0.000678 | 0.008107 | GO:0060346 | b | 4 | bone trabecula formation | 1277; 7480 | WNT10B; COL1A1 | 10 |
| 0.000676 | 0.008429 | GO:0048736 | b | 3 | appendage development | 257; 655; 1749; 60529; 130497 | ALX3; BMP7; ALX4; OSR1; DLX5 | 177 |
| 0.000678 | 0.008429 | GO:0072110 | b | 3 | glomerular mesangial cell proliferation | 655; 7490 | BMP7; WT1 | 10 |
| 0.000767 | 0.008848 | GO:0030855 | b | 4 | epithelial cell differentiation | 655; 1001; 1749; 2828; 5105; 7480; 7490; 7827; 56033; 130497 | BMP7; WT1; PCK1; GPR4; WNT10B; OSR1; BARX1; NPHS2; DLX5; CDH3 | 758 |
| 0.000975 | 0.009065 | GO:0045667 | b | 5 | regulation of osteoblast differentiation | 655; 7480; 338773; 115908 | BMP7; TMEM119; WNT10B; CTHRC1 | 111 |
| 0.000989 | 0.009065 | GO:0042340 | b | 5 | keratan sulfate catabolic process | 2331; 4958 | FMOD; OMD | 12 |
| 0.000812 | 0.009502 | GO:0010033 | b | 3 | response to organic substance | 440; 655; 1036; 1277; 1749; 2042; 2167; 2331; 2335; 3563; 3598; 5105; 5167; 6517; 7123; 7409; 7480; 7490; 5740; 10631; 284297; 85363; 54829; 130497 | PCK1; EPHA3; CDO1; FABP4; CLEC3B; WNT10B; FMOD; DLX5; WT1; SSC5D; OSR1; PTGIS; ENPP1; POSTN; IL3RA; BMP7; IL13RA2; FN1; VAV1; SLC2A4; ASNS; COL1A1; TRIM5; ASPN | 3166 |
| 0.000859 | 0.009502 | GO:0051239 | b | 3 | regulation of multicellular organismal process | 655; 1001; 1277; 2042; 2167; 2335; 2828; 4958; 5167; 7135; 7480; 7490; 5740; 10398; 10631; 338773; 144347; 115908; 284297; 54829; 55079; 26471; 130497 | EPHA3; TMEM119; FABP4; MYL9; FEZF2; WNT10B; TNNI1; CDH3; ASPN; WT1; SSC5D; RFLNA; NUPR1; OSR1; PTGIS; ENPP1; POSTN; BMP7; FN1; GPR4; OMD; COL1A1; CTHRC1 | 2983 |
| 0.001166 | 0.009863 | GO:1901722 | b | 5 | regulation of cell proliferation involved in kidney development | 655; 7490 | BMP7; WT1 | 13 |
| 0.001166 | 0.009863 | GO:2000194 | b | 5 | regulation of female gonad development | 7490; 26471 | WT1; NUPR1 | 13 |
| 0.000955 | 0.010005 | GO:0061383 | b | 3 | trabecula morphogenesis | 655; 1277; 7480 | BMP7; WNT10B; COL1A1 | 49 |
| 0.001319 | 0.010507 | GO:0030111 | b | 5 | regulation of Wnt signaling pathway | 1001; 1277; 1749; 7480; 56033; 115908 | WNT10B; BARX1; COL1A1; DLX5; CDH3; CTHRC1 | 306 |
| 0.001337 | 0.010507 | GO:0090183 | b | 5 | regulation of kidney development | 655; 7490; 130497 | BMP7; WT1; OSR1 | 55 |
| 0.001001 | 0.010843 | GO:0072001 | b | 4 | renal system development | 655; 2018; 2828; 7490; 7827; 130497 | BMP7; WT1; GPR4; EMX2; OSR1; NPHS2 | 290 |
| 0.001008 | 0.010843 | GO:0043200 | b | 4 | response to amino acid | 440; 1036; 1277; 5105 | PCK1; COL1A1; ASNS; CDO1 | 112 |

| | | | | | | | | |
|---|---|---|---|---|---|---|---|---|
| 0.001041 | 0.010843 | GO:0051093 | b | 4 | negative regulation of developmental process | 655; 1001; 2828; 5167; 7480; 7490; 10631; 144347; 54829; 55079; 130497 | BMP7; WT1; FEZF2; GPR4; WNT10B; OSR1; ENPP1; POSTN; CDH3; ASPN; RFLNA | 933 |
| 0.001166 | 0.011409 | GO:0061430 | b | 4 | bone trabecula morphogenesis | 1277; 7480 | WNT10B; COL1A1 | 13 |
| 0.001166 | 0.011409 | GO:0038065 | b | 4 | collagen-activated signaling pathway | 1277; 126014 | OSCAR; COL1A1 | 13 |
| 0.001304 | 0.011761 | GO:0071229 | b | 4 | cellular response to acid chemical | 1277; 2042; 5105; 7480; 130497 | WNT10B; PCK1; EPHA3; OSR1; COL1A1 | 205 |
| 0.001342 | 0.011761 | GO:0002062 | b | 4 | chondrocyte differentiation | 1303; 7480; 144347; 130497 | WNT10B; RFLNA; COL12A1; OSR1 | 121 |
| 0.00139 | 0.011761 | GO:0048762 | b | 4 | mesenchymal cell differentiation | 655; 1277; 2042; 2335; 130497 | BMP7; EPHA3; OSR1; FN1; COL1A1 | 208 |
| 0.001394 | 0.011761 | GO:0042060 | b | 4 | wound healing | 1001; 1277; 2335; 2828; 7409; 7480; 10398; 10631 | FN1; GPR4; VAV1; WNT10B; POSTN; COL1A1; CDH3; MYL9 | 547 |
| 0.001413 | 0.011761 | GO:0048568 | b | 4 | embryonic organ development | 257; 655; 1749; 60529; 6913; 115908; 130497 | BMP7; ALX3; OSR1; ALX4; TBX15; DLX5; CTHRC1 | 424 |
| 0.00142 | 0.011761 | GO:0060348 | b | 4 | bone development | 1277; 1303; 1749; 338773; 144347 | TMEM119; DLX5; RFLNA; COL12A1; COL1A1 | 209 |
| 0.001483 | 0.011976 | GO:0044273 | b | 4 | sulfur compound catabolic process | 1036; 2331; 4958 | CDO1; FMOD; OMD | 57 |
| 0.001558 | 0.012277 | GO:0042476 | b | 4 | odontogenesis | 655; 1277; 54829; 130497 | BMP7; ASPN; OSR1; COL1A1 | 126 |
| 0.00178 | 0.013501 | GO:0072112 | b | 5 | glomerular visceral epithelial cell differentiation | 7490; 7827 | WT1; NPHS2 | 16 |
| 0.00178 | 0.013686 | GO:0061318 | b | 4 | renal filtration cell differentiation | 7490; 7827 | WT1; NPHS2 | 16 |
| 0.001846 | 0.01387 | GO:0001655 | b | 4 | urogenital system development | 655; 2018; 2828; 7490; 7827; 130497 | BMP7; WT1; GPR4; EMX2; OSR1; NPHS2 | 327 |
| 0.001986 | 0.01444 | GO:0051094 | b | 4 | positive regulation of developmental process | 655; 121340; 1277; 2042; 2335; 7480; 7490; 5740; 338773; 115908; 83719; 55079; 130497 | BMP7; WT1; YPEL3; FN1; WNT10B; EPHA3; OSR1; PTGIS; SP7; TMEM119; COL1A1; FEZF2; CTHRC1 | 1331 |
| 0.002012 | 0.01444 | GO:0048820 | b | 4 | hair follicle maturation | 1001; 7480 | CDH3; WNT10B | 17 |
| 0.001596 | 0.015876 | GO:0009790 | b | 3 | embryo development | 257; 655; 1277; 1303; 1749; 2335; 60529; 6913; 7490; 115908; 130497 | BMP7; WT1; FN1; ALX3; OSR1; ALX4; TBX15; COL1A1; DLX5; CTHRC1; COL12A1 | 984 |
| 0.000417 | 0.016254 | GO:0030020 | m | 3 | extracellular matrix structural constituent conferring tensile strength | 1277; 1298; 1303 | COL9A2; COL12A1; COL1A1 | 37 |
| 0.002231 | 0.016361 | GO:0072073 | b | 5 | kidney epithelium development | 655; 7490; 7827; 130497 | BMP7; WT1; NPHS2; OSR1 | 139 |
| 0.002371 | 0.016647 | GO:0071560 | b | 4 | cellular response to transforming growth factor beta stimulus | 1277; 2331; 7123; 10631; 54829 | CLEC3B; POSTN; ASPN; FMOD; COL1A1 | 235 |
| 0.00178 | 0.016864 | GO:0021542 | b | 3 | dentate gyrus development | 2018; 55079 | EMX2; FEZF2 | 16 |
| 0.002517 | 0.017295 | GO:1905939 | b | 4 | regulation of gonad development | 7490; 26471 | WT1; NUPR1 | 19 |
| 0.002517 | 0.017588 | GO:0051797 | b | 5 | regulation of hair follicle development | 1001; 7480 | CDH3; WNT10B | 19 |
| 0.002558 | 0.017588 | GO:0071345 | b | 5 | cellular response to cytokine stimulus | 1277; 2167; 2335; 3563; 3598; 5105; 6517; 7409; 5740; 10631; 85363 | IL13RA2; PCK1; FN1; VAV1; SLC2A4; PTGIS; POSTN; FABP4; IL3RA; COL1A1; TRIM5 | 1045 |
| 0.002789 | 0.018594 | GO:0060039 | b | 5 | pericardium development | 655; 7490 | BMP7; WT1 | 20 |
| 0.002898 | 0.01875 | GO:0060350 | b | 5 | endochondral bone morphogenesis | 1277; 1303; 1749 | DLX5; COL12A1; COL1A1 | 72 |
| 0.002868 | 0.019301 | GO:0071456 | b | 4 | cellular response to hypoxia | 655; 5105; 6517; 5740 | BMP7; SLC2A4; PCK1; PTGIS | 149 |
| 0.002932 | 0.019328 | GO:0060429 | b | 4 | epithelium development | 655; 1001; 1749; 60529; 2828; 5105; 7480; 7490; 7827; 56033; 115908; 130497 | BMP7; WT1; PCK1; GPR4; WNT10B; OSR1; ALX4; BARX1; NPHS2; DLX5; CDH3; CTHRC1 | 1225 |
| 0.002312 | 0.020295 | GO:0003002 | b | 3 | regionalization | 2018; 60529; 7490; 56033; 55079; 130497 | WT1; FEZF2; EMX2; OSR1; ALX4; BARX1 | 342 |
| 0.002346 | 0.020295 | GO:0070482 | b | 3 | response to oxygen levels | 655; 1277; 5105; 6517; 5740; 10631 | BMP7; PCK1; SLC2A4; PTGIS; POSTN; COL1A1 | 343 |
| 0.003374 | 0.020791 | GO:0033688 | b | 5 | regulation of osteoblast proliferation | 338773; 115908 | TMEM119; CTHRC1 | 22 |
| 0.003598 | 0.020791 | GO:0034612 | b | 5 | response to tumor necrosis factor | 1277; 2167; 5105; 6517; 10631 | POSTN; FABP4; PCK1; SLC2A4; COL1A1 | 259 |
| 0.00362 | 0.020791 | GO:0036294 | b | 5 | cellular response to decreased oxygen levels | 655; 5105; 6517; 5740 | BMP7; SLC2A4; PCK1; PTGIS | 159 |
| 0.003686 | 0.020791 | GO:0034505 | b | 5 | tooth mineralization | 1277; 54829 | ASPN; COL1A1 | 23 |
| 0.003686 | 0.020791 | GO:1903010 | b | 5 | regulation of bone development | 338773; 144347 | TMEM119; RFLNA | 23 |

| | | | | | | | | |
|---|---|---|---|---|---|---|---|---|
| 0.002529 | 0.020966 | GO:0071495 | b | 3 | cellular response to endogenous stimulus | 440; 655; 1277; 1749; 2331; 5105; 5167; 6517; 7123; 7480; 7490; 10631; 54829 | BMP7; WT1; PCK1; CLEC3B; WNT10B; SLC2A4; ASNS; ENPP1; POSTN; FMOD; COL1A1; DLX5; ASPN | 1368 |
| 0.002691 | 0.021421 | GO:0071559 | b | 3 | response to transforming growth factor beta | 1277; 2331; 7123; 10631; 54829 | CLEC3B; POSTN; ASPN; FMOD; COL1A1 | 242 |
| 0.003374 | 0.021794 | GO:0010447 | b | 4 | response to acidic pH | 2828; 5105 | PCK1; GPR4 | 22 |
| 0.00401 | 0.022057 | GO:0070168 | b | 5 | negative regulation of biomineral tissue development | 144347; 54829 | RFLNA; ASPN | 24 |
| 0.004183 | 0.022447 | GO:0010717 | b | 5 | regulation of epithelial to mesenchymal transition | 655; 1277; 2042 | BMP7; EPHA3; COL1A1 | 82 |
| 0.003686 | 0.023342 | GO:0007413 | b | 4 | axonal fasciculation | 2042; 55079 | EPHA3; FEZF2 | 23 |
| 0.003935 | 0.024441 | GO:0060537 | b | 4 | muscle tissue development | 655; 7135; 7480; 7490; 26471; 130497 | BMP7; WT1; WNT10B; NUPR1; OSR1; TNNI1 | 381 |
| 0.00401 | 0.024441 | GO:0021871 | b | 4 | forebrain regionalization | 2018; 55079 | EMX2; FEZF2 | 24 |
| 0.003181 | 0.024583 | GO:0048646 | b | 2 | anatomical structure formation involved in morphogenesis | 655; 1277; 1303; 1749; 2335; 2828; 7480; 7490; 5740; 115908; 130497 | BMP7; WT1; FN1; GPR4; WNT10B; OSR1; PTGIS; COL1A1; DLX5; CTHRC1; COL12A1 | 1075 |
| 0.004698 | 0.024611 | GO:0072202 | b | 5 | cell differentiation involved in metanephros development | 7827; 130497 | NPHS2; OSR1 | 26 |
| 0.004488 | 0.024675 | GO:0043565 | m | 5 | sequence-specific DNA binding | 257; 121340; 1749; 2018; 60529; 165; 6913; 7490; 56033; 55079; 130497 | WT1; FEZF2; ALX3; EMX2; OSR1; ALX4; BARX1; AEBP1; TBX15; DLX5; SP7 | 1125 |
| 0.006271 | 0.024675 | GO:0005509 | m | 5 | calcium ion binding | 1001; 5167; 5321; 7123; 8639; 10398; 54829; 60681 | AOC3; FKBP10; CLEC3B; ASPN; ENPP1; CDH3; PLA2G4A; MYL9 | 700 |
| 0.009826 | 0.024675 | GO:0044212 | m | 5 | transcription regulatory region DNA binding | 121340; 1749; 60529; 165; 6913; 7490; 56033; 55079; 130497 | WT1; FEZF2; OSR1; ALX4; BARX1; AEBP1; TBX15; DLX5; SP7 | 916 |
| 0.001363 | 0.02544 | GO:0005788 | c | 4 | endoplasmic reticulum lumen | 1277; 1298; 1303; 2335; 493869; 60681 | FN1; FKBP10; COL9A2; COL1A1; GPX8; COL12A1 | 308 |
| 0.002361 | 0.02544 | GO:0032432 | c | 4 | actin filament bundle | 4082; 10398; 144347 | RFLNA; MARCKS; MYL9 | 67 |
| 0.003547 | 0.02544 | GO:0070062 | c | 4 | extracellular exosome | 1303; 1675; 2167; 2335; 4082; 4958; 165; 5105; 5349; 5375; 5480; 6517; 7123; 7827; 8857; 23452; 126014 | PCK1; FXYD3; FN1; OMD; PMP2; OSCAR; CFD; SLC2A4; FCGBP; CLEC3B; FABP4; MARCKS; AEBP1; ANGPTL2; NPHS2; PPIC; COL12A1 | 2139 |
| 0.003634 | 0.02544 | GO:0005811 | c | 4 | lipid droplet | 2167; 5321; 63924 | FABP4; PLA2G4A; CIDEC | 78 |
| 0.00526 | 0.026912 | GO:0098773 | b | 5 | skin epidermis development | 1001; 60529; 7480 | CDH3; WNT10B; ALX4 | 89 |
| 0.005437 | 0.027184 | GO:0018146 | b | 5 | keratan sulfate biosynthetic process | 2331; 4958 | FMOD; OMD | 28 |
| 0.003634 | 0.027286 | GO:0072028 | b | 3 | nephron morphogenesis | 655; 7490; 130497 | BMP7; WT1; OSR1 | 78 |
| 0.003702 | 0.027286 | GO:0071695 | b | 3 | anatomical structure maturation | 1001; 7480; 144347; 342035 | CDH3; RFLNA; WNT10B; GLDN | 160 |
| 0.004566 | 0.02731 | GO:0001822 | b | 4 | kidney development | 655; 2828; 7490; 7827; 130497 | BMP7; WT1; NPHS2; OSR1; GPR4 | 274 |
| 0.004702 | 0.027611 | GO:0034097 | b | 4 | response to cytokine | 1277; 2167; 2335; 3563; 3598; 5105; 6517; 7409; 5740; 10631; 85363 | IL13RA2; PCK1; FN1; VAV1; SLC2A4; PTGIS; POSTN; FABP4; IL3RA; COL1A1; TRIM5 | 1132 |
| 0.004937 | 0.02793 | GO:0001942 | b | 4 | hair follicle development | 1001; 60529; 7480 | CDH3; WNT10B; ALX4 | 87 |
| 0.005061 | 0.02793 | GO:0042634 | b | 4 | regulation of hair cycle | 1001; 7480 | CDH3; WNT10B | 27 |
| 0.005061 | 0.02793 | GO:0001958 | b | 4 | endochondral ossification | 1277; 1749 | DLX5; COL1A1 | 27 |
| 0.005188 | 0.02793 | GO:0007369 | b | 4 | gastrulation | 655; 1303; 2335; 130497 | BMP7; FN1; COL12A1; OSR1 | 176 |
| 0.005188 | 0.02793 | GO:0071453 | b | 4 | cellular response to oxygen levels | 655; 5105; 6517; 5740 | BMP7; SLC2A4; PCK1; PTGIS | 176 |
| 0.005437 | 0.028789 | GO:0034698 | b | 4 | response to gonadotropin | 440; 7490 | WT1; ASNS | 28 |
| 0.004108 | 0.029095 | GO:0009719 | b | 2 | response to endogenous stimulus | 440; 655; 1036; 1277; 1749; 2331; 5105; 5167; 6517; 7123; 7480; 7490; 10631; 54829 | BMP7; WT1; PCK1; CLEC3B; WNT10B; SLC2A4; ASNS; CDO1; POSTN; FMOD; COL1A1; DLX5; ENPP1; ASPN | 1622 |
| 0.004097 | 0.029121 | GO:0009636 | b | 3 | response to toxic substance | 440; 655; 1036; 1277; 493869; 6517; 26471 | BMP7; NUPR1; SLC2A4; ASNS; CDO1; COL1A1; GPX8 | 513 |
| 0.005595 | 0.02915 | GO:0033273 | b | 4 | response to vitamin | 655; 1277; 10631 | BMP7; POSTN; COL1A1 | 91 |
| 0.00443 | 0.030397 | GO:0009611 | b | 3 | response to wounding | 1001; 1277; 2335; 2828; 7409; 7480; 10398; 10631 | FN1; GPR4; VAV1; WNT10B; POSTN; COL1A1; CDH3; MYL9 | 660 |
| 0.005947 | 0.030488 | GO:0022603 | b | 4 | regulation of anatomical structure morphogenesis | 655; 2335; 2828; 7490; 5740; 10631; 115908; 54829; 55079; 130497 | BMP7; WT1; FEZF2; FN1; GPR4; ASPN; OSR1; PTGIS; POSTN; CTHRC1 | 1004 |

| | | | | | | | | |
|---|---|---|---|---|---|---|---|---|
| 0.006042 | 0.030493 | GO:0051241 | b | 4 | negative regulation of multicellular organismal process | 655; 1001; 2335; 2828; 7480; 7490; 144347; 284297; 54829; 55079; 130497 | BMP7; WT1; FEZF2; FN1; GPR4; WNT10B; SSC5D; OSR1; CDH3; ASPN; RFLNA | 1171 |
| 0.004698 | 0.03072 | GO:0033687 | b | 2 | osteoblast proliferation | 338773; 115908 | TMEM119; CTHRC1 | 26 |
| 0.004698 | 0.030786 | GO:0060343 | b | 3 | trabecula formation | 1277; 7480 | WNT10B; COL1A1 | 26 |
| 0.005061 | 0.030786 | GO:0036075 | b | 3 | replacement ossification | 1277; 1749 | DLX5; COL1A1 | 27 |
| 0.005175 | 0.030786 | GO:0009725 | b | 3 | response to hormone | 440; 655; 1036; 1277; 5105; 5167; 6517; 7480; 7490; 10631 | BMP7; WT1; PCK1; WNT10B; SLC2A4; ASNS; ENPP1; POSTN; COL1A1; CDO1 | 984 |
| 0.00526 | 0.030786 | GO:0022405 | b | 3 | hair cycle process | 1001; 60529; 7480 | CDH3; WNT10B; ALX4 | 89 |
| 0.00526 | 0.030786 | GO:0060021 | b | 3 | roof of mouth development | 1749; 60529; 130497 | DLX5; OSR1; ALX4 | 89 |
| 0.006304 | 0.031326 | GO:0060993 | b | 4 | kidney morphogenesis | 655; 7490; 130497 | BMP7; WT1; OSR1 | 95 |
| 0.006489 | 0.031724 | GO:0001657 | b | 5 | ureteric bud development | 655; 7490; 130497 | BMP7; WT1; OSR1 | 96 |
| 0.0065 | 0.031813 | GO:0043434 | b | 4 | response to peptide hormone | 655; 1036; 1277; 5105; 5167; 6517 | BMP7; PCK1; SLC2A4; CDO1; ENPP1; COL1A1 | 423 |
| 0.00526 | 0.031936 | GO:0022404 | b | 2 | molting cycle process | 1001; 60529; 7480 | CDH3; WNT10B; ALX4 | 89 |
| 0.006677 | 0.032191 | GO:0072164 | b | 4 | mesonephric tubule development | 655; 7490; 130497 | BMP7; WT1; OSR1 | 97 |
| 0.00398 | 0.033828 | GO:0043230 | c | 2 | extracellular organelle | 1303; 1675; 2167; 2335; 4082; 4958; 165; 5105; 5349; 5375; 5480; 6517; 7123; 7827; 8857; 23452; 126014 | PCK1; FXYD3; FN1; OMD; PMP2; OSCAR; CFD; SLC2A4; FCGBP; CLEC3B; FABP4; MARCKS; AEBP1; ANGPTL2; NPHS2; PPIC; COL12A1 | 2163 |
| 0.007498 | 0.035617 | GO:0042339 | b | 4 | keratan sulfate metabolic process | 2331; 4958 | FMOD; OMD | 33 |
| 0.007463 | 0.035693 | GO:0001823 | b | 5 | mesonephros development | 655; 7490; 130497 | BMP7; WT1; OSR1 | 101 |
| 0.007856 | 0.036774 | GO:0045595 | b | 4 | regulation of cell differentiation | 655; 121340; 1277; 2042; 2335; 5167; 7480; 10398; 10631; 338773; 144347; 115908; 55079; 130497 | BMP7; FEZF2; FN1; TMEM119; WNT10B; EPHA3; OSR1; ENPP1; POSTN; CTHRC1; COL1A1; SP7; RFLNA; MYL9 | 1747 |
| 0.003942 | 0.036793 | GO:1903561 | c | 3 | extracellular vesicle | 1303; 1675; 2167; 2335; 4082; 4958; 165; 5105; 5349; 5375; 5480; 6517; 7123; 7827; 8857; 23452; 126014 | PCK1; FXYD3; FN1; OMD; PMP2; OSCAR; CFD; SLC2A4; FCGBP; CLEC3B; FABP4; MARCKS; AEBP1; ANGPTL2; NPHS2; PPIC; COL12A1 | 2161 |
| 0.002837 | 0.038306 | GO:1901681 | m | 2 | sulfur compound binding | 655; 2335; 5167; 7123; 10631 | BMP7; CLEC3B; POSTN; FN1; ENPP1 | 245 |
| 0.002361 | 0.038564 | GO:0005518 | m | 3 | collagen binding | 2335; 165; 54829 | ASPN; AEBP1; FN1 | 67 |
| 0.003374 | 0.038564 | GO:0030021 | m | 3 | extracellular matrix structural constituent conferring compression resistance | 2331; 54829 | ASPN; FMOD | 22 |
| 0.003955 | 0.038564 | GO:0008201 | m | 3 | heparin binding | 655; 2335; 7123; 10631 | BMP7; CLEC3B; FN1; POSTN | 163 |
| 0.008566 | 0.039526 | GO:0036293 | b | 4 | response to decreased oxygen levels | 655; 5105; 6517; 5740; 10631 | BMP7; POSTN; SLC2A4; PCK1; PTGIS | 319 |
| 0.008959 | 0.039882 | GO:0048863 | b | 4 | stem cell differentiation | 655; 121340; 2335; 130497 | BMP7; OSR1; FN1; SP7 | 206 |
| 0.008959 | 0.039882 | GO:0010810 | b | 4 | regulation of cell-substrate adhesion | 1277; 2042; 2335; 10631 | POSTN; EPHA3; FN1; COL1A1 | 206 |
| 0.009014 | 0.039882 | GO:0042127 | b | 4 | regulation of cell proliferation | 655; 1001; 1749; 1805; 2335; 5321; 7480; 7490; 338773; 115908; 55079; 26471; 130497 | BMP7; WT1; DPT; FEZF2; FN1; WNT10B; NUPR1; OSR1; CTHRC1; TMEM119; DLX5; CDH3; PLA2G4A | 1592 |
| 0.007062 | 0.040151 | GO:0060325 | b | 3 | face morphogenesis | 1277; 1749 | DLX5; COL1A1 | 32 |
| 0.009321 | 0.040687 | GO:0043436 | b | 4 | oxoacid metabolic process | 440; 1036; 2331; 4958; 5105; 5167; 5321; 5740; 10941; 641371 | PCK1; OMD; ASNS; PTGIS; ENPP1; UGT2A1; FMOD; PLA2G4A; CDO1; ACOT1 | 1073 |
| 0.007422 | 0.041029 | GO:0001666 | b | 3 | response to hypoxia | 655; 5105; 6517; 5740; 10631 | BMP7; POSTN; SLC2A4; PCK1; PTGIS | 308 |
| 0.008879 | 0.041195 | GO:0033762 | b | 5 | response to glucagon | 1036; 5105 | PCK1; CDO1 | 36 |
| 0.009193 | 0.041195 | GO:0001676 | b | 5 | long-chain fatty acid metabolic process | 5321; 5740; 641371 | PLA2G4A; PTGIS; ACOT1 | 109 |
| 0.00926 | 0.041195 | GO:2000027 | b | 5 | regulation of animal organ morphogenesis | 655; 7490; 115908; 54829 | BMP7; WT1; ASPN; CTHRC1 | 208 |
| 0.009362 | 0.041195 | GO:0070741 | b | 5 | response to interleukin-6 | 5105; 5740 | PCK1; PTGIS | 37 |
| 0.009823 | 0.041473 | GO:0043009 | b | 5 | chordate embryonic development | 257; 655; 1277; 60529; 6913; 115908; 130497 | BMP7; ALX3; OSR1; ALX4; TBX15; COL1A1; CTHRC1 | 605 |
| 0.009914 | 0.041473 | GO:0016055 | b | 5 | Wnt signaling pathway | 1001; 1277; 1749; 7480; 56033; 115908 | WNT10B; BARX1; COL1A1; DLX5; CDH3; CTHRC1 | 463 |
| 0.009991 | 0.041473 | GO:0071396 | b | 5 | cellular response to lipid | 655; 1277; 2042; 5105; 7480; 85363; 130497 | BMP7; PCK1; WNT10B; EPHA3; OSR1; COL1A1; TRIM5 | 607 |

| | | | | | | | | |
|---|---|---|---|---|---|---|---|---|
| 0.009897 | 0.042121 | GO:0060349 | b | 4 | bone morphogenesis | 1277; 1303; 1749 | DLX5; COL12A1; COL1A1 | 112 |
| 0.009959 | 0.042121 | GO:0033993 | b | 4 | response to lipid | 655; 1036; 1277; 2042; 5105; 7480; 10631; 85363; 130497 | BMP7; PCK1; WNT10B; EPHA3; CDO1; OSR1; POSTN; COL1A1; TRIM5 | 918 |
| 0.008137 | 0.043763 | GO:0046677 | b | 3 | response to antibiotic | 1036; 1277; 5321; 6517; 8639 | PLA2G4A; SLC2A4; CDO1; AOC3; COL1A1 | 315 |
| 0.008789 | 0.046028 | GO:1901700 | b | 3 | response to oxygen-containing compound | 440; 655; 1036; 1277; 2042; 5105; 5167; 6517; 7480; 7490; 10631; 85363; 130497 | BMP7; WT1; PCK1; WNT10B; SLC2A4; EPHA3; ASNS; CDO1; POSTN; COL1A1; OSR1; ENPP1; TRIM5 | 1587 |
| 0.009857 | 0.048086 | GO:0060323 | b | 3 | head morphogenesis | 1277; 1749 | DLX5; COL1A1 | 38 |

Table 5-8. Gene SNPs in responder and nonresponder mice.

| | | | | No. of R mice (genotype) | | | No. of N mice (genotype) | | | | | | | | | |
|---|---|---|---|---|---|---|---|---|---|---|---|---|---|---|---|---|
| Chromosome | Position | Ref | Alt | Ref/Ref | Ref/Alt | Alt/Alt | Ref/Ref | Ref/Alt | Alt/Alt | Pvalue | Bonferroni | Annotation | Impact | Gene_Name | Variant_transcript | Variant_protein |
| 1 | 171234963 | A | G | 0 | 6 | 0 | 0 | 0 | 6 | NA | NA | 3_prime_UTR_variant | MODIFIER | Ndufs2 | c.*87T>C | |
| 1 | 171237185 | C | T | 0 | 6 | 0 | 6 | 0 | 0 | NA | NA | synonymous_variant | LOW | Ndufs2 | c.1071G>A | p.Lys357Lys |
| 1 | 171238004 | G | A | 0 | 6 | 0 | 6 | 0 | 0 | NA | NA | synonymous_variant | LOW | Ndufs2 | c.927C>T | p.Asp309Asp |
| 1 | 171238331 | G | A | 0 | 6 | 0 | 6 | 0 | 0 | NA | NA | synonymous_variant | LOW | Ndufs2 | c.822C>T | p.Asp274Asp |
| 1 | 171280157 | A | G | 0 | 6 | 0 | 0 | 0 | 6 | NA | NA | synonymous_variant | LOW | Ppox | c.321T>C | p.Pro107Pro |
| 1 | 171280454 | C | T | 0 | 6 | 0 | 6 | 0 | 0 | NA | NA | synonymous_variant | LOW | Ppox | c.156G>A | p.Ala52Ala |
| 1 | 171285578 | G | A | 0 | 6 | 0 | 6 | 0 | 0 | NA | NA | synonymous_variant | LOW | Usp21 | c.624C>T | p.Gly208Gly |
| 1 | 171294700 | G | A | 0 | 6 | 0 | 6 | 0 | 0 | NA | NA | synonymous_variant | LOW | Ufc1 | c.57C>T | p.Asn19Asn |
| 1 | 171387531 | C | T | 0 | 6 | 0 | 6 | 0 | 0 | NA | NA | 3_prime_UTR_variant | MODIFIER | Nectin4 | c.*726C>T | |
| 1 | 171387799 | T | C | 0 | 6 | 0 | 6 | 0 | 0 | NA | NA | 3_prime_UTR_variant | MODIFIER | Nectin4 | c.*994T>C | |
| 1 | 171416853 | C | T | 0 | 6 | 0 | 6 | 0 | 0 | NA | NA | missense_variant | MODERATE | Usf1 | c.320C>T | p.Thr107Met |
| 1 | 171463512 | A | G | 0 | 6 | 0 | 6 | 0 | 0 | NA | NA | 3_prime_UTR_variant | MODIFIER | F11r | c.*418A>G | |
| 1 | 171463558 | C | T | 0 | 6 | 0 | 6 | 0 | 0 | NA | NA | 3_prime_UTR_variant | MODIFIER | F11r | c.*464C>T | |
| 1 | 171504107 | A | G | 0 | 6 | 0 | 0 | 0 | 6 | NA | NA | missense_variant | MODERATE | Alyref2 | c.451A>G | p.Thr151Ala |
| 1 | 171504171 | G | A | 0 | 6 | 0 | 6 | 0 | 0 | NA | NA | missense_variant | MODERATE | Alyref2 | c.515G>A | p.Ser172Asn |
| 1 | 171504187 | C | T | 0 | 6 | 0 | 0 | 0 | 6 | NA | NA | synonymous_variant | LOW | Alyref2 | c.531C>T | p.Gly177Gly |
| 2 | 32095631 | A | G | 6 | 0 | 0 | 0 | 6 | 0 | NA | NA | upstream_gene_variant | MODIFIER | Plpp7 | c.-181A>G | |
| 2 | 32109665 | G | A | 6 | 0 | 0 | 0 | 6 | 0 | NA | NA | missense_variant | MODERATE | Plpp7 | c.508G>A | p.Gly170Ser |
| 2 | 32110154 | A | G | 6 | 0 | 0 | 0 | 6 | 0 | NA | NA | 3_prime_UTR_variant | MODIFIER | Plpp7 | c.*181A>G | |
| 2 | 32233700 | C | T | 0 | 0 | 6 | 0 | 6 | 0 | NA | NA | 3_prime_UTR_variant | MODIFIER | Prrc2b | c.*2958C>T | |
| 2 | 32402585 | C | A | 0 | 0 | 6 | 0 | 6 | 0 | NA | NA | 3_prime_UTR_variant | MODIFIER | Ptges2 | c.*257C>A | |
| 2 | 32607091 | G | C | 0 | 0 | 6 | 0 | 6 | 0 | NA | NA | 5_prime_UTR_variant | MODIFIER | St6galnac6 | c.-50G>C | |
| 2 | 32614942 | C | T | 0 | 0 | 6 | 0 | 6 | 0 | NA | NA | synonymous_variant | LOW | St6galnac6 | c.459C>T | p.Phe153Phe |
| 2 | 32619572 | T | A | 0 | 0 | 6 | 0 | 6 | 0 | NA | NA | synonymous_variant | LOW | St6galnac6 | c.1041T>A | p.Ala347Ala |
| 2 | 32619599 | T | C | 0 | 0 | 6 | 0 | 6 | 0 | NA | NA | synonymous_variant | LOW | St6galnac6 | c.1068T>C | p.His356His |
| 2 | 32619983 | G | A | 0 | 0 | 6 | 0 | 6 | 0 | NA | NA | 3_prime_UTR_variant | MODIFIER | St6galnac6 | c.*243G>A | |
| 2 | 32620397 | C | T | 0 | 0 | 6 | 0 | 6 | 0 | NA | NA | 3_prime_UTR_variant | MODIFIER | St6galnac6 | c.*657C>T | |
| 2 | 32620721 | A | G | 0 | 0 | 6 | 0 | 6 | 0 | NA | NA | 3_prime_UTR_variant | MODIFIER | St6galnac6 | c.*981A>G | |
| 2 | 32646834 | G | T | 6 | 0 | 0 | 0 | 6 | 0 | NA | NA | 5_prime_UTR_variant | MODIFIER | Eng | c.-124G>T | |
| 2 | 32683377 | G | A | 0 | 6 | 0 | 6 | 0 | 0 | NA | NA | synonymous_variant | LOW | Fpgs | c.1446C>T | p.Ala482Ala |
| 2 | 91712576 | C | T | 0 | 6 | 0 | 6 | 0 | 0 | NA | NA | synonymous_variant | LOW | Harbi1 | c.381C>T | p.Ala127Ala |
| 2 | 91767089 | C | A | 0 | 6 | 0 | 6 | 0 | 0 | NA | NA | synonymous_variant | LOW | Ambra1 | c.168C>A | p.Thr56Thr |
| 2 | 91940998 | G | A | 0 | 6 | 0 | 6 | 0 | 0 | NA | NA | synonymous_variant | LOW | Dgkz | c.1191C>T | p.Ile397Ile |
| 3 | 10015080 | T | C | 0 | 6 | 0 | 6 | 0 | 0 | NA | NA | synonymous_variant | LOW | Fabp5 | c.189T>C | p.Thr63Thr |
| 3 | 10180239 | T | C | 0 | 6 | 0 | 6 | 0 | 0 | NA | NA | 3_prime_UTR_variant | MODIFIER | Pmp2 | c.*511A>G | |
| 4 | 98756431 | A | C | 6 | 0 | 0 | 0 | 6 | 0 | NA | NA | 3_prime_UTR_variant | MODIFIER | Kank4 | c.*32T>G | |
| 4 | 98778499 | C | T | 6 | 0 | 0 | 0 | 6 | 0 | NA | NA | synonymous_variant | LOW | Kank4 | c.1710G>A | p.Val570Val |
| 4 | 99079749 | G | A | 6 | 0 | 0 | 0 | 6 | 0 | NA | NA | missense_variant | MODERATE | Dock7 | c.490C>T | p.Pro164Ser |
| 4 | 155900130 | G | A | 0 | 6 | 0 | 6 | 0 | 0 | NA | NA | synonymous_variant | LOW | Acap3 | c.609G>A | p.Gln203Gln |
| 7 | 116124812 | T | C | 0 | 0 | 6 | 0 | 6 | 0 | NA | NA | missense_variant | MODERATE | Plekha7 | c.3787A>G | p.Ile1263Val |
| 8 | 88134330 | A | T | 0 | 0 | 6 | 0 | 6 | 0 | NA | NA | 3_prime_UTR_variant | MODIFIER | Cnep1r1 | c.*501A>T | |
| 8 | 88139920 | G | A | 0 | 0 | 6 | 0 | 6 | 0 | NA | NA | missense_variant | MODERATE | Heatr3 | c.340G>A | p.Val114Ile |
| 8 | 88141697 | A | C | 0 | 0 | 6 | 0 | 6 | 0 | NA | NA | missense_variant | MODERATE | Heatr3 | c.425A>C | p.Glu142Ala |
| 8 | 88171037 | C | T | 0 | 0 | 6 | 0 | 6 | 0 | NA | NA | 3_prime_UTR_variant | MODIFIER | Heatr3 | c.*56C>T | |
| 9 | 105412506 | A | C | 0 | 6 | 0 | 6 | 0 | 0 | NA | NA | 3_prime_UTR_variant | MODIFIER | Atp2c1 | c.*553T>G | |
| 11 | 33117567 | G | T | 0 | 6 | 0 | 6 | 0 | 0 | NA | NA | 3_prime_UTR_variant | MODIFIER | Fgf18 | c.*198C>A | |
| 11 | 33117637 | T | C | 0 | 6 | 0 | 0 | 0 | 6 | NA | NA | 3_prime_UTR_variant | MODIFIER | Fgf18 | c.*128A>G | |
| 11 | 33201203 | T | C | 0 | 6 | 0 | 0 | 0 | 6 | NA | NA | 3_prime_UTR_variant | MODIFIER | Tlx3 | c.*76A>G | |
| 11 | 33202605 | T | C | 0 | 6 | 0 | 0 | 0 | 6 | NA | NA | synonymous_variant | LOW | Tlx3 | c.489A>G | p.Pro163Pro |
| 11 | 102841413 | T | C | 0 | 0 | 6 | 0 | 6 | 0 | NA | NA | synonymous_variant | LOW | Eftud2 | c.2247A>G | p.Thr749Thr |
| 11 | 102841747 | G | A | 0 | 0 | 6 | 0 | 6 | 0 | NA | NA | synonymous_variant | LOW | Eftud2 | c.2067C>T | p.Ala689Ala |
| 11 | 102994138 | G | A | 0 | 0 | 6 | 0 | 6 | 0 | NA | NA | 3_prime_UTR_variant | MODIFIER | Dcakd | c.*743C>T | |
| 11 | 102994168 | G | A | 0 | 0 | 6 | 0 | 6 | 0 | NA | NA | 3_prime_UTR_variant | MODIFIER | Dcakd | c.*713C>T | |
| 11 | 102994363 | T | C | 0 | 0 | 6 | 0 | 6 | 0 | NA | NA | 3_prime_UTR_variant | MODIFIER | Dcakd | c.*518A>G | |
| 11 | 102994662 | A | G | 0 | 0 | 6 | 0 | 6 | 0 | NA | NA | 3_prime_UTR_variant | MODIFIER | Dcakd | c.*219T>C | |
| 11 | 103000178 | T | C | 0 | 0 | 6 | 0 | 6 | 0 | NA | NA | synonymous_variant | LOW | Dcakd | c.84A>G | p.Val28Val |
| 11 | 103000196 | T | C | 0 | 0 | 6 | 0 | 6 | 0 | NA | NA | synonymous_variant | LOW | Dcakd | c.66A>G | p.Gln22Gln |
| 11 | 103060125 | A | C | 0 | 0 | 6 | 0 | 6 | 0 | NA | NA | synonymous_variant | LOW | Nmt1 | c.1041A>C | p.Pro347Pro |
| 11 | 103403893 | G | A | 0 | 6 | 0 | 6 | 0 | 0 | NA | NA | 5_prime_UTR_variant | MODIFIER | Plekhm1 | c.-24C>T | |
| 12 | 29535104 | G | A | 0 | 6 | 0 | 0 | 0 | 6 | NA | NA | 5_prime_UTR_variant | MODIFIER | Myt1l | c.-619G>A | |
| 12 | 29893377 | G | A | 0 | 6 | 0 | 6 | 0 | 0 | NA | NA | synonymous_variant | LOW | Myt1l | c.2793G>A | p.Pro931Pro |
| 12 | 29895177 | A | G | 0 | 6 | 0 | 0 | 0 | 6 | NA | NA | synonymous_variant | LOW | Myt1l | c.2946A>G | p.Ala982Ala |
| 12 | 29895264 | C | T | 0 | 6 | 0 | 6 | 0 | 0 | NA | NA | synonymous_variant | LOW | Myt1l | c.3033C>T | p.Pro1011Pro |
| 12 | 29920622 | C | T | 0 | 6 | 0 | 6 | 0 | 0 | NA | NA | 3_prime_UTR_variant | MODIFIER | Myt1l | c.*141C>T | |
| 12 | 29920914 | A | T | 0 | 6 | 0 | 6 | 0 | 0 | NA | NA | 3_prime_UTR_variant | MODIFIER | Myt1l | c.*433A>T | |
| 12 | 29922176 | G | A | 0 | 6 | 0 | 6 | 0 | 0 | NA | NA | 3_prime_UTR_variant | MODIFIER | Myt1l | c.*1695G>A | |
| 13 | 70637653 | G | A | 6 | 0 | 0 | 0 | 6 | 0 | NA | NA | upstream_gene_variant | MODIFIER | Ice1 | c.-56C>T | |
| 13 | 76579767 | G | C | 0 | 6 | 0 | 6 | 0 | 0 | NA | NA | intron_variant | MODIFIER | Mctp1 | c.-54-61945G>C | |
| 13 | 91022608 | T | G | 0 | 6 | 0 | 6 | 0 | 0 | NA | NA | synonymous_variant | LOW | Atg10 | c.400A>C | p.Arg134Arg |
| 13 | 93498461 | T | G | 0 | 6 | 0 | 6 | 0 | 0 | NA | NA | synonymous_variant | LOW | Jmy | c.846A>C | p.Val282Val |
| 13 | 93498667 | G | A | 0 | 6 | 0 | 6 | 0 | 0 | NA | NA | missense_variant | MODERATE | Jmy | c.640C>T | p.Pro214Ser |
| 13 | 93498749 | G | A | 0 | 6 | 0 | 6 | 0 | 0 | NA | NA | synonymous_variant | LOW | Jmy | c.558C>T | p.Val186Val |
| 13 | 93498831 | G | A | 0 | 6 | 0 | 6 | 0 | 0 | NA | NA | missense_variant | MODERATE | Jmy | c.476C>T | p.Ala159Val |
| 13 | 93498849 | G | A | 0 | 6 | 0 | 6 | 0 | 0 | NA | NA | missense_variant | MODERATE | Jmy | c.458C>T | p.Thr153Ile |
| 13 | 93499687 | A | T | 0 | 6 | 0 | 6 | 0 | 0 | NA | NA | 5_prime_UTR_variant | MODIFIER | Jmy | c.-381T>A | |
| 13 | 93862239 | A | T | 0 | 6 | 0 | 6 | 0 | 0 | NA | NA | missense_variant | MODERATE | Arsb | c.1054A>T | p.Thr352Ser |
| 14 | 8039847 | C | T | 0 | 6 | 0 | 0 | 0 | 6 | NA | NA | synonymous_variant | LOW | Abhd6 | c.219C>T | p.Ser73Ser |
| 14 | 8041211 | G | A | 0 | 6 | 0 | 0 | 0 | 6 | NA | NA | synonymous_variant | LOW | Abhd6 | c.369G>A | p.Gly123Gly |
| 14 | 8045478 | C | T | 0 | 6 | 0 | 0 | 0 | 6 | NA | NA | synonymous_variant | LOW | Abhd6 | c.534C>T | p.Tyr178Tyr |
| 14 | 8164250 | C | T | 0 | 6 | 0 | 0 | 0 | 6 | NA | NA | 3_prime_UTR_variant | MODIFIER | Pxk | c.*95C>T | |

| | | | | | | | | | | | | | | | | |
|---|---|---|---|---|---|---|---|---|---|---|---|---|---|---|---|---|
| 14 | 8164690 | T | C | 0 | 6 | 0 | 0 | 0 | 6 | NA | NA | 3_prime_UTR_variant | MODIFIER | Pxk | c.*535T>C | |
| 14 | 8166059 | C | T | 0 | 6 | 0 | 0 | 0 | 6 | NA | NA | 3_prime_UTR_variant | MODIFIER | Pdhb | c.*277G>A | |
| 14 | 8166076 | C | T | 0 | 6 | 0 | 0 | 0 | 6 | NA | NA | 3_prime_UTR_variant | MODIFIER | Pdhb | c.*260G>A | |
| 14 | 8166786 | A | G | 0 | 6 | 0 | 0 | 0 | 6 | NA | NA | synonymous_variant | LOW | Pdhb | c.861T>C | p.Asn287Asn |
| 14 | 93015454 | C | T | 0 | 6 | 0 | 6 | 0 | 0 | NA | NA | 3_prime_UTR_variant | MODIFIER | Pcdh9 | c.*58G>A | |
| 14 | 93886528 | G | T | 0 | 6 | 0 | 6 | 0 | 0 | NA | NA | synonymous_variant | LOW | Pcdh9 | c.2205C>A | p.Ile735Ile |
| 14 | 93886945 | C | T | 0 | 6 | 0 | 0 | 0 | 6 | NA | NA | synonymous_variant | LOW | Pcdh9 | c.1788G>A | p.Val596Val |
| 14 | 93888562 | C | T | 0 | 6 | 0 | 0 | 0 | 6 | NA | NA | synonymous_variant | LOW | Pcdh9 | c.171G>A | p.Gly57Gly |
| 14 | 93890344 | T | C | 0 | 6 | 0 | 6 | 0 | 0 | NA | NA | 5_prime_UTR_variant | MODIFIER | Pcdh9 | c.-380A>G | |
| 15 | 41819925 | G | A | 0 | 6 | 0 | 6 | 0 | 0 | NA | NA | missense_variant | MODERATE | Oxr1 | c.950G>A | p.Arg317His |
| 15 | 41820171 | G | A | 0 | 6 | 0 | 6 | 0 | 0 | NA | NA | missense_variant | MODERATE | Oxr1 | c.1196G>A | p.Gly399Glu |
| 15 | 42426872 | T | C | 0 | 6 | 0 | 6 | 0 | 0 | NA | NA | 3_prime_UTR_variant | MODIFIER | Angpt1 | c.*98A>G | |
| 15 | 42426962 | G | T | 0 | 6 | 0 | 6 | 0 | 0 | NA | NA | 3_prime_UTR_variant | MODIFIER | Angpt1 | c.*8C>A | |
| 15 | 96412655 | T | G | 6 | 0 | 0 | 0 | 6 | 0 | NA | NA | 3_prime_UTR_variant | MODIFIER | Scaf11 | c.*2133A>C | |
| 15 | 96572017 | C | T | 6 | 0 | 0 | 0 | 6 | 0 | NA | NA | 3_prime_UTR_variant | MODIFIER | Slc38a1 | c.*4694G>A | |
| 15 | 96572046 | C | T | 6 | 0 | 0 | 0 | 6 | 0 | NA | NA | 3_prime_UTR_variant | MODIFIER | Slc38a1 | c.*4665G>A | |
| 15 | 96572419 | T | G | 6 | 0 | 0 | 0 | 6 | 0 | NA | NA | 3_prime_UTR_variant | MODIFIER | Slc38a1 | c.*4292A>C | |
| 15 | 96572656 | A | C | 6 | 0 | 0 | 0 | 6 | 0 | NA | NA | 3_prime_UTR_variant | MODIFIER | Slc38a1 | c.*4055T>G | |
| 15 | 96572664 | G | A | 6 | 0 | 0 | 0 | 6 | 0 | NA | NA | 3_prime_UTR_variant | MODIFIER | Slc38a1 | c.*4047C>T | |
| 15 | 96572774 | G | T | 6 | 0 | 0 | 0 | 6 | 0 | NA | NA | 3_prime_UTR_variant | MODIFIER | Slc38a1 | c.*3937C>A | |
| 15 | 96572808 | A | G | 6 | 0 | 0 | 0 | 6 | 0 | NA | NA | 3_prime_UTR_variant | MODIFIER | Slc38a1 | c.*3903T>C | |
| 15 | 96572867 | T | G | 6 | 0 | 0 | 0 | 6 | 0 | NA | NA | 3_prime_UTR_variant | MODIFIER | Slc38a1 | c.*3844A>C | |
| 15 | 96572927 | C | T | 6 | 0 | 0 | 0 | 6 | 0 | NA | NA | 3_prime_UTR_variant | MODIFIER | Slc38a1 | c.*3784G>A | |
| 15 | 96572967 | C | T | 6 | 0 | 0 | 0 | 6 | 0 | NA | NA | 3_prime_UTR_variant | MODIFIER | Slc38a1 | c.*3744G>A | |
| 15 | 96572994 | T | A | 6 | 0 | 0 | 0 | 6 | 0 | NA | NA | 3_prime_UTR_variant | MODIFIER | Slc38a1 | c.*3717A>T | |
| 15 | 96573001 | T | G | 6 | 0 | 0 | 0 | 6 | 0 | NA | NA | 3_prime_UTR_variant | MODIFIER | Slc38a1 | c.*3710A>C | |
| 15 | 96573065 | C | T | 6 | 0 | 0 | 0 | 6 | 0 | NA | NA | 3_prime_UTR_variant | MODIFIER | Slc38a1 | c.*3646G>A | |
| 15 | 96573088 | T | C | 6 | 0 | 0 | 0 | 6 | 0 | NA | NA | 3_prime_UTR_variant | MODIFIER | Slc38a1 | c.*3623A>G | |
| 15 | 96573113 | A | C | 6 | 0 | 0 | 0 | 6 | 0 | NA | NA | 3_prime_UTR_variant | MODIFIER | Slc38a1 | c.*3598T>G | |
| 15 | 96573115 | T | C | 6 | 0 | 0 | 0 | 6 | 0 | NA | NA | 3_prime_UTR_variant | MODIFIER | Slc38a1 | c.*3596A>G | |
| 15 | 96573121 | A | G | 6 | 0 | 0 | 0 | 6 | 0 | NA | NA | 3_prime_UTR_variant | MODIFIER | Slc38a1 | c.*3590T>C | |
| 15 | 96573190 | C | T | 6 | 0 | 0 | 0 | 6 | 0 | NA | NA | 3_prime_UTR_variant | MODIFIER | Slc38a1 | c.*3521G>A | |
| 15 | 96573446 | T | C | 6 | 0 | 0 | 0 | 6 | 0 | NA | NA | 3_prime_UTR_variant | MODIFIER | Slc38a1 | c.*3265A>G | |
| 15 | 96573696 | G | T | 6 | 0 | 0 | 0 | 6 | 0 | NA | NA | 3_prime_UTR_variant | MODIFIER | Slc38a1 | c.*3015C>A | |
| 15 | 96573786 | T | C | 6 | 0 | 0 | 0 | 6 | 0 | NA | NA | 3_prime_UTR_variant | MODIFIER | Slc38a1 | c.*2925A>G | |
| 15 | 96573806 | G | A | 6 | 0 | 0 | 0 | 6 | 0 | NA | NA | 3_prime_UTR_variant | MODIFIER | Slc38a1 | c.*2905C>T | |
| 15 | 96573810 | T | G | 6 | 0 | 0 | 0 | 6 | 0 | NA | NA | 3_prime_UTR_variant | MODIFIER | Slc38a1 | c.*2901A>C | |
| 15 | 96574110 | T | G | 6 | 0 | 0 | 0 | 6 | 0 | NA | NA | 3_prime_UTR_variant | MODIFIER | Slc38a1 | c.*2601A>C | |
| 15 | 96574275 | C | G | 6 | 0 | 0 | 0 | 6 | 0 | NA | NA | 3_prime_UTR_variant | MODIFIER | Slc38a1 | c.*2436G>C | |
| 15 | 96574948 | C | T | 6 | 0 | 0 | 0 | 6 | 0 | NA | NA | 3_prime_UTR_variant | MODIFIER | Slc38a1 | c.*1763G>A | |
| 15 | 96575108 | G | A | 6 | 0 | 0 | 0 | 6 | 0 | NA | NA | 3_prime_UTR_variant | MODIFIER | Slc38a1 | c.*1603C>T | |
| 15 | 96575306 | C | T | 6 | 0 | 0 | 0 | 6 | 0 | NA | NA | 3_prime_UTR_variant | MODIFIER | Slc38a1 | c.*1405G>A | |
| 15 | 96588947 | C | T | 6 | 0 | 0 | 0 | 6 | 0 | NA | NA | synonymous_variant | LOW | Slc38a1 | c.627G>A | p.Leu209Leu |
| 15 | 96592565 | C | G | 6 | 0 | 0 | 0 | 6 | 0 | NA | NA | synonymous_variant | LOW | Slc38a1 | c.330G>C | p.Ser110Ser |
| 15 | 96624089 | G | A | 6 | 0 | 0 | 0 | 6 | 0 | NA | NA | 5_prime_UTR_variant | MODIFIER | Slc38a1 | c.-12C>T | |
| 15 | 96624099 | G | A | 6 | 0 | 0 | 0 | 6 | 0 | NA | NA | 5_prime_UTR_variant | MODIFIER | Slc38a1 | c.-22C>T | |
| 15 | 96642208 | T | C | 6 | 0 | 0 | 0 | 6 | 0 | NA | NA | 5_prime_UTR_variant | MODIFIER | Slc38a1 | c.-173A>G | |
| 15 | 96642564 | G | T | 6 | 0 | 0 | 0 | 6 | 0 | NA | NA | 5_prime_UTR_variant | MODIFIER | Slc38a1 | c.-166C>A | |
| 15 | 96642638 | T | C | 6 | 0 | 0 | 0 | 6 | 0 | NA | NA | 5_prime_UTR_variant | MODIFIER | Slc38a1 | c.-240A>G | |
| 15 | 96642818 | G | C | 6 | 0 | 0 | 0 | 6 | 0 | NA | NA | 5_prime_UTR_variant | MODIFIER | Slc38a1 | c.-420C>G | |
| 15 | 96688724 | G | A | 6 | 0 | 0 | 0 | 6 | 0 | NA | NA | 3_prime_UTR_variant | MODIFIER | Slc38a2 | c.*1403C>T | |
| 15 | 97696518 | C | T | 6 | 0 | 0 | 0 | 6 | 0 | NA | NA | synonymous_variant | LOW | Rpap3 | c.528G>A | p.Ser176Ser |
| 15 | 97750077 | T | C | 6 | 0 | 0 | 0 | 6 | 0 | NA | NA | synonymous_variant | LOW | Rapgef3 | c.2169A>G | p.Thr723Thr |
| 15 | 97791912 | A | T | 6 | 0 | 0 | 0 | 6 | 0 | NA | NA | 3_prime_UTR_variant | MODIFIER | Slc48a1 | c.*1189A>T | |
| 15 | 97792164 | T | C | 6 | 0 | 0 | 0 | 6 | 0 | NA | NA | 3_prime_UTR_variant | MODIFIER | Slc48a1 | c.*1441T>C | |
| 15 | 97800737 | A | G | 6 | 0 | 0 | 0 | 6 | 0 | NA | NA | synonymous_variant | LOW | Hdac7 | c.1938T>C | p.Ser646Ser |
| 15 | 97806529 | G | T | 6 | 0 | 0 | 0 | 6 | 0 | NA | NA | missense_variant | MODERATE | Hdac7 | c.1504C>A | p.Leu502Met |
| 17 | 87797664 | G | A | 6 | 0 | 0 | 0 | 6 | 0 | NA | NA | 5_prime_UTR_variant | MODIFIER | Kcnk12 | c.-210C>T | |
| 17 | 87984510 | C | T | 6 | 0 | 0 | 0 | 6 | 0 | NA | NA | missense_variant | MODERATE | Msh6 | c.692C>T | p.Ala231Val |
| 17 | 87984778 | C | G | 6 | 0 | 0 | 0 | 6 | 0 | NA | NA | synonymous_variant | LOW | Msh6 | c.960C>G | p.Gly320Gly |
| 17 | 87984805 | C | G | 6 | 0 | 0 | 0 | 6 | 0 | NA | NA | synonymous_variant | LOW | Msh6 | c.987C>G | p.Leu329Leu |
| 17 | 87985219 | G | A | 6 | 0 | 0 | 0 | 6 | 0 | NA | NA | synonymous_variant | LOW | Msh6 | c.1401G>A | p.Arg467Arg |
| 17 | 87985261 | A | G | 6 | 0 | 0 | 0 | 6 | 0 | NA | NA | synonymous_variant | LOW | Msh6 | c.1443A>G | p.Arg481Arg |
| 17 | 87985576 | T | A | 6 | 0 | 0 | 0 | 6 | 0 | NA | NA | synonymous_variant | LOW | Msh6 | c.1758T>A | p.Ala586Ala |
| 17 | 87986018 | G | T | 6 | 0 | 0 | 0 | 6 | 0 | NA | NA | missense_variant | MODERATE | Msh6 | c.2200G>T | p.Ala734Ser |
| 17 | 87986242 | C | T | 6 | 0 | 0 | 0 | 6 | 0 | NA | NA | synonymous_variant | LOW | Msh6 | c.2424C>T | p.Leu808Leu |
| 17 | 87986521 | C | T | 6 | 0 | 0 | 0 | 6 | 0 | NA | NA | synonymous_variant | LOW | Msh6 | c.2703C>T | p.Asp901Asp |
| 17 | 87986590 | C | T | 6 | 0 | 0 | 0 | 6 | 0 | NA | NA | synonymous_variant | LOW | Msh6 | c.2772C>T | p.Ile924Ile |
| 17 | 87986686 | C | G | 6 | 0 | 0 | 0 | 6 | 0 | NA | NA | synonymous_variant | LOW | Msh6 | c.2868C>G | p.Arg956Arg |
| 17 | 87990553 | C | T | 6 | 0 | 0 | 0 | 6 | 0 | NA | NA | synonymous_variant | LOW | Msh6 | c.3918C>T | p.Leu1306Leu |
| 17 | 87992100 | G | A | 6 | 0 | 0 | 0 | 6 | 0 | NA | NA | synonymous_variant | LOW | Fbxo11 | c.2739C>T | p.Asp913Asp |
| 17 | 87992109 | G | A | 6 | 0 | 0 | 0 | 6 | 0 | NA | NA | synonymous_variant | LOW | Fbxo11 | c.2730C>T | p.His910His |
| 17 | 87992142 | A | G | 6 | 0 | 0 | 0 | 6 | 0 | NA | NA | synonymous_variant | LOW | Fbxo11 | c.2697T>C | p.Ser899Ser |
| 17 | 87992148 | C | T | 6 | 0 | 0 | 0 | 6 | 0 | NA | NA | synonymous_variant | LOW | Fbxo11 | c.2691G>A | p.Thr897Thr |
| 17 | 87996142 | G | A | 6 | 0 | 0 | 0 | 6 | 0 | NA | NA | synonymous_variant | LOW | Fbxo11 | c.2130C>T | p.Asn710Asn |
| 17 | 88014516 | C | T | 6 | 0 | 0 | 0 | 6 | 0 | NA | NA | synonymous_variant | LOW | Fbxo11 | c.486G>A | p.Leu162Leu |
| 17 | 88065979 | T | C | 6 | 0 | 0 | 0 | 6 | 0 | NA | NA | upstream_gene_variant | MODIFIER | Fbxo11 | c.-727A>G | |
| 19 | 29719566 | C | T | 6 | 0 | 0 | 0 | 6 | 0 | NA | NA | synonymous_variant | LOW | 9930021J03Rik | c.2727G>A | p.Arg909Arg |
| 19 | 42256084 | C | A | 0 | 6 | 0 | 6 | 0 | 0 | NA | NA | 5_prime_UTR_variant | MODIFIER | Golga7b | c.-88C>A | |
| 19 | 42269112 | C | T | 0 | 6 | 0 | 6 | 0 | 0 | NA | NA | 3_prime_UTR_variant | MODIFIER | Golga7b | c.*650C>T | |
| 19 | 42284139 | G | A | 0 | 6 | 0 | 6 | 0 | 0 | NA | NA | synonymous_variant | LOW | Crtac1 | c.1806C>T | p.Asp602Asp |
| 19 | 42298358 | A | G | 0 | 6 | 0 | 6 | 0 | 0 | NA | NA | synonymous_variant | LOW | Crtac1 | c.1245T>C | p.Gly415Gly |
